# Multi-Omics Clustering Differentiates the Total and Intact HIV Reservoirs and Related Host Immune Mechanisms

**DOI:** 10.64898/2026.06.24.734029

**Authors:** Victoria Rios-Vazquez, Mareva Delporte, Twan Otten, Vasiliki Matzaraki, Wilhelm A.J.W. Vos, Marc J.T. Blaauw, Louise E. van Eekeren, Albert L. Groenendijk, Rainer Knoll, Anna C. Aschenbrenner, Rob J.W. Arts, Jéssica C. dos Santos, Adriana Navas, Sarah Gerlo, Jan van Lunzen, Leo A.B. Joosten, Mihai G. Netea, Andre J.A.M. van der Ven, Linos Vanderkerckhove

**Affiliations:** Department of Internal Medicine and Radboud Center of Infectious Diseases, Radboudumc, Radboud University, Nijmegen, The Netherlands; HIV Cure Research Center, Department of Internal Medicine and Pediatrics, Ghent University Hospital, Ghent University, Ghent, Belgium; Department of Laboratory Medicine, Radboudumc, Radboud University, Nijmegen, The Netherlands; Department of Internal Medicine and Infectious Diseases, OLVG, Amsterdam, The Netherlands; Department of Internal Medicine and Infectious Diseases, Elizabeth-Tweesteden Ziekenhuis; Department of Internal Medicine and Department of Medical Microbiology and Infectious Diseases, ErasmusMC, Erasmus University, Rotterdam, The Netherlands; Systems Medicine, German Center for Neurodegenerative Diseases (DZNE), Bonn, Germany; Department of Translational Immunology, Iuliu Hatieganu University of Medicine and Pharmacy, Cluj-Napoca, Romania; Department of Immunology and Metabolism, Life and Medical Sciences Institute, University of Bonn, Bonn, Germany

## Abstract

HIV reservoirs are heterogeneous across individuals, yet host determinants of this variability remain unclear. Applying multi-omics clustering to 1,230 people with HIV, integrating omics and functional data from circulating immune cells (transcriptomics, DNA methylation, immune phenotyping, ex-vivo cytokine production capacity), plasma proteomics, and CD4+ T-cell reservoir measurements (total and intact HIV-DNA copies), revealed three immunologically distinct endotypes: *All Low* (low total/low intact reservoir size), *All High* (high total/high intact reservoir size) and *Mixed* (high total/low intact reservoir size). Per endotype, distinct immune landscapes were noticed in single-layer analyses as well as differences in clinical signatures. Applying non-linear machine learning across all layers, key predictors not captured by linear single-layer approaches showed IFN-γ production and *TCF7/AK5* expression across T-cell and NK-cell populations as well as IL-1β/MCP-1 production after 24h stimulation and *MAN1C1/EDAR* expression on T-cell populations, linked to intact and total reservoir size, respectively. This host-virus integrative multi-omics framework provides a systems-level resource that may help to personalize reservoir-reducing intervention studies aiming for HIV cure and/or comorbidity reductions.

**Graphical Abstract:** **a) Multi-omics clustering** of 1,230 virally suppressed PLHIV identified three immunologically distinct endotypes: All Low (n=428), Mixed (n=483), and All High (n=280), with distinct total and intact HIV reservoir sizes. **b) Single-layer analyses** reveal characteristic immune landscapes: *All Low* Th1-skewed, naïve T cells (immunophenotyping), high IFN-γ production (7-day), downregulated Type I IFN pathways (bulk transcriptome); *Mixed* intermediate Th1/Th2, CD8 and CD4 effector memory T cells (immunophenotyping), IL-5 production (7-day), downregulated Type I IFN pathways (bulk transcriptome); *All High* Th2-skewed, CD4 effector memory T cells (immunophenotyping), high IL-5 production (7-day), upregulated Type I IFN pathways (bulk transcriptome). **c) Endotypes carry distinct comorbidity burdens:** *All Low* lower carotid plaque, AIDS-defining malignancies, and residual viremia; *Mixed* intermediate burden; *All High* highest carotid plaque, AIDS-defining malignancies, opportunistic infections, and residual viremia. **d) Non-linear classification models (XGBoost) prioritize top host predictive markers:** IFN-γ response (7-day), *TCF7* and *AK5* expression in CD4/CD8 T-cells and NK-cells (characterize *All Low*; IL-1β response (24-hour), *MAN1C1* and *EDAR* in CD4/CD8 T-cells, and MCP1 response (24-hour) characterize *All High*. Created in BioRender.

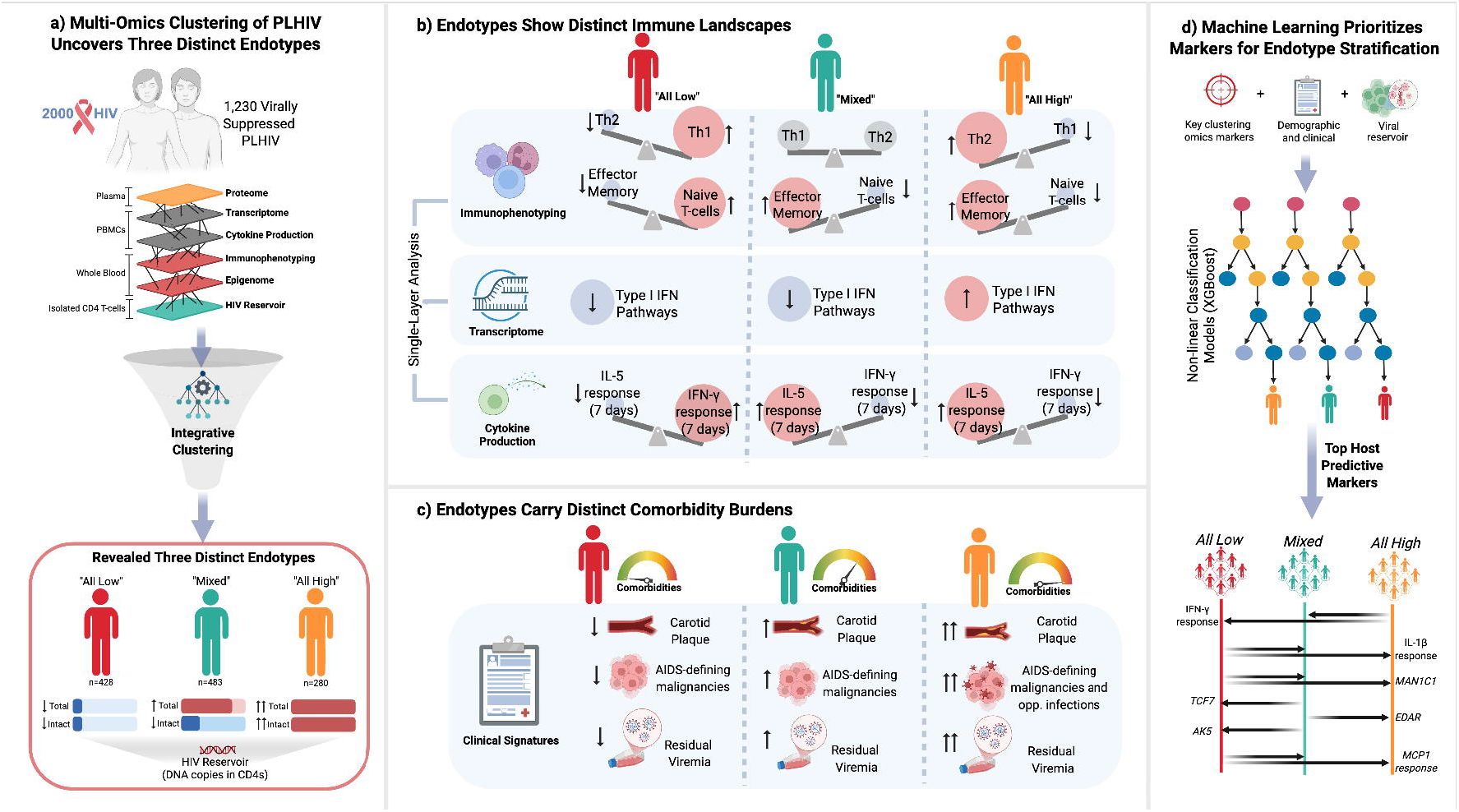

## Main Text

Despite the success of antiretroviral therapy (ART) in suppressing HIV replication, viral persistence in latent reservoirs remains a critical barrier to a functional cure^1–3^. People living with HIV (PLHIV) exhibit marked heterogeneity in clinical outcomes, attributed to factors such as immune status, comorbidities, and demographic characteristics^4,5^. This variability extends to their latent HIV reservoirs, that is primarily composed of proviral DNA integrated into host DNA of memory CD4+ T cells^3,6^. Reservoir metrics such as total and intact HIV DNA levels in circulating CD4 cells provide insights into reservoir size^7,8^. Total HIV DNA levels are associated with faster disease progression^9,10^ and persistent immune activation that impedes reservoir clearance^11^, while intact HIV DNA levels predict viral rebounds after ART interruption in chronically treated PLHIV^12^. The establishment and persistence of HIV reservoirs are influenced by host factors such as immune cell phenotypes, gene expression, and epigenetic modifications^6,13–15^. Recent studies have applied unsupervised clustering approaches to identify subgroups of PLHIV based on reservoir-associated features, highlighting the potential of data-driven stratification to dissect reservoir heterogeneity^16,17^. However, host characterization of these subgroups has largely been limited to immune phenotyping and targeted immunological assays, and it remains unclear whether integrating reservoir burden and composition with in-depth host multi-omics and functional immune profiling reveals biologically and clinically distinct endotypes. Furthermore, whether such endotypes associate with clinically meaningful comorbidity profiles and reproducible across independent clinical centers remains unknown.

Here, we address this gap by applying unsupervised multi-omics clustering, an approach widely used for molecular subtyping in cancer^18,19^, to one of the largest and most deeply characterized cohorts of virally suppressed PLHIV to date (the 2000HIV study^20^, NCT03994835, n = 1,230), comprising discovery (n=1027) and validation (n=203) sub-cohorts recruited from independent clinical centers (**Figure 1**). By integrating data from circulating immune cells (transcriptomics, DNA methylation, immune phenotyping, and *ex vivo* cytokine production), plasma proteomics data, and CD4-associated HIV DNA reservoir quantification, we aimed to (1) identify endotypes of PLHIV with distinct reservoir-host immune profiles using integrative multi-omics clustering and the predictive markers stratifying them, (2) characterize the immunological profiles of these endotypes while accounting for demographic and technical confounders using single-layer analyses and (3) assess their clinical relevance using comprehensive clinical and comorbidity signatures. This integrative framework bridges viral persistence and host immune complexity, providing a systems-level multi-omics resource for the field and a foundation for precision medicine strategies for reservoir-reducing interventions.

**Figure 1.**
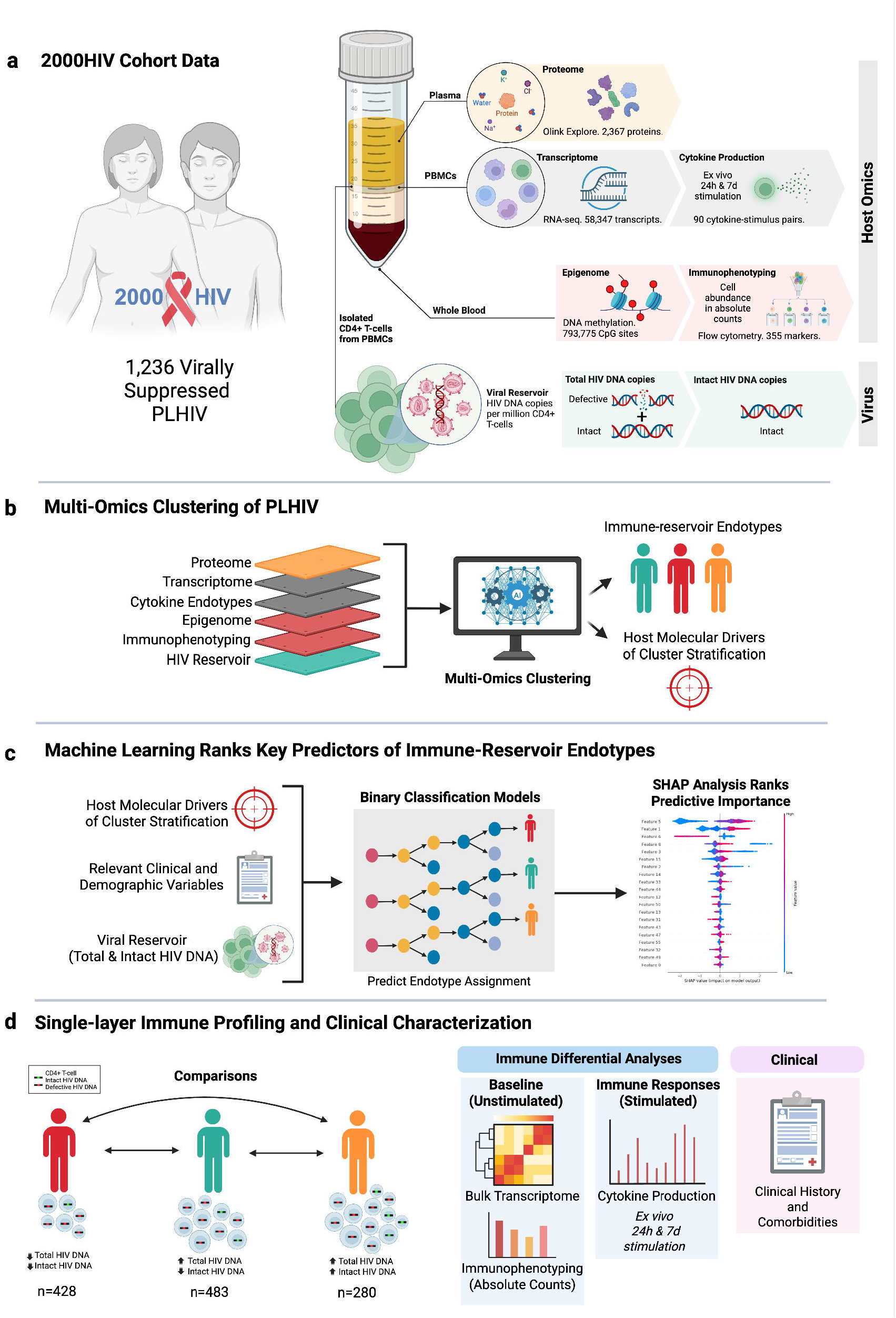
Overview of Multi-Omics Analysis in PLHIV. **a**, Overview of multi-omics and clinical data collected from 1,230 PLHIV in the 2000HIV cohort (NCT03994835) across four Dutch centers, split into discovery (n = 1,027) and validation (n = 203) sub-cohorts. Data included host omics and HIV DNA quantification (total/intact copies per million CD4⁺ T cells), alongside clinical history. Only the top 5,000 most variable transcripts and top 1% most variable CpG sites were included during integration to minimize noise. **b**, Multi-omics clustering of PLHIV using the MOVICS pipeline on preprocessed, high-variability, and log2-transformed features, identifying three endotypes. Endotype assignments were validated across cohorts and corresponded to distinct total and intact HIV DNA profiles. **c**, Prediction modeling and SHAP analysis. XGBoost binary models classified endotypes based on multi-omics, viral reservoir, and clinical features, with SHAP values highlighting top predictive drivers. Schematics were created in BioRender. **d**, Single-layer immune and clinical characterization across endotypes. Baseline and stimulated immune data (flow cytometry, bulk RNA-seq, cytokines) and clinical history (e.g., HIV duration, CD4 nadir) were analyzed.

## Multi-Omics Clustering Identifies Three Immune-Reservoir Endotypes in Virally Suppressed PLHIV

To investigate the heterogeneity of HIV reservoir size and composition while accounting for host immune variation, we applied unsupervised multi-omics clustering within the MOVICS framework^19^ to 1,230 in virally suppressed PLHIV from the 2000HIV cohort^20^ (NCT03994835), split into discovery (disc, n = 1,027) and validation (val, n = 203) sub-cohorts (see **Methods**). We integrated extensive host molecular and immune datasets, including immune phenotyping (whole blood flow cytometry), transcriptomics (peripheral blood mononuclear cells, PBMC), DNA methylation (whole blood), *ex vivo* cytokine production (PBMC) upon diverse bacterial/fungal pathogens, viral antigens, and pattern recognition receptor ligands stimuli, proteomics (plasma), and HIV reservoir parameters (total and intact HIV DNA copies per million circulating CD4⁺ T cells). The reservoir layer was intentionally included to identify host profiles co-varying with reservoir burden and composition without imposing predefined reservoir thresholds. Unsupervised integrative clustering identified three endotypes (k = 3), supported by the Consensus Partition Index and Gap statistics (**Supplementary Figure 1**), and consistent across discovery and validation sub-cohorts (see **Methods, Supplementary Fig. 2-6**).

The identified endotypes: *All Low*, *Mixed*, and *All High*, displayed distinct reservoir and host molecular/immune profiles (**Figure 2**). The *All Low* endotype (n_disc._ = 345, n_val._ = 83) exhibited the lowest total (median, *m_disc._* = 165, *m_val._* = 283) and intact HIV DNA (*m_disc._* = 4, *m_val._* = 7) levels (copies per million CD4⁺ T cells), whereas the *All High* endotype (n_disc._ = 241, n_val._ = 39) showed the highest total (*m_disc._*= 1,492, *m_val._*= 1,677) and intact (*m_disc._* = 144, *m_val._* = 168) proviral DNA levels. The *Mixed* endotype (n_disc._ = 416, n_val._ = 67) was intermediate, characterized by high total (*m_disc._* = 840, *m_val._* = 1,039), but low intact HIV DNA (*m_disc._* = 20, *m_val._* = 26) levels, revealing a potential decoupling between the quantity and quality of the reservoir.

Although total and intact HIV DNA levels were the primary drivers of endotype differentiation, the resulting endotypes were also defined by joint biological profiles across the integrated molecular and immune layers. Notably, the *Mixed* endotype exhibited partial decoupling of total and intact HIV DNA levels, challenging the assumption that total reservoir size closely reflects replication-competent virus levels^21^. While *All Low* and *All High* endotypes align with this assumption^21,22^, the newly-characterized *Mixed* endotype reveals distinct multi-omics signatures, highlighting the potential for cure strategies tailored to distinct host molecular and reservoir profiles.

## Host Molecular Drivers of Endotype Stratification

To identify the host molecular and immune factors driving endotype stratification, we examined features independently selected by the *MoCluster* integrative algorithm^22^ (**Figure 2**). Features consistently identified across both sub-cohorts were visualized and correlated with total and intact HIV DNA (see **Methods**).

**Figure 2.**
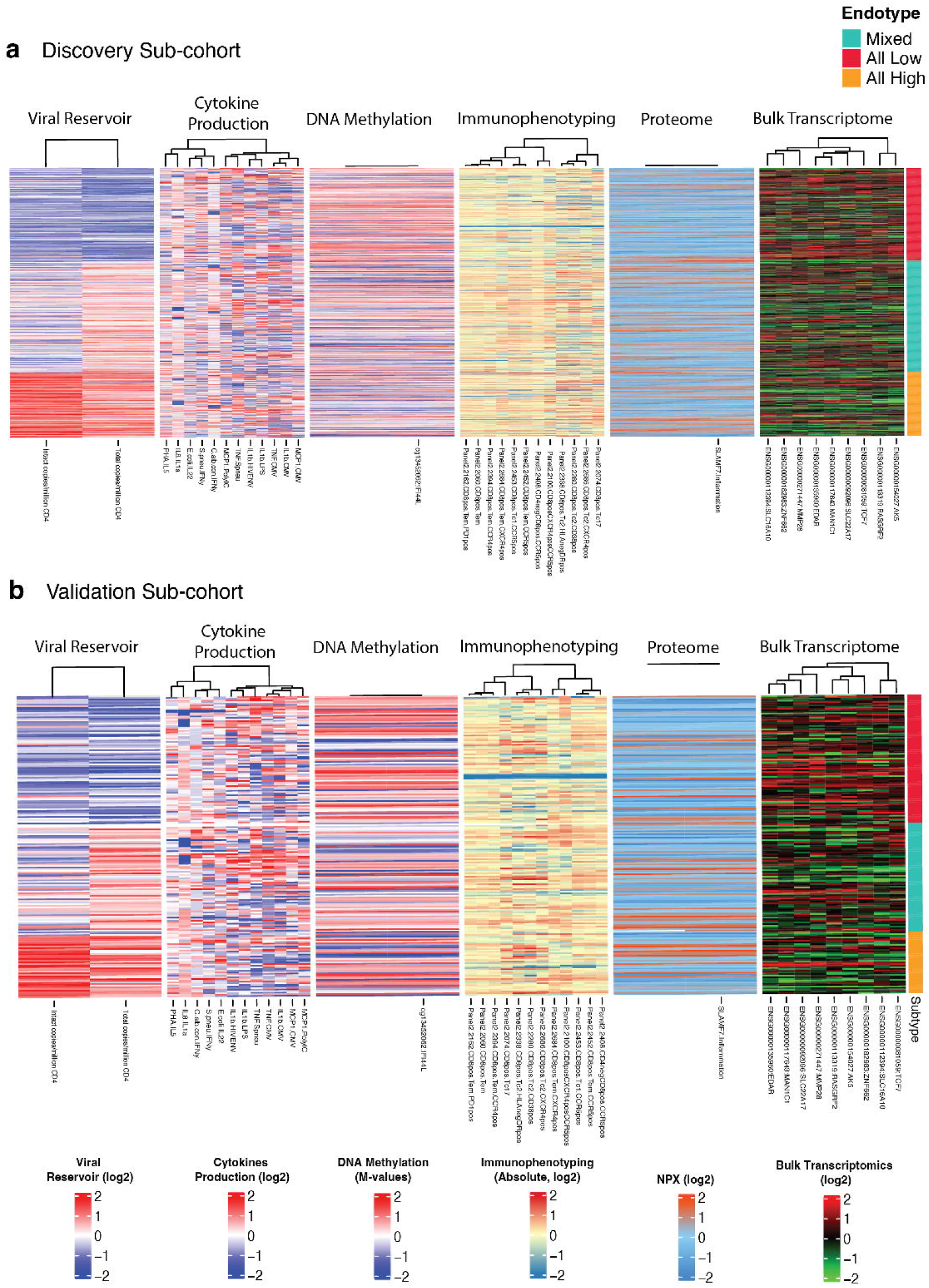
Heatmap of Multi-Omics Clustering Results in the 2000HIV Cohort. **a**, Heatmap of MoCluster multi-omics clustering for the discovery sub-cohort (n = 1,002), revealing three endotypes: All Low (Low Total, Low Intact HIV DNA), Mixed (High Total, Low Intact HIV DNA), and All High (High Total, High Intact HIV DNA), driven significantly by viral reservoir measurements (total and intact HIV DNA copies per million CD4+ T cells). **b**, Validation cohort heatmap (n = 189), confirming the same three endotypes (All Low, Mixed, All High) as in the discovery cohort, confirming the robustness of the stratification.

Beyond reservoir parameters, 43 host features stratified the three endotypes (**Supplementary Figs. 7-14**). In bulk PBMCs, nine transcripts were identified. Complementary scRNA-seq provided cellular context for these markers (**Supplementary Figs. 8 and 9**): *TCF7* and *AK5* showed the broadest lymphocyte-associated expression across naïve and memory CD4^+^/CD8^+^ T cells and NKCD56^bright/dim^ subsets; *RASGRF2, SLC16A10, SLC22A17, EDAR, MAN1C1*, and *MMP28* were also prominent across T-cells, especially across naïve and activated CD4^+^/CD8^+^ T-cell populations; and ZNF662 was most prominent in dendritic cells. Immune phenotyping revealed twelve CD8⁺ T-cell subsets positive for activation and exhaustion markers. *Ex vivo* production identified twelve cytokine-stimulus pairs reflecting both pro-inflammatory and regulatory responses. Systemic layers showed more limited contributions, with plasma proteome revealing only SLAMF7 and whole-blood DNA methylation showing a single CpG site (cg13452062) located in the 51UTR (gene body) of *IFI44L* within an open-chromatin region. Bulk RNA-seq confirmed reduced *IFI44L* expression in individuals with cg13452062 hypermethylation, and scRNA-seq localized *IFI44L* transcription to classical antiviral monocytes (**Supplementary Fig. 13**). Effect-size estimates (standardized mean differences and Cohen’s1d) across endotypes for all markers are provided in **Supplementary Table 1**.

Correlation analyses revealed cross-omics relationships with the HIV reservoir (P<0.05, **Supplementary Table 2**). In bulk PBMCs, all nine transcripts were inversely correlated with total HIV DNA across both sub-cohorts (ρ ≈ -0.06 to -0.24). Among immune phenotyping features, CD8⁺ Tem (PD-1⁺ and CCR4⁺) subsets showed consistent positive associations with total HIV DNA (ρ ≈ +0.16 to +0.23), but not with intact HIV DNA. For cytokine production, only IFN-γ production induced by *S. pneumoniae* (after 7-days) showed a consistent inverse association with intact HIV DNA in *All Low*, reaching significance in the validation sub-cohort (ρ ≈ -0.26). In DNA methylation, hypermethylation of cg13452062 was negatively associated with both total and intact HIV DNA (ρ ≈ -0.17 to -0.33), whereas in plasma proteome SLAMF7 showed weak, non-significant positive correlations.

Together, these findings define a focused set of host immune features that stratify the three endotypes and associate with reservoir size and integrity. To formally quantify the relative importance of each feature and evaluate their predictive power beyond unsupervised clustering, we next applied supervised machine-learning approaches.

## Machine Learning Ranks Key Predictors of Immune-Reservoir Endotypes

Building on the host stratifying markers defined by *MoCluster* integrative clustering algorithm, we trained supervised classification models (XGBoost) using these validated immune features, HIV reservoir parameters (total and intact HIV DNA), and relevant clinical covariates to quantify their relative predictive importance in distinguishing the three immune-reservoir endotypes (**Methods, Supplementary Fig. 15**). Model performance was evaluated in held-out test sets comprising 20% of the data and showed modest-to-moderate discrimination (test-set AUCs: ∼0.65). These models were therefore used primarily for feature prioritization rather than as stand-alone clinical classifiers.

Model interpretability was achieved using SHapley Additive exPlanations (SHAP) analysis^23^, which quantifies how much each variable contributes to the model’s prediction for an individual. The magnitude of the SHAP value reflects the strength of that effect: larger values correspond to greater shifts in predicted probability.

### 1) Predictors comparing All high and All Low endotypes

In *All Low vs All High* endotypes, positive SHAP values indicates higher levels in *All High,* negative SHAP values higher in *All Low*. The model revealed intact HIV DNA (SHAP: +0.46) and total HIV DNA (+0.43) levels as leading predictors, followed by IFN-γ (*S. pneumoniae* stimulation, -0.16), Tenofovir Alafenamide (TAF) exposure (−0.13), Ritonavir (RTV) exposure (+0.12), CD4 count (−0.11), IL-1β (CMV stimulation, +0.09), CD8⁺CXCR4⁺CCR5⁺ T-cells (−0.07), MAN1C1 (+0.06), and MCP-1 (Poly I:C stimulation, -0.06) (**Figure 3a**). Mechanistically, MCP-1 and IFN-γ have been linked to recruitment and activation of antiviral effector cells^24,25^, likely contributing to clearance of infected cells, while heightened IL-1β production to CMV in *All High* is associated with inflammatory signatures consistent with via chronic inflammation CMV-driven immune activation^26,27^. *MAN1C1*, involved in N-glycan processing, may contribute to HIV persistence in *All High* by promoting efficient Env glycan maturation, reducing exposure of conserved high-mannose epitopes targeted by broadly neutralizing antibodies^28–31^. Higher CD4 counts and CD8⁺CXCR4⁺CCR5⁺ T cells in *All Low* align with observations that higher percentages of CD4 cells associate with lower HIV reservoir^16^ while CD8⁺CXCR4⁺CCR5⁺ T cells represent cytotoxic subsets that traffic to lymphoid sanctuaries and support the elimination of infected targets^32,33^. Differences in TAF and RTV exposure likely reflect differences in start of treatment and cART duration.

**Figure 3.**
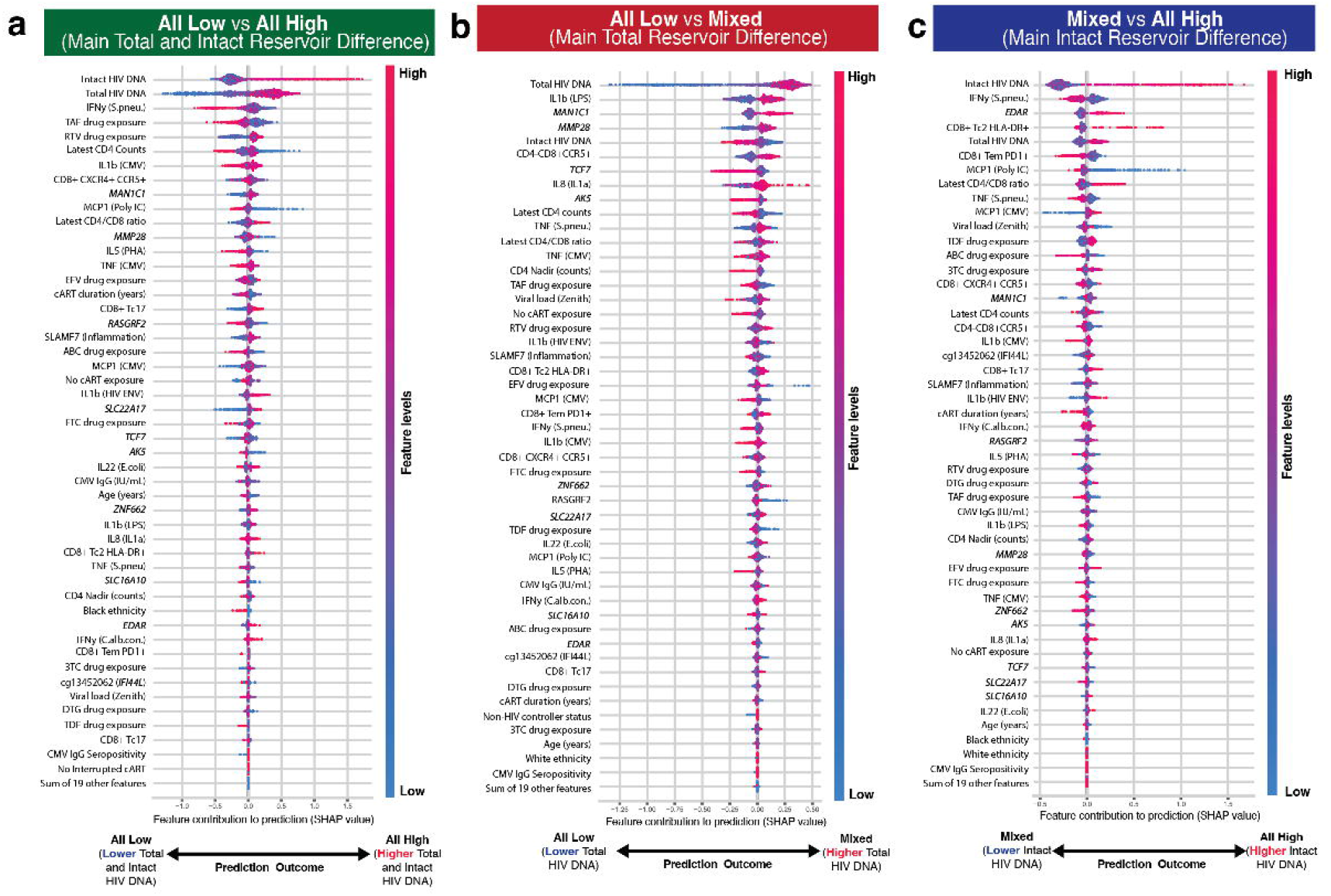
SHAP analysis identifies key predictors of HIV reservoir endotypes. SHAP (SHapley Additive exPlanations) values from XGBoost models quantify the contribution of each feature to predicting the three endotypes of PLHIV. Bee swarm plots show individual participants (color-coded by feature value: blue = low, red = high), with features ordered from most (top) to least predictive (bottom). (a) Features distinguishing *All Low* vs *All High* endotypes, with positive SHAP values supporting *All High* and negative values supporting *All Low*. (b) Features distinguishing *All Low* vs *Mixed endotypes*, with total HIV DNA levels as top predictor. (c) Features separating Mixed vs *All High* endotypes, with intact HIV DNA levels as top predictor. Italicized labels indicate gene symbols; cytokines are represented as cytokine (stimulus). Cytokines measured after 24-hour stimulation were IL-1β, IL-1Ra, IL-6, IL-8, IL-10 (LPS), MCP-1, MIP-1α and TNF, whereas IFN-γ, IL-5, IL-10 (PHA), IL-17 and IL-22 were measured after 7-day stimulation. Antiretroviral cumulative exposure is expressed in years and includes: TAF (tenofovir alafenamide), TDF (tenofovir disoproxil fumarate), EFV (efavirenz), RTV (ritonavir), ABC (abacavir), FTC (emtricitabine), 3TC (lamivudine), and DTG (dolutegravir). See Methods for details.

### 2) Predictors comparing *All Low* and *Mixed* endotypes

In *All Low* versus *Mixed* endotypes, positive SHAP values indicate high levels in *Mixed*. The top-ranked predictor is total HIV DNA (SHAP = +0.38) confirming that these endotypes primarily differ by reservoir size, followed by IL-1β production (LPS stimulation, +0.11), MMP28 (+0.08), TCF7 (−0.08), intact HIV DNA (−0.08), *MAN1C1* (+0.08), CD4⁻CD8⁺CCR5⁺ T-cells (+0.07), IL-8 (IL-1α stimulation, +0.07), AK5 (−0.05), and CD4 count (−0.05) (**Figure 3b**). Features enriched in *Mixed* partially overlapped with those in *All High* such as total HIV DNA, IL-1β, and *MAN1C1.* Mechanistically, heightened IL-1β and IL-8 production suggest a more inflammatory response profile, consistent with immune activation states associated with HIV reservoir maintenance^34,35^. Conversely, higher Transcription factor 7 (*TCF7)* expression in low-reservoir states aligns with its known role in maintaining TCF-1⁺ naïve and less differentiated CD4 and CD8⁺ T cells that sustain antiviral proliferation and limit proviral integration^36–38^. The role of Matrix Metallopeptidase 28 (MMP28), involved in breakdown extracellular matrix, and Adenylate cyclase 5 (AK5), a nucleoside monophosphate kinase, has not been specifically studied in HIV. Notably, these analyses can reveal trends that differ from those observed when inspecting individual markers in isolation (e.g., *MAN1C1*, *MMP28*). This arises because the SHAP-based multi-omic models incorporate interactions among features (**Supplementary Fig. 16**), allowing association patterns to shift once the broader molecular context is considered, an effect known as Simpson’s paradox^39^. Rather than indicating contradiction, this highlights how integrated modeling can capture system-level relationships that are not apparent in isolation.

### 3) Predictors comparing Mixed and All High endotypes

When comparing the *Mixed* vs *All High* endotypes, positive SHAP values indicating higher levels in *All High.* Top predictor markers is intact HIV DNA (SHAP: +0.48), confirming these endotypes primarily reflect differences in reservoir intactness, followed by IFN-γ (*S. pneumoniae* stimulation, -0.09), EDAR (+0.07), CD8+ Tc2 HLA-DR+ (+0.06), total HIV DNA (+0.08), CD8+ Tem PD1+ (−0.08), MCP-1 (Poly I:C stimulation, -0.09), CD4/CD8 ratio (+0.07), TNF (*S. pneumoniae* stimulation, -0.04), and MCP-1 (CMV stimulation, +0.05) (**Figure 3c**). *Mixed* overlapped with *All Low* with high IFN-γ (S. pneumoniae) and MCP-1 (Poly I:C) production. Remarkably, high TNF production (S. pneumoniae) is seen in *Mixed*, however this pro-inflammatory cytokine can also show anti-inflammatory functions by activating regulatory T cells (Tregs)^40,41^. Heightened Ectodysplasin A Receptor (*EDAR)* expression in *All High* may disrupt lymphoid homeostasis via dysregulated NF-κB signaling^42^. Expanded CD8⁺ Tc2 HLA-DR⁺ subsets, linked to Th2-skewed CD8 dysfunction in progressive HIV, correlate with larger reservoirs through impaired cytotoxic clearance^43^. Lower CD4/CD8 ratios in *All High* reflect disease progression^44^.

Altogether, these markers indicate that stronger IFN-γ and MCP-1 production, reflective of innate and adaptive immune system activity^45,46^, favor low intact HIV DNA levels, while T cell differentiation, as shown by *TCF7* expression, seems important for total HIV DNA. The total HIV DNA levels are linked to stronger inflammatory production, as shown by IL-1b levels, but also glycosylation and NF-KB signaling seem relevant, because of MAN1C1 and EDAR. Different activated cytotoxic T cells may play different roles among the endotypes. The *Mixed* endotype integrates features of both extremes, reinforcing a continuum model of immune-reservoir heterogeneity.

The markers presented above, are the drivers of endotype differentiation in the multi-omics clustering analysis. A more detailed profile of the three endotypes can be obtained by single-layer analysis.

## Single-layer Immune Profiling Reveals Immune Gradient Underlying HIV Reservoir Heterogeneity

Single-layer analyses were performed using data from phenotyping (absolute counts and percentages) circulating immune cells^47^, PBMC transcriptomics and cytokine production after ex-vivo PBMC stimulation^20^, capturing baseline immune composition and functional responses across endotypes, while adjusting for demographic, clinical, and technical confounders (**Methods**). Pairwise comparisons captured distinct contributions of total and intact HIV DNA levels: *All High* versus *All Low* reflected the combined extremes (log₂FC Total = 3.18 disc., 2.57 val.; log₂FC Intact = 5.17 disc., 4.58 val.). *All Low* versus *Mixed* showed moderately higher differences in total HIV DNA (log₂FC Total = 2.35 disc., 1.87 val.) with similar shifts in intact HIV DNA (log₂FC Intact = 2.32 disc., 1.89 val.). *Mixed* versus *All High* primarily reflected intact HIV DNA effects (log₂FC Total = 0.83 disc., 0.69 val.; log₂FC Intact = 2.32 disc., 1.89 val.). When plotted with total and intact HIV DNA as orthogonal axes, these patterns become visually apparent: *All Low* and *Mixed* overlap along the intact HIV DNA axis, whereas *Mixed* and *All High* overlap along the total HIV DNA axis (**Supplementary Figure 5**).

### 1) Detailed characteristics of All High compared to All Low

Unstimulated circulating immune cells of participants in the *All High* endotype exhibited a more differentiated, higher Tfh2 CD4+ proportions, and type I interferon-enriched profile relative to *All Low*. Absolute counts revealed enrichment of naïve B cells in *All High* (normalized estimates = 0.2-0.5, FDR_discovery_ < 0.05, P_validation_ < 0.05), consistent with impaired B-cell maturation linked to persistent type I interferon signaling and disrupted Tfh-B-cell interactions in germinal centers^48^. Proportion of effector-memory (CCR4^+^) and Tfh2 CD4⁺ subsets were also higher in this endotype (normalized estimates of 0.5-0.6 and 0.2-0.5, respectively; **Figure 4a**, **Supplementary Table 3a**). In contrast, *All Low* showed enriched absolute counts of CD4⁺ Th1 population (normalized estimates = 0.4-0.5; FDR < 0.05, P < 0.05), while elevated proportions of CD4^+^ Tfh1 confirmed a Th1/Tfh1-skewed profile (normalized estimates = 0.27-0.43).

**Figure 4.**
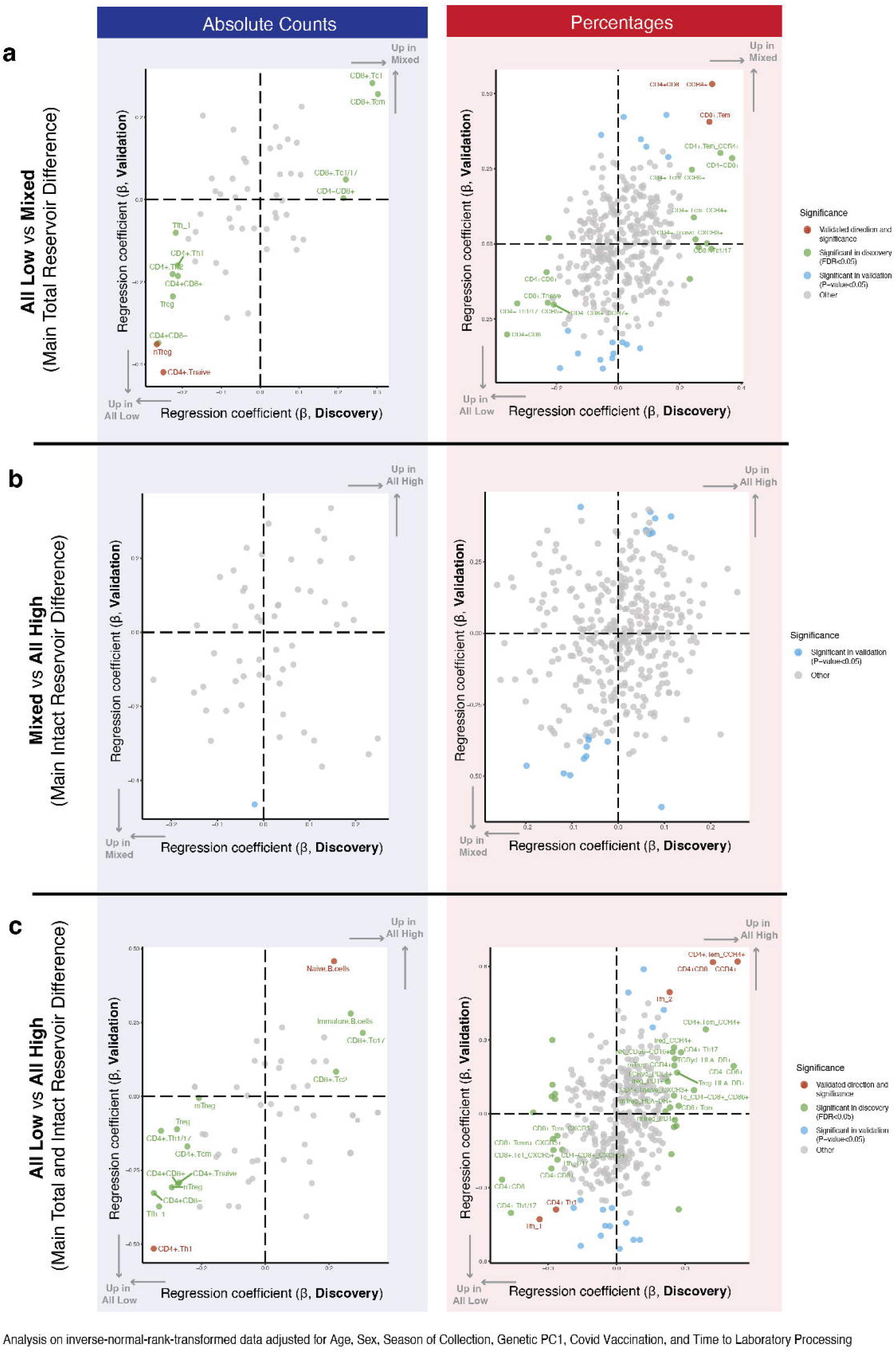
Consistency of Immune Cell Populations in Discovery and Validation Sub-Cohorts. **a-c**, Scatter plots comparing effect estimates for flow cytometry markers between the discovery (x-axis) and validation (y-axis) cohorts. Left panels show absolute cell counts; right panels show percentages of each subset. Analyses were adjusted for age, sex, season, first genetic principal component, COVID-19 vaccination status, and time to laboratory processing (see **Methods**). Red points highlight markers where both the direction of effect (same sign) and statistical significance (FDR < 0.05 in discovery, P < 0.05 in validation) were replicated, emphasizing robust, reproducible immune associations across cohorts.

At the transcriptional level, five genes were differentially expressed (P < 0.05 across sub-cohorts). Complementary scRNA-seq provided cellular context for bulk RNA-seq findings across comparisons (**Supplementary Figs. 17 and 18**). *NAB1,* detected across dendritic cells, monocytes and selected T-cell populations, is a corepressor of EGR-1 that modulates immune activation and inflammation^49^ and was modestly upregulated in *All High* (Log₂FC = 0.07), while *AL096816.1*, *CCT4*, *ANKRD18EP*, and *NAP1L3* were downregulated (Log₂FC = -0.03 to -0.25; **Figure 5a**, **Supplementary Table 4a**). Downregulation of *CCT4,* prominent in proliferating T-cell populations, and *NAP1L3,* expressed across CD4⁺ and CD8⁺ T-cell subsets, key mediators of protein folding and nucleosome assembly, points to disrupted cellular homeostasis and chromatin organization^50,51^, whereas *ANKRD18EP,* detected predominantly in plasmablasts and proliferating T-cell populations, have been implicated in immune signaling^52^. Gene-set enrichment analysis (GSEA) identified type I interferon IFN-α/β signaling as the top upregulated pathway enriched in *All High* (NES = 2.1, adj. P < 0.05; **Figure 5a, Supplementary Table 5a**), with contributing genes showing strongest expression across monocyte and dendritic-cell populations, alongside broader expression in selected lymphoid subsets (**Supplementary Figure 18a**), and peptide chain elongation as the top pathway enriched in *All Low* (NES = -3.3; adj. P < 0.05), with contributing genes broadly expressed across immune-cell populations (**Supplementary Figure 18b**).

**Figure 5.**
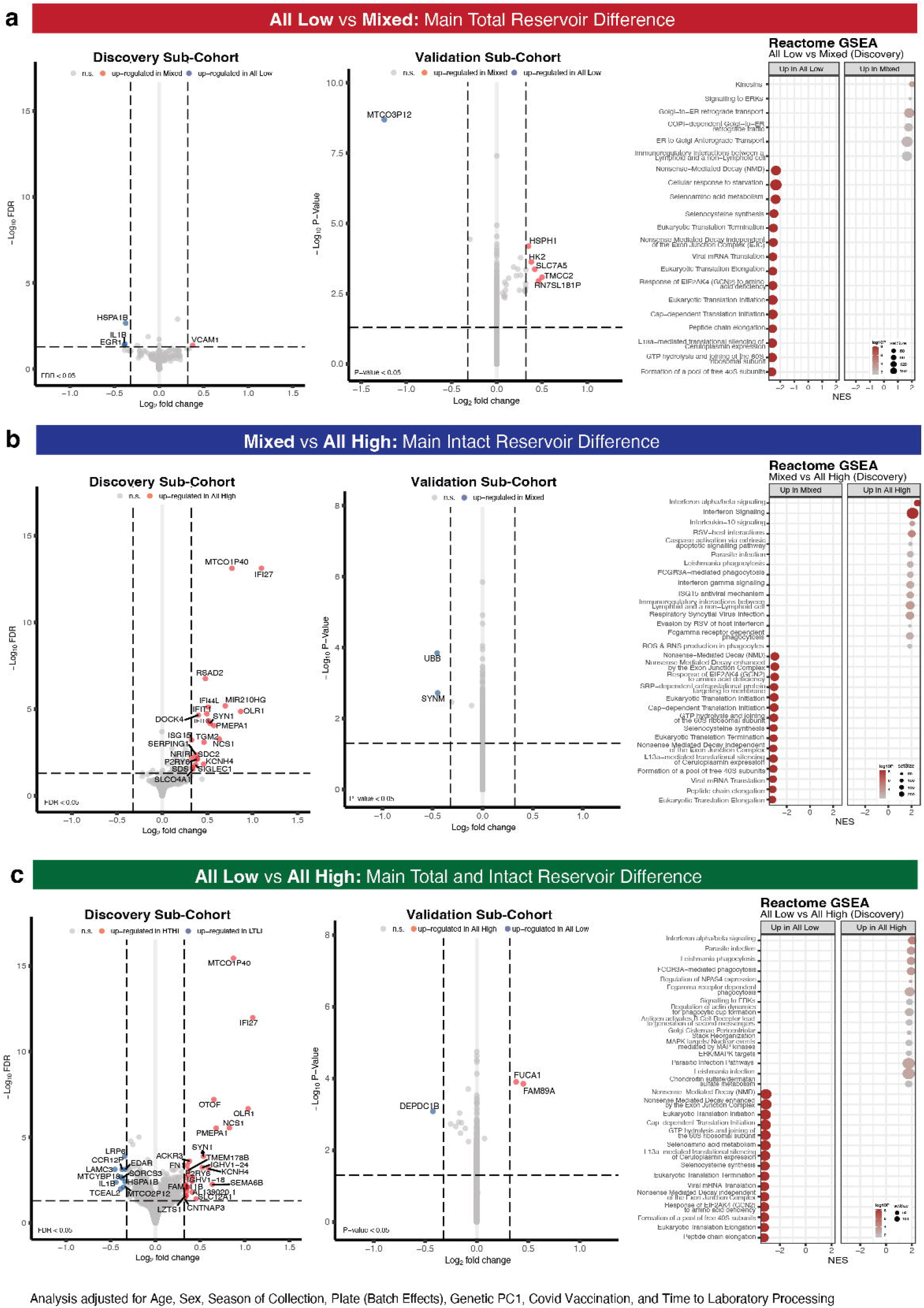
Consistency of Unstimulated Transcriptome Signatures in Discovery and Validation Sub-Cohorts. Panels a-c show differential expression and pathway enrichment PBMCs bulk transcriptomics across endotypes. In each panel, the left plot shows the discovery cohort volcano plot, highlighting genes significant after BH correction (FDR < 0.05) and exceeding a log₂ fold-change of 1.25. The middle plot shows the validation cohort volcano plot, labelling genes that surpass the same fold-change threshold and reach nominal significance (P < 0.05). The right plot shows Reactome gene set enrichment analysis (GSEA) in the discovery cohort of genes ranked by -log₁₀(nominal P-value) × sign of fold change and displaying the top significantly enriched pathways (up- or down-regulated) after BH correction. All analyses were adjusted for age, sex, season, first genetic principal component, plate (batch effect), COVID-19 vaccination status, and sample processing time (see **Methods**).

Ex vivo cytokine production suggested opposing Th1/Th2-associated functional profiles between extreme endotypes. Because cytokine responses were measured after both 24-hour and 7-day stimulation with a validated panel of microbial, viral and pattern-recognition receptor stimuli^20,53^, stimulus-specific cytokine patterns provide insight into predominantly monocyte-associated short-term responses and longer-term T-helper-associated cytokine-production capacity in mixed PBMC cultures across endotypes (see **Methods**). *All High showed* elevated IL-5 production (PHA, broad, polyclonal T-cell receptor stimulus) following 7-day stimulation (P < 0.05 in discovery, normalized estimates=0.06-0.10, validated estimate direction; **Supplementary Table 6, Figure 6a; Supplementary Fig. 19a**). This finding is consistent with a more Th2-associated functional profile^54^ in *All High*. *All Low* showed higher IFN-γ (*S. pneumoniae, E. coli, M. tuberculosis* stimulation; normalized estimates=0.01-0.07), IL-22 (*E. coli*; normalized estimates = 0.01-0.06), IL-10 (PHA; normalized estimates = 0.01-0.13), and IL-17 (PHA, S. pneumoniae; normalized estimates = 0.01-0.11) production following 7-day stimulation, suggesting greater Th1/Th17-associated cytokine-production capacity in profiles characterized by lower intact proviral burden^25,55^, while IL-22 reinforce mucosal defense and antiviral activity^56,57^ and IL-10 may contribute to regulation of excessive inflammation^58,59^. Among the short-term (24-hour) stimulations revealed only MIP-1α production (S. pneumoniae) enriched in *All Low* (normalized estimates = 0.11-0.18; **Supplementary Fig. 20a**), a predominantly monocyte-associated inflammatory response, consistent with its role in recruiting effector cells linked to HIV-1 inhibition^60,61^.

**Figure 6.**
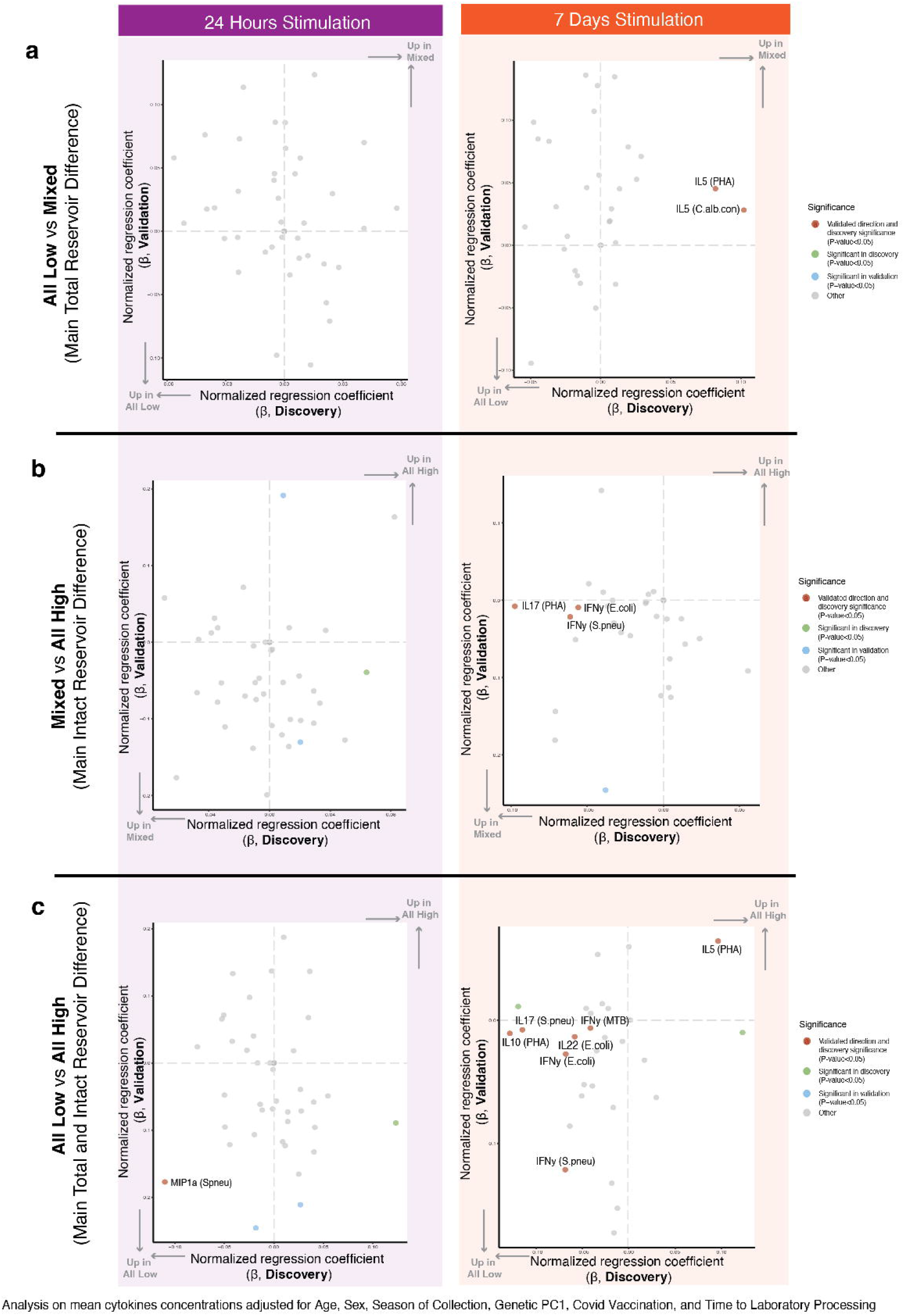
Consistency of Stimulated Ex Vivo Cytokine Production in Discovery and Validation Sub-Cohorts. **a-c**, four-quadrant scatter plots comparing effect estimates from ex vivo cytokine production capacity upon stimulation in the discovery cohort (x-axis) versus the validation cohort (y-axis) across distinct cytokine endotypes. In each panel, the left plot shows results after 24-hour stimulation, while the right plot shows results after 7-day stimulation. Markers highlighted in red indicate consistent direction (same sign of normalized effect estimates) and statistical significance (P < 0.05 in both cohorts). All analyses were adjusted for age, sex, season of sample collection, first genetic principal component, COVID-19 vaccination status, and time to laboratory processing (see **Methods**).

Together, these findings define *All High* as a more differentiated, type I interferon-enriched, and Tfh2-biased immune profile associated with greater reservoir burden, and *All Low* as a less differentiated, Th1/Tfh1-skewed endotype maintaining translational and protein homeostasis programs, consistent with IFN-γ-associated antiviral immunity and lower reservoir burden in this group^62–64^.

### 2) Detailed characteristics of Mixed vs All Low

Comparing the *Mixed* vs All Low endotypes exhibited an intermediate immune landscape. *Mixed* showed lower absolute counts of CD4⁺ naïve and natural regulatory T cells (normalized estimates = 0.3-0.4; FDR_discovery_ < 0.05, P_validation_ < 0.05), populations associated with flexible, less differentiated states limiting proviral inducibility^65–67^. In contrast, proportions of effector-memory CD8⁺ and CCR4⁺ CD4⁺ subsets were relatively higher (normalized estimates ≈ 0.3-0.4) in *Mixed*, mirroring the differentiated endotype seen in *All High* (**Figure 4b, Supplementary Table 3b**). Transcriptomics analyses identified *ASB2* mainly expressed in dendritic cells (**Supplementary Figure 17f**), involved in cytoskeletal remodeling and dendritic-cell antigen cross-presentation^68^, as the only significantly upregulated in *Mixed* versus *All Low* (Log₂FC = 0.1-0.3; P<0.05; **Fig. 5b**, **Supplementary Table 4b**). GSEA highlighted kinesin-related pathways (Normalized Enrichment Scores, NES = 2.0; adj. P < 0.05) as enriched in *Mixed,* with its contributing genes distributed mainly across monocytes and dendritic cells (**Supplementary Figure 18c**), whereas translational-control pathways (NES ≈ -2.6) remained dominant in *All Low* (**Fig. 5b**, **Supplementary Table 5b**) and were broadly expressed across immune-cell subsets (**Supplementary Figure 18b**).

Exploring functional capacities of *Mixed and All Low,* little functional differences were noticed, apart from IL-5 production following PHA and *C. albicans* stimulation (normalized estimates = 0.03-0.08, 7 days; **Figure 6b, Supplementary Fig. 19b**) in *Mixed*, mirroring the observed Th2 skew in this endotype.

### 3) Detailed characteristics of Mixed vs All High

Comparing *Mixed* and *All High* endotypes showed broadly similar immune compositions and transcriptomic profiles with no validated differences (Fig. 4c and 5c, **Supplementary Table 3c and 4c**). Nonetheless, subtle transcriptional programs emerged from pathway analysis in the discovery sub-cohort: *All High* was enriched for type I interferon IFN-α/β signaling (NES = 2.6), IFN signaling (NES = 2.1), and IL-10 signaling (NES = 2.1), with genes contributing to these pathways most prominently expressed across monocyte and dendritic-cell populations (**Supplementary Figure 18a**), whereas *Mixed* preserved enrichment of translational control pathways with eukaryotic translation elongation (NES = -3.26) as top pathway (**Figure 5c, Supplementary Table 5c**) with its contributing genes broadly expressed across PBMC populations (**Supplementary Figure 18b**). Thus, while *Mixed* and *All High* share overall similar immune composition, their transcriptional pathways offer more details, with *Mixed* preserving the translational control observed in *All Low*, while *All High* exhibits stronger type I interferon and inflammatory signaling.

*Mixed* compared to *All High*, exhibited higher IFN-γ (*S. pneumoniae, E. coli*) and IL-17 (PHA) production after 7-day stimulation (normalized estimates = 0.01-0.09; **Figure 6c, Supplementary Fig. 19c**), indicating enriched Th1/Th17-type antiviral activity^69^ resembling that of *All Low.* Thus, *Mixed* integrates Th2-associated and Th1/Th17-associated functional traits, reflecting its immunologic position between the two extremes.

## Distinct Clinical Signatures Associated with the Immune-Reservoir Endotypes

To explore if the three endotypes exhibit distinct clinical or comorbidity signatures, we first compared demographic and HIV-related history from clinical records at inclusion separately across both sub-cohorts (**Methods** and **Supplementary Tables 7-9**). Next, to ensure adequate event counts and stable effect estimates for less frequent outcomes, we evaluated endotype-comorbidity associations using logistic regression in the pooled cohort.

*Demographic and clinical differences.* Individuals in the *All High* endotype were ∼4 years older than those in *All Low* (mean 52 vs. 48 years in discovery, 56 vs. 52 in validation, P=0.001 and 0.18) and displayed higher CMV IgG antibodies (median Δ ≈ +15 and +145 IU/mL; P=0.002 and 0.97). They also had lower CD4 nadirs (median Δ ≈ -0.13 and -0.18×10⁹ cells/L; P<0.001 and 0.018), higher viral load zeniths (mean Δ ≈ +1 and +2 log₂ copies/mL; P<0.001 and 0.017), and lower latest CD4 counts (P<0.001 in discovery). In contrast, *All Low* individuals showed the most favorable profile, with younger age, higher CD4 nadirs and CD4 counts. The *Mixed* endotype most closely resembled *All High* in age, but exhibited longer HIV infection and treatment durations relative to both *All Low* (Δ=+1.6 and +2.5 years, P<0.001 and P=0.35) and *All High* (Δ=+1.3 and +3.17 years, P=0.040 and P=0.07). The *Mixed* group also had the lowest prevalence of early cART initiation in discovery (Δ=-7.2%, P=0.002 vs. *All Low*; Δ=-0.6%, P=0.913 vs. *All High*), though this was not confirmed in validation. No significant differences were observed across endotypes for sex, ethnicity, collection season, COVID-19 history, hepatitis co-infection, or treatment history across endotypes.

*Comorbidity associations.* To assess whether endotype membership carried distinct comorbidity profiles, we fitted logistic-regression models between endotypes and comorbidities across the full cohort adjusting for relevant confounders, only significant results are reported (see **Methods, Supplementary Table 10, Supplementary Figure 21**). Relative to *All Low*, individuals in *All High* showed higher odds of carotid plaque (adjusted OR=1.51 *P*=0.022), residual viremia at baseline and follow-up (adjusted OR=3.27-3.47, P<0.0001), musculoskeletal disorders composed of osteoporosis, osteoarthritis and inflammatory arthritis (adjusted OR=1.69 *P=*0.035), and AIDS-defining malignancies, opportunistic infections, and other events such as HIV encephalopathy and wasting syndrome (adjusted OR=2.04-2.71 *P*<0.05). Relative to *All Low*, the *Mixed* endotype displayed higher odds of carotid plaque (adjusted OR=1.41 *P*=0.024), residual viremia (adjusted OR=2.65-2.66, P<0.0001), AIDS-defining malignancies (adjusted OR=2.11 *P*=0.010), post-cART opportunistic infections (adjusted OR=1.61 *P*=0.037), and gastrointestinal disease composed of liver steatosis, fibrosis, cirrhosis, gastro-esophageal reflux disease, or irritable bowel disease (adjusted OR=1.40 *P*=0.042). *All High*, relative to *Mixed,* showed higher odds of AIDS-defining opportunistic infections and other AIDS-defining events (adjusted OR=1.93-1.97 *P*<0.007). These patterns align with another report from the 2000HIV cohort where higher total HIV DNA levels in PLHIV were associated with higher odds in carotid plaque, cardiovascular disease, and malignancies^70^.

To assess the extent to which endotype-comorbidity associations reflected historical disease severity rather than contemporary biological profiles, we performed a series of sensitivity analyses adjusting for demographic, clinical, HIV-related, vaccination, and study-centre variables individually (**Supplementary Table 11**). Most associations, including carotid plaque, residual viremia, gastrointestinal disease, and musculoskeletal disorders, retained the same direction of effect across models. In contrast, several AIDS-defining outcomes were attenuated after adjustment for CD4 nadir, indicating that historical immunological disease severity contributes to these associations. We further repeated the analyses in participants diagnosed after 2015 who initiated ART immediately under contemporary treatment guidelines, thereby minimizing variation in pre-ART disease accumulation (**Supplementary Table 12**). Effect estimates remained broadly directionally consistent with those observed in the full cohort, although statistical power was reduced because of the smaller sample size, while AIDS-related outcomes remained the most sensitive to adjustment for CD4 nadir.

*Mediation analyses of total HIV DNA.* Because total HIV DNA has previously been linked to comorbidity risk in this cohort^70^, we tested whether it mediated the significant associations between endotype membership and comorbidity prevalence. We first assessed the impact of adding total HIV DNA as a covariate to the unadjusted and adjusted endotype-comorbidity models. Across most outcomes, inclusion of total HIV DNA levels substantially attenuated the endotype effect and often removed statistical significance (e.g., *All High* vs. *All Low* for carotid plaque: OR dropped from 1.69 to 1.16, P=0.56; **Supplementary Table 13**). This attenuation suggests shared variance between endotype identity and reservoir size. However, attenuation does not establish full reservoir mediation, meaning a scenario where adjusting for total HIV DNA would eliminate the endotype-comorbidity association entirely. In contrast, partial mediation would imply that the endotype effect is reduced but not fully removed after adjustment, indicating that both endotype membership (likely underlying biology) and reservoir size contribute to comorbidity.

To confirm this, formal mediation analyses were performed both unadjusted and adjusted for confounders (see **Methods**, **Supplementary Table 13**); here we report the adjusted estimates. For *All High* versus *All Low*, total HIV DNA did not mediate the higher odds of carotid plaque, musculoskeletal disorders, or AIDS-defining malignancies (estimates *P*>0.05). In contrast, AIDS-defining opportunistic infections (effect estimate: 0.16 *P*<0.001) and residual viremia (baseline 0.19 and follow-up 0.21, *P*<0.001) showed significant mediation via the reservoir, but not via direct endotype effect (P>0.10). Other AIDS-defining events showed partial reservoir mediation, supported by significant mediated reservoir effect (0.17 *P*<0.001) and direct endotype effect (−0.09 *P*=0.042). For *Mixed* versus *All Low*, carotid plaque displayed significant reservoir mediation only before adjustment (0.10 *P*=0.010), which disappeared after adjusting for age and current smoking effects (*P*=0.143), suggesting they confound the endotype-reservoir-plaque pathway. AIDS-defining malignancies (0.08, *P*=0.002), residual viremia (baseline 0.15 and follow-up 0.14, *P*<0.001), and post-cART opportunistic infections (0.067 *P*=0.020) showed full reservoir mediation, while gastrointestinal disease did not (*P*=0.53). For *All High* versus *Mixed*, residual viremia (baseline 0.04 and follow-up 0.06, P<0.005), as well as AIDS-defining opportunistic infections and other AIDS-defining events (0.02-0.04, P<0.005) showed full reservoir mediation.

In summary, total HIV DNA mediated only some endotype-comorbidity associations, indicating that total reservoir burden contributes significantly to, but does not fully explain, differences in comorbidity prevalence across the three endotypes. Reservoir levels progressively increased from *All Low* to *Mixed* to *All High*, largely composed by defective proviruses (86-92%), mirroring increasing odds of atherosclerosis, residual viremia, and AIDS-defining: malignancies, opportunistic infections, and related events. These results link the three endotypes to distinct clinical signatures and underscore that both total reservoir burden and underlying host biological variation contribute to comorbidity outcomes. Insights into their underlying biological pathways may identify specific interventions for each endotype.

## Discussion

Integrative multi-omics profiling of 1,230 virally suppressed PLHIV identified three biologically and clinically distinct immune-reservoir endotypes, distinguished by total and intact proviral DNA, together with host immune polarization (Th1/Th2), differentiation states (naïve-like/memory), and cytokine response capacity (type I/II interferon). By jointly integrating total and intact HIV DNA with host multi-omics, HIV reservoir heterogeneity emerged as distinct immune-reservoir endotypes rather than a continuum of reservoir size or individual immune features. This approach, not taken in prior studies^16,17^, revealed a previously unrecognized *Mixed* endotype: individuals with high total but low intact HIV DNA, demonstrating that immune profiles associated with reservoir burden distinct to those associated with reservoir intactness. Importantly, comorbidities also differed among these endotypes: individuals with larger total reservoir levels (*All High* and *Mixed*) showed higher odds of carotid plaque, AIDS-defining malignancies, and residual viremia (at inclusion and 2-year follow-up) compared to *All Low,* with *Mixed* showing an intermediate burden. Total HIV DNA levels mediated only part of these associations, indicating that immune programs remain clinically relevant beyond reservoir size alone, and that *Mixed* and *All High* individuals are not clinically equivalent. This distinction has important implications for HIV cure strategies. Current interventions primarily target intact HIV DNA, yet the *Mixed* endotype, representing one third of our cohort, retains high total HIV DNA despite low intact HIV DNA. After “cure” of subjects in the *Mixed* endotype, relatively large total HIV DNA levels could remain capable of adverse effects if transcriptionally reactivated, underscoring the importance of considering immune-reservoir endotypes rather than reservoir burden alone for designing effective cure interventions.

The *All Low* endotype, with small and less intact reservoirs, displayed a Th1/Tfh1-skewed and less differentiated naïve-like immune landscape enriched for *TCF7* (TCF-1) expression, translational control programs, and stronger IFN-γ production that underscores the role of type II interferon in antiviral surveillance^25,71^. This endotype mirrors characteristics of elite and post-treatment controllers^72,73^, in whom TCF-1-associated maintenance of naïve and stem-like HIV-specific CD8 T-cells restrict proviral burdens^37^, while allowing effective type II IFN-mediated immune responses linked to CD8+ T cells in controllers^74^. Remarkably, a recent bNAb analytical treatment interruption study^75^ showed that pre-intervention CD8⁺ T-cell stemness, marked by *TCF7*-high cells with superior proliferative/cytolytic capacity, reduced exhaustion, and interferon-responses gene enrichment is the strongest predictor of post-intervention control of viremia. This parallels the coordinated stemness and interferon-driven immunity that hallmark our *All Low* endotype and suggests that these individuals may be biologically favored for HIV control following intervention. Together with reduced inflammatory and glycosylation markers (IL-1β, *MAN1C1*, *MMP28*), proviral accumulation may be limited independently of replication competence.

In contrast, the *All High* endotype, marked by large and intact reservoirs, showed Th2/Tfh2 polarization, type I interferon (IFN-I) enrichment (*IFI44L*, SLAMF7), and evidence of chronic activation and exhaustion, signatures paralleling the persistent IFN-I signaling seen in chronic HIV and lymphocytic choriomeningitis virus (LCMV) models, such as Clone 13 infection, where sustained IFN-I signaling drives T-cell exhaustion, suppresses Th1 responses, and promotes viral persistence^71,76,77^. Glycosylation and matrix-remodeling signatures (*MAN1C1*, *MMP28*) further suggest adaptive immune evasion, paralleling viral strategies observed in chronic infection models^29,78^.

The *Mixed* endotype challenged the assumption that total HIV DNA simply reflects replication-competent reservoirs^21^. Despite high total DNA, *Mixed* individuals harbored fewer intact proviruses and combined features of both extremes: inflammatory and glycosylation signals like *All High* yet preserved Th1/IFN-γ function like *All Low*. This intermediate state captured a biological gradient, possibly shaped by immune pressures^15^, linking total reservoir burden to T-cell differentiation and IFN-I activity, and reduced intactness to Th1/Tfh1 and type II interferon (IFN-γ)-mediated immune control.

Our findings both corroborate and substantially extend recent reservoir-based stratification studies in PLHIV. Consistent with Semenova et al.^16^ and, in part, Wang et al.^17^, several immune features distinguishing our *All Low* and *All High* endotypes, including naïve T-cell depletion, reduced CD127 expression, increased activation and exhaustion, and inflammatory leukocyte-vascular activation programs, have been reported previously, supporting the existence of a reproducible immune-reservoir axis across independent cohorts. However, the principal contribution of this study is not the identification of additional immune markers associated with HIV reservoir burden, but the demonstration that joint integration of total and intact HIV DNA with complementary host multi-omics reveals biologically and clinically meaningful immune-reservoir endotypes that cannot be inferred from individual immune features or reservoir measurements in isolation.

Our study extends these observations in three important ways. First, whereas Semenova et al.^16^ clustered on total reservoir-associated immune features and assessed intact HIV DNA post hoc, our joint integration of total and intact HIV DNA enabled identification of the *Mixed* endotype, characterized by high total but low intact HIV DNA, a subgroup not identified in previous clustering studies. This endotype demonstrates that immune programs associated with reservoir burden can differ from those associated with reservoir intactness, revealing biological heterogeneity that would remain hidden when these dimensions are considered independently. Second, jointly integrating reservoir measurements with six host molecular and functional immune layers, including transcriptomic, epigenetic, proteomic, immunophenotypic and functional cytokine-response data, in a seven-to ten-fold larger discovery-validation cohort demonstrated that these endotypes represent coordinated biological states involving multiple molecular layers rather than isolated immunological differences, while revealing biological dimensions not captured in previous studies. Third, Semenova et al.^16^ did not assess comorbidities, whereas Wang et al.^17^ included baseline mental health and substance-use characteristics but did not establish associations between reservoir phenotypes and major HIV-associated comorbidities or longitudinal clinical outcomes. Our comprehensive assessment linked immune-reservoir endotypes to distinct comorbidity and virologic burden, among them carotid plaque, residual viremia and AIDS-defining events, with associations remaining broadly consistent after adjustment for disease severity and restriction to post-cART era events. These findings establish the clinical significance of immune-reservoir endotyping and provide a translational framework for personalized reservoir-targeted strategies.

Collectively, our findings support an immune-reservoir framework in which endotype burden of reservoir size and intactness connect to opposing coordinated immune programs: naïve/stem-like subsets, translational control, and Th1/Th17/type II interferon dominance linked to smaller and less intact reservoirs, versus more differentiated Th2-skewed subsets, type I interferon activity, and glycosylation^28,30^ (*MAN1C1*), especially present in CD4 cells, linked to larger reservoirs. This supports an immune-reservoir gradient underlying the clinical and virologic heterogeneity in our cohort. This framework is consistent with previous observations that persistent IFN-I signaling promotes chronic activation and reservoir stability, whereas type II IFN-driven Th1 immunity contributes to functional restriction^28,71^. It also aligns with emerging evidence that optimal antiviral control depends on a combined axis of CD8⁺ T-cell stemness and interferon pathways, as observed in post-intervention controllers. Importantly, the *Mixed* endotype adds a previously unrecognized dimension to this framework by combining immune characteristics of both extremes

Reservoir size and composition are a central biological axis of HIV persistence, and their integration with host data allowed cut-points in total and intact HIV DNA to emerge from shared virus-host biological structure rather than arbitrary thresholds. This approach revealed that three, rather than two or four, immune-reservoir endotypes best captured the underlying biology and increased sensitivity to biologically relevant immune pathways. Notably, clinical outcomes were not used in clustering yet aligned strongly with the resulting groups, supporting the biological and clinical relevance of the uncovered pathways.

Therapeutically, these mechanistic gradients open the door to further research on the potential of distinct intervention points. *All Low*-like profiles, enriched for TCF-1⁺ stemness akin to predictors of post-intervention control^75^, may benefit from strategies that preserve or amplify this naïve/stem-like compartment. Strategies enhancing IFN-γ surveillance (e.g., IL-12/IL-18 stimulation or checkpoint blockade, linked to reservoir size), while preserving TCF-1⁺ naïve-like pools (e.g., Wnt agonists, linked to total reservoir intactness) may shift *Mixed* or *All High* endotypes towards more effective immune control. Glycosylation modifiers (e.g., *MAN1C1* targets) could disrupt survival niches sustaining proviruses infectivity and immune evasion. The *Mixed* endotype, between both extremes, exemplifies individuals where interventions must balance immune surveillance enhancement with control of chronic inflammation.

The large, discovery-validation cohort and integrative design strengthen these insights, though causal inference remains limited by the cross-sectional setting. While internal validation was applied throughout, establishing generalizable predictors of immune-reservoir endotypes will require confirmation in independent cohorts. Modest model accuracy and the Dutch-centric cohort emphasize the need for diverse, longitudinal, and tissue-based studies. Pneumococcal vaccination and tuberculosis-related history should also be considered in future studies when interpreting stimulus-specific cytokine responses. Nonetheless, this integrative multi-omics framework links determinants of HIV reservoir heterogeneity, revealing mechanistic gradients that connect host immune differentiation, interferon balance, glycosylation, and reservoir composition, offering a conceptual and analytical foundation for personalized HIV cure strategies targeting both the size and the quality of the reservoir in virally suppressed PLHIV.

## Methods

### Study Population

We analyzed 1,230 people living with HIV (PLHIV) from the 2000HIV cohort (NCT03994835) with available HIV DNA data, enrolled across four clinical centers in the Netherlands. Written informed consent was received from participants prior to inclusion in the study. All experiments with human samples were conducted according to the principles expressed in the Declaration of Helsinki. Participants were divided into discovery (n = 1,027; three centers) and validation (n = 203; one center) sub-cohorts prior to analytical preprocessing and downstream analyses to ensure robustness and reproducibility of the results. Eligibility criteria included age ≥18 years, confirmed HIV-1 positivity, combination antiretroviral therapy (cART) for ≥6 months (if treated), and plasma viral load <200 copies/mL. Full inclusion criteria and study design details have been described previously^20^.

### Analytical use of discovery and validation sub-cohorts

The discovery and validation sub-cohorts were defined a priori by clinical centre before preprocessing and analysis, comprising three clinical centers as the discovery sub-cohort (n=1,027) and an independent clinical center as the validation sub-cohort (n=203) participants, respectively. After removal of consensus outliers, 1,002 participants in discovery and 189 in validation were included in the final clustering and downstream analyses. The discovery sub-cohort was used to determine the optimal number of endotypes (*k = 3*) and select the integrative clustering solution. The same analytical framework was then applied independently in the validation sub-cohort to assess replication of the three-endotype structure. Single-layer flow-cytometry, transcriptome and ex vivo cytokine-responses analyses were performed separately in the two sub-cohorts. Findings in the single-layer omic analyses were considered replicated when they met the prespecified significance threshold in discovery (false-discovery-rate-adjusted *P < 0.05*, where applicable), showed nominal significance in validation (*P < 0.05*), and had a consistent direction of effect in both sub-cohorts. Pathway-enrichment analyses in bulk transcriptome were restricted to the discovery sub-cohort (*n = 1,002*) because of the limited validated markers across sub-cohorts and were interpreted as supportive rather than independently replicated findings. Descriptive demographic and clinical characteristics were evaluated separately in discovery and validation. Endotype-comorbidity and mediation analyses were conducted in the pooled post-outlier cohort (*n = 1,191*) to enable analysis in less frequent clinical outcomes. Similarly, for the supervised predictive classification analyses, data were partitioned using a stratified 80% training and 20% held-out test split; model tuning was performed within the training set, and final performance was evaluated in the held-out test set that was not used during model development.

### Sample Processing

Biological body fluids were collected during daytime hours at the participating centers and sent overnight to the Radboudumc. The overnight shipping was at room temperature and samples were processed immediately in the morning the day after the day of baseline visit. The samples collected in the coordinating center, Radboudumc, were also stored overnight at room temperature to ensure identical handling across all centers. Upon arrival of the samples in the lab, whole blood obtained from EDTA tubes was used for hemocytometric analysis with the XN Sysmex haematology analyzer (Sysmex, Kobe), and for immunophenotyping using flow cytometry (CytoFLEX-LX, Beckman Coulter). Peripheral blood mononuclear cells (PBMC) were isolated using density gradient separation (Ficoll-paque) in SepMate™ tubes, as previously described^79^. Whole blood was used for DNA isolation. Plasma (citrate and EDTA) and serum were aliquoted and stored at -80°C for future assessment.

### Multi-omics Data Acquisition and Preprocessing

Multi-omics profiling was performed on samples from 1,230 PLHIV, encompassing bulk RNA sequencing (RNA-seq, 58,347 genes, PBMCs), single-cell RNA-seq (scRNA-seq, PBMCs), plasma proteomics (Olink Explore, 2,367 proteins, plasma), DNA methylation (793,775 CpG sites, EDTA whole blood), flow cytometry (355 immune-cell subpopulations, whole blood), ex vivo cytokine production (90 markers, PBMCs), and HIV DNA quantification (total/intact copies per million CD4^+^ T cells, CD4^+^ T cells from PBMCs). Prior to integrative clustering, highly variable features were selected to reduce noise in high-dimensionality datasets where top 5,000 genes (RNA-seq, based on mean absolute deviation), 7,938 CpG sites (methylation, top 1% variable) were selected, as well as log2-transformed data for all proteomics, flow cytometry, cytokines, and HIV DNA measurements. Single-layer analyses used all available features per *omic* layer, adjusted for confounders (see *Confounder Selection for Single-Layer Analyses* and *Immune Characterization* sections).

### Bulk RNA-seq

Peripheral blood mononuclear cells (PBMCs) were isolated and viably frozen as previously described^20,80^. Briefly, PBMCs were isolated by Ficoll-Paque density gradient centrifugation, frozen in 90% fetal calf serum and 10% DMSO using a CoolCell container at −80°C, and stored in liquid nitrogen. For RNA sequencing, PBMCs were thawed and cultured in RPMI 1640 medium supplemented with 10% FCS, 2 mM GlutaMAX, 1 mM pyruvate, and 50 µg/mL gentamicin. RNA was extracted and sequenced on the Illumina platform, generating >30 million paired-end reads per sample. Reads were aligned to the human genome (GRCh38, Gencode v33) using STAR (v2.7.3a), and gene-level quantification was performed with HTSeq-count. Genes with <5 counts in ≥50% of samples were removed. Normalization and variance-stabilizing transformation (VST) were performed using DESeq2 (v1.34.0). The top 5,000 most variable genes (excluding genes on the Y chromosome), ranked by mean absolute deviation (MAD), were selected for unsupervised clustering.

### Single-cell RNA-seq

Single-cell RNA sequencing (scRNA-seq) was performed on PBMCs from a subset of 200 individuals, using the BD Rhapsody™ Single-Cell Analysis System (BD Biosciences). PBMCs were isolated and cryopreserved as described above and thawed in RPMI 1640 medium supplemented with 10% FCS prior to single-cell suspension preparation. Extensive details on single-cell workflow and library preparation are described elsewhere^20,80^. Sequencing data were processed to obtain UMI count matrices, which were imported into R (v4.1.0) and analyzed using Seurat (v4.0.4). Cells with >10% mitochondrial RNA content or <200 expressed genes were excluded. Genes expressed in fewer than three cells, as well as MT-RNR1 and MT-RNR2, were removed. Gene expression data were log-normalized and scaled, regressing out the number of detected transcripts per cell. Dimensionality reduction was performed via PCA using the top 2,000 most variable genes (vst method), followed by UMAP embedding using the first 30 principal components. Cell types were annotated based on clustering and canonical marker gene expression.

### Plasma Proteomics

Circulating plasma protein expression was measured using a commercially available multiplex proximity extension assay (PEA) from Olink® proteomics AB (Uppsala, Sweden), as previously described^81^. Depending on the batch, this study utilized either the Olink® Explore 1536 (1,472 proteins) or Explore 3072 (3,072 proteins) platform, which included targeted proteins organized into panels focusing on inflammatory, oncological, cardiometabolic, and neurological proteins. Protein measurements are delivered as Normalized Protein Expression (NPX) values following a quality control (QC) and normalization process developed and provided by Olink. Plasma proteins were measured in multiple batches, and therefore, bridging normalization was performed to remove any batch effect between the panels. Bridging normalization was performed by following the next steps for each protein: (1) we first calculated the median of the bridging samples for each protein in the two batches; (2) subsequently, we calculated the median difference keeping one batch as a reference; (3) finally, we subtracted the median difference from each protein in the non-reference batch. Limit of detection (LOD) values per protein were re-adjusted by the same adjustment factor as the respective protein measurements after bridging normalization. After removing batch effects using bridging normalization, standard QC per protein and sample was performed prior to statistical data analysis. Technical duplicates (e.g., IL-6, TNF, CXCL8) were measured across panels for QC purposes. Strong correlations were observed between the technical duplicates among panels (Spearman rho correlation r > 0.9), and therefore, we selected the measurements from one panel. Next, we excluded proteins with lower limit of detection (LOD) in >25% of the samples. Finally, samples were defined as outliers using PCA analysis of the NPX values. Outlier samples were defined as those that fell more than 3 standard deviations from the mean of principal component one (PC1) and/or two (PC2). After QC, data were log2-transformed and adjusted for plate controls and proximity extension effects for follow-up analysis.

### DNA Methylation

Genomic DNA was extracted from EDTA whole blood using the ChemagicStar system (Hamilton Robotics, with Chemagen Magnetic Separation Module 1) and M-PVA beads (PerkinElmer). DNA was normalized to 50 ng/µL in TE buffer, and 500 ng was bisulfite-converted using the EZ DNA Methylation Kit (Zymo Research, no. D5001). Methylation was profiled with the Illumina Infinium MethylationEPIC BeadChip (v1.0 B5, no. 850K). Raw IDAT files were processed with minfi (v4.2.0), excluding samples with gender mismatches, poor call rates (<99%), or low-quality probes. Probes with >10% missing values, sex chromosome mapping, or polymorphisms (MAF >5%, European populations) were removed. Stratified quantile normalization yielded β-values, transformed to M-values [log2(β/(1−β))] for analysis. Top 1% variable CpG sites (7,938) were selected for clustering based on mean absolute deviation.

### Flow Cytometry

Peripheral blood samples were immunophenotyped using three high-dimensional flow cytometry panels on a 21-color, six-laser CytoFLEX LX instrument (Beckman Coulter). Data were acquired using CytExpert software (v2.3) and analyzed with Kaluza software (v2.1.2) using conventional gating strategies. Antibodies were selected to identify 355 immune-cell subpopulations, covering major innate, T-, and B-cell populations. Markers including HLA-DR, CD38, PD-1, PD-L1, CD40, CD307d, and CD81 were included to assess activation, exhaustion, maturation status, and intercellular communication. Daily quality control was performed using CytoFLEX Daily QC Fluorospheres (Beckman Coulter, #B53230), CytoFLEX Daily IR QC Fluorospheres (Beckman Coulter, #C06147), and SPHERO™ Rainbow Calibration Particles, 6-peak (Spherotech, #RCP-30-5A-6). Absolute immune cell counts were calculated based on white blood cell counts per mL of whole blood. Data were log2-transformed for integration in the multi-omics clustering analysis. Antibody selection and gating strategies have been described in detail elsewhere^47^.

### Ex Vivo Cytokine Production

PBMCs were isolated and stimulated with a panel of whole inactivated pathogens, pattern-recognition receptor (PRR) ligands, and viral antigens using the validated Human Functional Genomics Project (HFGP) protocol^53^, as previously detailed in the 2000HIV cohort design^20^. Consistent with the validated HFGP framework, cytokine readouts were interpreted according to the combination of stimulation duration, cytokine identity and stimulus context. Cytokines measured after 24 hours, including IL-1β, IL-1Ra, IL-6, IL-8, IL-10, MCP-1, MIP-1α and TNF, predominantly reflect short-term monocyte-associated inflammatory responses. Cytokines measured after 7 days, including IFN-γ, IL-5, IL-10, IL-17 and IL-22, were interpreted as reflecting longer-term T-helper-associated cytokine-production capacity in mixed PBMC cultures. These classifications do not imply that the cultures contain only one responding cell type or that the readouts represent purified cell-specific responses. For peptide-based stimuli (CMV pp65, HIV Env), monocyte-associated 24-hour cytokines are interpreted as reflecting monocyte activation secondary to rapid antigen-specific memory T-cell recognition, rather than direct pattern-recognition receptor engagement.

For 24-hour stimulations, PBMCs were exposed to Poly I (100 µg/mL, TLR3), LPS (10 ng/mL, TLR4), imiquimod (5 µg/mL, TLR7), recombinant IL-1α (10 ng/mL), CMV pp65 peptide pool (1 µg/mL), HIV Env peptide pool (1 µg/mL), and heat-inactivated S. pneumoniae (5×10⁶/mL). For 7-day stimulations, PBMCs were incubated with E. coli (1×10⁶/mL), S. aureus (1×10⁶/mL), S. pneumoniae (5×10⁶/mL), M. tuberculosis H37Rv (5 µg/mL), C. albicans conidia and hyphae (1×10⁶/mL), and phytohaemagglutinin (10 µg/mL). Supernatants were quantified by ELISA. Because these assays used mixed PBMC cultures and, for several conditions, whole-microbial stimuli containing both antigens and innate ligands the resulting measurements represent integrated ex vivo cytokine-production capacity in mixed PBMC cultures rather than responses attributable exclusively to a single cell population or direct measures of pathogen-specific immunity. Absolute cytokine concentrations were log₂-transformed and integrated into the multi-omics clustering analysis.

### HIV DNA Quantification

HIV-1 DNA was quantified in CD4+ T cells from PBMCs using the Rainbow proviral HIV-1 DNA digital PCR (dPCR) assay, as described in detail elsewhere^82^. Briefly, CD4+ T cells were enriched by negative selection using the EasySep Human CD4+ T Cell Isolation Kit on the RoboSep-S platform (Stemcell Technologies, Vancouver, Canada). Genomic DNA was extracted using the QIAamp DNA Mini Kit on the QIAcube (Qiagen, Hilden, Germany), and DNA concentrations were measured using the SpectraMax Quant AccuBlue HiRange dsDNA Assay Kit on the SpectraMax i3x platform (Molecular Devices, San Jose, CA, USA). HIV-1 DNA was quantified in triplicate by dPCR on the QIAcuity Four platform (Qiagen, version 1.2). Total HIV-1 DNA was quantified using the RU5 region. Intact HIV-1 DNA was defined by the detection of at least two (PSI and ENV) and up to five (RU5, PSI, GAG, POL, and ENV) HIV-1 target regions according to a predefined decision-tree algorithm. Positivity thresholds were determined using the Rainbow Shiny tool and manually adjusted when required. HIV-1 DNA levels were normalized to copies per million CD4+ T cells using the RPP30 duplex assay and corrected for DNA shearing using the DNA Shearing Index (DSI; median 35.9%, IQR 33.6-38.3%). The assay limit of detection (LoD95) was 10 copies per reaction. For samples with undetectable intact HIV-1 DNA, values were imputed as one intact copy divided by the total number of cells tested and corrected for DNA shearing.Total and intact HIV-1 DNA measurements were reported as number of copies per million CD4^+^ T-cells and were log2-transformed for multi-omics clustering.

### Missing Data

Multi-omics data missingness assessment was performed prior to multi-omics clustering. Features with ≥80% missing values were excluded. K-nearest neighbor imputation (impute.knn, R, k = 5) was applied separately within each omics layer and within each sub-cohort (discovery and validation), yielding 15,752 markers (90 cytokines, 355 flow cytometry, 5,000 RNA-seq, 7,938 methylation, 2,367 proteomics, 2 HIV DNA).

### Multi-omics Clustering

Multi-omics clustering was performed to identify host molecular and immune profiles that co-varied with HIV reservoir burden and composition. Each dataset was preprocessed separately and entered as a distinct data layer, with total and intact HIV DNA included as a dedicated reservoir layer rather than converted into predefined high- or low-reservoir categories. During development of the analytical framework, preliminary clustering using the host layers alone produced alternative immune groupings but did not identify participant groups with meaningful differences in reservoir size or intactness and therefore did not address the central biological objective of the study. We consequently integrated the reservoir layer with the host multi-omics layers, allowing reservoir patterns and their associated host states to emerge jointly from the data. The MOVICS pipeline evaluated eight algorithms (*MoCluster*, *NEMO*, *COCA*, *SNF*, *PINSPlus*, *CMOIC*, *CIMLR*, *Consensus*) in the discovery sub-cohort on imputed data (15,752 markers, n=1,230). The optimal cluster number (k=3) was determined using Consensus Partition Index and Gap Statistics on the discovery cohort (k=2-8, **Supplementary Fig. 1**) in discovery and subsequently applied independently in the validation sub-cohort using the same analytical framework. *MoCluster* was selected for significant HIV DNA differences across endotypes (P<0.05, Kruskal-Wallis for continuous variables, Chi-square for categorical variables, **Supplementary Fig. 2**). Before integration, *MoCluster* applied its default consensus PCA block-weighting procedure, in which each data matrix was weighted by the inverse of its first eigenvalue. This block-level normalization places layers with different dimensionalities and variance structures on a more comparable scale and limits the influence of high-dimensional layers arising solely from their larger number of measured features. No additional manual weighting was applied to the reservoir or any other layer, and no variable-specific weights were prespecified. Within each layer, variable loadings on the joint latent components were estimated empirically from the covariance structure of the data. After clustering algorithm selection, fine-tuning of *MoCluster* was performed. *MoClusters’* hyperparameter *k* was optimized using Silhouette, Davies-Bouldin, and Calinski-Harabasz scores. Outliers (25 discovery, 14 validation) were identified using univariate IQR (1.5×IQR), robust Mahalanobis distance (MCD, α=0.75, 99% chi-squared, df=2), Mahalanobis distance on integrative factor scores, and Silhouette scores (Euclidean, threshold<0.1). Consensus outliers (flagged by ≥2 methods) were removed, yielding a final discovery (n=1,002) and validation (n=189) sub-cohorts that were used for final *MoCluster* assessment and downstream analyses. Endotypes were validated across cohorts and visualized via violin plots, silhouette plots, 3D scatterplots, and 2D projections (**Supplementary Figures 3-5, Supplementary Table 14**). Replication was assessed by determining whether the validation sub-cohort reproduced the three-endotype structure, the corresponding patterns of total and intact HIV DNA, and the principal host-layer features observed in discovery.

### Driver Marker Visualization and Partial Correlation Analysis Between Host-Markers and HIV Reservoir

Driver markers consistently separating endotypes in both discovery and validation cohorts were selected. For each omics layer, marker expression values were visualized in boxplots (with violin overlays and jittered points) to observe the distribution of marker values across the three endotypes (“All Low”, “Mixed”, “All High”) in both cohorts. Pairwise endotype differences were tested using two-sided t-tests with Benjamini-Hochberg correction. To control for demographic and technical confounders, marker values were adjusted using linear models. Specifically, expression values were regressed on age and sex at birth). Both unadjusted and residualized values were retained for visualization and subsequent analyses. Associations with viral reservoir size (total and intact HIV DNA per 10⁶ CD4⁺ T cells) were evaluated using Spearman correlations and partial Spearman correlations (adjusting for age and sex at birth) implemented in the ppcor R package (v1.1). Confidence intervals were estimated via Fisher’s z-transformation. Analyses were performed separately in discovery and validation cohorts, with p-values FDR-adjusted in the discovery cohort and reported as nominal in the validation cohort. Endotype-specific associations (Mixed, All Low, All High) were assessed to capture heterogeneity across groups. Results were visualized as scatterplots with regression fits (per marker) and as heatmaps summarizing effect sizes and significance (∗FDR or nominal p < 0.05, ∗∗p < 0.01).

### Clinical and Comorbidity Characterization

Demographic (age, sex, ethnicity, smoking status, CMV serology), clinical (BMI, HIV duration, cART duration, CD4 nadir, viral load zenith, latest CD4/CD8 counts, residual viremia, HIV-1 DNA reservoir), ART exposure (drug classes, specific drugs, treatment interruptions), and comorbidity data (cardiovascular disorders, hypertension, hypercholesterolemia, hypertriglyceridemia, myocardial infarction, stroke, angina pectoris, carotid plaque, intima-media thickness, non-AIDS cancer, endocrine-metabolic, pulmonary, gastrointestinal, CNS, psychiatric, and musculoskeletal disorders) were extracted from baseline clinical records and measurements (NCT03994835). These variables were compared across multi-omics endotypes in discovery and validation cohorts. Continuous variables were analyzed using T-tests (normally distributed) or Kruskal-Wallis tests (non-normally distributed), and categorical variables were analyzed using Chi-square tests, with multiple testing correction (Bonferroni, P<0.05) using the compareGroups package in R (v4.3.2). Results are reported in **Supplementary Tables 6-8**.

*Odds Ratios Calculations.* To further assess associations between endotype membership and clinical comorbidities, we performed logistic regression analyses in the pooled discovery and validation cohort (n = 1,191). Potential covariates were selected *a priori* based on established HIV and cardiovascular risk factors (Centers for Disease Control and Prevention, 2023; Trickey et al., 2021), including age, sex, BMI, smoking status, HIV duration, ART duration, clinical center, CMV IgG levels, ethnicity, hepatitis C status, and CD4 nadir. Binary outcomes were encoded as factors (“No”/“Yes”). Only complete cases were analyzed per model. Right-skewed continuous covariates (e.g., CD4 nadir) were log-transformed to approximate normality. Confounder selection followed a data-driven change-in-estimate strategy, where variables were retained if they altered the odds ratio by ≥10% in any exposure group, as recommended in epidemiological research (Greenland, 1989; Lee et al., 2019). Covariates were screened for outcome and exposure association (*p*<0.05), and multicollinearity was addressed by removing highly correlated variables (Pearson *r*>0.5). Final models were assessed for collinearity using variance inflation factors (VIF<5), and their performance was evaluated using five-fold cross-validation and area under the receiver operating characteristic curve (AUC). Rare outcomes (e.g., hyperthyroidism, *n*=4) were excluded from primary analyses due to model instability. Unadjusted and adjusted odds ratios (ORs) with 95% confidence intervals were calculated for all pairwise endotype comparisons (*All High* vs. *All Low*, *All High* vs. *Mixed*, *Mixed* vs. *All Low*). Forest plots were generated for each comparison, presenting odds ratios and p-values of the adjusted and unadjusted models. All analyses were conducted in R (v4.3.0).

### Mediation Analysis

To evaluate whether total HIV DNA levels mediated the association between endotype membership and clinical comorbidities, we performed causal mediation analyses using both model-based (a×b) estimates and the nonparametric bootstrap framework implemented in the *mediate()* function (Imai et al., 2010). Analyses were conducted using the pooled cohort (n=1,191) and included all binary clinical outcomes that were significantly associated with endotype membership in adjusted odds-ratio models. For each pairwise endotype comparison, the exposure was coded as a binary variable, the mediator was log-transformed total HIV DNA, and outcomes were modeled using logistic regression. Confounders were prespecified based on the same HIV and cardiometabolic risk factors used in the odds-ratio analyses and were applied per outcome based on the corresponding adjusted model. The mediation workflow consisted of three components. (1) *Mediator model (a-path):* linear regression of total HIV DNA on endotype. (2) *Outcome model (b-and cll-paths):* logistic regression of each binary comorbidity on endotype and total HIV DNA, with and without confounders. (3) *Estimation of indirect and direct effects:* we computed the model-based indirect effect (a×b) on the log-odds scale, its delta-method confidence interval, and optional nonparametric bootstrap confidence intervals (5,000 resamples). In parallel, the *mediate()* function provided the average causal mediation effect (ACME), average direct effect (ADE), total effect, and proportion mediated, using 5,000 bootstrap simulations and the nonparametric potential-outcomes framework. Significance of mediation was determined using ACME (p<0.05 or 95% CI excluding 0). Cases where ACME was significant but ADE remained non-zero (p<0.05) were classified as *partial mediation*, whereas non-significant ADE indicated *complete mediation*. All mediation results were compiled across adjusted and unadjusted models, with complete-case data used for each mediator-outcome pair. Analyses were conducted in R (v4.3.0).

### Confounder Selection for Single-Layer Analyses

Confounders for single-layer analyses (bulk RNA-seq, flow cytometry, and ex vivo cytokine production) were selected to adjust for potential biases in discovery (n=1,002) and validation (n=189) cohorts. A two-step approach was used: (1) Principal component analysis (PCA) was performed on raw omics data from the discovery sub-cohort using FactoMineR (v2.4) in R (v4.3.0), with the first 10 principal components regressed against potential confounders (e.g. age, sex, ethnicity, center, seasonality (sine/cosine functions), time to lab, COVID-19 vaccination, HIV-related variables (e.g., cART duration, CD4 nadir)) via linear regression. (2) Confounders causing >10% change in beta coefficients for omics-clinical variable associations were selected. Selected confounders included age, sex, seasonality (sine and cosine scores), collection center, time to laboratory preprocessing, COVID-19 vaccination status, and first genetic principal component accounting for ethnicity, applied consistently across sub-cohorts. COVID-19 vaccination status was considered because participant sampling overlapped the COVID-19 pandemic and vaccination rollout, and vaccination could contribute to broader variation in circulating immune-cell, transcriptional and inflammatory profiles, independently of SARS-CoV-2-specific responses. SARS-CoV-2-specific immune responses were not measured in this study. See **Supplementary Figures 22-24**.

### Immune Characterization Analyses

The discovery sub-cohort was used to identify significant single-layer features, and the validation sub-cohort was used to assess their replication according to the criteria specified below. Baseline and stimulated immune profiles were analyzed separately in discovery (n=1,002) and validation (n=189) cohorts, adjusting for confounders via linear models (beta coefficient increase >10%).

### Baseline Immune Landscape

Flow cytometry (355 immune-cell subpopulations, absolute counts/percentages) used linear regression with inverse rank-based transformation. Data were inverse rank-transformed using the qnorm function (R v4.3.0) to achieve normality, then analyzed via linear regression, adjusting for season of sample collection, age, sex, the first genetic principal component, COVID-19 vaccination status, and time from sample collection to laboratory processing (see *Confounder Selection for Single-Layer Analyses*). Significant differences required FDR-adjusted P<0.05 (discovery), nominal P<0.05 (validation), and consistent effect estimates direction. RNA-seq data (58,347 genes, unstimulated PBMCs) were analyzed using DESeq2 (v1.34.0) with negative binomial models, Benjamini-Hochberg correction, and apeglm shrinkage. Models were adjusted for season of sample collection, age, sex, the first genetic principal component, COVID-19 vaccination status, and time from sample collection to laboratory processing. Differentially expressed genes needed FDR-adjusted P<0.05 (discovery), nominal P<0.05 (validation), and consistent log2 fold change direction.

### Gene Set Enrichment Analysis (GSEA)

To assess pathway-level differences in bulk transcriptomic data, we performed gene set enrichment analysis (GSEA) on differential expression results from the discovery cohort (n=1,002), given its larger sample size. Genes were ranked by a signed statistic defined as −log_10_(P) × sign(log_2_ fold change), where P corresponds to the nominal DESeq2 P-value. Ensembl gene identifiers were mapped to Entrez IDs and HGNC symbols via biomaRt (v2.58.0). For Entrez IDs mapping to multiple Ensembl IDs, the transcript with the most extreme ranking metric was retained. Gene set enrichment was performed using clusterProfiler (v4.10.0) and ReactomePA (v1.46.0) against Reactome pathways (minimum 10, maximum 500 genes per set), with significance defined as Benjamini-Hochberg adjusted P<0.05. Normalized enrichment scores (NES), adjusted P-values, and leading-edge genes were extracted for interpretation. Results were visualized with custom functions built on ggplot2 (v3.5.0), enrichplot (v1.22.0), and cowplot (v1.1.3), highlighting the most significantly enriched upregulated and downregulated pathways per comparison. Pathway-enrichment results were therefore interpreted as discovery-cohort supportive evidence and were not considered independently replicated findings.

### Stimulated Immune Functionality

Ex vivo cytokine production was assessed in discovery (n=1,002) and validation (n=189) cohorts, comparing multi-omics endotypes. Peripheral blood was stimulated for 24h (Poly I:C, LPS, Imiquimod, IL-1α, HIV-ENV, CMV, S. pneumoniae) or 7d (E. coli, S. aureus, S. pneumoniae, M. tuberculosis, C. albicans conidia/hyphae, PHA). Cytokine levels were analyzed using rank-based regression (rfit), R package Rfit v0.24.6; Bent1 transformation for skewness) in R (v4.3.0), adjusting for season of sample collection, age, sex, the first genetic principal component, COVID-19 vaccination status, and time from sample collection to laboratory processing. Significant associations required FDR-adjusted P<0.05 (discovery), nominal P<0.05 (validation), and consistent effect direction. Missing cytokine-stimulus pairs (e.g., IL-10\PolyIC, S.aureus\IL-5) were excluded from analysis.

### Predictive Modeling

Binary classification models distinguished multi-omics endotypes (*All Low* vs. *All High*, *Mixed* vs. *All High*, *All Low* vs. *Mixed*) using features from bulk RNA-seq, flow cytometry, ex vivo cytokine production, DNA methylation, HIV-1 DNA quantification, and clinical/demographic data (e.g., age, CD4 nadir, cART duration, CMV serostatus) in discovery (n=1,002) and validation (n=189) sub-cohorts. For each comparison, data were divided using a stratified 80% training and 20% held-out test split with a fixed random seed (random_state=42). Multicollinearity was addressed by iteratively removing features with high variance inflation factor (VIF) >10 or an absolute pairwise correlation |r|>0.8 until no features met either criterion (**Supplementary Figures 25-26**). Numerical features were k-nearest-neighbors (KNN)-imputed (n_neighbors=5), winsorized (1% limits), log-transformed (e.g., CD4 nadir, CMV IgG), and scaled (RobustScaler for omics, Yeo-Johnson for clinical). Categorical variables were mode-imputed and one-hot encoded (**Supplementary Figures 27-28**). Random Forest, XGBoost, Logistic Regression, and KNN models were trained (80%/20% train/test split, stratified, random_state=42). Hyperparameters were optimized within the training data using RandomizedSearchCV with 50 iterations and weighted F1-score and area under the receiver operating characteristic curve (ROC AUC) as evaluation metrics. Final model performance was assessed in the held-out test set. XGBoost was selected based on its combined held-out weighted F1-score and AUC performance (**Supplementary Figure 15**), and feature contributions were subsequently evaluated using SHAP analysis (**Figure 3**). The models were used primarily for feature prioritization and comparison of predictive contributions across data layers rather than as stand-alone clinical classifiers. Analyses used scikit-learn (v1.2.2), xgboost (v1.7.3), and shap (v0.41.0) in Python (v3.9.12).

### Software

Analyses used R (v4.3.2; dplyr 1.1.4, robustbase 0.99-4, cluster 2.1.6, minfi 1.40.0, DESeq2 1.34.0, compareGroups 4.6.0, FactoMineR 2.4, ppcor 1.1, Rfit 0.24.6, biomaRt 2.58.0, clusterProfiler 4.10.0, ReactomePA 1.46.0, ggplot2 3.5.0, enrichplot 1.22.0, cowplot 1.1.3, rms) and Python (v3.9.12; scikit-learn 1.2.2, xgboost 1.7.3, shap 0.41.0, seaborn). Seurat (v4.0.4).

## Data Availability

Bulk RNA-seq, DNA methylation, and ex vivo cytokine production data generated in this study are available upon approved request in the Radboud Data Repository (RDR) collection^83^ at https://doi.org/10.34973/p96d-kz55. Plasma proteomics data are available in a separate RDR collection^84^ at https://doi.org/10.34973/qk29-f305. Flow cytometry data are available via controlled access through the RDR collection (ID: ru.rumc.2000hiv_t0000321a_dsc_002). HIV-1 DNA quantification data are available in the RDR collection^85^ at https://doi.org/10.34973/2bf5-yc03. In accordance with the 2000HIV consortium data governance policy and ethical approvals, all data access is subject to data sharing agreements and approval by the 2000HIV data access committee. The 2000HIV cohort is registered at ClinicalTrials.gov (NCT03994835).

## Code Availability

All analyses were performed using established, widely available peer-reviewed software packages cited in the Methods, and no new analytical methods were developed as part of this study. Key analytical scripts used in this study are being made available at: https://github.com/victoriariosvzz2/Multiomics-Clusters-Reservoir-2000HIV. Additional scripts and workflow details are available from the corresponding author upon reasonable request.

