## Supplementary Figures for "Multi-Omics Clustering Differentiates the Total and Intact HIV Reservoirs and Related Host Immune Mechanisms"

**Supplementary Figure 1 |** Identification of Optimal Number of Multi-Omics Clusters

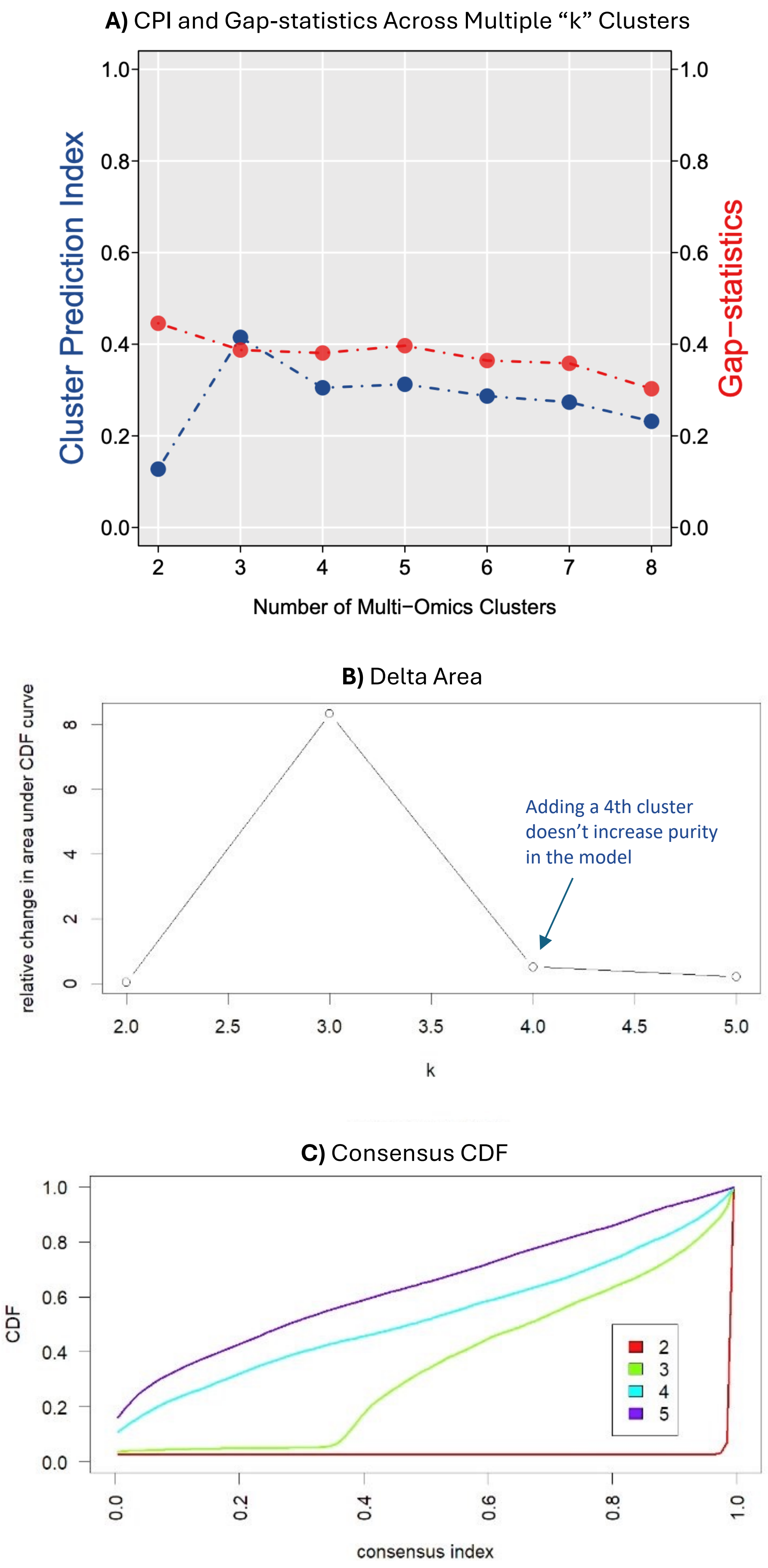


**Supplementary Figure 1. Identification of the Optimal Number of Multi-Omics Clusters. a,**Consensus Partition Index (CPI) plot showing the optimal number of clusters ($k$=3) identified for the discovery cohort. **b,** Relative change in the area under the CDF curve, indicating that adding more than three clusters does not improve cluster purity. **c,** Consensus CDF plot illustrating the stability of clusters, confirming that $k$=3 yields the most stable and interpretable cluster solution.

**Supplementary Figure 2 |** Evaluation of Clustering Consistency Using Different Algorithms

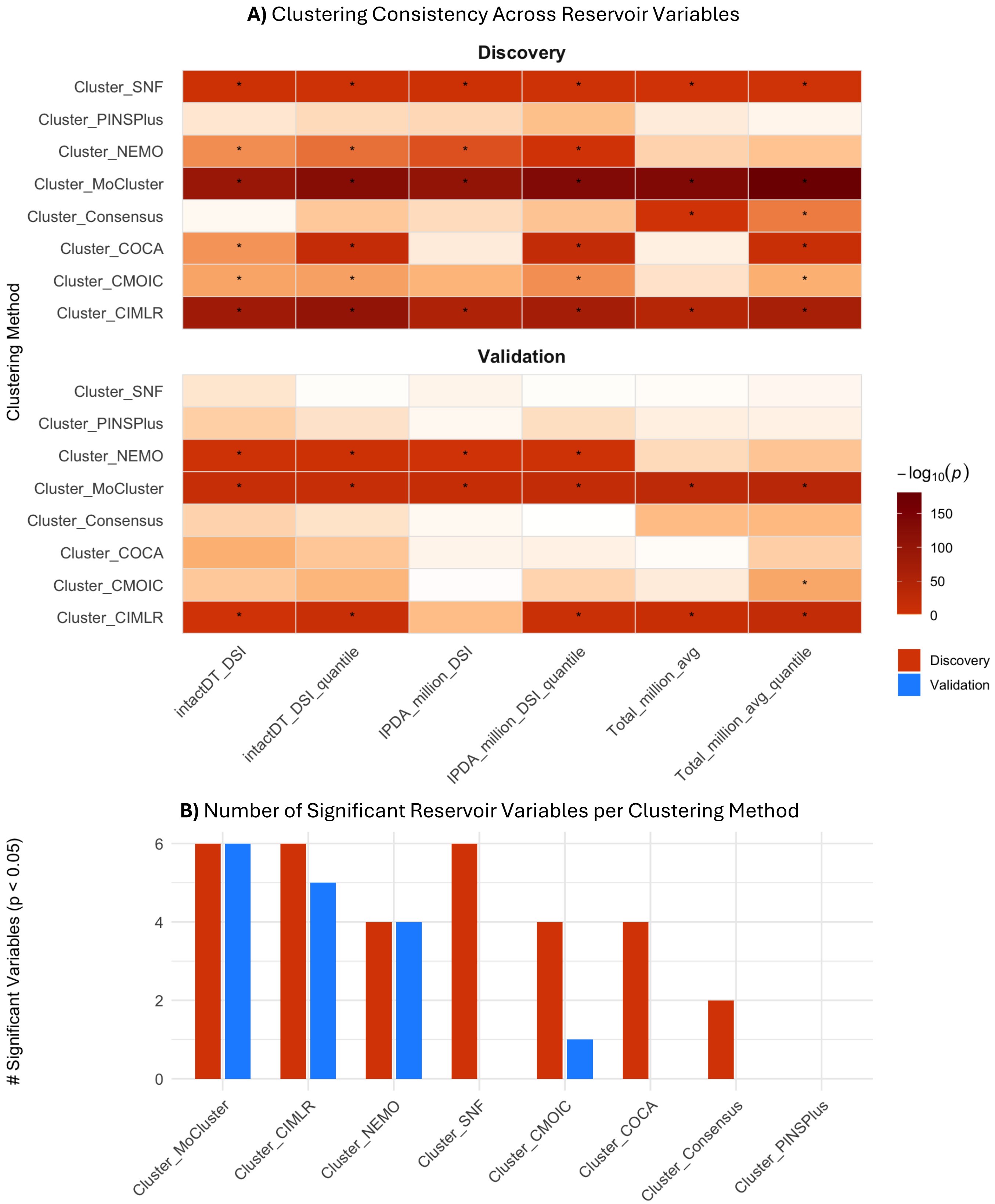


**Supplementary Figure 2. Evaluation of Clustering Consistency Using Different Algorithms. a,** Heatmap of $-\log_{10}$ P-values from statistical tests assessing associations between cluster assignments (from eight multi-omics integration algorithms in the MOVICS framework: SNF, PINSPlus, NEMO, COCA, ConsensusClustering, CIMLR, CMOIC, and MoCluster) and viral reservoir/clinical variables. Continuous variables (Total HIV DNA, Intact HIV DNA, IPDA) were tested with the Kruskal–Wallis rank-sum test; corresponding categorical quartile variables (e.g., Total HIV DNA quantiles) were tested with chi-square tests. The x-axis shows viral reservoir features, and the y-axis shows clustering methods. Heatmap colors represent $-\log_{10}(P)$, with deeper red indicating stronger associations; asterisks denote $P < 0.05$. MoCluster, NEMO, and CIMLR exhibited the strongest associations across most variables, with MoCluster consistently performing well for both continuous and categorical traits. **b,** Bar plot showing the number of significantly associated variables ($P < 0.05$) for each clustering method in the discovery and validation cohorts. Bars are ordered by decreasing number of significant associations. MoCluster had the highest counts and greatest cross-cohort consistency, followed by CIMLR and NEMO.

#### **Supplementary Figure 3 |** Consensus Outliers Assessment in Multi-Omics Factors Space

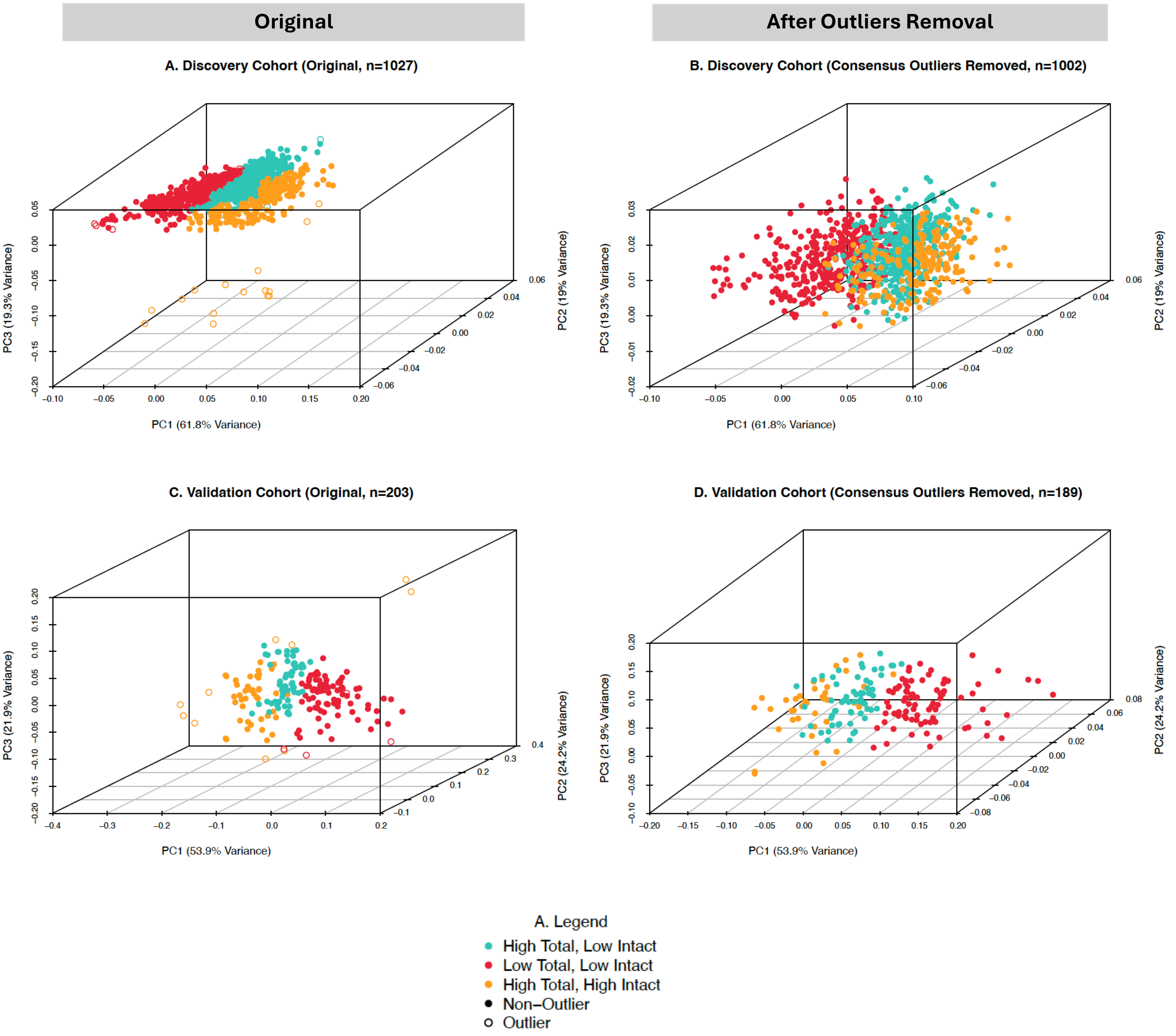


**Supplementary Figure 3: Multi-Omics Factor Scores and Cluster Fit Assessment.**
This figure displays 3D scatterplots of MoCluster-derived multi-omics factor scores (PC1, PC2, PC3) to visualize cluster structure and outlier detection in both the discovery and validation cohorts. Points are colored by cluster assignment and shaped by consensus outlier status (filled circle: non-outlier; open circle: outlier).

(A) and (C) show the original datasets (including outliers), while (B) and (D) present the datasets after removal of consensus outliers. Consensus outliers were identified using an iterative approach combining three methods applied to the integrated latent space (PC1, PC2, PC3): 1. Univariate IQR (Multi-Omics Only): Outliers detected by applying the 1.5 × IQR rule to the z-scores of each principal component. 2. Mahalanobis Distance (Factor Scores): Outliers identified using the Minimum Covariance Determinant (MCD)-based Mahalanobis distance on integrated factor scores. 3. Silhouette Scores (Cluster Fit): Outliers flagged based on low silhouette scores, indicating poor fit to any cluster. A sample was classified as a consensus outlier if flagged by at least two of the three methods. This process was repeated iteratively until no further consensus outliers remained.

**Supplementary Figure 4 |** Consensus Outliers Assessment Silhouette Scores

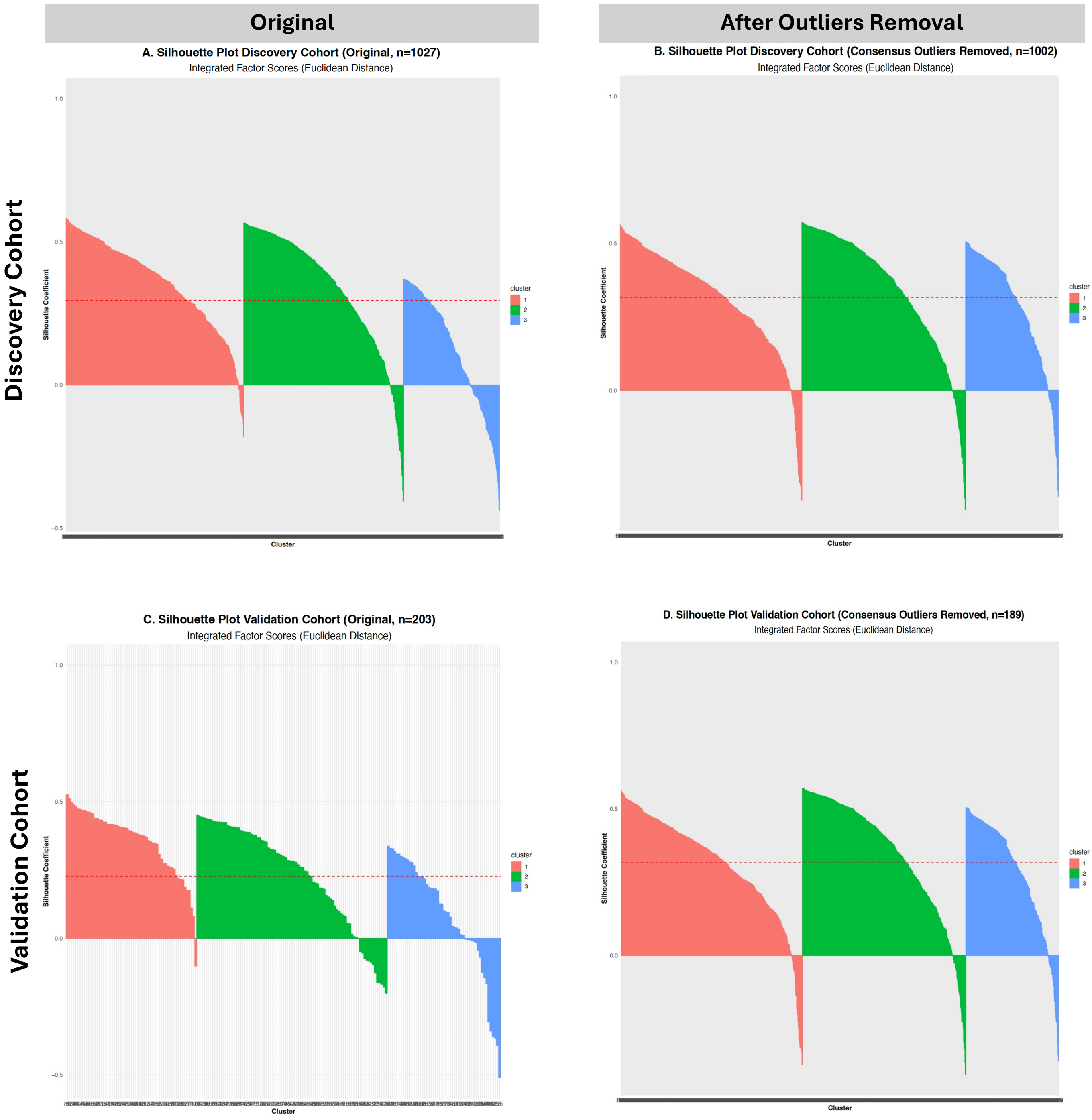


**Supplementary Figure 4:** ***Silhouette Scores and Cluster Fit Assessment.***

(A) and (C) Silhouette plots for the original datasets, assessing cluster quality based on integrated factor scores using Euclidean distance.

(B) and (D) Silhouette plots for dataset after consensus outlier removal, showing improved cluster fit. Samples with coefficients < 0.1 indicate potential outliers or poor cluster assignment.

#### **Supplementary Figure 5 |** Clusters and the HIV Reservoir

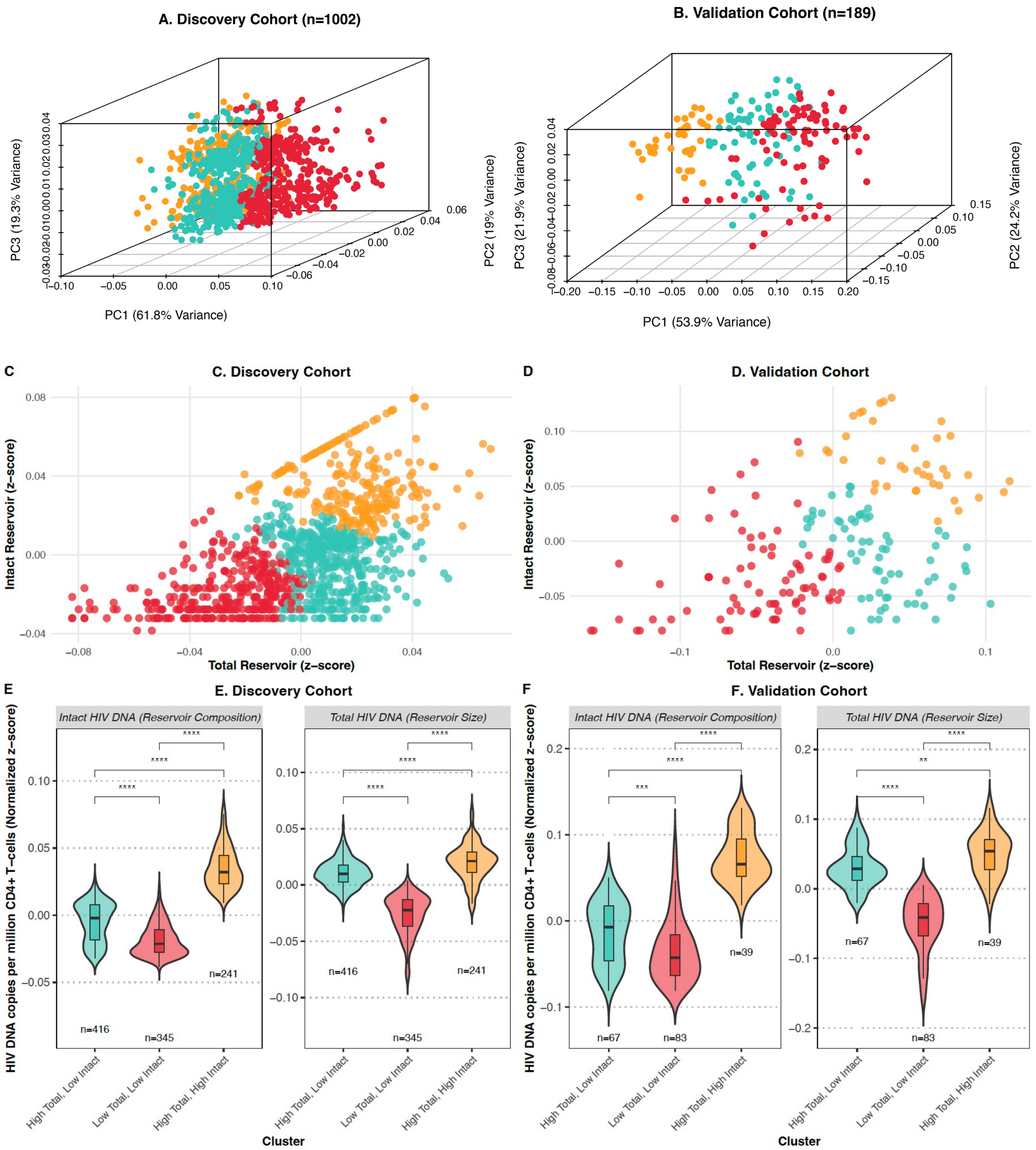


**Supplementary Figure 5: Clusters and HIV DNA viral reservoir**

(A–B) 3D scatterplots of sample factor scores from MoCluster clustering analysis (PC1, PC2, PC3) in the discovery and validation cohorts, colored by final cluster assignment ("High Total, Low Intact" (Mixed), "Low Total, Low Intact" (All Low), "High Total, High Intact" (All High)). Axes are labeled with the proportion of variance explained by each component in the discovery cohort.

(C–D) Scatterplots showing the relationship between *Total HIV DNA* (reservoir size) and *Intact HIV DNA* (reservoir composition), z-score normalized, for the discovery and validation cohorts. Each point represents one participant, colored by cluster.

(E–F) Violin plots of *Total* and *Intact HIV DNA* (z-scores) for each cluster in the discovery and validation cohorts. Distributions are overlaid with boxplots; sample sizes per cluster are indicated, and statistically significant pairwise differences between clusters were assessed using the Wilcoxon rank-sum test ($P<0.05$, adjusted), with significance levels annotated on the plots.

#### **Supplementary Figure 6 |** Linear and nonlinear visualization of clusters

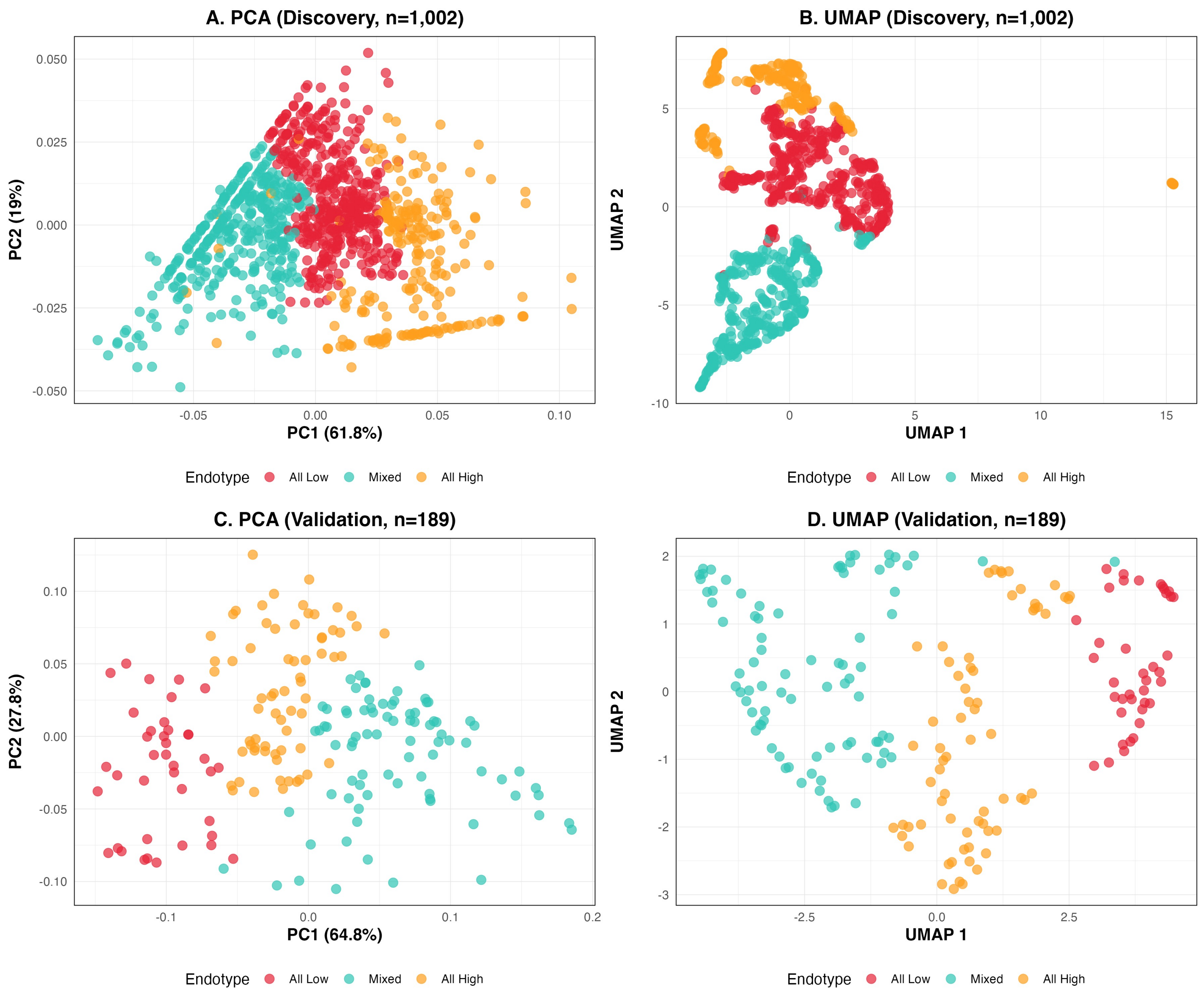


**Supplementary Figure 6. PCA and UMAP projections of the MoCluster consensus scores in the discovery and validation cohorts.** PCA projections are shown for the discovery (A) and validation (C) cohorts, with corresponding UMAP projections shown in B and D. Points represent individual participants and are coloured by endotype assignment. The UMAP projection provides a complementary nonlinear visualization and shows consistent separation of the three endotypes in both cohorts.

#### **Supplementary Figure 7 |** Bulk Transcriptomics Driver Markers and Reservoir Correlation Across Clusters

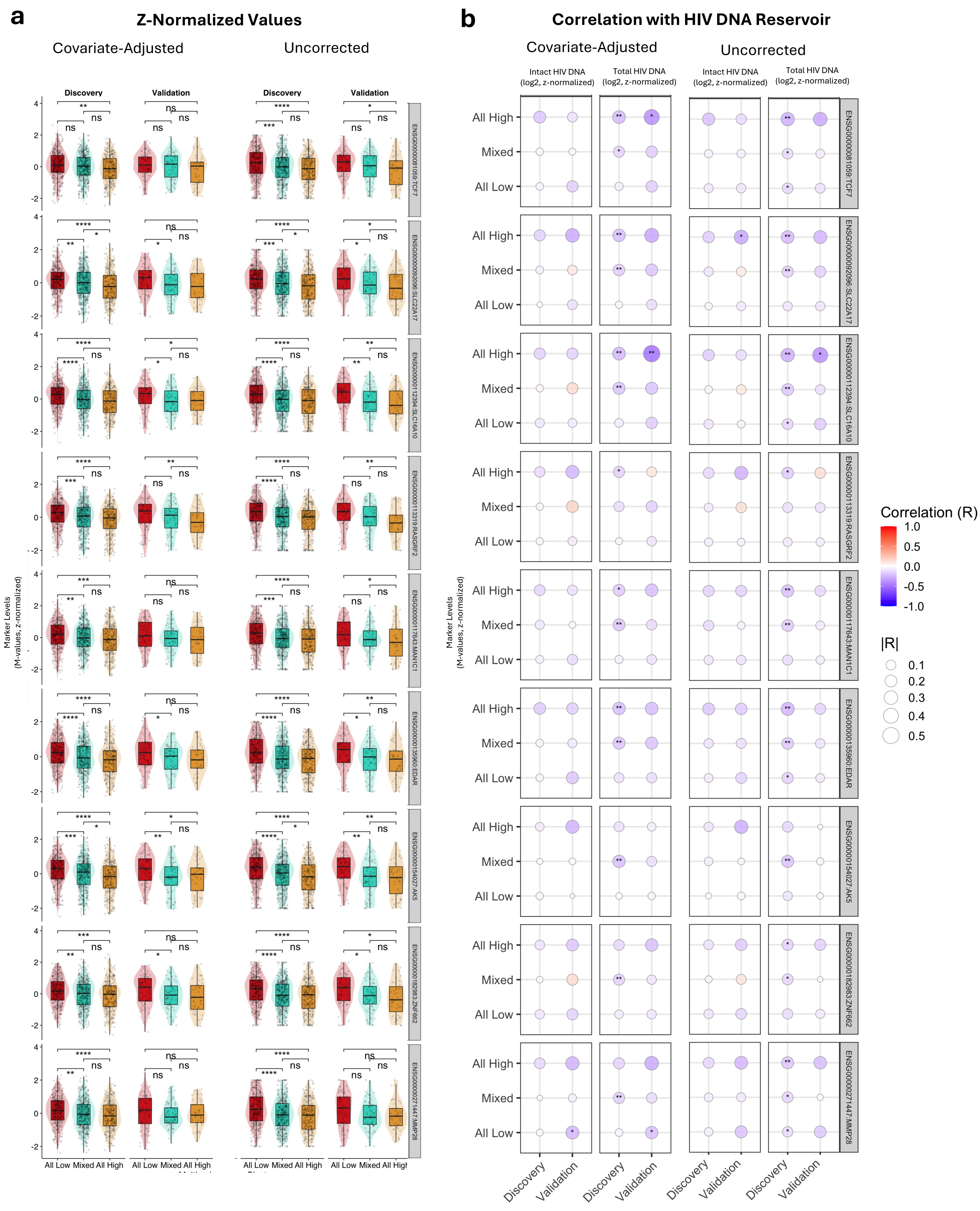


**Supplementary Figure 7. Bulk Transcriptomics Driver Markers and Reservoir Correlation Across Clusters. a,** Violin–boxplots of validated bulk transcriptomics driver markers extracted from MoCluster (VST-transformed, top 5,000 variable features by MAD) across clusters (Mixed, All Low, All High) in discovery and validation cohorts (n = 1,002 and n = 189, respectively). Left panels show age- and sex-adjusted levels; right panels show unadjusted data. Inter-cluster t-tests were used to address significance. **b,** Correlation plot of driver markers with total and intact HIV DNA (reservoir size and intactness). Left panels display partial correlations (age- and sex-adjusted); right panels show unadjusted correlations. Dot size and color intensity reflect correlation magnitude; color indicates direction. Significance is denoted by asterisks (*P < 0.05; **P < 0.01; ***P < 0.001; ****P < 0.0001).

### **Supplementary Figure 8 |** Single-Cell RNA-seq Localization of Validated Transcriptomics Markers

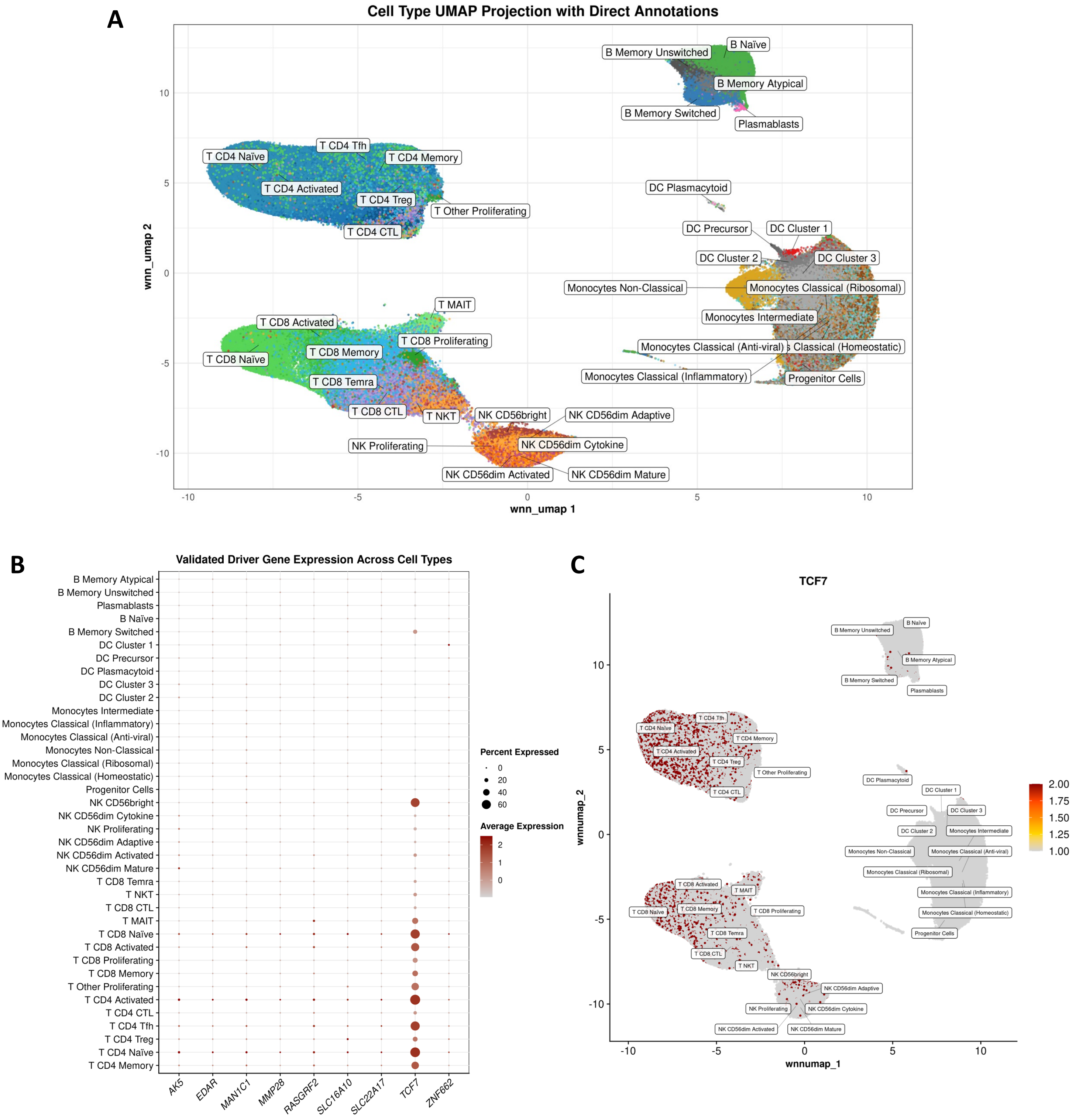


**Supplementary Figure 8 | Single-cell RNA-seq localization of validated transcriptomic markers.** Single-cell RNA-seq analysis was used to characterize the cellular expression of validated bulk-transcriptomic markers identified in the MoCluster analysis.

**(A)** UMAP projection of single cells coloured by annotated immune-cell subtype. Cell-population labels are positioned near the corresponding clusters to facilitate identification of the major lymphoid and myeloid populations.

**(B)** Dot plot showing the expression of nine validated bulk-transcriptomic markers (*AK5, EDAR, MAN1C1, MMP28, RASGRF2, SLC16A10, SLC22A17, TCF7* and *ZNF662)* across annotated cell types. Dot colour indicates average expression within each cell type, and dot size indicates the proportion of cells expressing the gene. Among these markers, *TCF7* showed the broadest expression across lymphocyte populations, with prominent expression in naïve and memory CD4$^{+}$ and CD8$^{+}$ T-cell subsets and NK-cell populations.

**(C)** UMAP projection showing single-cell *TCF7* expression, with annotated cell populations overlaid. *TCF7* expression was concentrated predominantly within T-cell and selected NK-cell populations, consistent with the distribution observed in panel B.

### **Supplementary Figure 9 |** Cell-type Localization of Validated Bulk-Transcriptomic Driver Genes at Gene-specific Resolution

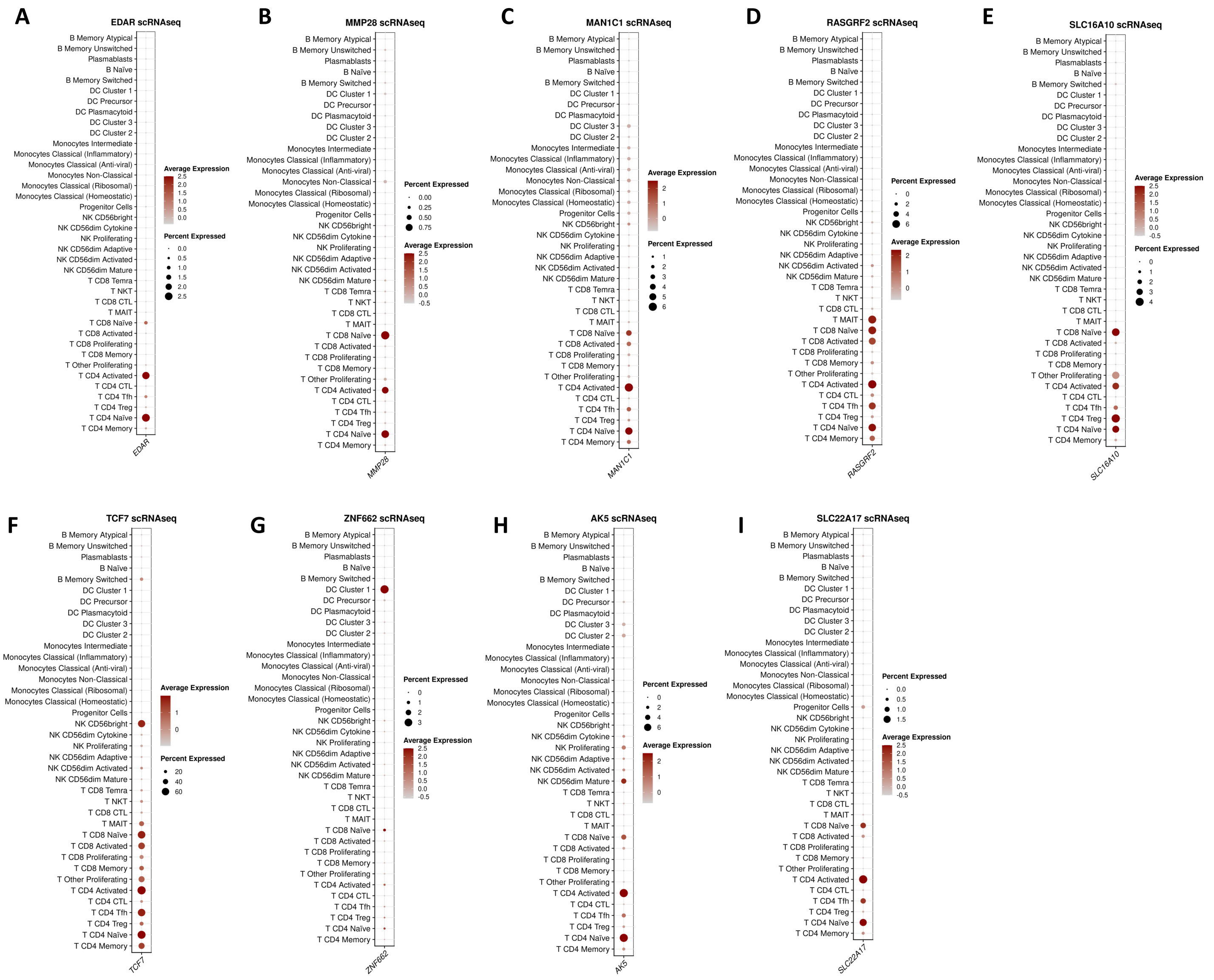


**Supplementary Figure 9 | Cell-type distribution of validated bulk-transcriptomic driver genes at gene-specific resolution.** Dot plots show the expression of nine validated bulk-PBMC transcriptomic driver genes across annotated immune-cell populations in the complementary single-cell RNA-seq dataset: **A**, *EDAR*; **B**, *MMP28*; **C**, *MAN1C1*; **D**, *RASGRF2*; **E**, *SLC16A10*; **F**, *TCF7*; **G**, *ZNF662*; **H**, *AK5*; and **I**, *SLC22A17*. Each gene is displayed using an independent expression scale to preserve cell-type-specific patterns that may be obscured in a shared-scale representation. Dot size indicates the percentage of cells expressing each gene, and colour intensity indicates scaled average expression within each cell population. *TCF7* showed the broadest lymphocyte-associated expression, whereas *AK5*, *EDAR, MMP28, MAN1C1*, *RASGRF2*, *SLC16A10* and *SLC22A17* were also most evident across T-cell populations. *ZNF662* was most prominent in dendritic cells.

**Supplementary Figure 10 |** Flow Cytometry Driver Markers and Reservoir Correlation Across Clusters

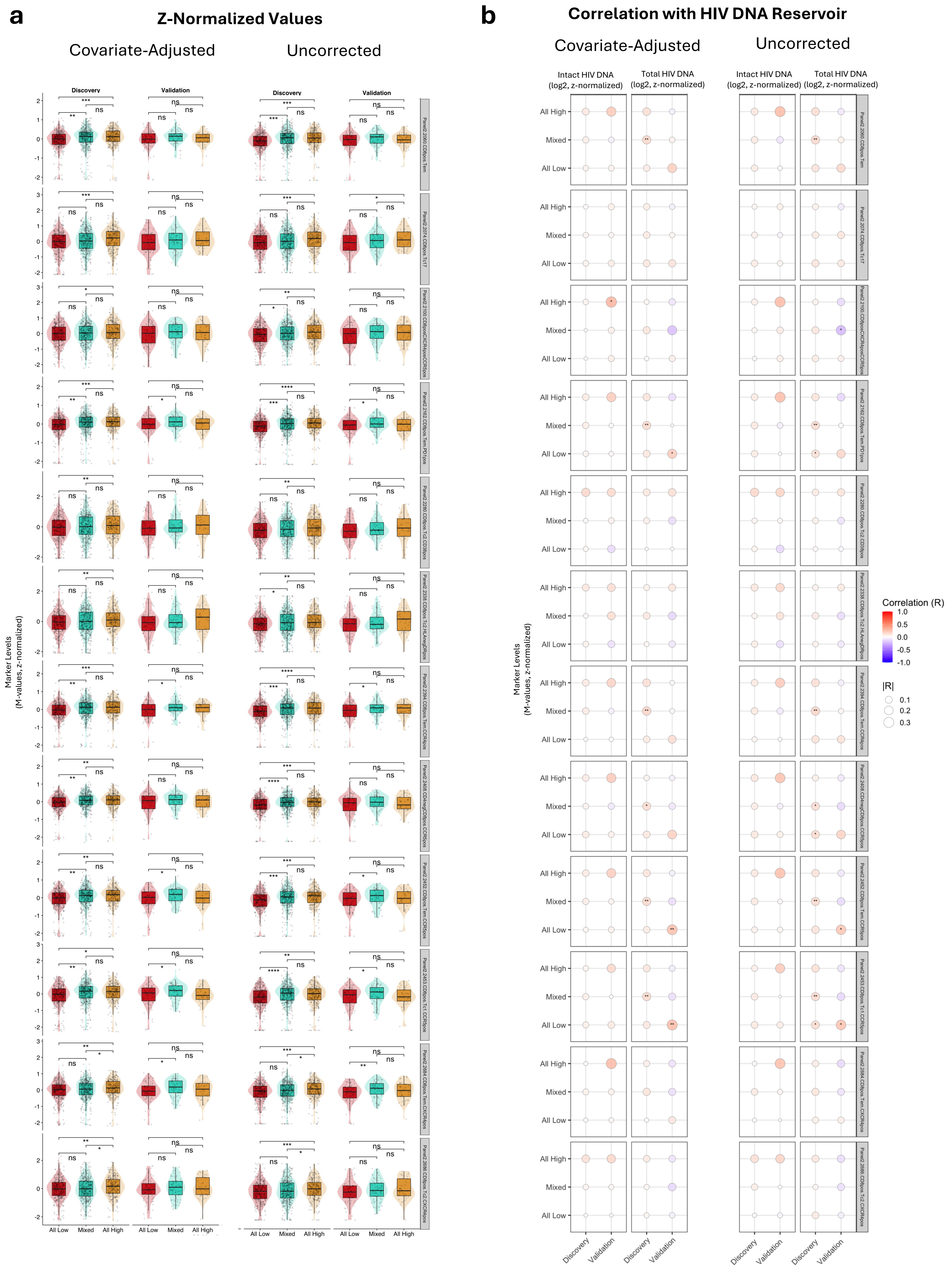


**Supplementary Figure 10. Flow Cytometry Driver Markers and Reservoir Correlation Across Clusters. a,** Boxplots of 12 validated flow cytometry driver markers (log2-transformed absolute values, identified as per-layer MoCluster drivers post-outlier removal) across clusters (Mixed, All Low, All High) in discovery and validation cohorts (n = 1,002 and n = 189, respectively). Inter-cluster t-tests indicate significance (). **b,** Correlation plot of driver markers with total and intact HIV DNA (reservoir size and intactness). Left panels show age- and sex-adjusted partial correlations; right panels display unadjusted correlations. Dot size and color intensity reflect correlation magnitude; color indicates direction. Significance denoted by: *P < 0.05; **P < 0.01; ***P < 0.001; ****P < 0.0001.

#### **Supplementary Figure 11 |** Ex Vivo Cytokines Driver Markers and Reservoir Correlation Across Clusters

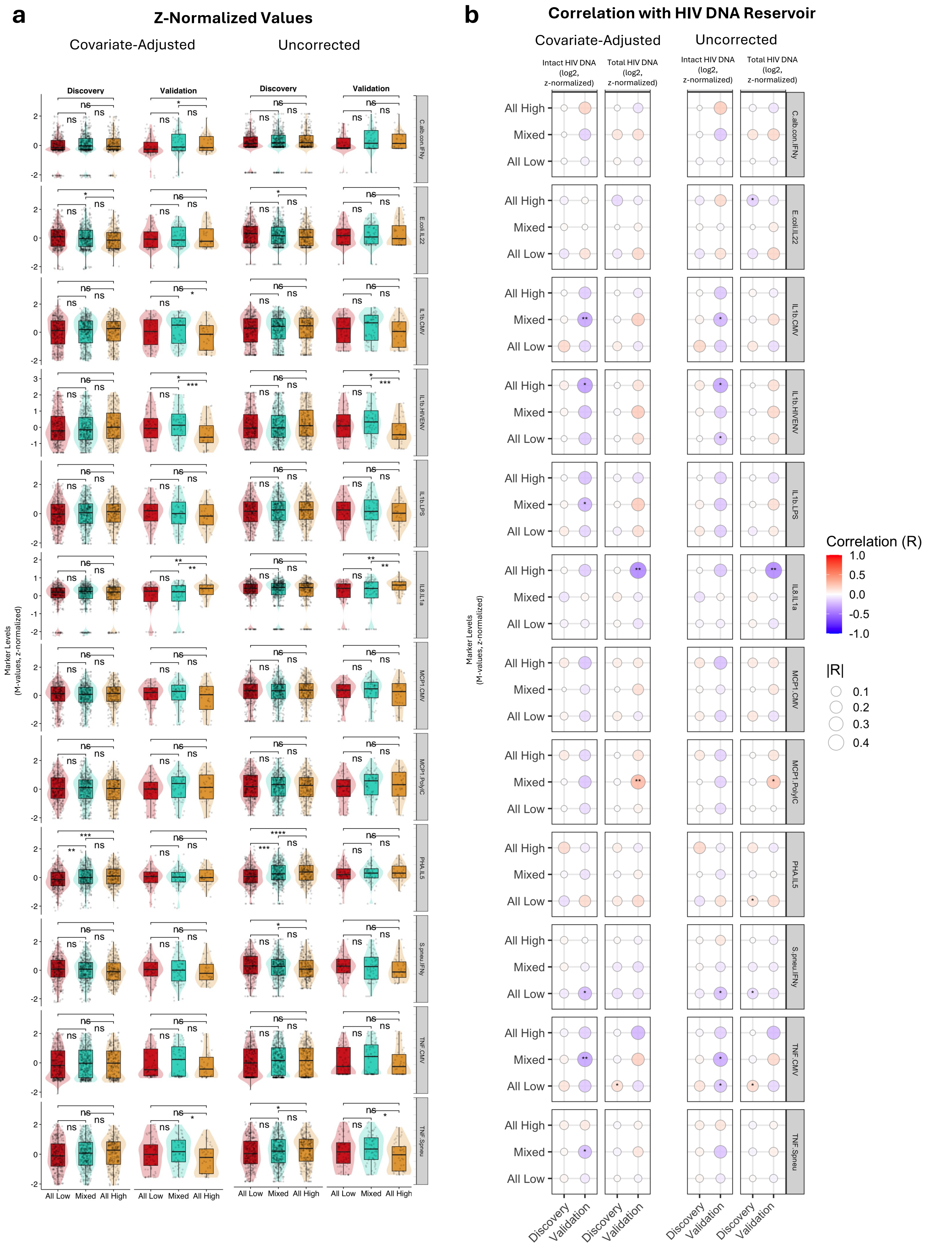


**Supplementary Figure 11. Ex Vivo Cytokines Driver Markers and Reservoir Correlation Across Clusters. a,** Boxplots of 12 validated ex vivo cytokine driver markers from MoCluster (measured after 24-hour and 7-day stimulations, see **Methods**) across clusters in discovery and validation cohorts (n = 1,002 and n = 189, respectively). Inter-cluster t-tests assessed significance. **b,** Correlation of driver markers with total and intact HIV DNA (reservoir size and intactness). Left panels show age- and sex-adjusted partial correlations; right panels display unadjusted correlations. Dot size and color intensity reflect correlation magnitude; color indicates direction. Significance denoted by: *P < 0.05; **P < 0.01; ***P < 0.001; ****P < 0.0001.

**Supplementary Figure 12 |** DNA Methylation Driver Markers and Reservoir Correlation Across Clusters

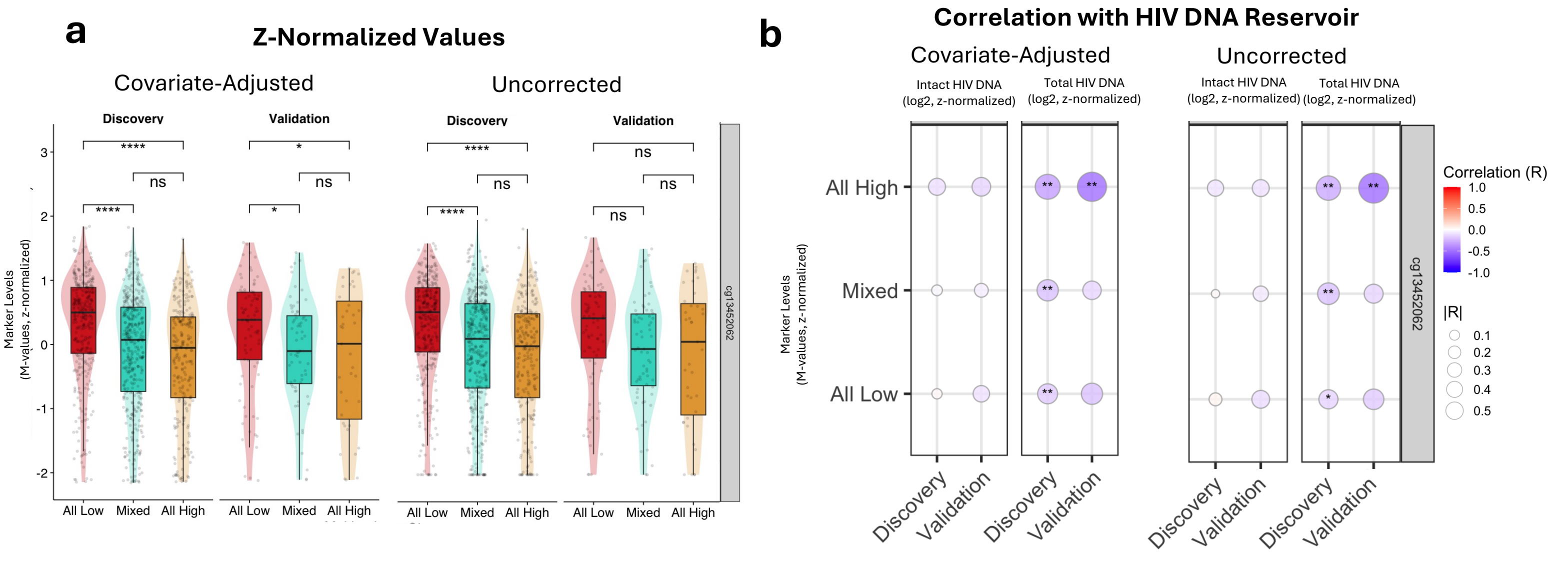


**Supplementary Figure 12. DNA Methylation Driver Markers Across Clusters and Reservoir Correlation Across Clusters**
**a,** Boxplots of validated DNA methylation driver markers (M-values shifted for positivity, top 1% variable CpG sites by MAD) across clusters (Mixed, All Low, All High) in discovery and validation cohorts (n = 1,002 and n = 189, respectively). Inter-cluster t-tests indicate significance. **b,** Correlation plot of driver markers with total and intact HIV DNA (reservoir size and intactness). Left panels show age- and sex-adjusted partial correlations; right panels display unadjusted correlations. Dot size and color intensity reflect correlation magnitude; color indicates direction. Significance denoted by: *P < 0.05; **P < 0.01; ***P < 0.001; ****P < 0.0001.

### **Supplementary Figure 13 |** Bulk and Single-Cell RNA-seq Cell Type Expression of *IFI44L*

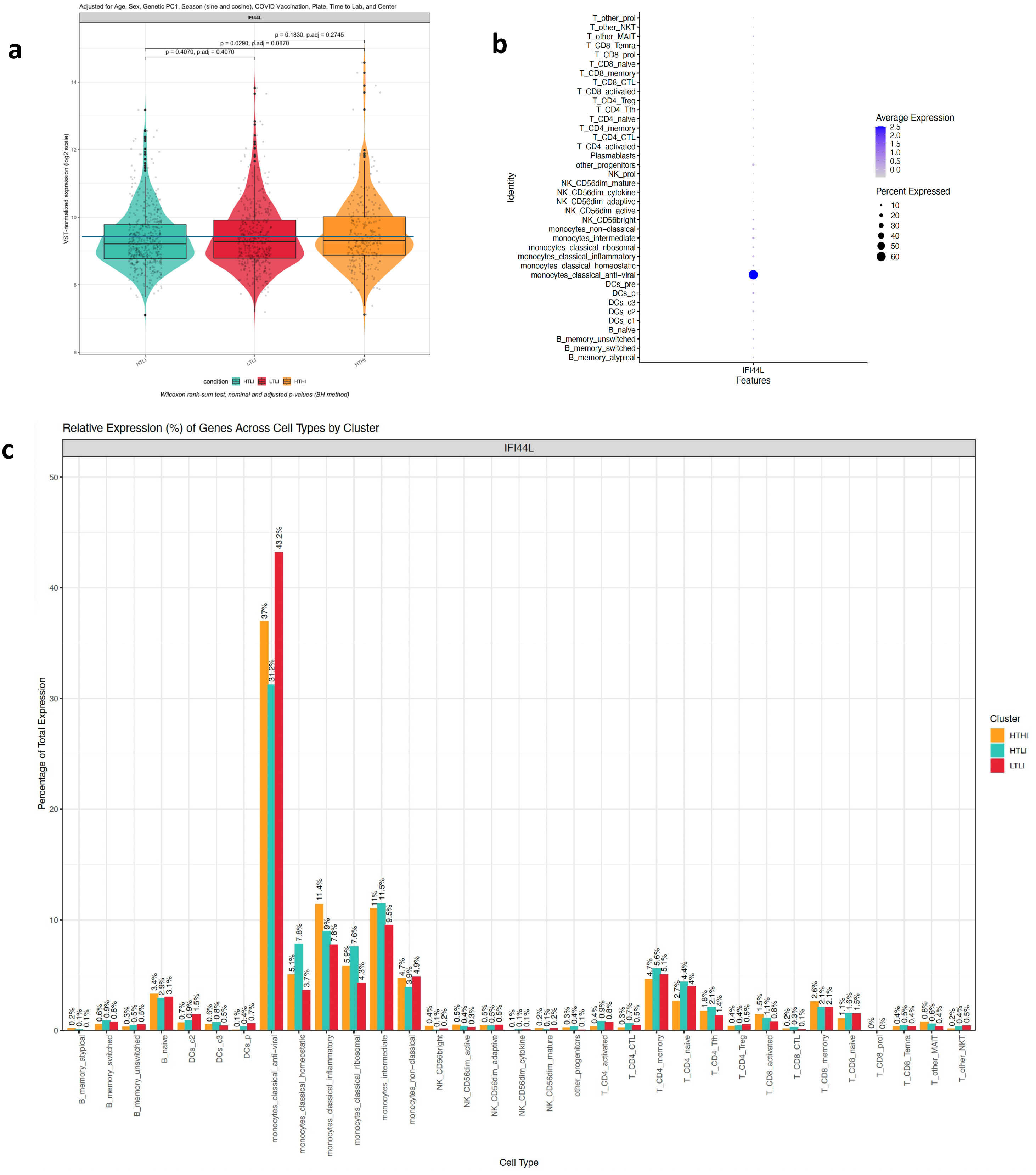


**Supplementary Figure 13 - Bulk and Single-Cell RNA-seq Cell Type Expression of *IFI44L***

(A) Violin plot of *IFI44L* bulk RNA-seq variance-stabilized transformed (VST) expression across clusters after adjusting for Age, Sex, Genetic PC1, Season (sine and cosine), COVID Vaccination, Plate, Time to Lab, and Center (see **Methods**). This visualization highlights that *IFI44L* exhibits trends of higher expression in All High and All Low clusters compared to Mixed.

(B) Dot plot where each row represents a cell type. The color intensity indicates the average expression level of each gene within a cell type, while the dot size reflects the percentage of cells expressing that gene. This visualization highlights that *IFI44L* exhibits notably higher expression in classical antiviral monocytes compared to other cell types.

(C) Stacked bar plots summarizing the expression patterns of *IFI44L* across cell types and clusters. The barplot is divided by cell type, with segment heights representing the proportion of cells expressing that gene within each cell type, relative to the total number of expressing cells for the gene. Each bar is further colored by cluster, allowing direct comparison of how gene expression is distributed across both cell types and clusters.

**Supplementary Figure 14 | ** Proteomics Driver Markers and Reservoir Correlation Across Clusters

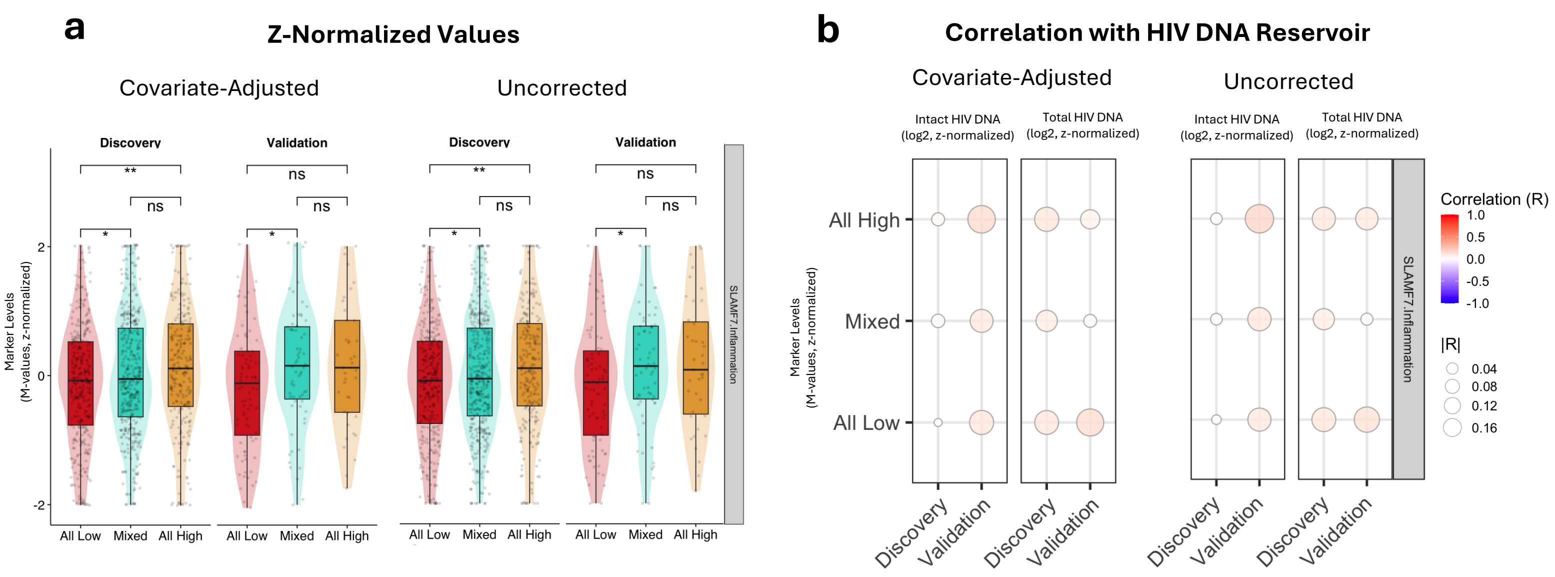


**Supplementary Figure 14. Proteomics Driver Markers and Reservoir Correlation Across Clusters.**
**a,** Boxplots of validated proteomics driver markers (log2-transformed NPX values) across clusters (Mixed, All Low, All High) in discovery and validation cohorts (n = 1,002 and n = 189, respectively). Inter-cluster t-tests indicate significance. **b,** Correlation plot of driver markers with total and intact HIV DNA (reservoir size and intactness). Left panels show age- and sex-adjusted partial correlations; right panels display unadjusted correlations. Dot size and color intensity reflect correlation magnitude; color indicates direction. Significance denoted by: *P < 0.05; **P < 0.01; ***P < 0.001; ****P < 0.0001.

**Supplementary Figure 15 |** Classification Models Performance In Held-Out Test Set

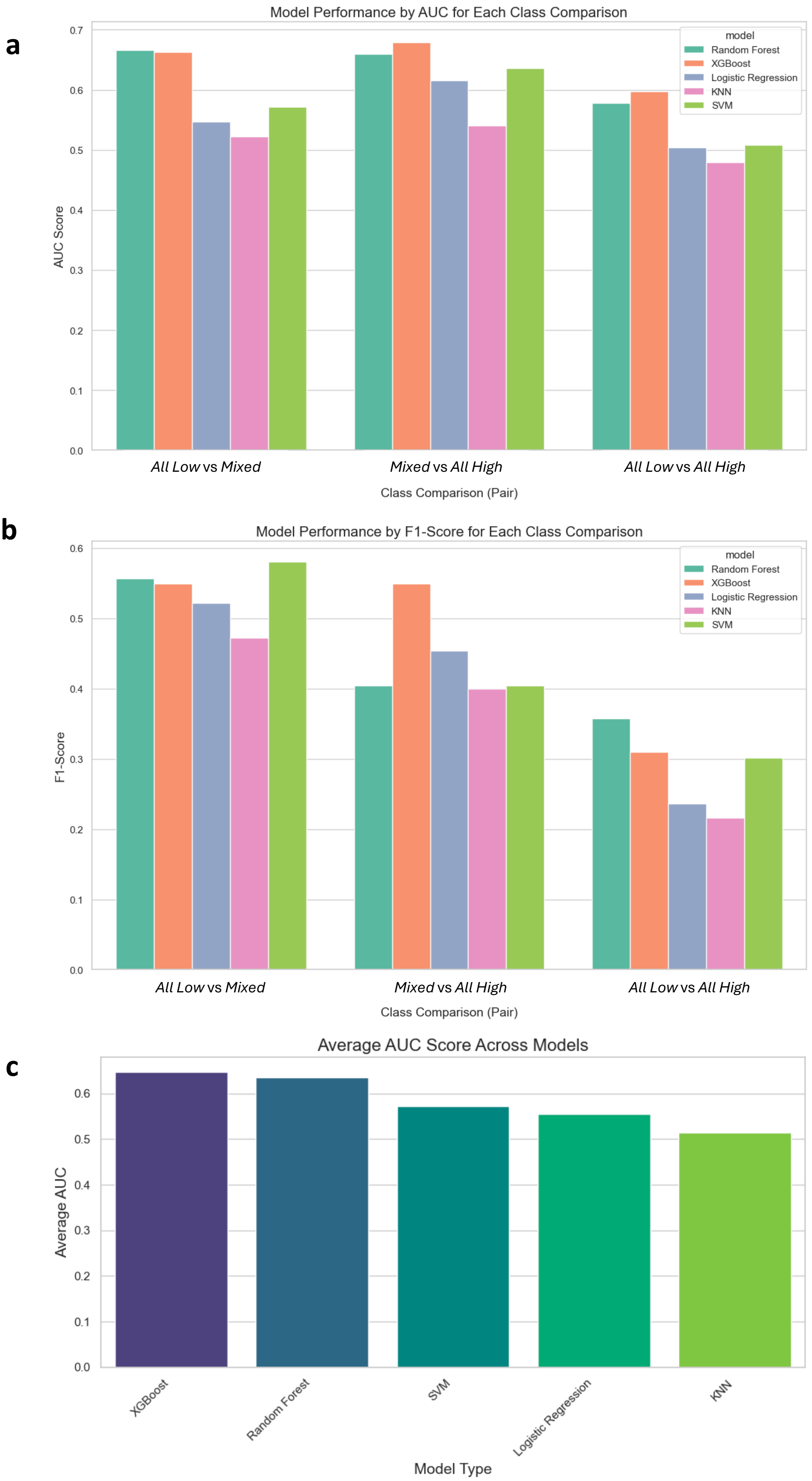


**Supplementary Figure 15 - Binary Classification Model Performance Across Clusters In Held-Out Test Set**
This figure summarizes the performance of binary classification models trained to distinguish between HIV reservoir clusters, using multiple evaluation metrics commonly applied in machine learning.
(A) Area Under the Receiver Operating Characteristic Curve (AUC) scores in the held-out test set for each binary classification model across different cluster comparisons. The AUC quantifies the model's ability to discriminate between classes, with values closer to 1 indicating better performance.
(B) F1-scores for the same models. The F1-score is the harmonic mean of precision and recall in the held-out test set, providing a balanced measure of model accuracy that is especially informative when class distributions are imbalanced.
(C) Presents the AUC score obtained from grid search cross-validation across all binary classification models, reflecting the models' generalizability and stability under different hyperparameter configurations.
Together, these panels provide a comprehensive overview of model performance, highlighting both discrimination ability (AUC) and balanced accuracy (F1-score) for each cluster comparison.

**Supplementary Figure 16 |** SHAP Interaction Heatmaps Across Binary Classification Models

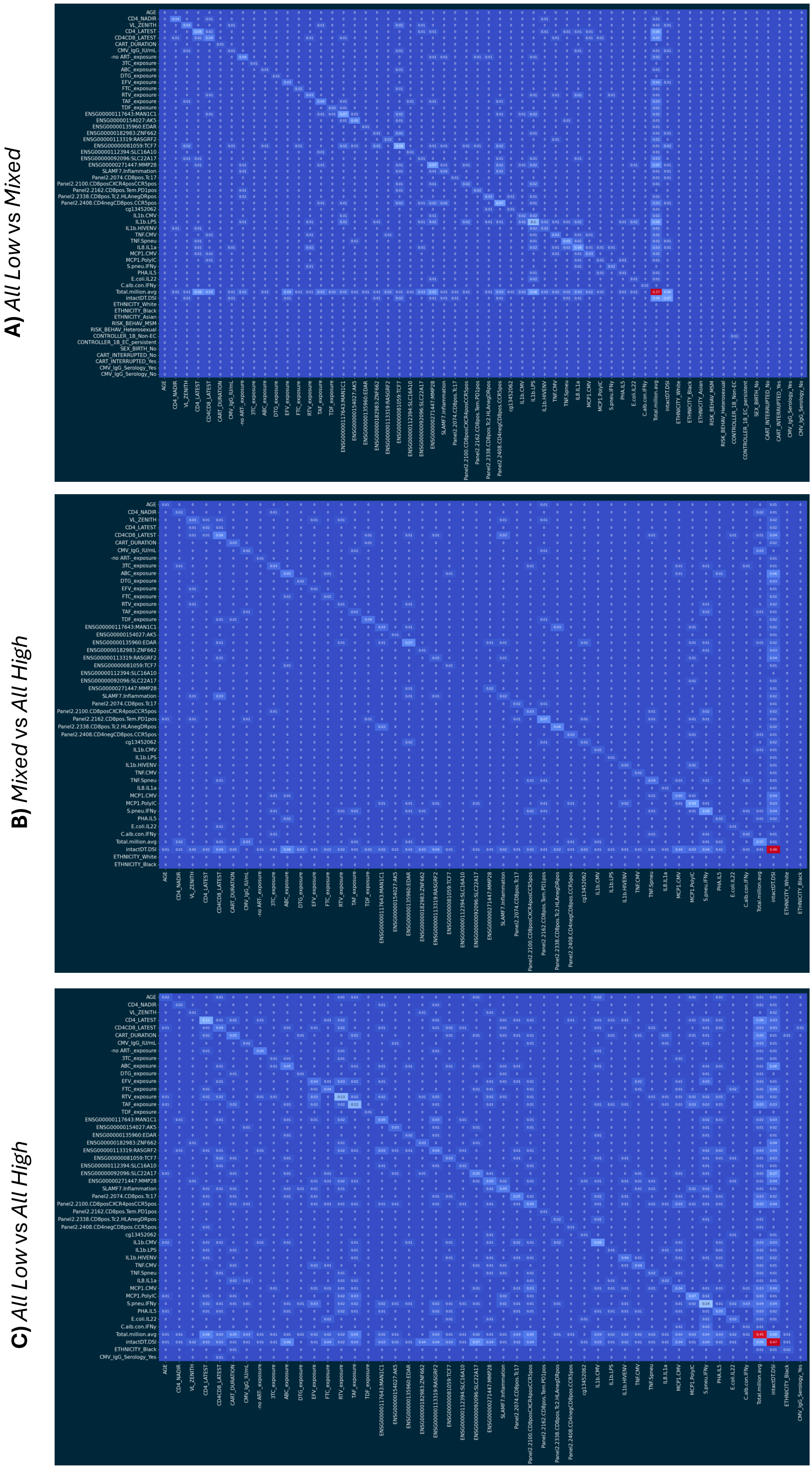


**Supplementary Figure 16. SHAP interaction heatmaps across binary classification models..** Panels **a–c** show SHAP dependence and interaction heatmaps for XGBoost classifiers distinguishing All Low vs Mixed (**a**), Mixed vs All High (**b**), and All Low vs All High (**c**) clusters. The diagonal represents the marginal contribution of individual features, while the off-diagonal entries capture pairwise interactions, with warmer colors indicating stronger influence on model predictions. Unlike correlation heatmaps, which depict statistical associations between features, SHAP values quantify model-based contributions, reflecting how features individually or jointly drive classification across clusters.

**Supplementary Figure 17 |** Single-cell localization of differentially expressed genes identified by bulk PBMC RNA-seq

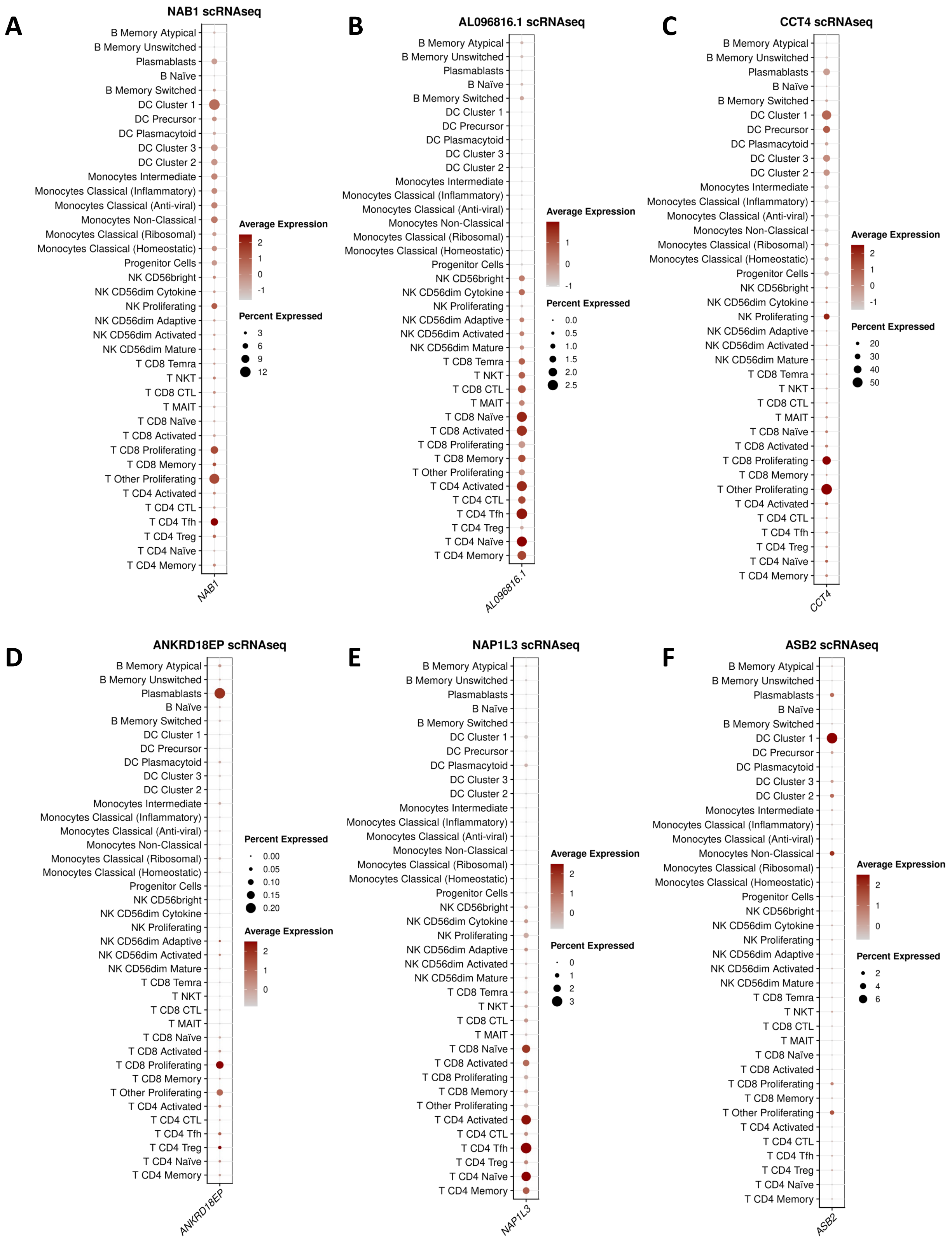


**Supplementary Figure 17 | Single-cell localization of differentially expressed genes identified by bulk PBMC RNA-seq.** Dot plots show the expression of selected differentially expressed genes across annotated peripheral blood mononuclear cell populations in the complementary single-cell RNA-seq dataset: **A**, *NAB1*; **B**, *AL096816.1*; **C**, *CCT4*; **D**, *ANKRD18EP*; **E**, *NAP1L3*; and **F**, *ASB2*. Dot colour indicates average expression within each cell type, and dot size indicates the proportion of cells expressing the gene.

#### **Supplementary Figure 18 |** Single-cell localization of leading-edge genes from bulk PBMC pathway-enrichment analyses

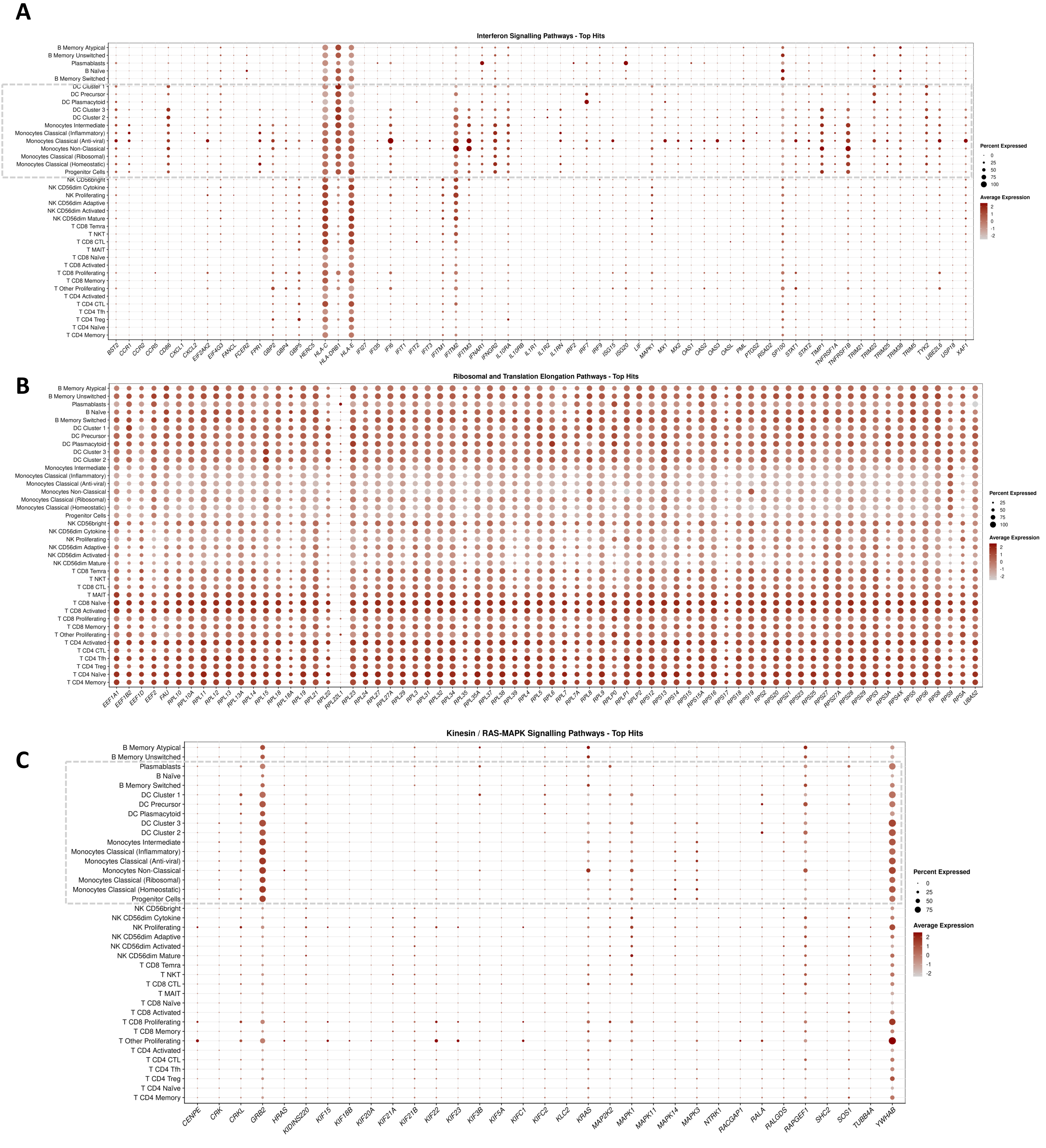


**Supplementary Figure 18 | Single-cell localization of leading-edge genes from bulk PBMC pathway-enrichment analyses.** Dot plots show the expression of genes contributing to selected pathway-enrichment signals across annotated peripheral blood mononuclear cell populations in the complementary single-cell RNA-seq dataset. **A**, interferon-signalling pathways; **B**, ribosomal and translation-elongation pathways; and **C**, kinesin- and RAS–MAPK-associated pathways. Dot size represents the percentage of cells expressing each gene, and colour intensity represents scaled average expression within each cell population. Interferon-associated genes and Kinesin- and RAS–MAPK-associated genes showed prominent expression across myeloid populations, particularly monocyte and dendritic-cell subsets, while also extending to selected lymphoid populations. Ribosomal and translation-elongation genes were broadly expressed across PBMC populations, consistent with a non-cell-restricted transcriptional programme. These patterns provide cellular context for the bulk pathway signals but do not assign pathway enrichment exclusively to a single cell population.

#### **Supplementary Figure 19 |** Ex-vivo Cytokine Production Capacity (7 days stimulation)

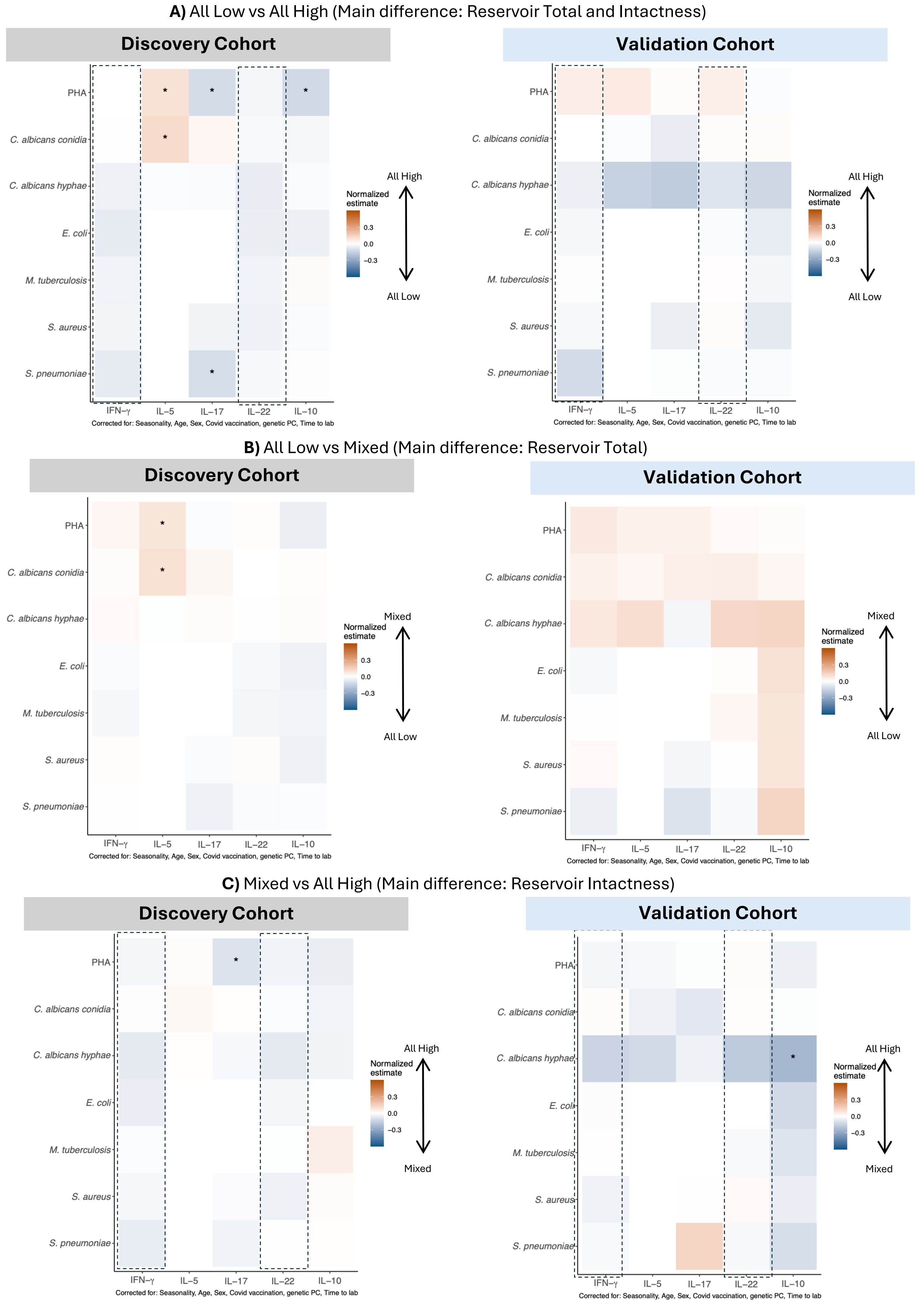


**Supplementary Figure 19.** ***Differential ex vivo cytokine production capacity across clusters after 7-day stimulation.*** Results from 7-day stimulation assays in discovery (left panels) and validation (right panels) cohorts, comparing cytokine production between: **(A)** All Low vs. Mixed (main difference: reservoir total), **(B)** Mixed vs. All High (main difference: reservoir intactness), and **(C)** All Low vs. All High (main difference: reservoir total and intactness). Cytokine production was assessed under multiple microbial and mitogenic stimuli (see Methods), and values were adjusted for confounders. Color scale represents normalized effect estimates from rank-based regression models, with higher production shown in red and lower in blue; significance is denoted as *"***" for P ≤ 0.0005, "**" for P ≤ 0.005, and "*"* for *P* ≤ 0.05.

#### **Supplementary Figure 20 |** Ex-vivo Cytokine Production Capacity (24 hours)

**A)** All Low vs All High (Main difference: Reservoir Total and Intactness)

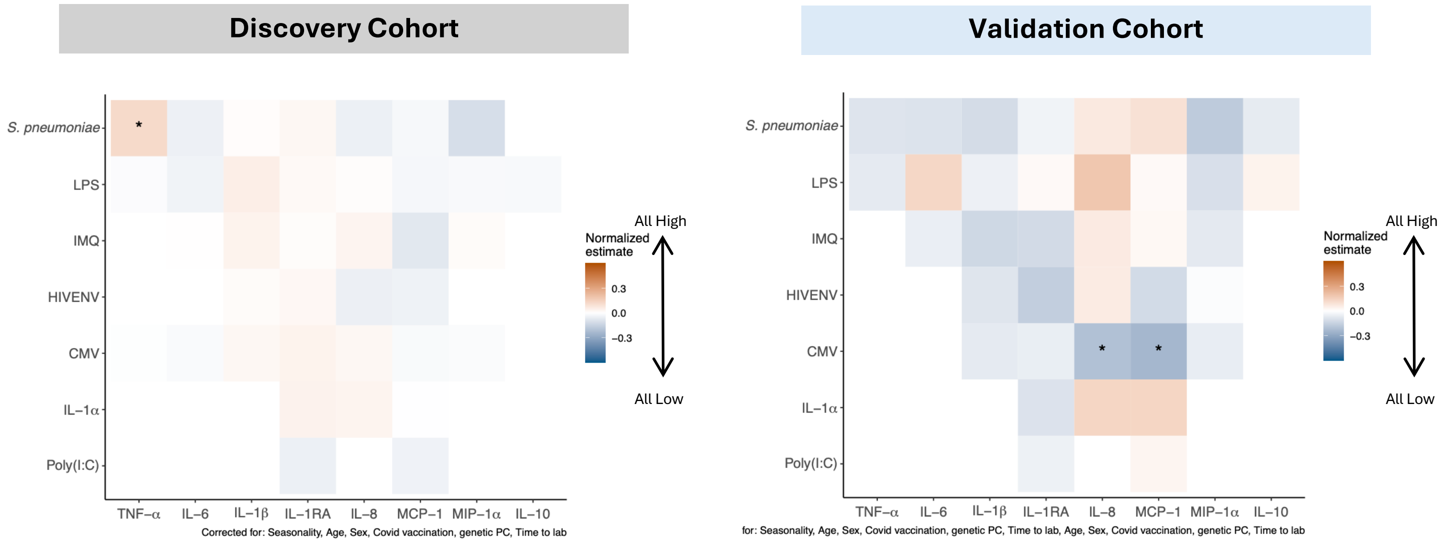


**B)** All Low vs Mixed (Main difference: Reservoir Total)

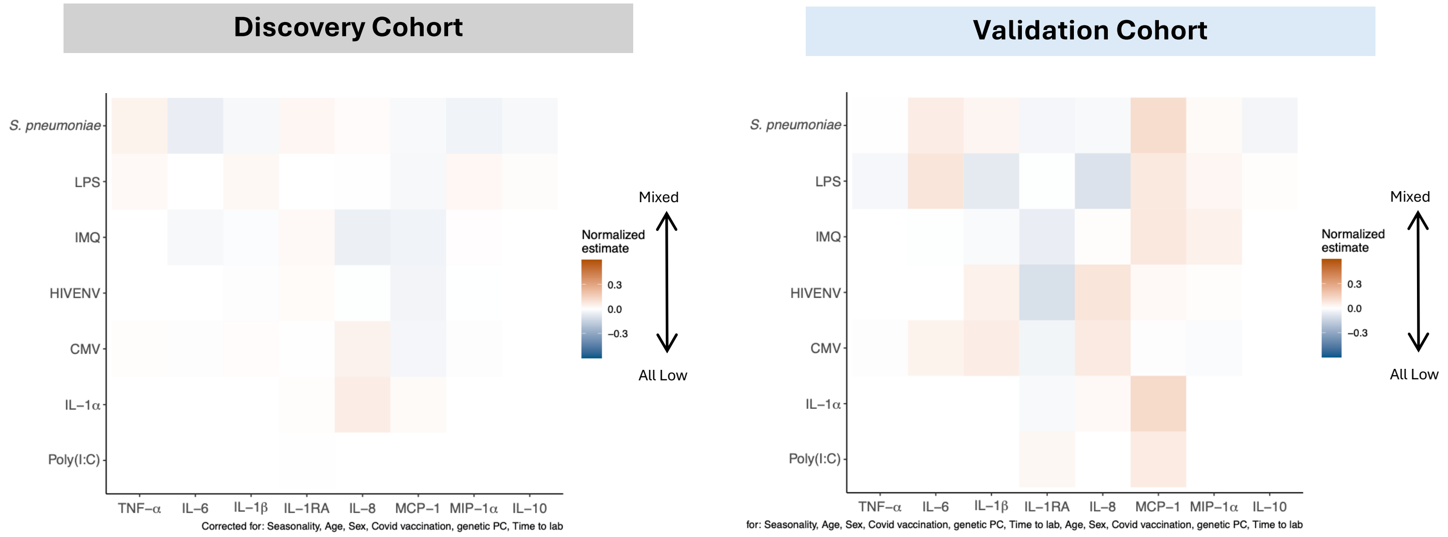


**C)** Mixed vs All High (Main difference: Reservoir Intactness)

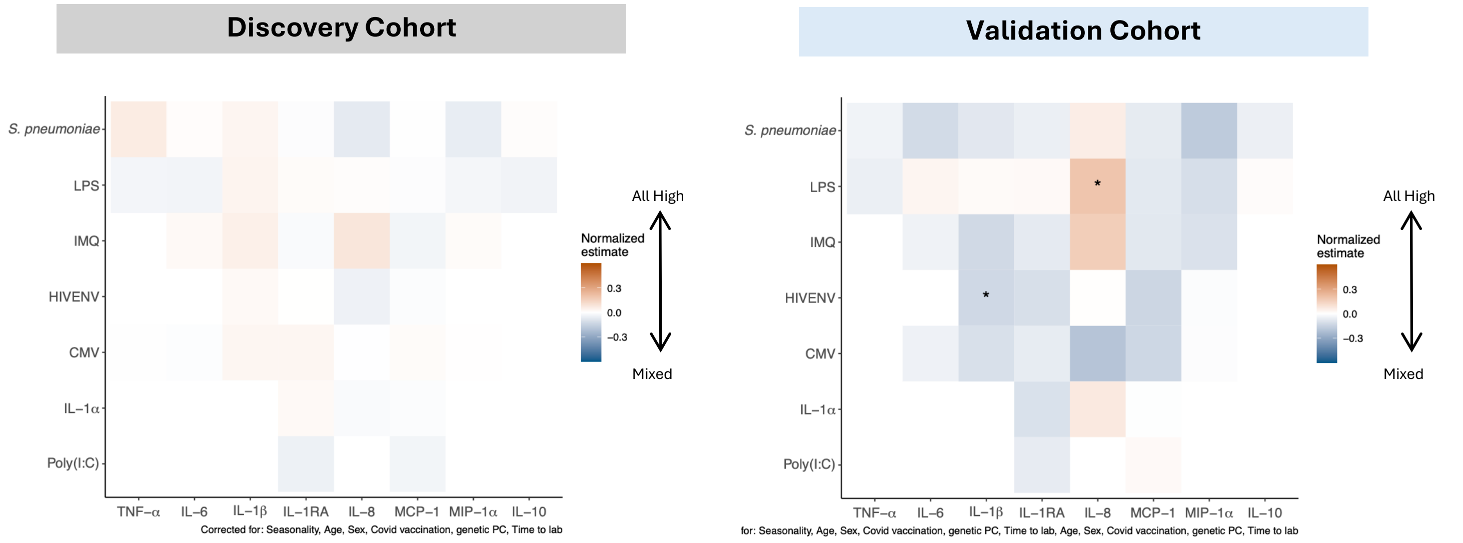


**Supplementary Figure 20.** ***Differential ex vivo cytokine production capacity across clusters after 24-hour stimulation.*** Results from 24-hour stimulation assays in discovery (left panels) and validation (right panels) cohorts, comparing cytokine production between: **(A)** All Low vs. Mixed (main difference: reservoir total), **(B)** Mixed vs. All High (main difference: reservoir intactness), and **(C)** All Low vs. All High (main difference: reservoir total and intactness). Cytokine production was assessed under multiple microbial and mitogenic stimuli (see Methods), and values were adjusted for confounders. Color scale represents normalized effect estimates from rank-based regression models, with higher production shown in red and lower in blue; significance is denoted as ”***" *for P ≤ 0.0005, ”**" for P ≤ 0.005, and ”*"* for P ≤ 0.05.

#### **Supplementary Figure 21 |** Comorbidity Associations Across Immune-Reservoir Clusters

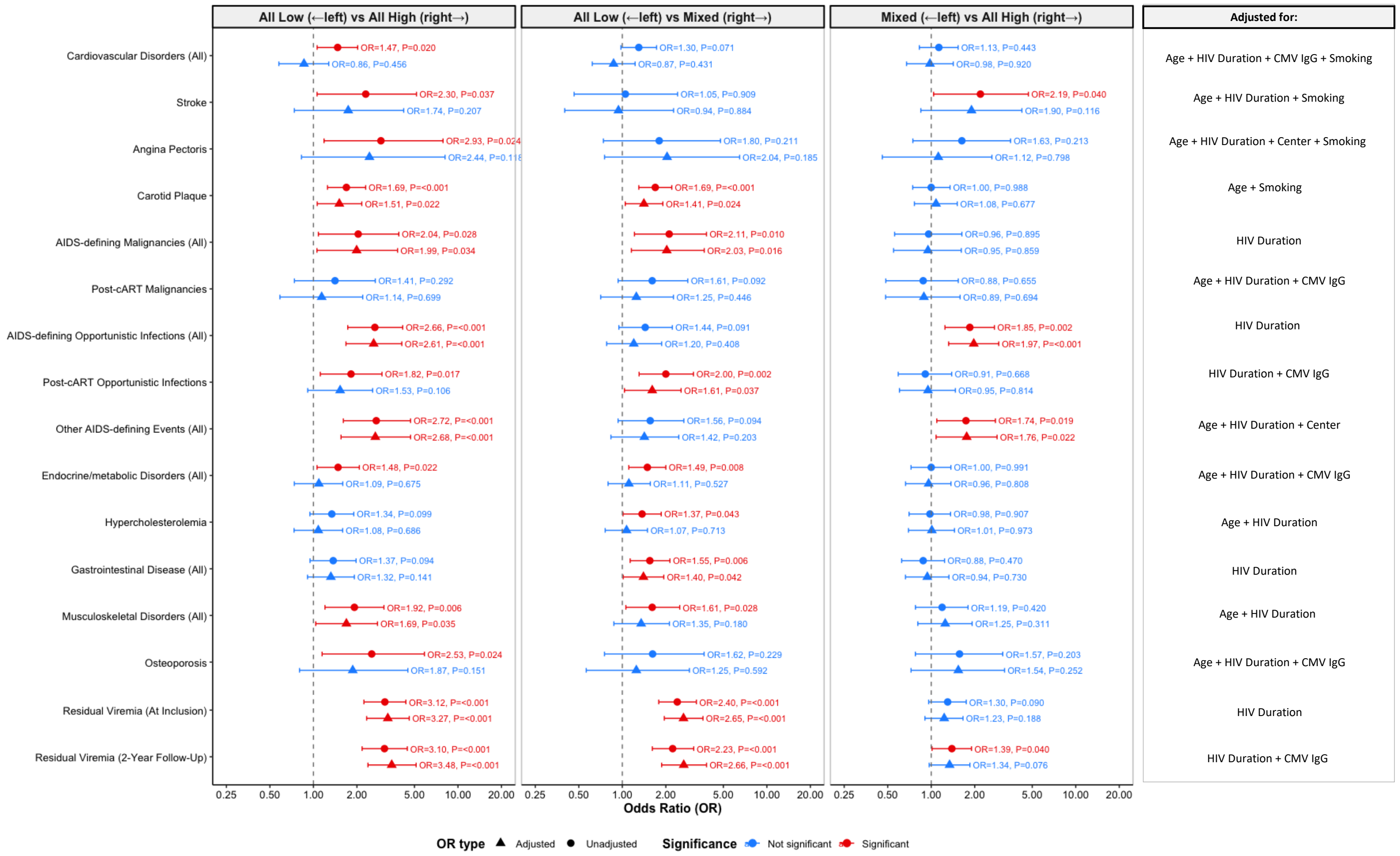


**Supplementary Figure 21.** ***Comorbidity Associations Across Immune-Reservoir Clusters*****.**
Unadjusted (circles) and adjusted (triangles) odds ratios (ORs) with 95% confidence intervals are shown for three pairwise comparisons of cluster membership (*All Low* vs. *All High*; *All Low* vs. *Mixed*; *Mixed* vs. *All High*). Analyses were performed using logistic regression with covariates selected per outcome using a change-in-estimate approach; the covariates included in each final adjusted model are indicated to the right of the plots. Significant associations (p<0.05) are shown in red. ORs >1 indicate increased odds of the comorbidity in the right-hand cluster. Rare outcomes were excluded due to model instability.

#### **Supplementary Figure 22 |** Confounders Assessment in Flow Cytometry Layer

##### **A)** Flow Cytometry: Absolute Counts

**Before Correction**

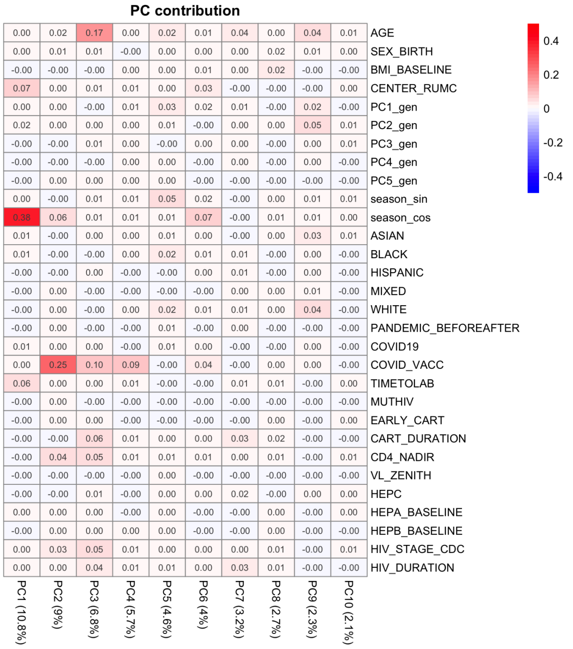


**After Confounders Correction**

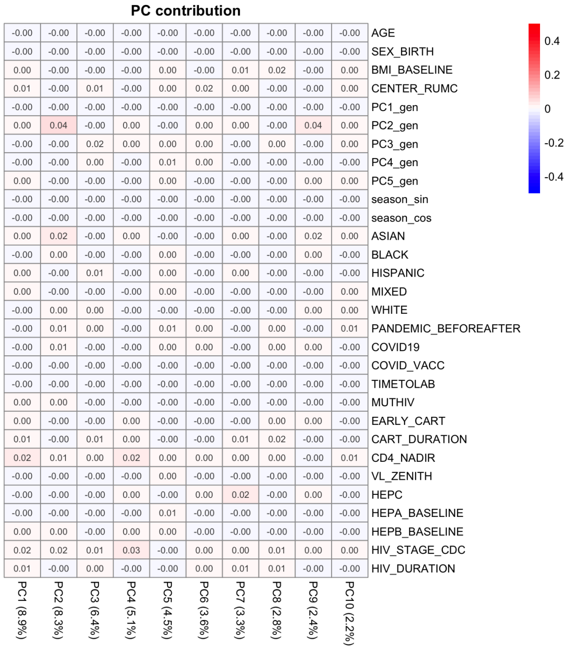


##### **B)** Flow Cytometry: Percentages

**Before Correction**

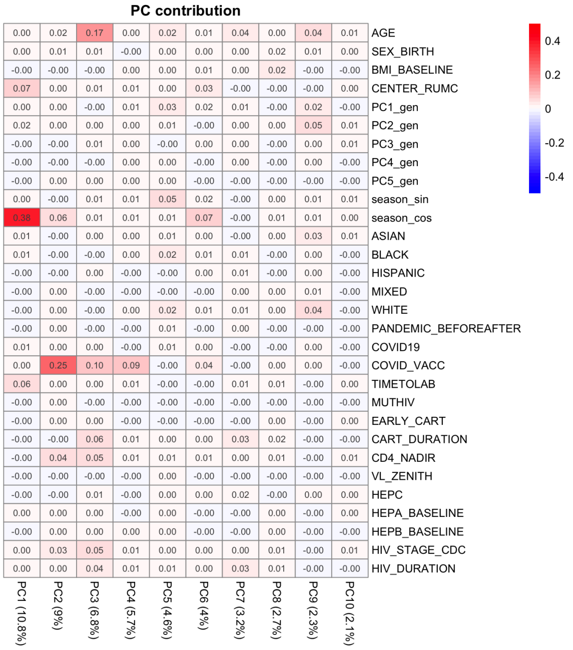


**After Confounders Correction**

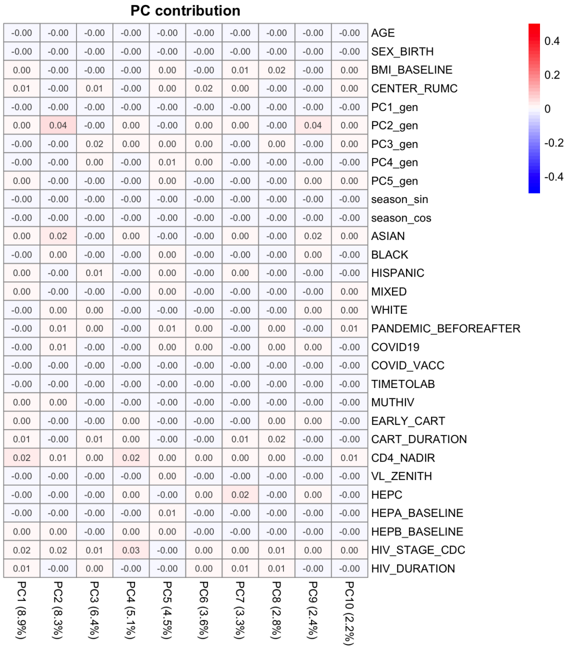


**Supplementary Figure 22. Assessment of confounding factors in the flow cytometry data.**
Heatmaps depict the associations between the first 10 principal components (PCs) of flow cytometry data and candidate confounders for (A) absolute cell counts and (B) cell population percentages. Left panels show unadjusted beta coefficients from linear regression models, while right panels present associations after adjustment for key covariates: season of sample collection, age, sex, the first genetic principal component, COVID-19 vaccination status, and sample processing time.
Color intensity corresponds to the magnitude and direction of beta estimates. Confounders were defined as variables increasing beta coefficients by >10% in a multi-step approach. Following adjustment, no confounder exceeded this threshold, demonstrating effective control of potential confounding effects in both data representations.

#### **Supplementary Figure 23 |** Confounders Assessment in Bulk Transcriptomics Layer

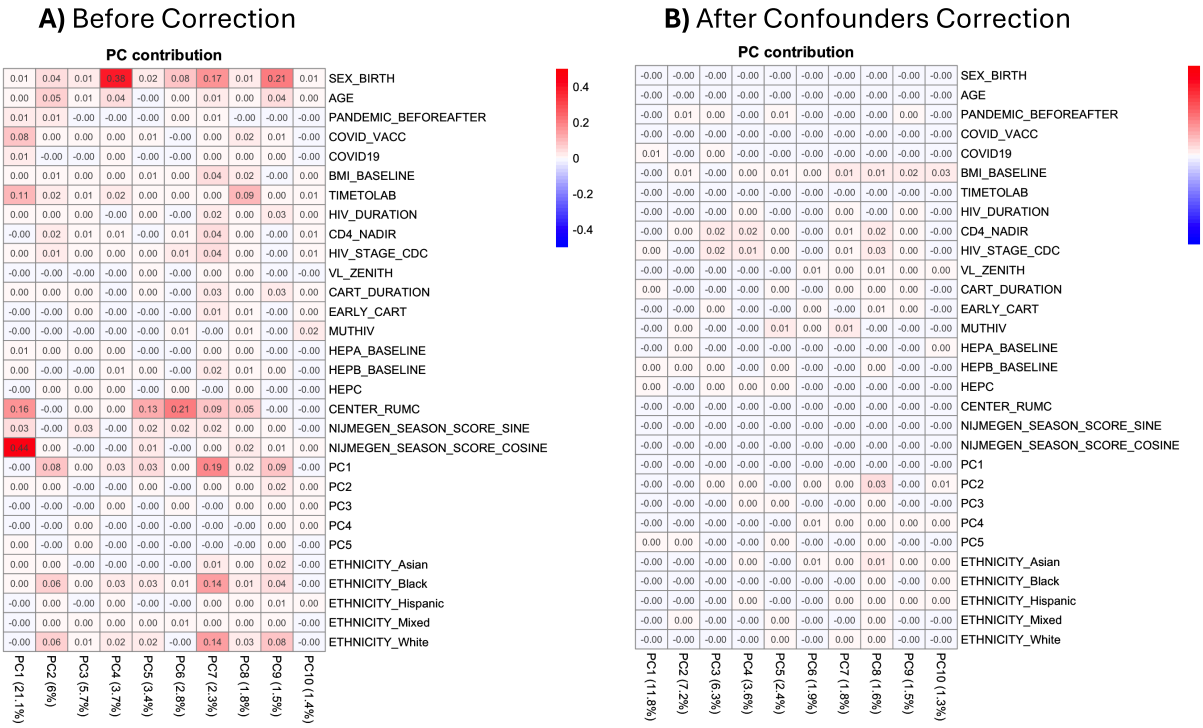


**Supplementary Figure 23.** ***Assessment of confounding factors in the bulk transcriptomics data.***
Heatmaps illustrate the associations between the first 10 principal components (PCs) of bulk RNA-seq data and potential confounders. The left panel shows beta coefficients from linear regression models prior to adjustment, while the right panel displays results following correction for key covariates: season of sample collection, age, sex, the first genetic principal component, COVID-19 vaccination status, and sample processing time. Color intensity represents the magnitude and direction of beta estimates. Confounders were defined as variables inducing >10% change in beta coefficients through a multi-step approach. Post-adjustment, no confounder surpassed this threshold, indicating effective mitigation of confounding influences.

### **Supplementary Figure 24 |** Confounders Assessment in Ex Vivo Cytokines Layer

#### **A)** Ex Vivo Cytokines (7-day Stimulation)

##### Before Correction

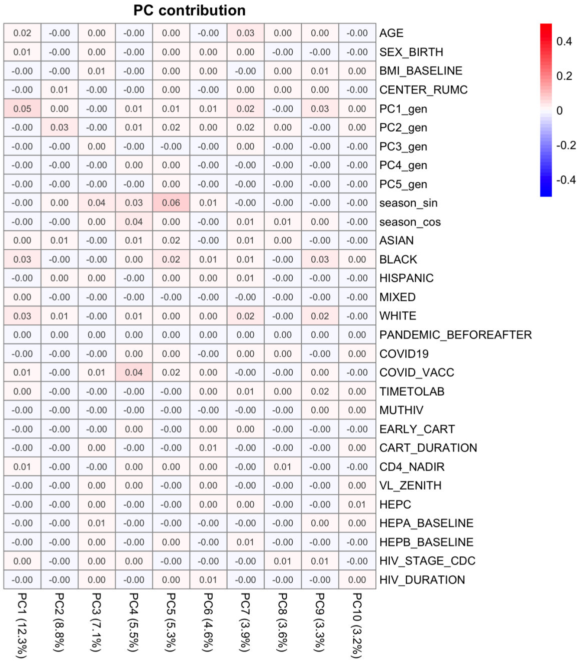


##### After Confounders Correction

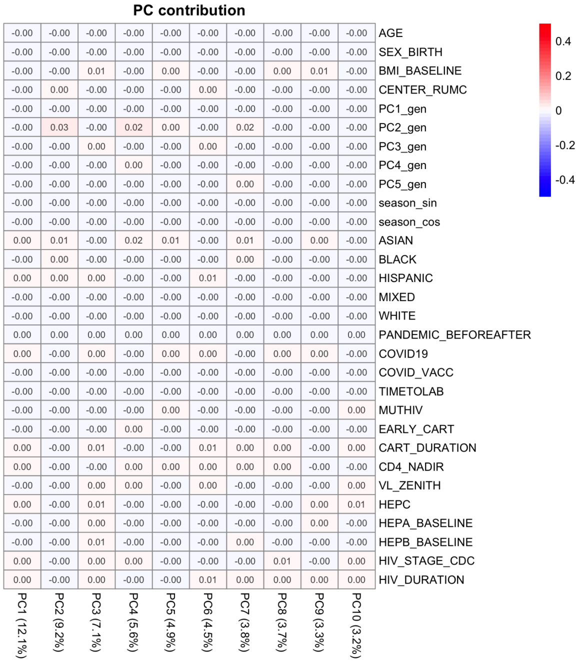


#### **B)** Ex Vivo Cytokines (24-hour Stimulation)

##### Before Correction

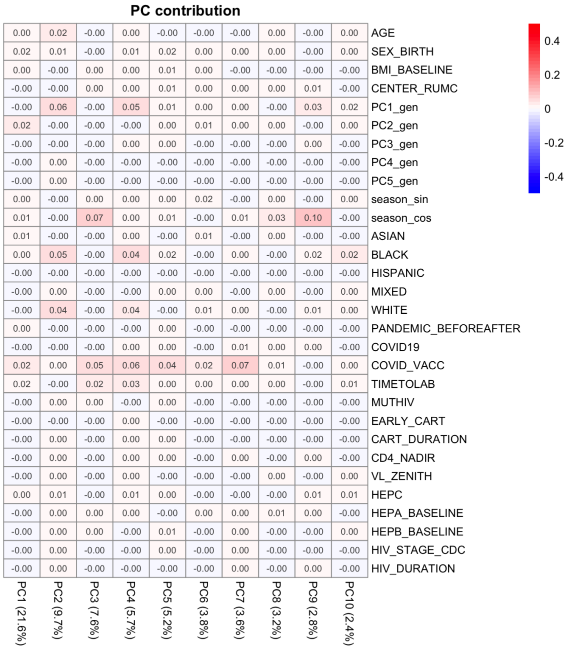


##### After Confounders Correction

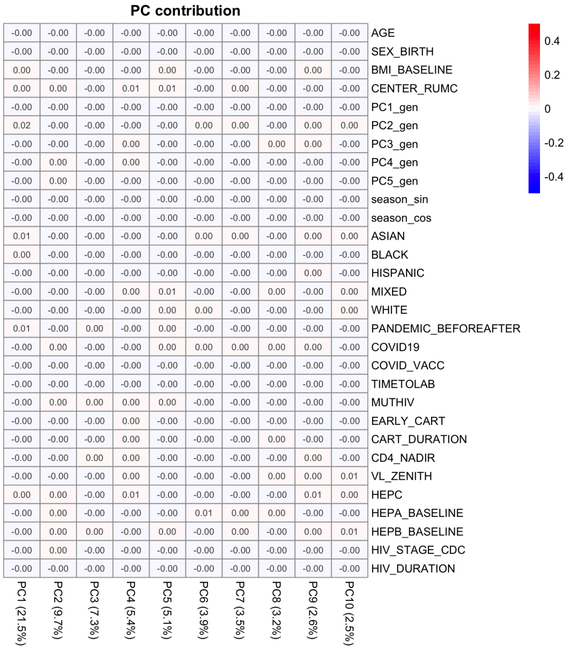


**Supplementary Figure 24.** ***Assessment of confounding factors in the ex vivo cytokine production data.*** (A) 7-day stimulation. (B) 24-hour stimulation. Heatmaps depict associations between the first 10 principal components (PCs) of cytokine profiles and candidate confounders. For each panel, the left heatmap shows beta coefficients from linear regression models prior to adjustment, and the right heatmap shows results after correction for key covariates: season of sample collection, age, sex, the first genetic principal component, COVID-19 vaccination status, and sample processing time.

Color intensity indicates the magnitude and direction of beta estimates. Confounders were defined as variables producing >10% change in beta coefficients in a multi-step regression framework. Following adjustment, no confounder exceeded this threshold, demonstrating effective control of potential confounding effects.

**Supplementary Figure 25 |** Variance Inflation Factor Assessment for Model Training

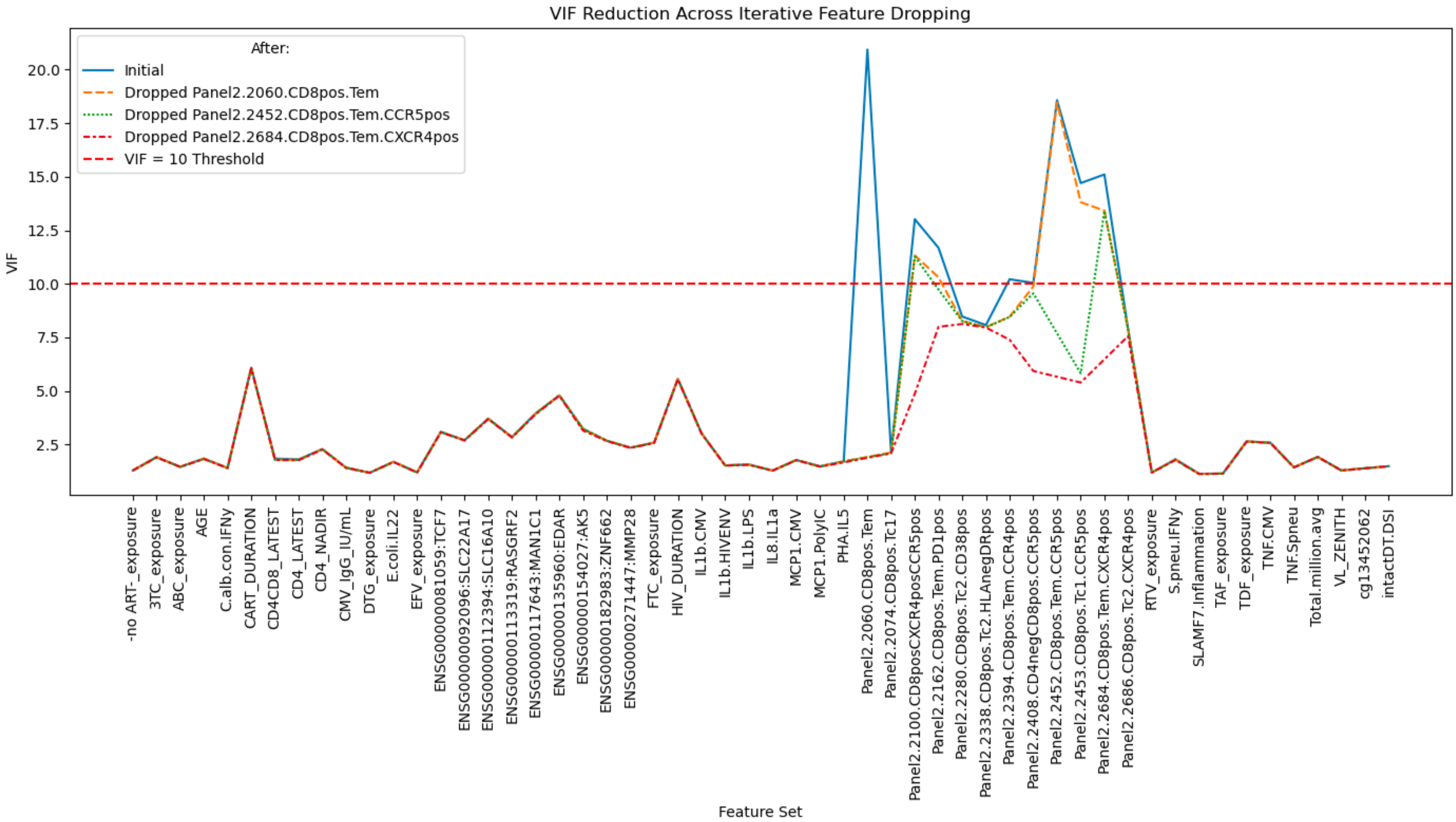


**Supplementary Figure 25. Variance Inflation Factor Assessment for Model Training**
This plot illustrates the variance inflation factor (VIF) values for all candidate features considered as inputs for the binary classification models across clusters. VIF quantifies the extent of multicollinearity among predictor variables, with higher values indicating greater redundancy and potential instability in regression estimates. To ensure robust model training, features with VIF values ≥ 10, indicative of serious multicollinearity, were iteratively removed: at each step, the feature with the highest VIF was dropped and VIFs were recalculated, continuing until all remaining features had VIF < 10. This approach minimizes multicollinearity, improving the interpretability and reliability of the model coefficients.

#### **Supplementary Figure 26 |** Spearman Correlation Heatmaps for Model Input Features

##### **A)** Before Assessing Correlated Features

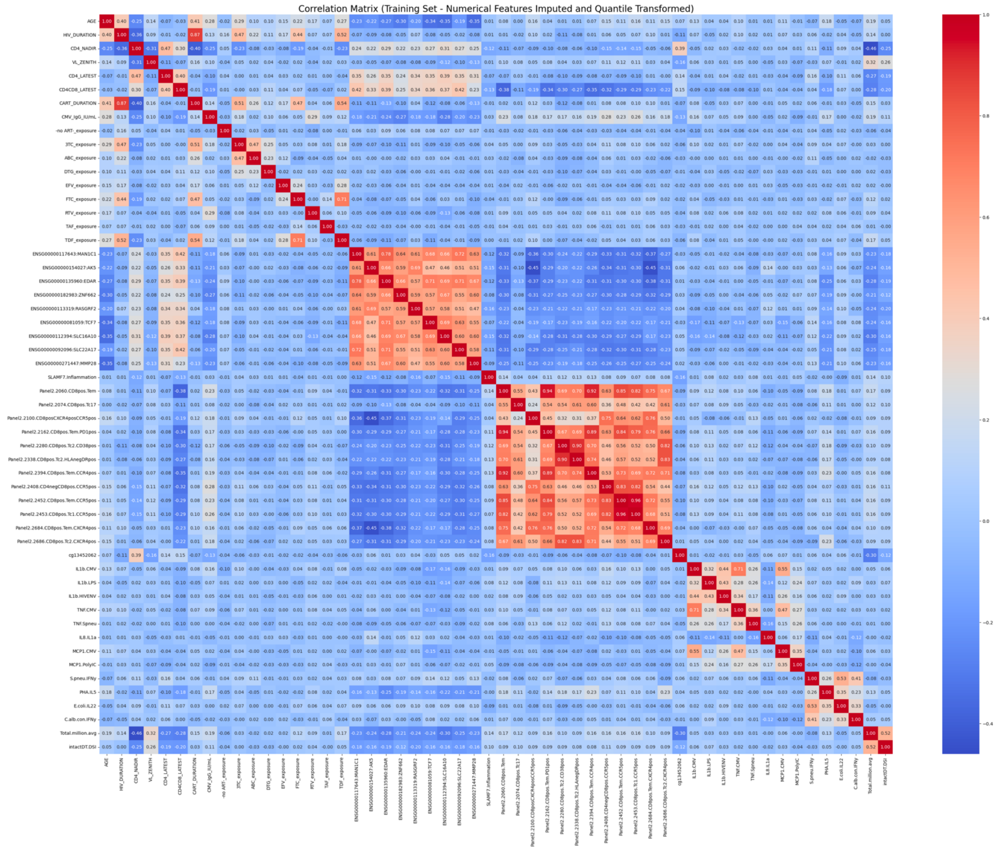


##### **B)** After Removal of Highly Correlated Features

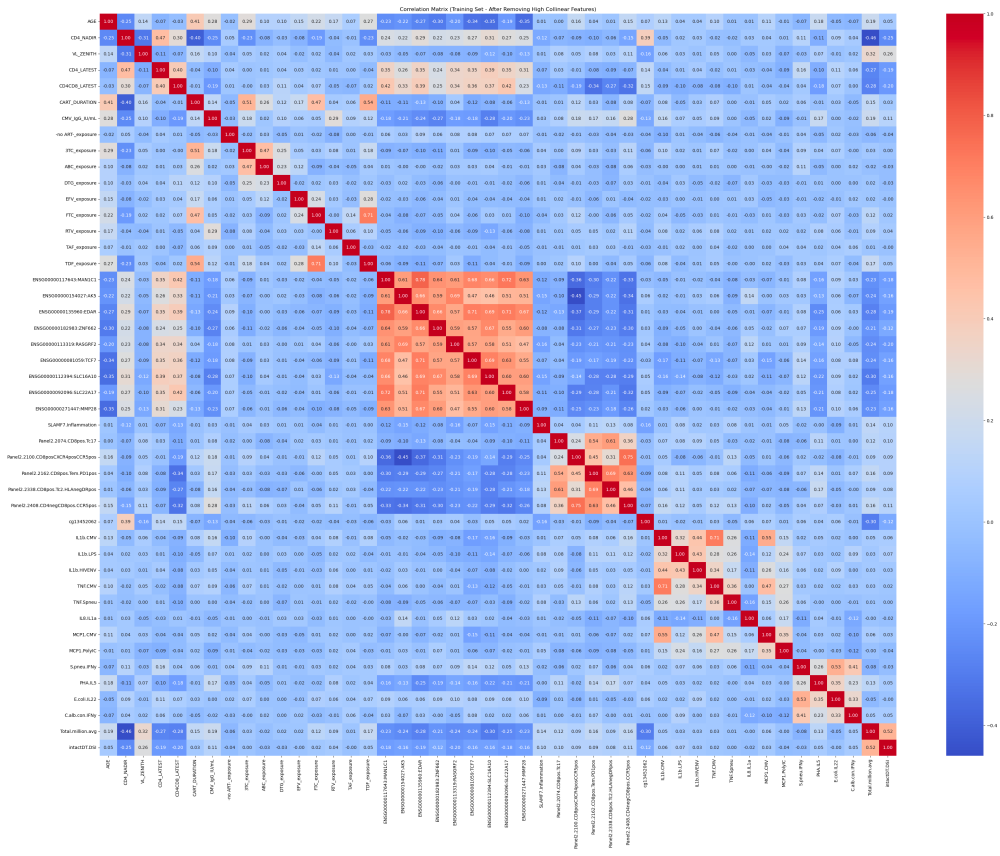


**Supplementary Figure 26. *Feature Spearman Correlations***
Spearman correlation heatmaps illustrating pairwise relationships among features considered for prediction model training. Spearman's rank correlation coefficient (ρ) quantifies the strength and direction of monotonic associations between features, with values ranging from -1 (perfect negative) to 1 (perfect positive).
**(A)** Correlation matrix for the original dataset, including all candidate features. Highly correlated feature pairs (|ρ| > 0.80) are identified as problematic due to potential redundancy and multicollinearity.
**(B)** Correlation matrix after removing features from highly correlated pairs (|ρ| > 0.80), retaining only one feature from each pair to improve model robustness and interpretability. The strongest remaining correlations are seen along the diagonal, as expected for self-correlation. Highly correlated features reduces redundancy and prepares the dataset for reliable model training.

#### **Supplementary Figure 27 | Feature Distribution Assessment for Model Training**

##### **A)** Before Feature Preprocessing

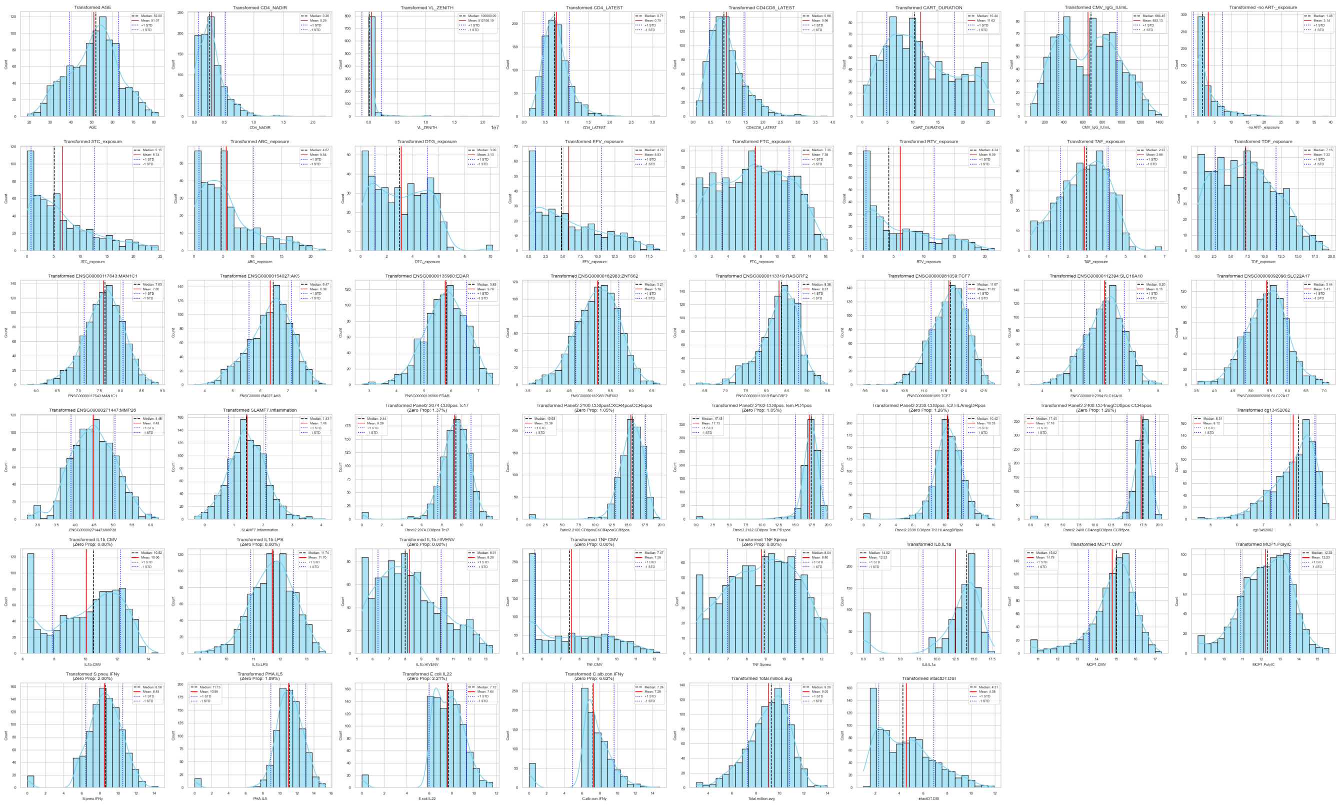

##### **B)** After Preprocessing

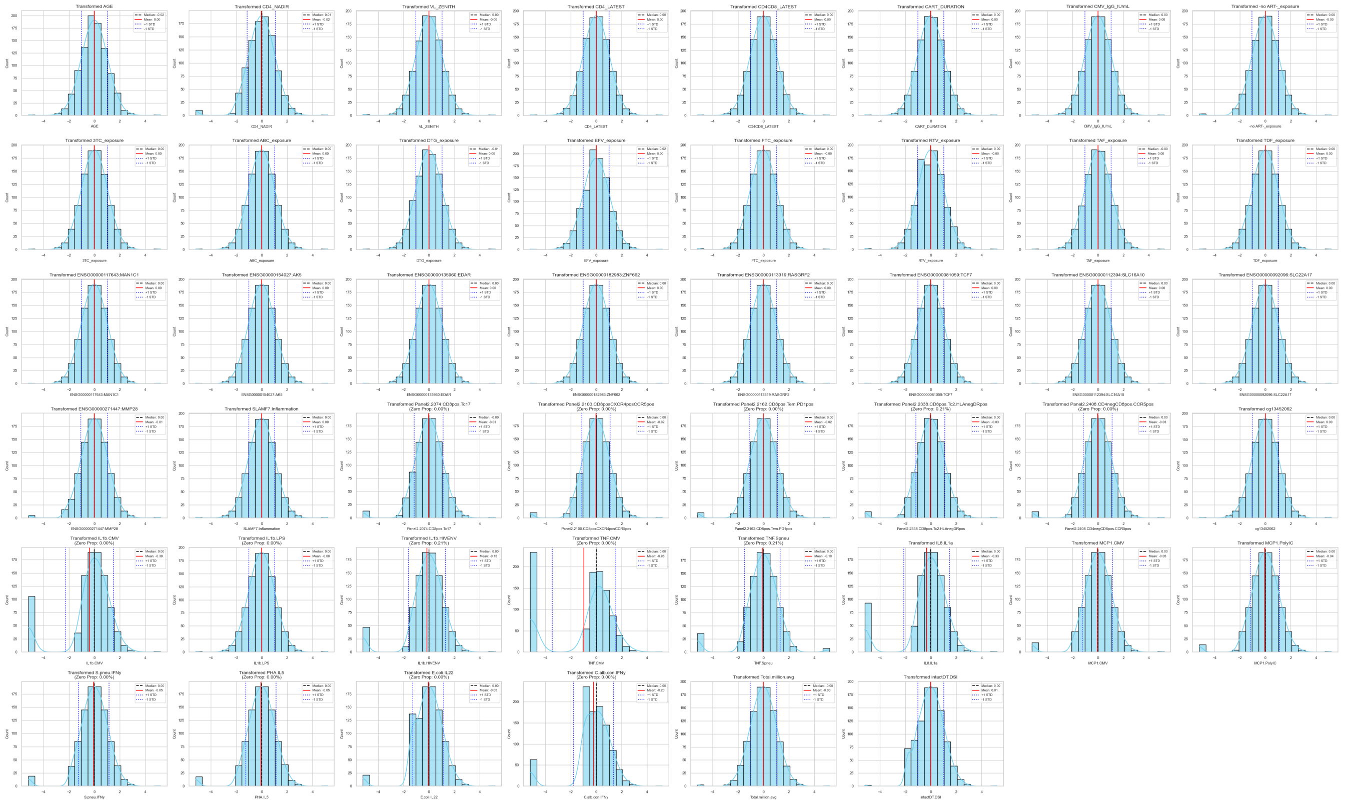

**Supplementary Figure 27.** ***Histogram of Features Distribution for Model Training***
This figure presents histograms illustrating the distribution of features used for model training, before and after data preprocessing steps.
**(A)** Raw distributions of all features prior to any data transformation. These histograms allow assessment of the initial range, central tendency, skewness, and presence of outliers for each variable, providing an overview of the dataset's original structure.
**(B)** Feature distributions after preprocessing, which included imputation of missing values, log transformation of skewed variables, and quantile transformation. These steps were applied to normalize distributions, reduce skewness, and make the features more suitable for machine learning model training.

### **Supplementary Figure 28 |** Q-Q Plots Assessing Feature Distributions Before and After Preprocessing

#### **A)** Before Feature Preprocessing

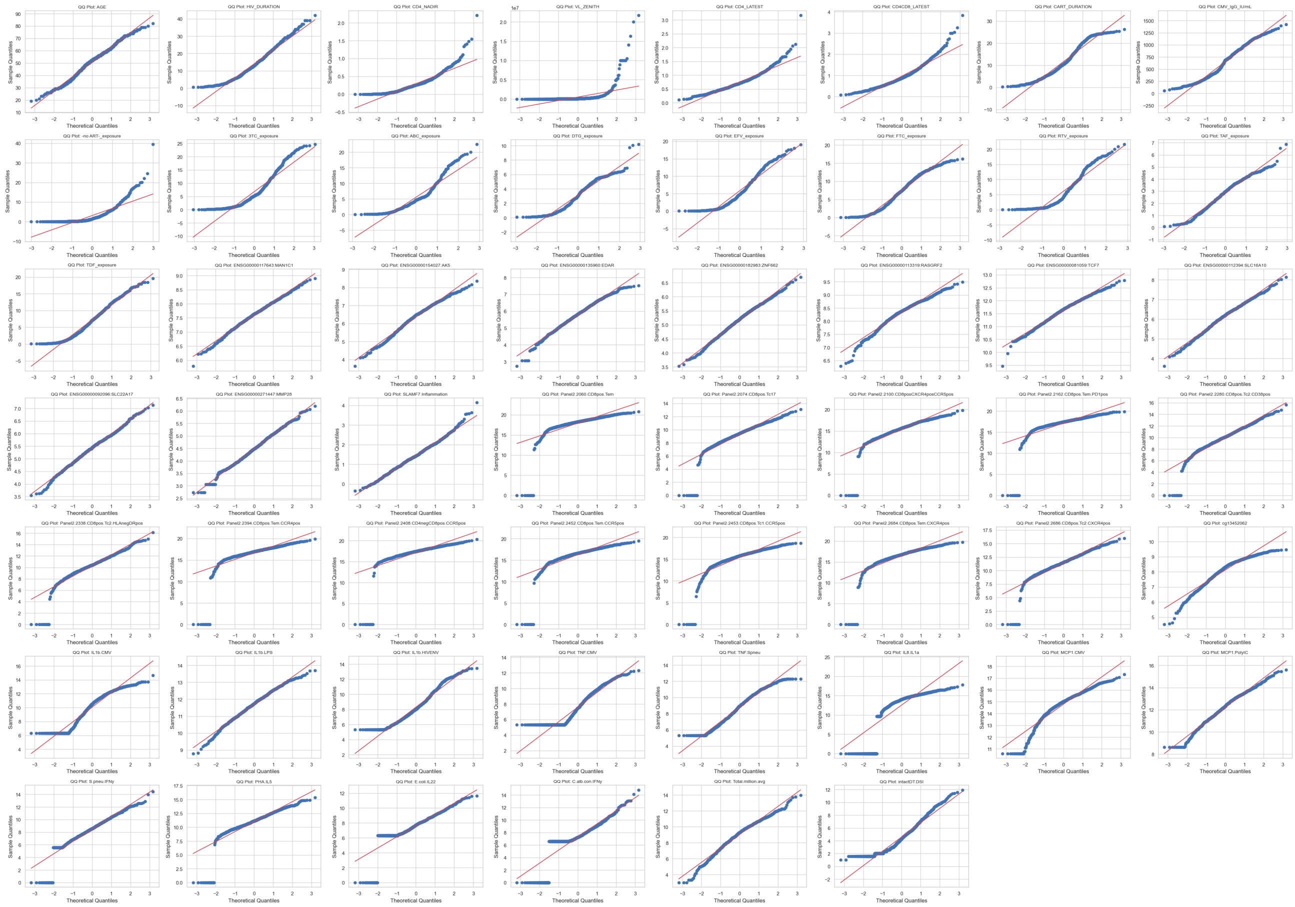

#### **B)** After Preprocessing

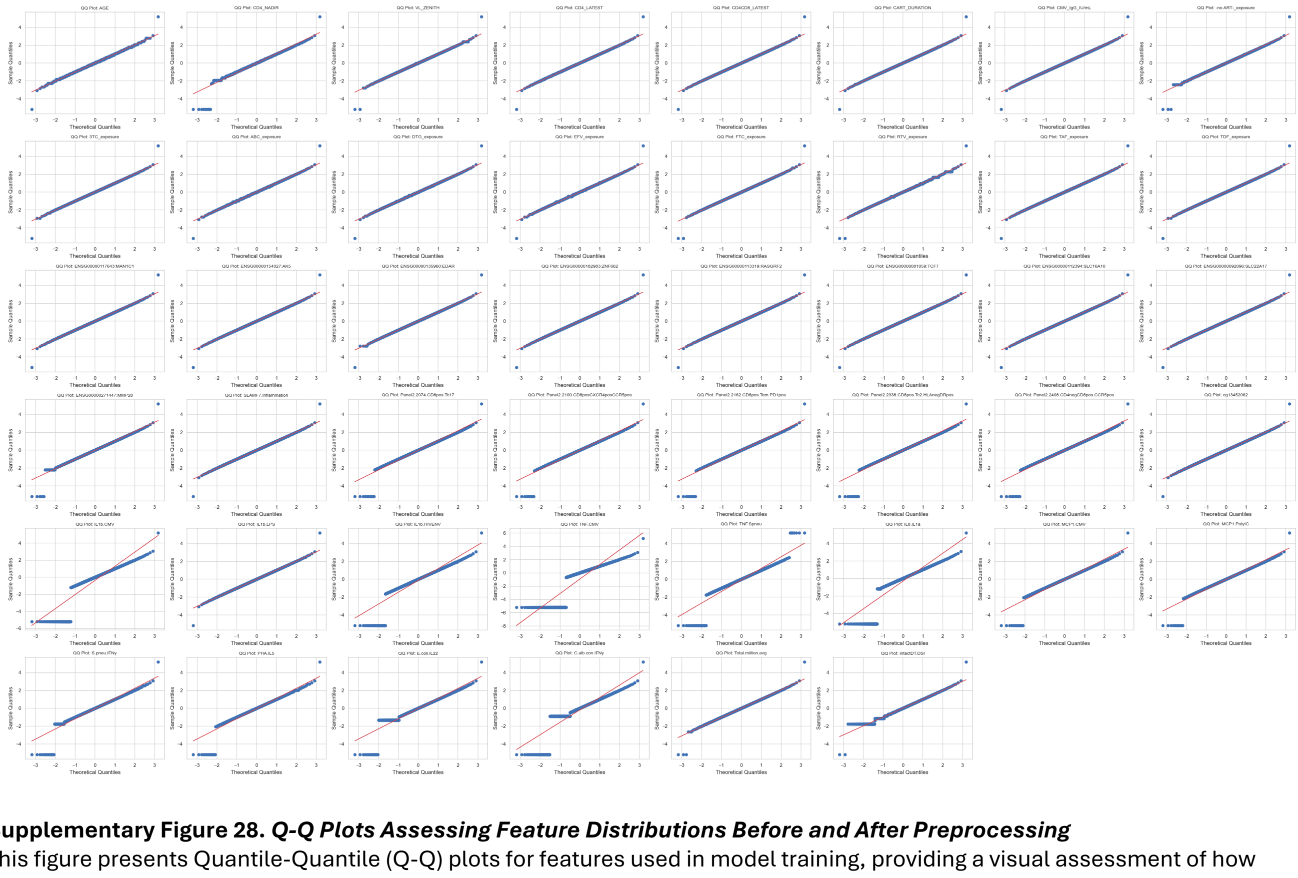

**Supplementary Figure 28.** ***Q-Q Plots Assessing Feature Distributions Before and After Preprocessing***
This figure presents Quantile-Quantile (Q-Q) plots for features used in model training, providing a visual assessment of how closely each feature's distribution approximates a theoretical normal distribution at different stages of data processing.
**(A)** Q-Q plots for all features in their raw, untransformed state. In these plots, deviations from the reference line indicate departures from normality, such as skewness, heavy tails, or outliers.
**(B)** Q-Q plots for the same features after data preprocessing, which included imputation of missing values, log transformation of skewed variables, and quantile transformation. Improved alignment of points along the reference line after preprocessing demonstrates increased normality and suitability for model training.
