## Supplementary Tables for "Multi-Omics Clustering Differentiates the Total and Intact HIV Reservoirs and Related Host Immune Mechanisms"

**Supplementary Table 1 | Standardized contrasts and effect sizes for validated host markers across reservoir-defined endotypes in the 2000HIV study.** This table reports mean standardized expression (z‑scores), standardized mean differences, and Cohen’s d effect sizes for validated host markers across the All Low, Mixed, and All High endotypes in the discovery (n=1,002) and validation (n=189) sub-cohorts. Markers span bulk transcriptomics (RNA‑seq genes) and ex vivo cytokine responses (cytokine–stimulus pairs), selected from single‑layer analyses and MoCluster results generated via the MOVICS pipeline. Z‑scores were derived from normalized data (variance‑stabilized counts for RNA‑seq; log₂‑transformed cytokine responses), enabling direct comparison across endotypes. Columns include marker identifier, cohort (discovery or validation), mean z‑score per endotype, standardized mean differences for each pairwise contrast, and Cohen’s d values for Mixed vs All Low, All High vs Mixed, and All High vs All Low. Effect sizes quantify the magnitude of between‑endotype differences and complement the statistical associations reported in the main text.

| Host Omic Layer | Marker | Sub-Cohort | Mean Z-score (All Low) | Mean Z-score (Mixed) | Mean Z-score (All High) | Standard Deviation Difference (Mixed vs All Low) | Standard Deviation Difference (All High vs Mixed) | Standard Deviation Difference (All High vs All Low) | Cohen's D (Mixed vs All Low) | Cohen's D (All High vs Mixed) | Cohen's D (All High vs All Low) |
| --- | --- | --- | --- | --- | --- | --- | --- | --- | --- | --- | --- |
| Bulk Transcriptomics | ENSG00000081059:TCF7 | Discovery | 0.155 | -0.030 | -0.145 | -0.186 | -0.115 | -0.300 | -0.21 | -0.13 | -0.33 |
| Bulk Transcriptomics | ENSG00000081059:TCF7 | Validation | 0.134 | 0.005 | -0.228 | -0.129 | -0.233 | -0.362 | -0.15 | -0.25 | -0.41 |
| Bulk Transcriptomics | ENSG00000092096:SLC22A17 | Discovery | 0.173 | -0.030 | -0.190 | -0.203 | -0.161 | -0.363 | -0.23 | -0.17 | -0.39 |
| Bulk Transcriptomics | ENSG00000092096:SLC22A17 | Validation | 0.205 | -0.127 | -0.185 | -0.332 | -0.058 | -0.390 | -0.37 | -0.06 | -0.42 |
| Bulk Transcriptomics | ENSG00000112394:SLC16A10 | Discovery | 0.234 | -0.074 | -0.192 | -0.308 | -0.118 | -0.426 | -0.35 | -0.13 | -0.48 |
| Bulk Transcriptomics | ENSG00000112394:SLC16A10 | Validation | 0.245 | -0.164 | -0.205 | -0.409 | -0.041 | -0.450 | -0.45 | -0.05 | -0.51 |
| Bulk Transcriptomics | ENSG00000113319:RASGRF2 | Discovery | 0.213 | -0.042 | -0.183 | -0.256 | -0.141 | -0.397 | -0.29 | -0.15 | -0.44 |
| Bulk Transcriptomics | ENSG00000113319:RASGRF2 | Validation | 0.214 | -0.061 | -0.313 | -0.275 | -0.252 | -0.528 | -0.3 | -0.28 | -0.57 |
| Bulk Transcriptomics | ENSG00000117643:MAN1C1 | Discovery | 0.174 | -0.039 | -0.165 | -0.213 | -0.127 | -0.339 | -0.24 | -0.14 | -0.36 |
| Bulk Transcriptomics | ENSG00000117643:MAN1C1 | Validation | 0.163 | -0.061 | -0.213 | -0.224 | -0.151 | -0.375 | -0.25 | -0.16 | -0.37 |
| Bulk Transcriptomics | ENSG00000135960:EDAR | Discovery | 0.237 | -0.063 | -0.200 | -0.300 | -0.138 | -0.437 | -0.34 | -0.15 | -0.48 |
| Bulk Transcriptomics | ENSG00000135960:EDAR | Validation | 0.208 | -0.126 | -0.198 | -0.334 | -0.072 | -0.406 | -0.37 | -0.08 | -0.44 |
| Bulk Transcriptomics | ENSG00000154027:AK5 | Discovery | 0.237 | -0.045 | -0.233 | -0.282 | -0.188 | -0.470 | -0.31 | -0.2 | -0.51 |
| Bulk Transcriptomics | ENSG00000154027:AK5 | Validation | 0.274 | -0.209 | -0.221 | -0.484 | -0.012 | -0.495 | -0.53 | -0.01 | -0.53 |
| Bulk Transcriptomics | ENSG00000182983:ZNF662 | Discovery | 0.193 | -0.068 | -0.143 | -0.260 | -0.075 | -0.335 | -0.29 | -0.08 | -0.36 |
| Bulk Transcriptomics | ENSG00000182983:ZNF662 | Validation | 0.213 | -0.119 | -0.239 | -0.332 | -0.119 | -0.451 | -0.36 | -0.12 | -0.45 |
| Bulk Transcriptomics | ENSG00000271447:MMP28 | Discovery | 0.202 | -0.068 | -0.163 | -0.270 | -0.095 | -0.365 | -0.3 | -0.11 | -0.4 |
| Bulk Transcriptomics | ENSG00000271447:MMP28 | Validation | 0.134 | -0.101 | -0.096 | -0.234 | 0.004 | -0.230 | -0.26 | 0.01 | -0.25 |
| Cytokine Responses | C.alb.con.IFNy | Discovery | -0.034 | 0.057 | 0.088 | 0.091 | 0.031 | 0.122 | 0.12 | 0.04 | 0.15 |
| Cytokine Responses | C.alb.con.IFNy | Validation | -0.088 | 0.127 | 0.143 | 0.215 | 0.016 | 0.230 | 0.28 | 0.02 | 0.38 |
| Cytokine Responses | E.coli.IL22 | Discovery | 0.090 | 0.019 | -0.048 | -0.071 | -0.067 | -0.139 | -0.09 | -0.09 | -0.18 |
| Cytokine Responses | E.coli.IL22 | Validation | -0.004 | 0.023 | 0.117 | 0.027 | 0.094 | 0.121 | 0.04 | 0.12 | 0.16 |
| Cytokine Responses | IL1b.CMV | Discovery | -0.052 | 0.013 | 0.052 | 0.065 | 0.039 | 0.104 | 0.07 | 0.04 | 0.1 |
| Cytokine Responses | IL1b.CMV | Validation | 0.002 | 0.148 | -0.259 | 0.145 | -0.406 | -0.261 | 0.15 | -0.41 | -0.27 |
| Cytokine Responses | IL1b.HIVENV | Discovery | -0.038 | -0.034 | 0.097 | 0.004 | 0.131 | 0.135 | 0 | 0.13 | 0.13 |
| Cytokine Responses | IL1b.HIVENV | Validation | -0.016 | 0.226 | -0.396 | 0.243 | -0.622 | -0.379 | 0.25 | -0.69 | -0.43 |
| Cytokine Responses | IL1b.LPS | Discovery | -0.039 | 0.016 | 0.059 | 0.055 | 0.043 | 0.098 | 0.06 | 0.05 | 0.11 |
| Cytokine Responses | IL1b.LPS | Validation | 0.056 | 0.022 | -0.114 | -0.034 | -0.135 | -0.169 | -0.04 | -0.14 | -0.19 |
| Cytokine Responses | IL8.IL1a | Discovery | 0.060 | 0.053 | 0.001 | -0.008 | -0.052 | -0.059 | -0.01 | -0.07 | -0.08 |
| Cytokine Responses | IL8.IL1a | Validation | -0.011 | -0.096 | 0.352 | -0.085 | 0.448 | 0.363 | -0.09 | 0.59 | 0.57 |
| Cytokine Responses | MCP1.CMV | Discovery | 0.000 | 0.012 | 0.072 | 0.012 | 0.059 | 0.072 | 0.01 | 0.07 | 0.08 |
| Cytokine Responses | MCP1.CMV | Validation | 0.040 | 0.089 | -0.153 | 0.049 | -0.242 | -0.192 | 0.06 | -0.24 | -0.2 |
| Cytokine Responses | MCP1.PolyIC | Discovery | 0.062 | 0.005 | -0.063 | -0.057 | -0.068 | -0.125 | -0.06 | -0.07 | -0.13 |
| Cytokine Responses | MCP1.PolyIC | Validation | -0.140 | 0.125 | 0.093 | 0.265 | -0.032 | 0.234 | 0.28 | -0.03 | 0.24 |
| Cytokine Responses | PHA.IL5 | Discovery | -0.100 | 0.079 | 0.138 | 0.179 | 0.060 | 0.238 | 0.25 | 0.08 | 0.34 |
| Cytokine Responses | PHA.IL5 | Validation | 0.008 | 0.030 | 0.122 | 0.022 | 0.092 | 0.114 | 0.03 | 0.15 | 0.18 |
| Cytokine Responses | S.pneu.IFNy | Discovery | 0.084 | 0.015 | -0.059 | -0.069 | -0.074 | -0.144 | -0.08 | -0.09 | -0.17 |
| Cytokine Responses | S.pneu.IFNy | Validation | 0.055 | 0.047 | -0.096 | -0.008 | -0.143 | -0.151 | -0.01 | -0.16 | -0.19 |
| Cytokine Responses | TNF.CMV | Discovery | -0.045 | 0.032 | 0.007 | 0.077 | -0.025 | 0.052 | 0.08 | -0.03 | 0.05 |
| Cytokine Responses | TNF.CMV | Validation | -0.047 | 0.143 | -0.157 | 0.190 | -0.300 | -0.110 | 0.19 | -0.3 | -0.12 |
| Cytokine Responses | TNF.Spneu | Discovery | -0.085 | 0.020 | 0.087 | 0.105 | 0.067 | 0.172 | 0.1 | 0.07 | 0.17 |
| Cytokine Responses | TNF.Spneu | Validation | -0.003 | 0.168 | -0.282 | 0.171 | -0.449 | -0.278 | 0.17 | -0.47 | -0.28 |
| Immunophenotyping | Panel2.2060.CD8pos.Tem | Discovery | -0.054 | 0.074 | 0.105 | 0.128 | 0.031 | 0.159 | 0.23 | 0.06 | 0.3 |
| Immunophenotyping | Panel2.2060.CD8pos.Tem | Validation | -0.029 | 0.127 | 0.041 | 0.156 | -0.086 | 0.070 | 0.32 | -0.22 | 0.14 |
| Immunophenotyping | Panel2.2074.CD8pos.Tc17 | Discovery | -0.066 | 0.037 | 0.153 | 0.103 | 0.116 | 0.219 | 0.14 | 0.16 | 0.29 |
| Immunophenotyping | Panel2.2074.CD8pos.Tc17 | Validation | -0.086 | 0.086 | 0.193 | 0.173 | 0.107 | 0.279 | 0.23 | 0.16 | 0.39 |
| Immunophenotyping | Panel2.2100.CD8posCXCR4posCCR5pos | Discovery | -0.036 | 0.038 | 0.115 | 0.074 | 0.077 | 0.151 | 0.11 | 0.11 | 0.22 |
| Immunophenotyping | Panel2.2100.CD8posCXCR4posCCR5pos | Validation | -0.066 | 0.124 | 0.079 | 0.190 | -0.044 | 0.145 | 0.27 | -0.07 | 0.2 |
| Immunophenotyping | Panel2.2162.CD8pos.Tem.PD1pos | Discovery | -0.060 | 0.075 | 0.114 | 0.135 | 0.039 | 0.175 | 0.25 | 0.07 | 0.33 |
| Immunophenotyping | Panel2.2162.CD8pos.Tem.PD1pos | Validation | -0.030 | 0.140 | 0.022 | 0.170 | -0.118 | 0.052 | 0.35 | -0.29 | 0.1 |
| Immunophenotyping | Panel2.2280.CD8pos.Tc2.CD38pos | Discovery | -0.071 | 0.033 | 0.133 | 0.104 | 0.099 | 0.203 | 0.13 | 0.12 | 0.25 |
| Immunophenotyping | Panel2.2280.CD8pos.Tc2.CD38pos | Validation | -0.077 | 0.104 | 0.098 | 0.181 | -0.006 | 0.175 | 0.24 | -0.01 | 0.2 |
| Immunophenotyping | Panel2.2338.CD8pos.Tc2.HLAnegDRpos | Discovery | -0.066 | 0.048 | 0.113 | 0.114 | 0.065 | 0.179 | 0.14 | 0.08 | 0.23 |
| Immunophenotyping | Panel2.2338.CD8pos.Tc2.HLAnegDRpos | Validation | -0.078 | 0.043 | 0.207 | 0.121 | 0.164 | 0.285 | 0.15 | 0.2 | 0.34 |
| Immunophenotyping | Panel2.2394.CD8pos.Tem.CCR4pos | Discovery | -0.058 | 0.073 | 0.114 | 0.131 | 0.041 | 0.172 | 0.23 | 0.07 | 0.32 |
| Immunophenotyping | Panel2.2394.CD8pos.Tem.CCR4pos | Validation | -0.053 | 0.141 | 0.065 | 0.194 | -0.076 | 0.118 | 0.38 | -0.18 | 0.23 |
| Immunophenotyping | Panel2.2408.CD4negCD8pos.CCR5pos | Discovery | -0.042 | 0.092 | 0.082 | 0.134 | -0.009 | 0.125 | 0.27 | -0.02 | 0.25 |
| Immunophenotyping | Panel2.2408.CD4negCD8pos.CCR5pos | Validation | -0.019 | 0.104 | 0.054 | 0.123 | -0.051 | 0.073 | 0.23 | -0.12 | 0.14 |
| Immunophenotyping | Panel2.2452.CD8pos.Tem.CCR5pos | Discovery | -0.072 | 0.088 | 0.098 | 0.160 | 0.010 | 0.170 | 0.26 | 0.02 | 0.28 |
| Immunophenotyping | Panel2.2452.CD8pos.Tem.CCR5pos | Validation | -0.050 | 0.154 | 0.025 | 0.204 | -0.128 | 0.076 | 0.37 | -0.25 | 0.13 |
| Immunophenotyping | Panel2.2453.CD8pos.Tc1.CCR5pos | Discovery | -0.077 | 0.101 | 0.081 | 0.179 | -0.020 | 0.159 | 0.27 | -0.03 | 0.24 |
| Immunophenotyping | Panel2.2453.CD8pos.Tc1.CCR5pos | Validation | -0.040 | 0.161 | -0.018 | 0.201 | -0.179 | 0.022 | 0.33 | -0.32 | 0.03 |
| Immunophenotyping | Panel2.2684.CD8pos.Tem.CXCR4pos | Discovery | -0.030 | 0.033 | 0.134 | 0.063 | 0.100 | 0.164 | 0.1 | 0.16 | 0.27 |
| Immunophenotyping | Panel2.2684.CD8pos.Tem.CXCR4pos | Validation | -0.071 | 0.168 | 0.044 | 0.239 | -0.124 | 0.115 | 0.42 | -0.22 | 0.19 |
| Immunophenotyping | Panel2.2686.CD8pos.Tc2.CXCR4pos | Discovery | -0.041 | 0.011 | 0.149 | 0.052 | 0.138 | 0.190 | 0.07 | 0.18 | 0.25 |
| Immunophenotyping | Panel2.2686.CD8pos.Tc2.CXCR4pos | Validation | -0.074 | 0.090 | 0.137 | 0.164 | 0.047 | 0.211 | 0.22 | 0.06 | 0.28 |
| DNA Methylation | cg13452062 | Discovery | 0.296 | -0.083 | -0.210 | -0.378 | -0.127 | -0.506 | -0.44 | -0.14 | -0.58 |
| DNA Methylation | cg13452062 | Validation | 0.192 | -0.094 | -0.179 | -0.286 | -0.085 | -0.371 | -0.33 | -0.09 | -0.38 |
| Proteomics (OLINK Explore) | SLAMF7.Inflammation | Discovery | -0.126 | 0.026 | 0.118 | 0.152 | 0.092 | 0.244 | 0.16 | 0.1 | 0.26 |
| Proteomics (OLINK Explore) | SLAMF7.Inflammation | Validation | -0.188 | 0.135 | 0.166 | 0.323 | 0.031 | 0.354 | 0.35 | 0.03 | 0.37 |
| HIV Viral Reservoir | Total.million.avg | Discovery | -0.998 | 0.398 | 0.776 | 1.396 | 0.378 | 1.774 | 2.56 | 0.66 | 2.92 |
| HIV Viral Reservoir | Total.million.avg | Validation | -0.836 | 0.551 | 0.875 | 1.387 | 0.324 | 1.712 | 2.42 | 0.59 | 2.89 |
| HIV Viral Reservoir | intactDT.DSI | Discovery | -0.735 | -0.188 | 1.340 | 0.547 | 1.529 | 2.075 | 1.02 | 2.82 | 4.34 |
| HIV Viral Reservoir | intactDT.DSI | Validation | -0.539 | -0.157 | 1.386 | 0.382 | 1.543 | 1.924 | 0.56 | 2.77 | 3.27 |

**Supplementary Table 2 | Partial correlation analyses between host markers and HIV reservoir in the 2000HIV study.** The table reports partial Spearman correlations between validated host markers and HIV reservoir metrics (total and intact HIV DNA per million CD4+ T cells) in the discovery (n=1,002) and validation (n=189) cohorts, stratified by endotypes: High-total Low-intact (MIXED), Low-total Low-intact (ALL LOW), and High-total High-intact (ALL HIGH). Markers include transcriptomic (e.g., RNA-seq genes), proteomic (e.g., inflammation markers), flow cytometry (absolute counts and percentages), ex vivo cytokine (e.g., cytokine/stimulus pairs), and DNA methylation (e.g., CpG sites) data, selected from single-layer analyses and MoCluster results via the MOVICS clustering pipeline. Data were transformed for analysis (variance-stabilized for RNA-seq; log-2 transformed for proteomics, flow cytometry, cytokines, and HIV DNA; shifted M-values for methylation). Columns include marker identifier, endotype, subset (discovery or validation), correlation coefficients (R Total, R Intact), nominal P-values (P-value Total, P-value Intact), 95% confidence intervals (CI Total Low, CI Total High, CI Intact Low, CI Intact High), FDR-adjusted P-values (P-value Adj. Total, P-value Adj. Intact), and significance indicators for total and intact reservoir (*P<0.05, **P<0.01). Partial correlations were calculated using the ppcor package (v1.1) in R (v4.3.2), adjusting for age and sex as confounders, with confidence intervals derived via Fisher’s z-transformation. In the discovery cohort, P-values were adjusted for multiple testing using the false discovery rate (FDR) method; in the validation cohort, nominal P-values were reported. Only markers with at least one significant association (Discovery *FDR<0.05 or **FDR<0.01 or Validation *P<0.05 or **P<0.01) in either total or intact reservoir metrics are included. Results were visualized as a heatmap using ggplot2 (v3.5.0) and cowplot (v1.1.3). cART, combination antiretroviral therapy; FDR, false discovery rate.

| **Marker** | **Endotype** | **Subset** | **R Total** | **P-value Total** | **CI Total Low** | **CI Total High** | **R Intact** | **P-value Intact** | **CI Intact Low** | **CI Intact High** | **P-value Adj. Total** | **P-value Adj. Intact** | | **Sign. Total** | | **Sign. Intact** |
| --- | --- | --- | --- | --- | --- | --- | --- | --- | --- | --- | --- | --- | --- | --- | --- | --- |
| BulkRNAseq_ENSG00000081059:TCF7 | MIXED | Disc. | -0.154 | 0.002 | -0.246 | -0.058 | 0.019 | 0.695 | -0.077 | 0.116 | 0.011 | 0.911 | * | |  | |
| BulkRNAseq_ENSG00000081059:TCF7 | ALL HIGH | Disc. | -0.247 | 0.000 | -0.363 | -0.124 | -0.219 | 0.001 | -0.337 | -0.094 | 0.003 | 0.208 | ** | |  | |
| BulkRNAseq_ENSG00000081059:TCF7 | ALL HIGH | Val. | -0.411 | 0.011 | -0.652 | -0.096 | -0.103 | 0.543 | -0.418 | 0.233 | 0.011 | 0.543 | * | |  | |
| BulkRNAseq_ENSG00000092096:SLC22A17 | MIXED | Disc. | -0.176 | 0.000 | -0.268 | -0.080 | -0.059 | 0.235 | -0.154 | 0.038 | 0.004 | 0.718 | ** | |  | |
| BulkRNAseq_ENSG00000092096:SLC22A17 | ALL HIGH | Disc. | -0.228 | 0.000 | -0.346 | -0.104 | -0.165 | 0.011 | -0.286 | -0.038 | 0.004 | 0.472 | ** | |  | |
| BulkRNAseq_ENSG00000112394:SLC16A10 | MIXED | Disc. | -0.211 | 0.000 | -0.301 | -0.117 | 0.020 | 0.681 | -0.076 | 0.117 | 0.001 | 0.903 | ** | |  | |
| BulkRNAseq_ENSG00000112394:SLC16A10 | ALL HIGH | Disc. | -0.250 | 0.000 | -0.366 | -0.127 | -0.163 | 0.012 | -0.284 | -0.036 | 0.003 | 0.472 | ** | |  | |
| BulkRNAseq_ENSG00000112394:SLC16A10 | ALL HIGH | Val. | -0.523 | 0.001 | -0.727 | -0.235 | -0.162 | 0.339 | -0.466 | 0.176 | 0.001 | 0.339 | ** | |  | |
| BulkRNAseq_ENSG00000113319:RASGRF2 | ALL HIGH | Disc. | -0.161 | 0.013 | -0.283 | -0.035 | -0.131 | 0.043 | -0.254 | -0.004 | 0.049 | 0.600 | * | |  | |
| BulkRNAseq_ENSG00000117643:MAN1C1 | MIXED | Disc. | -0.200 | 0.000 | -0.291 | -0.106 | -0.091 | 0.065 | -0.186 | 0.006 | 0.002 | 0.607 | ** | |  | |
| BulkRNAseq_ENSG00000117643:MAN1C1 | ALL HIGH | Disc. | -0.186 | 0.004 | -0.306 | -0.061 | -0.154 | 0.017 | -0.276 | -0.027 | 0.020 | 0.472 | * | |  | |
| BulkRNAseq_ENSG00000135960:EDAR | MIXED | Disc. | -0.203 | 0.000 | -0.294 | -0.109 | -0.056 | 0.255 | -0.152 | 0.041 | 0.002 | 0.719 | ** | |  | |
| BulkRNAseq_ENSG00000135960:EDAR | ALL HIGH | Disc. | -0.231 | 0.000 | -0.348 | -0.107 | -0.190 | 0.003 | -0.309 | -0.064 | 0.004 | 0.346 | ** | |  | |
| BulkRNAseq_ENSG00000138386:NAB1 | MIXED | Disc. | 0.138 | 0.005 | 0.042 | 0.231 | 0.110 | 0.025 | 0.014 | 0.204 | 0.024 | 0.501 | * | |  | |
| BulkRNAseq_ENSG00000154027:AK5 | MIXED | Disc. | -0.226 | 0.000 | -0.316 | -0.132 | -0.031 | 0.526 | -0.127 | 0.065 | 0.000 | 0.879 | ** | |  | |
| BulkRNAseq_ENSG00000182983:ZNF662 | MIXED | Disc. | -0.161 | 0.001 | -0.254 | -0.066 | 0.006 | 0.908 | -0.091 | 0.102 | 0.007 | 0.985 | ** | |  | |
| BulkRNAseq_ENSG00000186310:NAP1L3 | MIXED | Disc. | -0.155 | 0.002 | -0.248 | -0.059 | -0.022 | 0.658 | -0.118 | 0.075 | 0.010 | 0.895 | * | |  | |
| BulkRNAseq_ENSG00000186310:NAP1L3 | ALL HIGH | Disc. | -0.207 | 0.001 | -0.325 | -0.082 | -0.136 | 0.036 | -0.258 | -0.009 | 0.009 | 0.573 | ** | |  | |
| BulkRNAseq_ENSG00000186310:NAP1L3 | ALL HIGH | Val. | -0.060 | 0.726 | -0.381 | 0.274 | -0.326 | 0.049 | -0.591 | 0.003 | 0.726 | 0.049 |  | | * | |
| BulkRNAseq_ENSG00000217165:ANKRD18EP | MIXED | Disc. | -0.128 | 0.009 | -0.222 | -0.032 | 0.053 | 0.278 | -0.043 | 0.149 | 0.039 | 0.740 | * | |  | |
| BulkRNAseq_ENSG00000271447:MMP28 | MIXED | Disc. | -0.165 | 0.001 | -0.257 | -0.069 | -0.014 | 0.781 | -0.110 | 0.083 | 0.007 | 0.959 | ** | |  | |
| BulkRNAseq_ENSG00000271447:MMP28 | ALL LOW | Val. | -0.225 | 0.043 | -0.424 | -0.006 | -0.273 | 0.014 | -0.465 | -0.057 | 0.043 | 0.014 | * | | * | |
| DNAMethylation_cg13452062 | MIXED | Disc. | -0.210 | 0.000 | -0.301 | -0.116 | -0.029 | 0.555 | -0.125 | 0.068 | 0.001 | 0.879 | ** | |  | |
| DNAMethylation_cg13452062 | ALL LOW | Disc. | -0.174 | 0.001 | -0.275 | -0.069 | 0.026 | 0.627 | -0.080 | 0.132 | 0.009 | 0.891 | ** | |  | |
| DNAMethylation_cg13452062 | ALL HIGH | Disc. | -0.328 | 0.000 | -0.437 | -0.210 | -0.106 | 0.103 | -0.230 | 0.022 | 0.000 | 0.607 | ** | |  | |
| DNAMethylation_cg13452062 | ALL HIGH | Val. | -0.504 | 0.001 | -0.714 | -0.210 | -0.141 | 0.405 | -0.449 | 0.197 | 0.001 | 0.405 | ** | |  | |
| ExVivo_IL1b_CMV | MIXED | Val. | 0.219 | 0.080 | -0.029 | 0.441 | -0.380 | 0.002 | -0.572 | -0.148 | 0.080 | 0.002 |  | | ** | |
| ExVivo_IL1b_HIVENV | MIXED | Val. | 0.229 | 0.067 | -0.018 | 0.449 | -0.249 | 0.045 | -0.466 | -0.003 | 0.067 | 0.045 |  | | * | |
| ExVivo_IL1b_HIVENV | ALL HIGH | Val. | 0.126 | 0.456 | -0.211 | 0.437 | -0.374 | 0.023 | -0.625 | -0.051 | 0.456 | 0.023 |  | | * | |
| ExVivo_IL1b_LPS | MIXED | Val. | 0.233 | 0.062 | -0.014 | 0.453 | -0.284 | 0.022 | -0.495 | -0.041 | 0.062 | 0.022 |  | | * | |
| ExVivo_IL8_IL1a | ALL HIGH | Val. | -0.460 | 0.004 | -0.685 | -0.155 | -0.188 | 0.265 | -0.487 | 0.150 | 0.004 | 0.265 | ** | |  | |
| ExVivo_MCP1_PolyIC | MIXED | Val. | 0.321 | 0.009 | 0.082 | 0.525 | -0.210 | 0.094 | -0.433 | 0.038 | 0.009 | 0.094 | ** | |  | |
| ExVivo_MTB_IFNy | ALL HIGH | Disc. | -0.181 | 0.005 | -0.302 | -0.056 | -0.153 | 0.018 | -0.275 | -0.027 | 0.024 | 0.472 | * | |  | |
| ExVivo_MTB_IFNy | MIXED | Val. | 0.108 | 0.391 | -0.141 | 0.345 | -0.250 | 0.045 | -0.467 | -0.004 | 0.391 | 0.045 |  | | * | |
| ExVivo_PHA_IL10 | ALL HIGH | Disc. | -0.192 | 0.003 | -0.312 | -0.067 | -0.149 | 0.022 | -0.271 | -0.022 | 0.016 | 0.491 | * | |  | |
| ExVivo_PHA_IL10 | ALL HIGH | Val. | -0.295 | 0.077 | -0.568 | 0.038 | -0.465 | 0.004 | -0.688 | -0.161 | 0.077 | 0.004 |  | | ** | |
| ExVivo_S.pneu_IFNy | ALL LOW | Val. | -0.110 | 0.328 | -0.322 | 0.112 | -0.256 | 0.021 | -0.450 | -0.038 | 0.328 | 0.021 |  | | * | |
| ExVivo_TNF_CMV | ALL LOW | Disc. | 0.143 | 0.008 | 0.038 | 0.246 | 0.118 | 0.028 | 0.013 | 0.222 | 0.034 | 0.528 | * | |  | |
| ExVivo_TNF_CMV | MIXED | Val. | 0.213 | 0.088 | -0.034 | 0.436 | -0.384 | 0.002 | -0.576 | -0.153 | 0.088 | 0.002 |  | | ** | |
| FlowAbs_Panel2_2018_CD4+CD8- | MIXED | Disc. | -0.177 | 0.000 | -0.268 | -0.081 | -0.043 | 0.382 | -0.139 | 0.054 | 0.004 | 0.804 | ** | |  | |
| FlowAbs_Panel2_2025_nTreg | MIXED | Disc. | -0.162 | 0.001 | -0.254 | -0.066 | -0.077 | 0.120 | -0.172 | 0.020 | 0.007 | 0.607 | ** | |  | |
| FlowAbs_Panel2_2035_Tfh_1 | ALL HIGH | Disc. | -0.171 | 0.008 | -0.292 | -0.045 | -0.110 | 0.091 | -0.234 | 0.018 | 0.035 | 0.607 | * | |  | |
| FlowAbs_Panel2_2043_CD4+.Tcm | MIXED | Disc. | -0.124 | 0.012 | -0.218 | -0.028 | -0.016 | 0.747 | -0.112 | 0.081 | 0.047 | 0.947 | * | |  | |
| FlowAbs_Panel2_2043_CD4+.Tcm | ALL HIGH | Disc. | -0.179 | 0.005 | -0.299 | -0.053 | -0.091 | 0.163 | -0.215 | 0.037 | 0.026 | 0.651 | * | |  | |
| FlowAbs_Panel2_2045_CD4+.Temra | MIXED | Val. | 0.247 | 0.047 | 0.001 | 0.465 | -0.381 | 0.002 | -0.573 | -0.149 | 0.047 | 0.002 | * | | ** | |
| FlowAbs_Panel2_2046_CD4+.Tnaive | MIXED | Disc. | -0.179 | 0.000 | -0.271 | -0.084 | -0.057 | 0.248 | -0.153 | 0.040 | 0.004 | 0.718 | ** | |  | |
| FlowAbs_Panel2_2046_CD4+.Tnaive | ALL HIGH | Disc. | -0.185 | 0.004 | -0.305 | -0.059 | -0.051 | 0.433 | -0.177 | 0.077 | 0.021 | 0.856 | * | |  | |
| FlowAbs_Panel2_2086_CD4+HLA-DR-CD38+ | MIXED | Disc. | -0.189 | 0.000 | -0.280 | -0.094 | -0.055 | 0.268 | -0.150 | 0.042 | 0.003 | 0.738 | ** | |  | |
| FlowAbs_Panel2_2086_CD4+HLA-DR-CD38+ | ALL HIGH | Disc. | -0.165 | 0.010 | -0.286 | -0.039 | -0.034 | 0.599 | -0.161 | 0.093 | 0.043 | 0.891 | * | |  | |
| FlowAbs_Panel2_2095_CD4+CXCR4+CCR5- | MIXED | Disc. | -0.179 | 0.000 | -0.271 | -0.084 | -0.008 | 0.874 | -0.104 | 0.089 | 0.004 | 0.984 | ** | |  | |
| FlowAbs_Panel2_2100_CD8+CXCR4+CCR5+ | ALL HIGH | Val. | -0.084 | 0.622 | -0.401 | 0.252 | 0.331 | 0.045 | 0.003 | 0.595 | 0.622 | 0.045 |  | | * | |
| FlowAbs_Panel2_2140_CD4+.Tnaive_PD1+ | MIXED | Disc. | -0.145 | 0.003 | -0.238 | -0.049 | -0.048 | 0.331 | -0.144 | 0.049 | 0.017 | 0.751 | * | |  | |
| FlowAbs_Panel2_2140_CD4+.Tnaive_PD1+ | MIXED | Val. | 0.066 | 0.602 | -0.183 | 0.307 | -0.259 | 0.037 | -0.475 | -0.014 | 0.602 | 0.037 |  | | * | |
| FlowAbs_Panel2_2162_CD8+.Tem_PD1+ | MIXED | Disc. | 0.161 | 0.001 | 0.065 | 0.253 | 0.081 | 0.100 | -0.016 | 0.176 | 0.007 | 0.607 | ** | |  | |
| FlowAbs_Panel2_2162_CD8+.Tem_PD1+ | ALL LOW | Val. | 0.225 | 0.043 | 0.006 | 0.424 | 0.037 | 0.743 | -0.184 | 0.255 | 0.043 | 0.743 | * | |  | |
| FlowAbs_Panel2_2233_CD4+CD8-_CD38+ | MIXED | Disc. | -0.198 | 0.000 | -0.289 | -0.104 | -0.050 | 0.307 | -0.146 | 0.046 | 0.002 | 0.740 | ** | |  | |
| FlowAbs_Panel2_2240_Treg_CD38+ | MIXED | Disc. | -0.164 | 0.001 | -0.257 | -0.069 | -0.026 | 0.601 | -0.122 | 0.071 | 0.007 | 0.891 | ** | |  | |
| FlowAbs_Panel2_2242_nTreg_CD38+ | MIXED | Disc. | -0.162 | 0.001 | -0.255 | -0.067 | -0.076 | 0.123 | -0.171 | 0.021 | 0.007 | 0.607 | ** | |  | |
| FlowAbs_Panel2_2255_CD4+.Tcm_CD38+ | MIXED | Disc. | -0.152 | 0.002 | -0.245 | -0.057 | -0.016 | 0.753 | -0.112 | 0.081 | 0.012 | 0.950 | * | |  | |
| FlowAbs_Panel2_2255_CD4+.Tcm_CD38+ | ALL HIGH | Disc. | -0.195 | 0.002 | -0.314 | -0.069 | -0.107 | 0.100 | -0.231 | 0.021 | 0.015 | 0.607 | * | |  | |
| FlowAbs_Panel2_2255_CD4+.Tcm_CD38+ | ALL HIGH | Val. | -0.348 | 0.035 | -0.607 | -0.022 | -0.052 | 0.758 | -0.374 | 0.281 | 0.035 | 0.758 | * | |  | |
| FlowAbs_Panel2_2256_CD4+.Tnaive_CD38+ | MIXED | Disc. | -0.190 | 0.000 | -0.282 | -0.096 | -0.049 | 0.323 | -0.145 | 0.048 | 0.003 | 0.751 | ** | |  | |
| FlowAbs_Panel2_2259_Tfh_1_CD38+ | ALL HIGH | Disc. | -0.198 | 0.002 | -0.318 | -0.073 | -0.111 | 0.088 | -0.234 | 0.017 | 0.013 | 0.607 | * | |  | |
| FlowAbs_Panel2_2263_CD4+.Th1_CD38+ | MIXED | Disc. | -0.145 | 0.003 | -0.238 | -0.049 | -0.024 | 0.621 | -0.121 | 0.072 | 0.017 | 0.891 | * | |  | |
| FlowAbs_Panel2_2263_CD4+.Th1_CD38+ | ALL HIGH | Disc. | -0.173 | 0.007 | -0.294 | -0.047 | -0.065 | 0.316 | -0.191 | 0.063 | 0.033 | 0.749 | * | |  | |
| FlowAbs_Panel2_2263_CD4+.Th1_CD38+ | ALL HIGH | Val. | -0.387 | 0.018 | -0.635 | -0.066 | -0.018 | 0.914 | -0.345 | 0.312 | 0.018 | 0.914 | * | |  | |
| FlowAbs_Panel2_2314_CD4+.Tnaive_HLA-DR+ | MIXED | Val. | 0.127 | 0.313 | -0.122 | 0.362 | -0.325 | 0.008 | -0.529 | -0.086 | 0.313 | 0.008 |  | | ** | |
| FlowAbs_Panel2_2394_CD8+.Tem_CCR4+ | MIXED | Disc. | 0.182 | 0.000 | 0.087 | 0.274 | 0.085 | 0.085 | -0.012 | 0.180 | 0.004 | 0.607 | ** | |  | |
| FlowAbs_Panel2_2408_CD4-CD8+_CCR5+ | MIXED | Disc. | 0.155 | 0.002 | 0.059 | 0.248 | 0.065 | 0.185 | -0.031 | 0.161 | 0.010 | 0.651 | * | |  | |
| FlowAbs_Panel2_2440_CD4+.Th1/17_CCR5+ | ALL LOW | Disc. | -0.142 | 0.008 | -0.245 | -0.037 | -0.011 | 0.836 | -0.117 | 0.095 | 0.036 | 0.981 | * | |  | |
| FlowAbs_Panel2_2452_CD8+.Tem_CCR5+ | MIXED | Disc. | 0.168 | 0.001 | 0.073 | 0.261 | 0.080 | 0.104 | -0.017 | 0.175 | 0.006 | 0.607 | ** | |  | |
| FlowAbs_Panel2_2452_CD8+.Tem_CCR5+ | ALL LOW | Val. | 0.290 | 0.009 | 0.075 | 0.479 | 0.066 | 0.561 | -0.156 | 0.281 | 0.009 | 0.561 | ** | |  | |
| FlowAbs_Panel2_2453_CD8+.Tc1_CCR5+ | MIXED | Disc. | 0.168 | 0.001 | 0.073 | 0.260 | 0.083 | 0.091 | -0.013 | 0.178 | 0.006 | 0.607 | ** | |  | |
| FlowAbs_Panel2_2453_CD8+.Tc1_CCR5+ | ALL LOW | Val. | 0.302 | 0.006 | 0.088 | 0.489 | 0.101 | 0.370 | -0.121 | 0.314 | 0.006 | 0.370 | ** | |  | |
| FlowAbs_Panel2_2467_CD4+CD8+_CCR6+ | ALL LOW | Disc. | -0.191 | 0.000 | -0.292 | -0.087 | -0.025 | 0.640 | -0.131 | 0.081 | 0.004 | 0.891 | ** | |  | |
| FlowAbs_Panel2_2467_CD4+CD8+_CCR6+ | ALL HIGH | Val. | -0.237 | 0.158 | -0.525 | 0.099 | -0.331 | 0.045 | -0.595 | -0.003 | 0.158 | 0.045 |  | | * | |
| FlowAbs_Panel2_2523_CD4+CD8-_CCR7+ | MIXED | Disc. | -0.184 | 0.000 | -0.276 | -0.089 | -0.051 | 0.302 | -0.147 | 0.046 | 0.003 | 0.740 | ** | |  | |
| FlowAbs_Panel2_2523_CD4+CD8-_CCR7+ | ALL HIGH | Disc. | -0.222 | 0.001 | -0.339 | -0.097 | -0.067 | 0.304 | -0.192 | 0.061 | 0.006 | 0.740 | ** | |  | |
| FlowAbs_Panel2_2530_Treg_CCR7+ | MIXED | Disc. | -0.165 | 0.001 | -0.257 | -0.070 | -0.033 | 0.499 | -0.129 | 0.063 | 0.007 | 0.876 | ** | |  | |
| FlowAbs_Panel2_2531_mTreg_CCR7+ | MIXED | Disc. | -0.147 | 0.003 | -0.240 | -0.051 | -0.007 | 0.886 | -0.103 | 0.089 | 0.016 | 0.984 | * | |  | |
| FlowAbs_Panel2_2532_nTreg_CCR7+ | MIXED | Disc. | -0.170 | 0.001 | -0.262 | -0.074 | -0.078 | 0.112 | -0.173 | 0.018 | 0.006 | 0.607 | ** | |  | |
| FlowAbs_Panel2_2549_Tfh_1_CCR7+ | ALL HIGH | Disc. | -0.216 | 0.001 | -0.334 | -0.091 | -0.115 | 0.075 | -0.239 | 0.012 | 0.007 | 0.607 | ** | |  | |
| FlowAbs_Panel2_2553_CD4+.Th1_CCR7+ | MIXED | Disc. | -0.133 | 0.007 | -0.227 | -0.037 | 0.003 | 0.948 | -0.093 | 0.100 | 0.030 | 0.985 | * | |  | |
| FlowAbs_Panel2_2553_CD4+.Th1_CCR7+ | ALL HIGH | Disc. | -0.175 | 0.007 | -0.296 | -0.049 | -0.111 | 0.086 | -0.235 | 0.016 | 0.030 | 0.607 | * | |  | |
| FlowAbs_Panel2_2590_nTreg_CXCR3+ | MIXED | Disc. | -0.161 | 0.001 | -0.254 | -0.066 | -0.088 | 0.073 | -0.183 | 0.008 | 0.007 | 0.607 | ** | |  | |
| FlowAbs_Panel2_2605_CD4+.Temra_CXCR3+ | MIXED | Val. | 0.166 | 0.187 | -0.083 | 0.395 | -0.368 | 0.003 | -0.563 | -0.134 | 0.187 | 0.003 |  | | ** | |
| FlowAbs_Panel2_2639_CD4+CD8-_CXCR4+ | MIXED | Disc. | -0.179 | 0.000 | -0.271 | -0.084 | -0.013 | 0.786 | -0.110 | 0.083 | 0.004 | 0.959 | ** | |  | |
| FlowAbs_Panel2_2646_Treg_CXCR4+ | MIXED | Disc. | -0.157 | 0.001 | -0.250 | -0.062 | 0.011 | 0.830 | -0.086 | 0.107 | 0.009 | 0.981 | ** | |  | |
| FlowAbs_Panel2_2648_nTreg_CXCR4+ | MIXED | Disc. | -0.171 | 0.000 | -0.263 | -0.076 | -0.051 | 0.301 | -0.147 | 0.046 | 0.005 | 0.740 | ** | |  | |
| FlowAbs_Panel2_2661_CD4+.Tcm_CXCR4+ | MIXED | Disc. | -0.129 | 0.009 | -0.223 | -0.033 | 0.002 | 0.967 | -0.094 | 0.099 | 0.036 | 0.985 | * | |  | |
| FlowAbs_Panel2_2661_CD4+.Tcm_CXCR4+ | ALL HIGH | Disc. | -0.177 | 0.006 | -0.298 | -0.051 | -0.074 | 0.256 | -0.199 | 0.054 | 0.029 | 0.719 | * | |  | |
| FlowAbs_Panel2_2662_CD4+.Tnaive_CXCR4+ | MIXED | Disc. | -0.183 | 0.000 | -0.274 | -0.088 | -0.052 | 0.295 | -0.147 | 0.045 | 0.004 | 0.740 | ** | |  | |
| FlowAbs_Panel2_2662_CD4+.Tnaive_CXCR4+ | ALL HIGH | Disc. | -0.187 | 0.004 | -0.306 | -0.061 | -0.049 | 0.450 | -0.175 | 0.079 | 0.020 | 0.856 | * | |  | |
| FlowAbs_Panel2_2670_CD4+.Th2_CXCR4+ | MIXED | Disc. | -0.134 | 0.006 | -0.228 | -0.038 | -0.042 | 0.394 | -0.138 | 0.055 | 0.029 | 0.820 | * | |  | |
| FlowAbs_Panel2_2670_CD4+.Th2_CXCR4+ | ALL HIGH | Val. | -0.339 | 0.040 | -0.600 | -0.011 | -0.145 | 0.393 | -0.452 | 0.193 | 0.040 | 0.393 | * | |  | |
| FlowAbs_Panel2_2697_CD4+CD8-_CXCR5+ | MIXED | Disc. | -0.166 | 0.001 | -0.259 | -0.071 | -0.037 | 0.451 | -0.133 | 0.060 | 0.006 | 0.856 | ** | |  | |
| FlowAbs_Panel2_2719_CD4+.Tcm_CXCR5+ | MIXED | Disc. | -0.128 | 0.009 | -0.221 | -0.031 | -0.015 | 0.763 | -0.111 | 0.082 | 0.039 | 0.954 | * | |  | |
| FlowAbs_Panel2_2719_CD4+.Tcm_CXCR5+ | ALL HIGH | Disc. | -0.161 | 0.013 | -0.283 | -0.035 | -0.091 | 0.161 | -0.216 | 0.037 | 0.049 | 0.651 | * | |  | |
| FlowAbs_Panel2_2720_CD4+.Tnaive_CXCR5+ | MIXED | Disc. | -0.172 | 0.000 | -0.264 | -0.077 | -0.052 | 0.288 | -0.148 | 0.044 | 0.005 | 0.740 | ** | |  | |
| FlowPer_Panel2_2035_Tfh_1 | ALL HIGH | Disc. | -0.179 | 0.005 | -0.300 | -0.054 | -0.088 | 0.177 | -0.213 | 0.040 | 0.026 | 0.651 | * | |  | |
| FlowPer_Panel2_2035_Tfh_1 | MIXED | Val. | -0.214 | 0.087 | -0.437 | 0.034 | 0.277 | 0.025 | 0.034 | 0.490 | 0.087 | 0.025 |  | | * | |
| FlowPer_Panel2_2055_CD4+.Th1/17 | ALL LOW | Disc. | -0.192 | 0.000 | -0.292 | -0.088 | -0.117 | 0.030 | -0.221 | -0.011 | 0.004 | 0.528 | ** | |  | |
| FlowPer_Panel2_2060_CD8+.Tem | MIXED | Disc. | 0.177 | 0.000 | 0.082 | 0.269 | 0.084 | 0.089 | -0.013 | 0.179 | 0.004 | 0.607 | ** | |  | |
| FlowPer_Panel2_2349_CD4+CD8-_CCR4+ | MIXED | Disc. | 0.247 | 0.000 | 0.155 | 0.336 | 0.011 | 0.817 | -0.085 | 0.108 | 0.000 | 0.973 | ** | |  | |
| FlowPer_Panel2_2349_CD4+CD8-_CCR4+ | ALL LOW | Disc. | 0.163 | 0.002 | 0.058 | 0.264 | 0.086 | 0.114 | -0.021 | 0.190 | 0.015 | 0.607 | * | |  | |
| FlowPer_Panel2_2349_CD4+CD8-_CCR4+ | ALL HIGH | Disc. | 0.251 | 0.000 | 0.128 | 0.366 | 0.042 | 0.522 | -0.086 | 0.168 | 0.003 | 0.879 | ** | |  | |
| FlowPer_Panel2_2371_CD4+.Tcm_CCR4+ | MIXED | Disc. | 0.203 | 0.000 | 0.108 | 0.293 | -0.043 | 0.382 | -0.139 | 0.054 | 0.002 | 0.804 | ** | |  | |
| FlowPer_Panel2_2371_CD4+.Tcm_CCR4+ | ALL HIGH | Disc. | 0.214 | 0.001 | 0.089 | 0.332 | 0.036 | 0.581 | -0.092 | 0.162 | 0.007 | 0.891 | ** | |  | |
| FlowPer_Panel2_2371_CD4+.Tcm_CCR4+ | MIXED | Val. | 0.290 | 0.019 | 0.048 | 0.500 | -0.087 | 0.489 | -0.326 | 0.162 | 0.019 | 0.489 | * | |  | |
| FlowPer_Panel2_2374_CD4+.Tem_CCR4+ | MIXED | Disc. | 0.141 | 0.004 | 0.045 | 0.234 | -0.070 | 0.152 | -0.166 | 0.026 | 0.021 | 0.651 | * | |  | |

**Supplementary Table 3 | Flow cytometry results for immune cell populations in the 2000HIV study.**

The tables report flow cytometry data for immune cell subsets in the discovery (n=1,002) and validation (n=189) cohorts, comparing endotypes: 6A (ALL LOW vs. MIXED), 6B (MIXED vs. ALL HIGH), and 6C (ALL LOW vs. ALL HIGH). Each subtable is divided into absolute cell counts (cells/µL) and cell percentages (% of parent population) for immune cell subsets (356 markers), including CD4+ and CD8+ T cells (e.g., Th1, Th2, Th1/17, Tc1, Tc2, Tc17, Tcm, Tem, Tnaive, Temra), T follicular helper cells (Tfh), regulatory T cells (Treg, mTreg, nTreg), NK cells, B cells (e.g., naive, immature, switched memory, plasma cells), and non-classical monocytes, with markers such as PD1, CD38, HLA-DR, CCR4, CCR5, CCR6, CCR7, CXCR3, CXCR4, CXCR5, and CD81. Data were inverse rank-transformed using the qnorm function in R (v4.3.0) to achieve normality and analyzed via linear regression, adjusting for confounders selected through a two-step process: (1) principal component analysis (PCA) on raw omics data using FactoMineR (v2.4), with the first 10 principal components regressed against potential confounders (age, sex, ethnicity [first genetic principal component], collection center, seasonality [sine/cosine functions], time to laboratory preprocessing, COVID-19 vaccination status, HIV-related variables [e.g., cART duration, CD4 nadir]); (2) confounders causing >10% change in beta coefficients for omics-clinical associations were selected (age, sex, seasonality, collection center, time to laboratory preprocessing, first genetic principal component). Estimates represent standardized beta coefficients from linear regression. P-values are nominal (P<0.05) for the validation cohort and FDR-adjusted (P<0.05) for the discovery cohort, with false discovery rate (FDR) values reported for multiple testing correction. All analyses were performed in R (v4.3.0). cART, combination antiretroviral therapy; FDR, false discovery rate. For details, see **Methods.** Only markers with nominal P-values < 0.05 in either sub-cohorts are presented below.

**Supplementary Table 3A:** *All Low* vs *All High* (Main difference: Reservoir Total and Intactness)

*Absolute Cell Counts*

| **Cell Population** | **Estimate** | **P-value** | **FDR** | **Cohort** |
| --- | --- | --- | --- | --- |
| Panel2_2018_CD4+CD8- | -0.349386527 | 2.65156E-05 | 0.000811077 | Discovery |
| Panel2_2060_CD8+.Tem | 0.352891306 | 3.27862E-05 | 0.000811077 | Discovery |
| Panel2_2051_CD4+.Th1 | -0.349704797 | 3.42709E-05 | 0.000811077 | Discovery |
| Panel2_2035_Tfh_1 | -0.332762761 | 8.59304E-05 | 0.001331236 | Discovery |
| Panel2_2025_nTreg | -0.293065547 | 0.000134203 | 0.001639537 | Discovery |
| Panel2_2055_CD4+.Th1/17 | -0.326889075 | 0.000134234 | 0.001639537 | Discovery |
| Panel2_2074_CD8+.Tc17 | 0.305823963 | 0.000194771 | 0.001819572 | Discovery |
| Panel1_1031_NK_CD56-CD16+ | 0.289421993 | 0.000637547 | 0.004437826 | Discovery |
| Panel2_2046_CD4+.Tnaive | -0.269889208 | 0.000652143 | 0.004452128 | Discovery |
| Panel2_2021_Treg | -0.27666414 | 0.000717029 | 0.004676758 | Discovery |
| Panel2_2019_CD4+CD8+ | -0.274311565 | 0.001216162 | 0.007317585 | Discovery |
| Panel2_2017_CD4-CD8+ | 0.274549471 | 0.001300878 | 0.007696862 | Discovery |
| Panel2_2069_CD8+.Tc1 | 0.266877746 | 0.001721357 | 0.009284021 | Discovery |
| Panel3_3039_Immature.B.cells | 0.267904864 | 0.001726043 | 0.009284021 | Discovery |
| Panel2_2043_CD4+.Tcm | -0.244328141 | 0.003616509 | 0.016459751 | Discovery |
| Panel2_2070_CD8+.Tc2 | 0.222110364 | 0.00692577 | 0.027318315 | Discovery |
| Panel2_2052_CD4+.Th2 | -0.220852121 | 0.007099643 | 0.027696408 | Discovery |
| Panel3_3040_Naive.B.cells | 0.215547334 | 0.011691056 | 0.041838242 | Discovery |
| Panel2_2024_mTreg | -0.208187354 | 0.013382291 | 0.04612343 | Discovery |
| Panel2_2045_CD4+.Temra | -0.205261357 | 0.017156302 | 0.056920443 | Discovery |
| Panel2_2039_Tfh_1/17 | -0.196865611 | 0.020174073 | 0.061439555 | Discovery |
| Panel3_3028_Unswitched.Memory.B.cells | 0.190794505 | 0.026327934 | 0.074998798 | Discovery |
| Panel2_2044_CD4+.Tem | -0.187837013 | 0.030580112 | 0.083507229 | Discovery |
| Panel3_3010_CD19+ | 0.184732987 | 0.033098624 | 0.089015239 | Discovery |
| Panel2_2051_CD4+.Th1 | -0.515350437 | 0.012128017 | 0.777987986 | Validation |
| Panel3_3040_Naive.B.cells | 0.457871589 | 0.026967239 | 0.777987986 | Validation |

*Cell Percent*

| **Cell Population** | **Estimate** | **P-value** | **FDR** | **Cohort** |
| --- | --- | --- | --- | --- |
| Panel2_2018_CD4+CD8- | -0.500689971 | 8.48254E-10 | 1.32143E-07 | Discovery |
| Panel2_2017_CD4-CD8+ | 0.503594196 | 1.1167E-09 | 1.32143E-07 | Discovery |
| Panel2_2349_CD4+CD8-_CCR4+ | 0.420691011 | 3.09695E-07 | 2.19884E-05 | Discovery |
| Panel2_2374_CD4+.Tem_CCR4+ | 0.528266119 | 5.51354E-10 | 1.32143E-07 | Discovery |
| Panel2_2055_CD4+.Th1/17 | -0.462132093 | 2.16208E-08 | 1.91885E-06 | Discovery |
| Panel2_2035_Tfh_1 | -0.336832164 | 3.84184E-05 | 0.001704818 | Discovery |
| Panel2_2056_CD4+.Th17 | 0.281254317 | 0.000588853 | 0.018723487 | Discovery |
| Panel2_2019_CD4+CD8+ | -0.285228486 | 0.000611678 | 0.018723487 | Discovery |
| Panel2_2060_CD8+.Tem | 0.270999456 | 0.000818252 | 0.018723487 | Discovery |
| Panel2_2039_Tfh_1/17 | -0.258672975 | 0.00111489 | 0.019290865 | Discovery |
| Panel2_2051_CD4+.Th1 | -0.265126304 | 0.001802568 | 0.025596463 | Discovery |
| Panel1_1031_NK_CD56-CD16+ | 0.244943351 | 0.004099103 | 0.040883586 | Discovery |
| Panel2_2036_Tfh_2 | 0.231416353 | 0.004376271 | 0.040883586 | Discovery |
| Panel3_3029_Switched.Memory.B.cells | -0.224757288 | 0.008613493 | 0.069495231 | Discovery |
| Panel3_3010_CD19+ | 0.205903701 | 0.015098592 | 0.101132078 | Discovery |
| Panel3_3039_Immature.B.cells | 0.201702594 | 0.01659344 | 0.106549974 | Discovery |
| Panel2_2062_CD8+.Tnaive | -0.147476162 | 0.036427349 | 0.155803721 | Discovery |
| Panel2_2015_Tcells | -0.166499359 | 0.03847614 | 0.160969321 | Discovery |
| Panel2_2025_nTreg | -0.15778862 | 0.03944882 | 0.160969321 | Discovery |
| Panel2_2024_mTreg | 0.15778862 | 0.03944882 | 0.160969321 | Discovery |
| Panel2_2349_CD4+CD8-_CCR4+ | 0.616751261 | 0.003591147 | 0.424952428 | Validation |
| Panel2_2374_CD4+.Tem_CCR4+ | 0.619022823 | 0.002644051 | 0.424952428 | Validation |
| Panel3_3017_Plasma.Cells | -0.54917781 | 0.008148982 | 0.587334441 | Validation |
| Panel2_2036_Tfh_2 | 0.494807585 | 0.01094362 | 0.587334441 | Validation |
| Panel1_1018_IM_CD14++CD16+ | -0.495794654 | 0.011581243 | 0.587334441 | Validation |
| Panel2_2052_CD4+.Th2 | 0.493072156 | 0.024815841 | 0.694748521 | Validation |
| Panel2_2035_Tfh_1 | -0.428684738 | 0.025441495 | 0.694748521 | Validation |
| Panel2_2051_CD4+.Th1 | -0.389394374 | 0.042708313 | 0.828187433 | Validation |
| Panel3_3010_CD19+ | 0.422354696 | 0.043217454 | 0.828187433 | Validation |
| Panel2_2025_nTreg | -0.351397499 | 0.048991369 | 0.828187433 | Validation |
| Panel2_2024_mTreg | 0.351397499 | 0.048991369 | 0.828187433 | Validation |

**Supplementary Table 3B:** *All Low* vs *Mixed* (Main difference: Reservoir Total)

*Absolute Cell Counts*

| **Cell Population** | **Estimate** | **P-value** | **FDR** | **Cohort** |
| --- | --- | --- | --- | --- |
| Panel2_2060_CD8+.Tem | 0.302129486 | 5.19926E-05 | 0.002150399 | Discovery |
| Panel2_2069_CD8+.Tc1 | 0.287815721 | 0.000103201 | 0.00252484 | Discovery |
| Panel2_2025_nTreg | -0.265425025 | 0.000121608 | 0.002608193 | Discovery |
| Panel2_2018_CD4+CD8- | -0.261193518 | 0.000349034 | 0.004260384 | Discovery |
| Panel2_2046_CD4+.Tnaive | -0.248682028 | 0.000372034 | 0.004260384 | Discovery |
| Panel2_2052_CD4+.Th2 | -0.224758823 | 0.002025217 | 0.015679378 | Discovery |
| Panel2_2021_Treg | -0.22413054 | 0.002031694 | 0.015679378 | Discovery |
| Panel2_2035_Tfh_1 | -0.21677204 | 0.003150196 | 0.019969991 | Discovery |
| Panel2_2073_CD8+.Tc1/17 | 0.219796579 | 0.003230188 | 0.020117839 | Discovery |
| Panel2_2017_CD4-CD8+ | 0.213612449 | 0.004149812 | 0.023383859 | Discovery |
| Panel2_2051_CD4+.Th1 | -0.21185113 | 0.004290151 | 0.023557705 | Discovery |
| Panel2_2019_CD4+CD8+ | -0.211123469 | 0.005108102 | 0.026280813 | Discovery |
| Panel1_1049_TCRvd2 | -0.184175463 | 0.012345241 | 0.054105686 | Discovery |
| Panel2_2024_mTreg | -0.165921343 | 0.024123077 | 0.092484077 | Discovery |
| Panel2_2043_CD4+.Tcm | -0.166955855 | 0.024549321 | 0.092506139 | Discovery |
| Panel2_2055_CD4+.Th1/17 | -0.153112745 | 0.043593256 | 0.145996283 | Discovery |
| Panel2_2046_CD4+.Tnaive | -0.41923944 | 0.01268988 | 0.505288809 | Validation |
| Panel2_2025_nTreg | -0.351286408 | 0.027401522 | 0.590922501 | Validation |

*Cell Percent*

| **Cell Population** | **Estimate** | **P-value** | **FDR** | **Cohort** |
| --- | --- | --- | --- | --- |
| Panel2_2017_CD4-CD8+ | 0.370145136 | 4.3843E-07 | 0.000142382 | Discovery |
| Panel2_2018_CD4+CD8- | -0.360385926 | 8.02154E-07 | 0.000142382 | Discovery |
| Panel2_2060_CD8+.Tem | 0.296251974 | 4.27497E-05 | 0.00216802 | Discovery |
| Panel2_2073_CD8+.Tc1/17 | 0.288374622 | 0.000115854 | 0.005141029 | Discovery |
| Panel2_2062_CD8+.Tnaive | -0.22841569 | 0.000333467 | 0.010063587 | Discovery |
| Panel2_2019_CD4+CD8+ | -0.232416384 | 0.001843686 | 0.038500494 | Discovery |
| Panel2_2055_CD4+.Th1/17 | -0.211422598 | 0.003961491 | 0.070316464 | Discovery |
| Panel1_1031_NK_CD56-CD16+ | 0.213907284 | 0.004998712 | 0.084502038 | Discovery |
| Panel2_2035_Tfh_1 | -0.192859719 | 0.006803552 | 0.095679613 | Discovery |
| Panel1_1049_TCRvd2 | -0.185480357 | 0.010771429 | 0.117631227 | Discovery |
| Panel2_2025_nTreg | -0.162796506 | 0.016682296 | 0.141005121 | Discovery |
| Panel2_2024_mTreg | 0.162796506 | 0.016682296 | 0.141005121 | Discovery |
| Panel2_2070_CD8+.Tc2 | -0.155950191 | 0.031843678 | 0.207314516 | Discovery |
| Panel3_3029_Switched.Memory.B.cells | -0.151520545 | 0.043803122 | 0.259168471 | Discovery |
| Panel2_2060_CD8+.Tem | 0.406051735 | 0.004308173 | 0.58644808 | Validation |
| Panel2_2046_CD4+.Tnaive | -0.364463311 | 0.017030279 | 0.671749894 | Validation |
| Panel2_2044_CD4+.Tem | 0.348202419 | 0.027648525 | 0.794321014 | Validation |
| Panel2_2024_mTreg | 0.289139137 | 0.040275432 | 0.794321014 | Validation |
| Panel2_2025_nTreg | -0.289139137 | 0.040275432 | 0.794321014 | Validation |

**Supplementary Table 3C:** *Mixed* vs *All High* (Main difference: Reservoir Intactness)

*Absolute Cell Counts*

| **Cell Population** | **Estimate** | **P-value** | **FDR** | **Cohort** |
| --- | --- | --- | --- | --- |
| Panel1_1047_TCRvd1 | -0.238849803 | 0.004061468 | 0.360455274 | Discovery |
| Panel2_2014_NKcells | 0.228256332 | 0.005222324 | 0.370785024 | Discovery |
| Panel2_2074_CD8+.Tc17 | 0.195799828 | 0.013828989 | 0.655804802 | Discovery |
| Panel3_3039_Immature.B.cells | 0.191908058 | 0.01487585 | 0.655804802 | Discovery |
| Panel1_1031_NK_CD56-CD16+ | 0.176402174 | 0.030386695 | 0.666044347 | Discovery |
| Panel2_2070_CD8+.Tc2 | 0.159020255 | 0.046627114 | 0.666044347 | Discovery |
| Panel2_2069_CD8+.Tc1 | -0.464802969 | 0.019315683 | 0.978944992 | Validation |

*Cell Percent*

| **Cell Population** | **Estimate** | **P-value** | **FDR** | **Cohort** |
| --- | --- | --- | --- | --- |
| Panel2_2073_CD8+.Tc1/17 | -0.262218114 | 0.001599698 | 0.189842294 | Discovery |
| Panel2_2055_CD4+.Th1/17 | -0.254851989 | 0.001804258 | 0.189842294 | Discovery |
| Panel1_1047_TCRvd1 | -0.246336595 | 0.002496141 | 0.189842294 | Discovery |
| Panel1_1027_DC | -0.223350613 | 0.005973853 | 0.259461075 | Discovery |
| Panel2_2014_NKcells | 0.221446107 | 0.006577886 | 0.259461075 | Discovery |
| Panel2_2051_CD4+.Th1 | -0.198660768 | 0.016793429 | 0.427515926 | Discovery |
| Panel2_2074_CD8+.Tc17 | 0.186141859 | 0.023733314 | 0.427515926 | Discovery |
| Panel1_1028_Basophils | -0.186790557 | 0.02458383 | 0.427515926 | Discovery |
| Panel2_2013_CD56+CD3+ | -0.180184379 | 0.029510182 | 0.427515926 | Discovery |
| Panel2_2015_Tcells | -0.170484865 | 0.031642797 | 0.427515926 | Discovery |
| Panel2_2051_CD4+.Th1 | -0.463715954 | 0.027551155 | 0.750621622 | Validation |
| Panel2_2062_CD8+.Tnaive | 0.351915952 | 0.033599169 | 0.750621622 | Validation |
| Panel2_2052_CD4+.Th2 | 0.409457276 | 0.040660687 | 0.750621622 | Validation |
| Panel2_2060_CD8+.Tem | -0.3789451 | 0.045276788 | 0.750621622 | Validation |
| Panel2_2036_Tfh_2 | 0.402800972 | 0.048493513 | 0.750621622 | Validation |

**Supplementary Table 4 | Bulk transcriptomics results for differentially expressed genes in the 2000HIV study**.

The table reports gene expression data from unstimulated peripheral blood mononuclear cells (PBMCs) in the discovery (n=1,002) and validation (n=189) cohorts, comparing endotypes: 7A (ALL LOW vs. MIXED), 7B (MIXED vs. ALL HIGH), and 7C (ALL LOW vs. ALL HIGH). Each subtable includes genes with validated direction and significance, listing Ensembl gene ID (GENEID), gene symbol (SYMBOL), gene type (e.g., protein coding, processed pseudogene, TEC), chromosome (CHR), base mean expression, log2 fold change (Log2 FC), standard error of log2 fold change (lfcSE), nominal P-value (pvalue), and false discovery rate (FDR)-adjusted P-value (padj). RNA-seq data (58,347 genes) were analyzed using DESeq2 (v1.34.0) in R (v4.3.0) with negative binomial models, Benjamini–Hochberg correction, and apeglm shrinkage. Models were adjusted for confounders selected through a two-step process: (1) principal component analysis (PCA) on raw omics data using FactoMineR (v2.4), with the first 10 principal components regressed against potential confounders (age, sex, ethnicity [first genetic principal component], collection center, seasonality [sine/cosine functions], time to laboratory preprocessing, COVID-19 vaccination status, HIV-related variables [e.g., cART duration, CD4 nadir]); (2) confounders causing >10% change in beta coefficients for omics-clinical associations were selected (age, sex, seasonality, collection center, time to laboratory preprocessing, first genetic principal component). Differentially expressed genes required FDR-adjusted P<0.05 in the discovery cohort, nominal P<0.05 in the validation cohort, and consistent log2 fold change direction. cART, combination antiretroviral therapy; FDR, false discovery rate; TEC, to be experimentally confirmed. See **Methods** for details.

**Supplementary Table 4A:** *ALL LOW vs ALL HIGH (Main difference: Reservoir Total and Intactness)*

| GENEID | SYMBOL | GENE TYPE | CHR | Base Mean | Log2 FC | lfcSE | pvalue | padj | Cohort |
| --- | --- | --- | --- | --- | --- | --- | --- | --- | --- |
| ENSG00000115484.14 | CCT4 | Protein coding | 2 | 598.8511 | -0.030409 | 0.011374 | 0.002702 | 0.055173 | Discovery |
| ENSG00000138386.16 | NAB1 | Protein coding | 2 | 197.5068 | 0.066506 | 0.023963 | 0.000490 | 0.022875 | Discovery |
| ENSG00000186310.9 | NAP1L3 | Protein coding | X | 32.53048 | -0.25190 | 0.049478 | 1.38062E-08 | 2.66827E-05 | Discovery |
| ENSG00000217165.1 | ANKRD18EP | Processed pseudogene | 6 | 85.3828 | -0.038181 | 0.034568 | 0.031831 | 0.206361 | Discovery |
| ENSG00000280135.1 | AL096816.1 | TEC | 6 | 323.8889 | -0.038071 | 0.035850 | 0.032666 | 0.208780 | Discovery |
| ENSG00000115484.14 | CCT4 | Protein coding | 2 | 602.0861 | -0.083556 | 0.030583 | 0.000198 | 0.485541 | Validation |
| ENSG00000138386.16 | NAB1 | Protein coding | 2 | 207.4730 | 0.07489 |  | 0.004714 | 0.945682 | Validation |
| ENSG00000186310.9 | NAP1L3 | Protein coding | X | 33.23983 | -0.252923 | 0.148049 | 0.001859 | 0.945682 | Validation |
| ENSG00000217165.1 | ANKRD18EP | Processed pseudogene | 6 | 83.6642 | -0.175267 | 0.097366 | 0.001657 | 0.945682 | Validation |
| ENSG00000280135.1 | AL096816.1 | TEC | 6 | 299.3286 | -0.079884 |  | 0.005758 | 0.945682 | Validation |

**Supplementary Table 4B:** *ALL LOW vs MIXED (Main difference: Reservoir Total)*

| GENEID | SYMBOL | GENE TYPE | CHR | Base Mean | Log2 FC | lfcSE | pvalue | padj | Cohort |
| --- | --- | --- | --- | --- | --- | --- | --- | --- | --- |
| ENSG00000100628.11 | ASB2 | Protein coding | 14 | 21.1330 | 0.12264 | 0.06552 | 0.00141 | 0.21093 | Discovery |
| ENSG00000100628.11 | ASB2 | Protein coding | 14 | 20.4115 | 0.26369 | 0.12067 | 0.00070 | 0.42915 | Validation |

**Supplementary Table 4C:** *MIXED vs ALL HIGH (Main difference: Reservoir Intactness)*

No validated markers in validation and significance

**Supplementary Table 5 | Gene Set Enrichment Analysis (GSEA) for genes in the 2000HIV study discovery sub-cohort**.

**Supplementary Table 5A:** *ALL LOW vs ALL HIGH (Main difference: Reservoir Total and Intactness)*

| **ID** | **Description** | **Set Size** | **Enrichment Score (ES)** | **Normalized Enrichment Score (NES)** | **P-Value** | **Adjusted P-Value** | **Genes** |
| --- | --- | --- | --- | --- | --- | --- | --- |
| R-HSA-909733 | Interferon alpha/beta signaling | 63 | 6.43E-01 | 2.05 | 1.25E-06 | 5.03E-05 | IFI27; USP18; IFIT1; TYK2; RSAD2; IFITM2; MX2; ISG20; XAF1; IRF2; OAS1; IFIT3; HLA-C; OAS2; ISG15; IFI6; OAS3; STAT2; IFNAR1; IFITM1; OASL; EIF2AK2 |
| R-HSA-9664407 | Parasite infection | 57 | 6.19E-01 | 1.94 | 4.32E-05 | 1.21E-03 | MYO1C; MYO9B; VAV3; NCKAP1L; ARPC2; MAPK1; VAV2; HCK; CRK; VAV1; WIPF1; BAIAP2; ACTR2; WAS; BTK; MYH9; ARPC5; MAPK3; ARPC1B; CD3G; ABL1; FGR; SRC; LYN; SYK |
| R-HSA-9664417 | Leishmania phagocytosis | 57 | 6.19E-01 | 1.94 | 4.32E-05 | 1.21E-03 | MYO1C; MYO9B; VAV3; NCKAP1L; ARPC2; MAPK1; VAV2; HCK; CRK; VAV1; WIPF1; BAIAP2; ACTR2; WAS; BTK; MYH9; ARPC5; MAPK3; ARPC1B; CD3G; ABL1; FGR; SRC; LYN; SYK |
| R-HSA-9664422 | FCGR3A-mediated phagocytosis | 57 | 6.19E-01 | 1.94 | 4.32E-05 | 1.21E-03 | MYO1C; MYO9B; VAV3; NCKAP1L; ARPC2; MAPK1; VAV2; HCK; CRK; VAV1; WIPF1; BAIAP2; ACTR2; WAS; BTK; MYH9; ARPC5; MAPK3; ARPC1B; CD3G; ABL1; FGR; SRC; LYN; SYK |
| R-HSA-9768759 | Regulation of NPAS4 gene expression | 11 | 8.74E-01 | 1.90 | 3.98E-05 | 1.18E-03 | AGO1; AGO2; AGO4 |
| R-HSA-2029480 | Fcgamma receptor (FCGR) dependent phagocytosis | 82 | 5.55E-01 | 1.86 | 5.03E-05 | 1.39E-03 | MYO1C; LIMK1; PLCG2; MYO9B; VAV3; NCKAP1L; ARPC2; MAPK1; VAV2; AHCYL1; ITPR1; HCK; CRK; VAV1; FCGR2A; WIPF1; BAIAP2; PLPP5; ACTR2; WAS; BTK; MYH9; ARPC5; MAPK3; ARPC1B; CD3G; ABL1; FGR; SRC; PRKCE; LYN; SYK |
| R-HSA-187687 | Signalling to ERKs | 31 | 6.47E-01 | 1.83 | 6.61E-04 | 1.51E-02 | RAPGEF1; MAPK1; CRK; MAPK14; SOS1; NTRK1; MAPK3; RALA; SRC; RALB; KIDINS220; MAP2K2; MAPK12; KRAS; RAP1A; SHC2; GRB2 |
| R-HSA-2029482 | Regulation of actin dynamics for phagocytic cup formation | 59 | 5.75E-01 | 1.80 | 3.72E-04 | 9.37E-03 | MYO1C; LIMK1; MYO9B; VAV3; NCKAP1L; ARPC2; MAPK1; VAV2; CRK; VAV1; FCGR2A; WIPF1; BAIAP2; ACTR2; WAS; BTK; MYH9; ARPC5; MAPK3; ARPC1B; CD3G; ABL1 |
| R-HSA-983695 | Antigen activates B Cell Receptor (BCR) leading to generation of second messengers | 31 | 6.37E-01 | 1.80 | 9.88E-04 | 2.12E-02 | ORAI2; PLCG2; CD79A; CD79B; DAPP1; CD19; AHCYL1; PIK3AP1; ITPR1; VAV1; BLNK; BLK; CD22; SOS1; BTK |
| R-HSA-450282 | MAPK targets/ Nuclear events mediated by MAP kinases | 31 | 6.31E-01 | 1.79 | 1.16E-03 | 2.33E-02 | MEF2C; MAPK1; RPS6KA1; MEF2A; MAPK9; MAPK14; MAPK3; ATF1; VRK3; RPS6KA3; DUSP4 |
| R-HSA-198753 | ERK/MAPK targets | 22 | 6.81E-01 | 1.78 | 1.09E-03 | 2.20E-02 | MEF2C; MAPK1; RPS6KA1; MEF2A; MAPK14; MAPK3; VRK3; RPS6KA3; DUSP4 |
| R-HSA-162658 | Golgi Cisternae Pericentriolar Stack Reorganization | 14 | 7.65E-01 | 1.78 | 1.07E-03 | 2.20E-02 | GOLGA2; MAPK1; PLK1; RAB1B; BLZF1; CCNB2; MAPK3 |
| R-HSA-418990 | Adherens junctions interactions | 32 | 6.19E-01 | 1.76 | 1.43E-03 | 2.71E-02 | AGO1; AGO2; AGO4; CTNND1; SP1; NECTIN1; ADAM19; ZEB2 |
| R-HSA-1793185 | Chondroitin sulfate/dermatan sulfate metabolism | 37 | 6.05E-01 | 1.76 | 1.96E-03 | 3.62E-02 | CHST15; SDC2; CHST12; CHPF2; IDS; AGRN; CHSY1; DSE; B3GALT6; B3GAT1; CHST13; CHST11; CHST14 |
| R-HSA-451927 | Interleukin-2 family signaling | 34 | 6.10E-01 | 1.75 | 1.99E-03 | 3.64E-02 | INPPL1; HAVCR2; PTK2B; INPP5D; PIK3R3; STAT5B; SOS1; IL2RB; SYK; SOS2; LGALS9; IL21R; IL2RG; GAB2; CSF2RB; PTPN6; GRB2; STAT5A |
| R-HSA-9759476 | Regulation of Homotypic Cell-Cell Adhesion | 21 | 6.73E-01 | 1.74 | 2.40E-03 | 4.04E-02 | AGO1; AGO2; AGO4; CTNND1; SP1; ADAM19; ZEB2 |
| R-HSA-9764260 | Regulation of Expression and Function of Type II Classical Cadherins | 21 | 6.73E-01 | 1.74 | 2.40E-03 | 4.04E-02 | AGO1; AGO2; AGO4; CTNND1; SP1; ADAM19; ZEB2 |
| R-HSA-9658195 | Leishmania infection | 142 | 4.90E-01 | 1.74 | 3.36E-05 | 1.04E-03 | DVL3; MYO1C; PLCG2; ADCY4; RHBDF2; MYO9B; VAV3; NCKAP1L; ARPC2; PRKAR1A; MAPK1; CYBA; VAV2; GNG2; AHCYL1; GNB1; ITPR1; HCK; NFKB2; CRK; VAV1; MAPK14; C3AR1; FCGR2A; ADCY7; WIPF1; BAIAP2; MEFV; ACTR2; WAS; GNGT2; BTK; MYH9; ARPC5; MAPK3; ARPC1B; CD3G; FURIN; ABL1; FGR; DPEP3; SRC; GNG7; LYN; SYK |
| R-HSA-9824443 | Parasitic Infection Pathways | 142 | 4.90E-01 | 1.74 | 3.36E-05 | 1.04E-03 | DVL3; MYO1C; PLCG2; ADCY4; RHBDF2; MYO9B; VAV3; NCKAP1L; ARPC2; PRKAR1A; MAPK1; CYBA; VAV2; GNG2; AHCYL1; GNB1; ITPR1; HCK; NFKB2; CRK; VAV1; MAPK14; C3AR1; FCGR2A; ADCY7; WIPF1; BAIAP2; MEFV; ACTR2; WAS; GNGT2; BTK; MYH9; ARPC5; MAPK3; ARPC1B; CD3G; FURIN; ABL1; FGR; DPEP3; SRC; GNG7; LYN; SYK |
| R-HSA-933541 | TRAF6 mediated IRF7 activation | 16 | 7.23E-01 | 1.74 | 2.42E-03 | 4.04E-02 | RIGI; TRIM25; TANK; IFIH1; TBK1; IRF7; TRAF2 |
| R-HSA-9759475 | Regulation of CDH11 Expression and Function | 20 | 6.81E-01 | 1.73 | 2.44E-03 | 4.04E-02 | AGO1; AGO2; AGO4; CTNND1; SP1; ADAM19; ZEB2 |
| R-HSA-1630316 | Glycosaminoglycan metabolism | 90 | 5.13E-01 | 1.73 | 3.36E-04 | 8.75E-03 | CHST15; SLC35B3; SDC2; GLB1L2; CHST12; CHPF2; SLC9A1; IDS; AGRN; B4GALT5; CHST2; ST3GAL4; ST3GAL6; NDST1; CHSY1; HS3ST1; DSE; B3GALT6; B3GAT1; HGSNAT; ST3GAL2; GLB1; CHST13; B4GAT1; CHST11; HS6ST1; HPSE; B4GALT2; CHST14; GLB1L3; HYAL2 |
| R-HSA-198933 | Immunoregulatory interactions between a Lymphoid and a non-Lymphoid cell | 114 | 4.98E-01 | 1.73 | 1.71E-04 | 4.61E-03 | CD8A; NPDC1; SIGLEC12; KLRD1; TREML2; CD81; CD19; SLAMF6; CD8B; SIGLEC1; KLRK1; KLRK1; HLA-C; OSCAR; CD200; ITGA4; KLRC1; FCGR2B; IFITM1; SH2D1B; ITGB7; CD40; SELL; CD22; SIGLEC6; NCR1; LAIR1; CD3G; ITGAL; KIR2DL3; CD300A; SLAMF7; LILRB1; LILRA5; VCAM1; SIGLEC10; LILRB2; MICB |
| R-HSA-2022870 | Chondroitin sulfate biosynthesis | 11 | 7.89E-01 | 1.72 | 2.09E-03 | 3.77E-02 | CHST15; CHST12; CHPF2; CHSY1; CHST13; CHST11 |
| R-HSA-375276 | Peptide ligand-binding receptors | 81 | 5.10E-01 | 1.70 | 9.48E-04 | 2.06E-02 | ACKR3; CCR2; CX3CR1; CXCR2; PROK2; CCR9; FPR2; F2R; CXCL16; C3AR1; FPR1; NMUR1; CCR5; CXCR1; OPRL1; UTS2; ECE1; CXCL1; CXCL2; CCL5; EDN3; PNOC; C5AR2 |
| R-HSA-9833110 | RSV-host interactions | 79 | 5.09E-01 | 1.69 | 1.27E-03 | 2.47E-02 | CX3CR1; TYK2; SDC2; TLR6; MED22; MED12; AGRN; OAS2; RIGI; MED11; ISG15; MED20; STAT2; IFNAR1; TLR4; TRIM25; EIF2AK2; HERC5; IFIH1; MED13; MED25; MED8; MED14; MED1; MED15; CDK19; TLR7 |
| R-HSA-6798695 | Neutrophil degranulation | 448 | 4.29E-01 | 1.69 | 1.57E-08 | 6.54E-07 | OLR1; ADAM8; LPCAT1; NBEAL2; ALOX5; ITGAV; ITGAX; SLCO4C1; CXCR2; FCGR3B; RAB27A; P2RX1; RAB5B; DBNL; NCKAP1L; HSPA6; PTPRJ; IDH1; CD58; CEACAM3; FPR2; CD53; MAPK1; CYBA; ADGRG3; HVCN1; NCSTN; SLC2A5; IGF2R; PSEN1; IQGAP1; PECAM1; DOK3; DNAJC5; PTPRN2; HLA-C; OSCAR; NHLRC3; TNFAIP6; MMP25; FRMPD3; MAPK14; TNFRSF1B; VAT1; ARHGAP9; APAF1; OSTF1; MGAM; MNDA; PFKL; C3AR1; RAB10; FCGR2A; ADGRE3; FPR1; TCIRG1; ORM1; TMEM30A; CXCR1; RAB37; MME; SIRPA; IRAG2; TRAPPC1; CEACAM1; MMP9; CDA; ACLY; CHIT1; SELL; HGSNAT; TIMP2; FABP5; FTH1; UNC13D; NFAM1; ACTR2; SLC11A1; VNN1; ARPC5; KCNAB2; LAIR1; PADI2; GLB1; MLEC; SLC44A2; CR1; SNAP23; ITGAL; CYSTM1; TMT1A; CAB39; CNN2; CXCL1; CD300A; LAMP1; ACTR10; HPSE; TMEM179B; AMPD3; RAB31; FGR; SURF4; SYNGR1; DYNC1H1; FGL2; LRG1; QSOX1; MOSPD2; SIRPB1; GCA; CTSC; MCEMP1; ERP44; CSNK2B; MANBA; UBR4; CAP1; LILRB2; DIAPH1; CHI3L1; CLEC4D |
| R-HSA-416476 | G alpha (q) signalling events | 117 | 4.61E-01 | 1.60 | 1.22E-03 | 2.40E-02 | P2RY6; GPR132; TRIO; PROK2; FPR2; F2R; GNRH1; MAPK1; RPS6KA1; RGS3; LPAR5; GNG2; GNB1; ITPR1; PLCB3; PIK3R3; FFAR2; RGS19; ABHD6; NMUR1; UTS2; SOS1; TRPC3; GNGT2; BTK; MAPK3; RGS13; GRK2; DGKD; GNG7; EDN3; PRKCE; RPS6KA3 |
| R-HSA-877300 | Interferon gamma signaling | 84 | 4.77E-01 | 1.60 | 3.07E-03 | 4.72E-02 | TRIM38; MAPK1; TRIM68; IRF2; TRIM5; OAS1; SP100; HLA-C; OAS2; GBP3; OAS3; GBP4; CIITA; TRIM25; OASL; TRIM10; MAPK3; IRF7; VCAM1; TRIM21; RAF1; PTPN11; GBP5; TRIM26; IRF9; TRIM17; PML; HLA-E; PTPN6; TRIM22; TRIM8 |
| R-HSA-400206 | Regulation of lipid metabolism by PPARalpha | 105 | 4.64E-01 | 1.60 | 2.25E-03 | 3.95E-02 | PPARA; MTF1; FADS1; ABCB4; HELZ2; ESRRA; MED22; MED12; RXRA; SP1; RXRB; ARNT; MED11; AHRR; MED20; ACADM; SREBF2; CHD9; ACSL1; TBL1XR1; CPT1A; MED13; CPT2; MED25; MED8; MED14; TIAM2; PPARG; MED1; APOA2; ACOX1; MED15; G0S2; CDK19; HDAC3; NFYC; NCOR1; NCOA3; NCOR2; SLC27A1; TBL1X; MED18; ALAS1; MED29; PEX11A |
| R-HSA-373076 | Class A/1 (Rhodopsin-like receptors) | 146 | 4.43E-01 | 1.58 | 1.06E-03 | 2.20E-02 | ACKR3; P2RY6; CCR2; CX3CR1; CNR1; CXCR2; GPR132; PROK2; CCR9; FPR2; F2R; GNRH1; LPAR5; CXCL16; FFAR2; C3AR1; FPR1; NMUR1; CCR5; CXCR1; P2RY14; OPRL1; UTS2; CNR2; ECE1; CMKLR1; CXCL1; CXCL2; CCL5 |
| R-HSA-418594 | G alpha (i) signalling events | 175 | 4.18E-01 | 1.53 | 1.38E-03 | 2.65E-02 | ACKR3; ADCY4; CCR2; CX3CR1; GNAL; CNR1; CXCR2; CCR9; FPR2; PRKAR1A; PDE4A; MAPK1; RGS3; LPAR5; GNG2; AHCYL1; GNB1; ITPR1; RGS9; PLCB3; PPP1CA; CXCL16; C3AR1; FPR1; RGS19; ADCY7; NMUR1; CCR5; CXCR1; P2RY14; OPRL1; PPP3CA; CNR2; CAMKK2; GNGT2; RGS13; GRK2; CXCL1; CXCL2; CCL5; SRC; GNG7; CAMKK1; PNOC; RGS14 |
| R-HSA-913531 | Interferon Signaling | 233 | 4.06E-01 | 1.52 | 7.21E-04 | 1.62E-02 | IFI27; USP18; IFIT1; TRIM38; TYK2; RSAD2; IFITM2; MX2; FANCA; EIF4G3; ISG20; MAPK1; TRIM68; XAF1; IRF2; TRIM5; OAS1; TUBB4A; IFIT3; SP100; HLA-C; OAS2; RIGI; ISG15; IFI6; GBP3; OAS3; STAT2; GBP4; IFNAR1; IFITM1; CIITA; TRIM25; OASL; EIF2AK2; DUS2; EIF4E3; FAAP24; NUP62; NUP37; TRIM10; IFI35; MAPK3; FANCL; IFITM3; HERC5 |
| R-HSA-500792 | GPCR ligand binding | 198 | 4.10E-01 | 1.52 | 1.69E-03 | 3.16E-02 | ACKR3; P2RY6; GIPR; CCR2; CX3CR1; CNR1; CXCR2; GPR132; PROK2; CCR9; FPR2; F2R; GNRH1; LPAR5; GNG2; GNB1; ADGRE2; FZD4; CXCL16; FFAR2; C3AR1; ADGRE3; FPR1; NMUR1; PTCH1; CCR5; CXCR1; P2RY14; OPRL1; UTS2; RAMP1; CNR2; ECE1; GNGT2; CMKLR1; CXCL1; CXCL2; CCL5; GNG7; EDN3; PNOC |
| R-HSA-168898 | Toll-like Receptor Cascades | 156 | 4.20E-01 | 1.51 | 2.52E-03 | 4.08E-02 | LGMN; USP18; RIPK3; PLCG2; MEF2C; TLR10; MAPK1; RPS6KA1; TLR6; MAP2K4; IRAK2; DNM2; NFKB2; MEF2A; MAPK9; IKBIP; MAPK14; TLR1; TLR4; MYD88; AGER; TRAF3; NLRX1; TLR8; TANK; N4BP1; UNC93B1; BTK; NKIRAS2; BIRC2; MAPK3; ATF1; TBK1; IRF7; TRAF2; IKBKG; VRK3; FBXW11; TP53; RPS6KA3; IKBKB; DUSP4 |
| R-HSA-109582 | Hemostasis | 472 | 3.81E-01 | 1.51 | 2.02E-05 | 6.69E-04 | OLR1; FN1; DOCK4; ORAI2; PLCG2; ITGAV; CD84; ITGA1; ITGAX; SLC16A3; TOR4A; P2RX5; VAV3; SDC2; ACTN4; ATP2A3; P2RX1; TIMP1; AKT1; VPREB3; CD58; CD109; CEACAM3; SPN; PRKAR1A; F2R; SCCPDH; MAPK1; MANF; CABLES2; IRAG1; VAV2; IRF2; GNG2; KIF3B; GNB1; INPP5D; ITPR1; FAM3C; STXBP3; PHF21A; ATP2A1; PECAM1; THBS1; MFN1; MICAL1; CRK; TUBB4A; SELPLG; CDK2; PIK3R3; F11R; STX4; ZFPM1; VAV1; KIF2C; MAPK14; GTPBP2; NHLRC2; ITGA4; CYRIB; CTSW; ORM1; KIF11; ABHD6; PDE2A; DOCK5; SIRPA; VTI1B; CEACAM1; CD99L2; SELL; CSK; SOS1; CD9; TRPC3; GNGT2; TEX264; MAPK3; RACGAP1; SH2B1; KIF21B; TGFB1; CENPE; ITGAL; PPP2R5C; ABL1; DGKD; APBB1IP; RAC2; HGF; GNA12; PRKCB; FGR; KLC4; SRC; KIF23; GNG7; TP53; PRKCE; AAMP; LYN; QSOX1; PDPK1; PIK3R6; SYK; FLNA; HDAC1; WDR1; CABLES1; GATA2; CAP1; PIK3R5; FERMT3; LGALS3BP; KIF15; MAGED2; KIF18B; RAF1; DGKK; PPIL2; DOCK8; TLN1; PTPN11; DOK2; BSG; NFE2; KIF4A; ATP2B4; APLP2; CD244; PDE1B; SERPINE1; PAFAH2; WEE1; ITPR3; ORM2; CDK5; AKAP10; ARRB2; ARRB1; CAPZB; LAMP2 |
| R-HSA-388396 | GPCR downstream signalling | 355 | 3.86E-01 | 1.49 | 1.85E-04 | 4.90E-03 | ACKR3; FN1; P2RY6; GIPR; ABR; ADCY4; CCR2; CX3CR1; GNAL; CNR1; CXCR2; VAV3; GPR132; TRIO; PROK2; AKT1; CCR9; FPR2; PRKAR1A; F2R; PDE4A; GNRH1; MAPK1; RPS6KA1; RGS3; VAV2; LPAR5; GNG2; AHCYL1; GNB1; ITPR1; RGS9; GPR27; PREX1; PLCB3; PIK3R3; VAV1; PPP1CA; CXCL16; FFAR2; C3AR1; FPR1; RGS19; ADCY7; ABHD6; NMUR1; CCR5; PDE2A; CXCR1; P2RY14; OPRL1; UTS2; RAMP1; PPP3CA; CNR2; SOS1; CAMKK2; TRPC3; GNGT2; BTK; ARHGEF40; MAPK3; RGS13; ARHGEF12; GRK2; DGKD; CXCL1; CXCL2; GNA12; CCL5; PRKCB; GRK3; SRC; ARHGEF39; ARHGEF2; GNG7; CAMKK1; EDN3; PRKCE; RPS6KA3; PNOC; PDPK1; ARHGEF17; PIK3R6; RGS14; TIAM2; PCP2; SOS2 |
| R-HSA-166016 | Toll Like Receptor 4 (TLR4) Cascade | 137 | 4.19E-01 | 1.49 | 3.07E-03 | 4.72E-02 | USP18; RIPK3; PLCG2; MEF2C; MAPK1; RPS6KA1; TLR6; MAP2K4; IRAK2; DNM2; NFKB2; MEF2A; MAPK9; IKBIP; MAPK14; TLR1; TLR4; MYD88; AGER; TRAF3; NLRX1; TANK; N4BP1; BTK; NKIRAS2; BIRC2; MAPK3; ATF1; TBK1; IRF7; TRAF2; IKBKG; VRK3; FBXW11; TP53; RPS6KA3; IKBKB; DUSP4 |
| R-HSA-372790 | Signaling by GPCR | 394 | 3.77E-01 | 1.47 | 3.69E-04 | 9.37E-03 | ACKR3; FN1; P2RY6; GIPR; ABR; ADCY4; CCR2; CX3CR1; GNAL; CNR1; CXCR2; VAV3; GPR132; TRIO; PROK2; AKT1; CCR9; FPR2; PRKAR1A; F2R; PDE4A; GNRH1; MAPK1; RPS6KA1; RGS3; VAV2; LPAR5; GNG2; AHCYL1; GNB1; ITPR1; RGS9; GPR27; PREX1; PLCB3; ADGRE2; FZD4; PIK3R3; VAV1; PPP1CA; CXCL16; FFAR2; C3AR1; ADGRE3; FPR1; RGS19; ADCY7; ABHD6; NMUR1; PTCH1; CCR5; PDE2A; CXCR1; P2RY14; OPRL1; UTS2; RAMP1; PPP3CA; CNR2; SOS1; ECE1; CAMKK2; TRPC3; GNGT2; BTK; ARHGEF40; MAPK3; RGS13; ARHGEF12; GRK2; CMKLR1; DGKD; CXCL1; CXCL2; GNA12; CCL5; PRKCB; GRK3; SRC; ARHGEF39; ARHGEF2; GNG7; CAMKK1; EDN3; PRKCE; RPS6KA3; PNOC; PDPK1; ARHGEF17; PIK3R6; RGS14; TIAM2; C5AR2; PCP2; SOS2 |
| R-HSA-446203 | Asparagine N-linked glycosylation | 274 | 3.74E-01 | 1.41 | 2.72E-03 | 4.36E-02 | GOLGA2; ST8SIA4; SLC35C1; SEC24D; MAN1A1; PREB; ST6GALNAC2; TGFA; MIA2; STX5; ST8SIA1; MOGS; COPB2; B4GALT5; RAB1B; LMAN2; TUBB4A; ST3GAL4; ST3GAL6; SEL1L; PPP6R1; ACTR1A; SEC23IP; LMAN2L; UGGT1; SYVN1; TMED9; TRAPPC1; TBC1D20; MVD; SPTA1; SPTBN4; COL7A1; NAPB; RNF5; SEC16A; CSNK1D; ST3GAL2; TRAPPC2B; GLB1; MLEC; ACTR10; ST8SIA6; B4GALT2; BET1L; ALG14; ALG5; DYNC1H1; DPAGT1; KDELR2; CTSC; ST6GAL1; SEC24A; TRAPPC3; COPA; ARFGAP1; DYNC1LI2; EDEM2; ARFGAP3; MGAT3; ARF4; SPTBN5; FUCA1; ST6GALNAC6; MAGT1; LMAN1; CAPZB; ST6GALNAC4; TRAPPC6B; GMPPB; SLC35A1; STT3A; SLC17A5; CAPZA1; DCTN3; NPL; TMED10; TMEM115; GFPT2; MAN2A2; DCTN2; COPG1; CHST10; NSF; CMAS; MGAT1 |
| R-HSA-9679506 | SARS-CoV Infections | 367 | -2.99E-01 | -1.34 | 1.00E-03 | 2.12E-02 | SAP18; DDX20; SNRPB; RPS29; RPS17; UBA52; ATP1A4; YWHAG; HNRNPA1; RPS2; RIPK2; FAU; ATP1A1; RPS19; RPS27; RPS3A; RPS8; RPS25; IL17RC; RPS18; RPS20; RPS15A; ISCU; HSPG2; MAN2A1; RPS23; G3BP2; RPS21; GEMIN4; RPS3; RPS6; UBE2N; RPS12; FNTA; CHD3; RPS15; CRBN; RPS28; PTGES3; CHMP7; KDM1A; ST3GAL3; HSP90AA1; RPS27A; NPM1; SNRPE; RPSA; RPS5; RCAN3; RPS4X; RPS16; RPS9; HSP90AB1; EEF1A1; MTA3; PALS1; CRB3; RPS13; RPS14; IMPDH2 |
| R-HSA-9694516 | SARS-CoV-2 Infection | 260 | -3.25E-01 | -1.40 | 2.19E-03 | 3.90E-02 | NUP93; SNRPD2; NDC1; G3BP1; YWHAE; SNRPG; MASP2; RPS7; RPS24; TOMM70; ATG14; RPS11; NUP54; DDX20; SNRPB; RPS29; RPS17; UBA52; YWHAG; RPS2; RIPK2; FAU; RPS19; RPS27; RPS3A; RPS8; RPS25; IL17RC; RPS18; RPS20; RPS15A; ISCU; HSPG2; MAN2A1; RPS23; G3BP2; RPS21; GEMIN4; RPS3; RPS6; UBE2N; RPS12; RPS15; RPS28; CHMP7; ST3GAL3; HSP90AA1; RPS27A; SNRPE; RPSA; RPS5; RPS4X; RPS16; RPS9; HSP90AB1; PALS1; CRB3; RPS13; RPS14 |
| R-HSA-1428517 | Aerobic respiration and respiratory electron transport | 206 | -3.67E-01 | -1.55 | 4.48E-04 | 1.11E-02 | COX20; NDUFA7; TRAP1; COX6A1; COX7B; IDH3A; ATP5ME; ACAT1; ARMC8; GPT; ATP5PB; TACO1; UQCRC1; NDUFV2; SDHD; NDUFA2; ATP5MG; COX4I1; NDUFAF4; ATP5F1C; NUBPL; ATP5MC2; NDUFB5; SURF1; COX7A2L; UBA52; UQCRB; UQCRFS1; ECSIT; LRPPRC; ATP5MC3; SLC25A27; DLST; ISCU; NDUFA4; SUCLG2; NDUFS5; CYCS; ATP5F1A; RPS27A; MDH1; LYRM4; COX7C; PDK1; FXN; LDHB |
| R-HSA-163200 | Respiratory electron transport, ATP synthesis by chemiosmotic coupling, and heat production by uncoupling proteins. | 133 | -4.00E-01 | -1.61 | 5.12E-04 | 1.22E-02 | NDUFAB1; NDUFB4; CYC1; COX5B; COX6C; UQCRH; COX20; NDUFA7; TRAP1; COX6A1; COX7B; ATP5ME; ATP5PB; TACO1; UQCRC1; NDUFV2; SDHD; NDUFA2; ATP5MG; COX4I1; NDUFAF4; ATP5F1C; NUBPL; ATP5MC2; NDUFB5; SURF1; COX7A2L; UQCRB; UQCRFS1; ECSIT; LRPPRC; ATP5MC3; SLC25A27; NDUFA4; NDUFS5; CYCS; ATP5F1A; MDH1; COX7C |
| R-HSA-5610785 | GLI3 is processed to GLI3R by the proteasome | 57 | -4.73E-01 | -1.63 | 2.80E-03 | 4.40E-02 | GLI3; PSMA6; PSMB1; PSMA7; PSMD6; PSMA1; PSMA3; PSMD7; UBA52; PSMB4; BTRC; CUL1; PSMB5; RPS27A; SKP1 |
| R-HSA-195253 | Degradation of beta-catenin by the destruction complex | 81 | -4.37E-01 | -1.64 | 2.48E-03 | 4.07E-02 | PSMA6; CTNNB1; PSMB1; PSMA7; PSMD6; PSMA1; PSMA3; PSMD7; PPP2CA; UBA52; PSMB4; BTRC; CUL1; PSMB5; RPS27A; TCF7; TLE2; LEF1; SKP1 |
| R-HSA-72203 | Processing of Capped Intron-Containing Pre-mRNA | 280 | -3.85E-01 | -1.67 | 3.08E-06 | 1.12E-04 | THOC3; U2AF1; ; CWC22; CDC40; NUP88; HNRNPH2; C9orf78; MFAP1; PNN; SEH1L; SRSF5; POLR2K; PRPF40A; HNRNPA3; PRPF18; NUP85; SYMPK; PCF11; NXF1; DDX42; RBM25; SNRPF; IK; DDX5; CWC25; PRP4K; ACIN1; SRRM2; SMU1; SF3B6; PPIL4; NCBP1; STEEP1; CHTOP; POLR2L; THOC6; LUC7L3; THOC1; SNRPC; CDC5L; CWF19L2; PUF60; LSM7; DHX8; SRSF7; TRA2B; LSM8; NUP188; DNAJC8; SNRPD1; RBM17; SNRNP27; FUS; HNRNPD; LSM4; SF3A1; SNRPB2; POLR2D; RBM7; NUP50; HTATSF1; RANBP2; WBP11; GCFC2; PRPF8; SDE2; NUP160; DHX15; DDX39B; SF3A2; CRNKL1; YJU2; HNRNPH1; SNW1; PPIE; DDX39A; HNRNPK; PPIG; RAE1; SNRNP25; NKAP; EIF4E; NUP93; TCERG1; PDCD7; SNRPD2; NDC1; SRSF10; CWC15; U2SURP; SNRPG; HNRNPU; BCAS2; DHX35; WTAP; PCBP2; RBM39; SRSF11; NUP54; SNRNP70; SAP18; SYF2; PRPF3; SNRPB; CLP1; WBP4; SNRPA1; SRSF3; HNRNPA1; SNU13; CSTF2T; CWC27; WDR33; POLR2C; SRRM1; DHX9; PRPF19; SRSF6; PQBP1; SRSF8; NXT1; SF3B1; RBMX; LSM5; CPSF6; WDR70; SF1; SNRPE; HNRNPM; HSPA8 |
| R-HSA-3371556 | Cellular response to heat stress | 82 | -4.44E-01 | -1.67 | 2.33E-03 | 4.04E-02 | NUP50; RANBP2; HSPH1; NUP160; CAMK2D; DNAJB6; HSPA9; HSPA1A; BAG5; RAE1; NUP93; NDC1; YWHAE; DNAJC2; SIRT1; NUP54; ATM; ST13; PTGES3; HSP90AA1; HSP90AB1; EEF1A1; BAG3; HSPA8; HSPA1B; DNAJB1 |
| R-HSA-9705683 | SARS-CoV-2-host interactions | 180 | -4.17E-01 | -1.72 | 1.06E-05 | 3.59E-04 | NUP93; SNRPD2; NDC1; G3BP1; YWHAE; SNRPG; MASP2; RPS7; RPS24; TOMM70; ATG14; RPS11; NUP54; DDX20; SNRPB; RPS29; RPS17; UBA52; YWHAG; RPS2; RIPK2; FAU; RPS19; RPS27; RPS3A; RPS8; RPS25; IL17RC; RPS18; RPS20; RPS15A; RPS23; G3BP2; RPS21; GEMIN4; RPS3; RPS6; UBE2N; RPS12; RPS15; RPS28; HSP90AA1; RPS27A; SNRPE; RPSA; RPS5; RPS4X; RPS16; RPS9; HSP90AB1; PALS1; CRB3; RPS13; RPS14 |
| R-HSA-3371497 | HSP90 chaperone cycle for steroid hormone receptors (SHR) in the presence of ligand | 45 | -5.25E-01 | -1.72 | 2.78E-03 | 4.40E-02 | TUBB1; PTGES3; HSP90AA1; HSP90AB1; HSPA8; DNAJA4; HSPA1B; DNAJB1; NR3C2 |
| R-HSA-9675108 | Nervous system development | 470 | -3.84E-01 | -1.74 | 1.84E-10 | 7.88E-09 | PSMA1; EPHB6; PSMA3; ROBO3; DNM3; RPS11; NEO1; PSMD7; KALRN; ABLIM3; PRKCQ; ABL2; PLXNB3; RPL35; ABLIM2; SIAH1; RPS29; LAMB1; RPS17; UBA52; RPL22L1; PSMB4; RPLP1; RPL37; TUBB1; EPHA2; EPHB3; CDC42; RPS2; FAU; RPS19; RPS27; NRCAM; RPL11; RPS3A; RPS8; SRGAP3; EPHA1; RPS25; RPS18; RPL39; RPS20; DLG1; RPL27A; RPS15A; RPL6; SMARCA4; RPS23; PLCG1; ROBO1; RPL12; RPS21; PSMB5; RPL9; ITGB3; RPS3; RPS6; RPS12; SEMA5A; RPS15; RPS28; RPLP2; RPL14; DAB1; HSP90AA1; RPL27; COL5A2; RPL15; RPL24; RPS27A; RPL21; RPL18A; RPL29; RPL23; RPL35A; RPL10; RPSA; RPS5; RPL7A; RPLP0; RPL18; RPL13; RPL8; RPL13A; RPL32; RPL31; RPL22; RPS4X; RPS16; RPL19; RPL38; RPS9; HSP90AB1; PFN2; RPL7; RPL34; PRKCA; RGMB; TIAM1; RPL10A; HSPA8; RPL5; RPS13; RPL4; RPL3; RPS14; TRPC1; CACNA1I |
| R-HSA-72172 | mRNA Splicing | 213 | -4.13E-01 | -1.76 | 3.29E-06 | 1.17E-04 | U2AF1; ; CWC22; CDC40; HNRNPH2; C9orf78; MFAP1; PNN; SRSF5; POLR2K; PRPF40A; HNRNPA3; PRPF18; DDX42; RBM25; SNRPF; IK; DDX5; CWC25; PRP4K; ACIN1; SRRM2; SMU1; SF3B6; PPIL4; NCBP1; STEEP1; POLR2L; LUC7L3; SNRPC; CDC5L; CWF19L2; PUF60; LSM7; DHX8; SRSF7; TRA2B; LSM8; DNAJC8; SNRPD1; RBM17; SNRNP27; FUS; HNRNPD; LSM4; SF3A1; SNRPB2; POLR2D; RBM7; HTATSF1; WBP11; GCFC2; PRPF8; SDE2; DHX15; DDX39B; SF3A2; CRNKL1; YJU2; HNRNPH1; SNW1; PPIE; HNRNPK; PPIG; SNRNP25; NKAP; TCERG1; PDCD7; SNRPD2; SRSF10; CWC15; U2SURP; SNRPG; HNRNPU; BCAS2; DHX35; PCBP2; RBM39; SRSF11; SNRNP70; SAP18; SYF2; PRPF3; SNRPB; WBP4; SNRPA1; SRSF3; HNRNPA1; SNU13; CWC27; POLR2C; SRRM1; DHX9; PRPF19; SRSF6; PQBP1; SRSF8; SF3B1; RBMX; LSM5; WDR70; SF1; SNRPE; HNRNPM; HSPA8 |
| R-HSA-9678108 | SARS-CoV-1 Infection | 135 | -4.39E-01 | -1.77 | 3.90E-05 | 1.18E-03 | RPS29; RPS17; UBA52; YWHAG; HNRNPA1; RPS2; FAU; RPS19; RPS27; RPS3A; RPS8; RPS25; RPS18; RPS20; RPS15A; RPS23; RPS21; RPS3; RPS6; RPS12; RPS15; RPS28; CHMP7; ST3GAL3; RPS27A; NPM1; RPSA; RPS5; RCAN3; RPS4X; RPS16; RPS9; EEF1A1; PALS1; RPS13; RPS14 |
| R-HSA-72163 | mRNA Splicing - Major Pathway | 205 | -4.20E-01 | -1.78 | 3.08E-06 | 1.12E-04 | U2AF1; ; CWC22; CDC40; HNRNPH2; C9orf78; MFAP1; PNN; SRSF5; POLR2K; PRPF40A; HNRNPA3; PRPF18; DDX42; RBM25; SNRPF; IK; DDX5; CWC25; PRP4K; ACIN1; SRRM2; SMU1; SF3B6; PPIL4; NCBP1; STEEP1; POLR2L; LUC7L3; SNRPC; CDC5L; CWF19L2; PUF60; LSM7; DHX8; SRSF7; TRA2B; LSM8; DNAJC8; SNRPD1; RBM17; SNRNP27; FUS; HNRNPD; LSM4; SF3A1; SNRPB2; POLR2D; RBM7; HTATSF1; WBP11; GCFC2; PRPF8; SDE2; DHX15; DDX39B; SF3A2; CRNKL1; YJU2; HNRNPH1; SNW1; PPIE; HNRNPK; PPIG; NKAP; TCERG1; SNRPD2; SRSF10; CWC15; U2SURP; SNRPG; HNRNPU; BCAS2; DHX35; PCBP2; RBM39; SRSF11; SNRNP70; SAP18; SYF2; PRPF3; SNRPB; WBP4; SNRPA1; SRSF3; HNRNPA1; SNU13; CWC27; POLR2C; SRRM1; DHX9; PRPF19; SRSF6; PQBP1; SRSF8; SF3B1; RBMX; LSM5; WDR70; SF1; SNRPE; HNRNPM; HSPA8 |
| R-HSA-422475 | Axon guidance | 452 | -3.94E-01 | -1.79 | 1.00E-10 | 4.42E-09 | PSMA1; EPHB6; PSMA3; ROBO3; DNM3; RPS11; NEO1; PSMD7; KALRN; ABLIM3; PRKCQ; ABL2; PLXNB3; RPL35; ABLIM2; SIAH1; RPS29; LAMB1; RPS17; UBA52; RPL22L1; PSMB4; RPLP1; RPL37; TUBB1; EPHA2; EPHB3; CDC42; RPS2; FAU; RPS19; RPS27; NRCAM; RPL11; RPS3A; RPS8; SRGAP3; EPHA1; RPS25; RPS18; RPL39; RPS20; DLG1; RPL27A; RPS15A; RPL6; RPS23; PLCG1; ROBO1; RPL12; RPS21; PSMB5; RPL9; ITGB3; RPS3; RPS6; RPS12; SEMA5A; RPS15; RPS28; RPLP2; RPL14; DAB1; HSP90AA1; RPL27; COL5A2; RPL15; RPL24; RPS27A; RPL21; RPL18A; RPL29; RPL23; RPL35A; RPL10; RPSA; RPS5; RPL7A; RPLP0; RPL18; RPL13; RPL8; RPL13A; RPL32; RPL31; RPL22; RPS4X; RPS16; RPL19; RPL38; RPS9; HSP90AB1; PFN2; RPL7; RPL34; PRKCA; RGMB; TIAM1; RPL10A; HSPA8; RPL5; RPS13; RPL4; RPL3; RPS14; TRPC1; CACNA1I |
| R-HSA-1268020 | Mitochondrial protein import | 61 | -5.22E-01 | -1.83 | 5.27E-04 | 1.24E-02 | CHCHD7; CYC1; TIMM17A; HSPA9; GRPEL1; SAMM50; TOMM22; PMPCB; PMPCA; TIMM13; TIMM9; HSPD1; TOMM70; TIMM23; CHCHD2; TOMM7; ATP5F1A; GRPEL2; TOMM40; TOMM20; FXN |
| R-HSA-8949613 | Cristae formation | 31 | -6.29E-01 | -1.92 | 5.07E-04 | 1.22E-02 | ATP5PO; APOO; HSPA9; SAMM50; ATP5ME; ATP5PB; ATP5MG; TMEM11; ATP5F1C; ATP5MC2; ATP5MC3; ATP5F1A; CHCHD6; MICOS10 |
| R-HSA-9613829 | Chaperone Mediated Autophagy | 18 | -7.17E-01 | -1.94 | 8.95E-04 | 1.98E-02 | UBA52; PLIN2; HSP90AA1; RPS27A; HSP90AB1; EEF1A1; HSPA8 |
| R-HSA-9629569 | Protein hydroxylation | 19 | -7.30E-01 | -1.99 | 6.17E-04 | 1.43E-02 | ETF1; ASPH; DRG1; KDM8; ZC3H15; OGFOD1; RIOX1; RWDD1; RPL27A; RPS23; RPS6; RIOX2; RPL8 |
| R-HSA-6790901 | rRNA modification in the nucleus and cytosol | 60 | -5.82E-01 | -2.06 | 2.16E-05 | 7.01E-04 | PNO1; IMP3; MPHOSPH10; BUD23; RPS7; UTP14A; WDR36; UTP15; KRR1; NOP58; NAT10; NOP56; SNU13; RPS2; UTP20; NOL11; DKC1; RCL1; UTP4; RPS6; WDR43; FBL; RPS9; UTP18; RPS14 |
| R-HSA-9692914 | SARS-CoV-1-host interactions | 93 | -5.52E-01 | -2.11 | 1.28E-06 | 5.03E-05 | YWHAE; RPS7; RPS24; PCBP2; TOMM70; RPS11; RPS29; RPS17; UBA52; YWHAG; HNRNPA1; RPS2; FAU; RPS19; RPS27; RPS3A; RPS8; RPS25; RPS18; RPS20; RPS15A; RPS23; RPS21; RPS3; RPS6; RPS12; RPS15; RPS28; RPS27A; NPM1; RPSA; RPS5; RCAN3; RPS4X; RPS16; RPS9; EEF1A1; PALS1; RPS13; RPS14 |
| R-HSA-3371568 | Attenuation phase | 13 | -8.87E-01 | -2.18 | 2.43E-06 | 9.34E-05 | PTGES3; HSP90AA1; HSP90AB1; HSPA8; HSPA1B; DNAJB1 |
| R-HSA-3371571 | HSF1-dependent transactivation | 19 | -8.18E-01 | -2.23 | 5.83E-06 | 2.02E-04 | PTGES3; HSP90AA1; HSP90AB1; HSPA8; HSPA1B; DNAJB1 |
| R-HSA-71291 | Metabolism of amino acids and derivatives | 301 | -5.64E-01 | -2.47 | 1.00E-10 | 4.42E-09 | PSMD6; PSMA1; PSMA3; RPS11; PSMD7; SERINC1; RPL35; SECISBP2; RPS29; RPS17; UBA52; AHCY; RPL22L1; PSMB4; RPLP1; MCCC1; GLUD1; RPL37; RPS2; FAU; RPS19; NAALAD2; RPS27; RPL11; MTAP; RPS3A; RPS8; RPS25; RPS18; DLST; RPL39; RPS20; RPL27A; RPS15A; RPL6; AMD1; RPS23; RPL12; RPS21; PSMB5; RPL9; RPS3; RPS6; RPS12; RPS15; RPS28; DBH; RPLP2; RPL14; RPL27; RPL15; RPL24; RPS27A; RPL21; SARDH; RPL18A; RPL29; RPL23; RPL35A; RPL10; RPSA; RPS5; RPL7A; RPLP0; RPL18; ALDH7A1; RPL13; RPL8; RPL13A; RPL32; RPL31; RPL22; RPS4X; RPS16; RPL19; RPL38; GAMT; RPS9; RPL7; RPL34; SERINC5; AIMP1; RPL10A; RPL5; RPS13; RPL4; RPL3; KARS1; RPS14; AASS |
| R-HSA-9754678 | SARS-CoV-2 modulates host translation machinery | 50 | -7.73E-01 | -2.62 | 1.00E-10 | 4.42E-09 | SNRPD2; SNRPG; RPS7; RPS24; RPS11; DDX20; SNRPB; RPS29; RPS17; RPS2; FAU; RPS19; RPS27; RPS3A; RPS8; RPS25; RPS18; RPS20; RPS15A; RPS23; RPS21; GEMIN4; RPS3; RPS6; RPS12; RPS15; RPS28; RPS27A; SNRPE; RPSA; RPS5; RPS4X; RPS16; RPS9; RPS13; RPS14 |
| R-HSA-376176 | Signaling by ROBO receptors | 199 | -6.44E-01 | -2.69 | 1.00E-10 | 4.42E-09 | PSMD6; PSMA1; PSMA3; ROBO3; RPS11; PSMD7; ABL2; RPL35; RPS29; RPS17; UBA52; RPL22L1; PSMB4; RPLP1; RPL37; CDC42; RPS2; FAU; RPS19; RPS27; RPL11; RPS3A; RPS8; SRGAP3; RPS25; RPS18; RPL39; RPS20; RPL27A; RPS15A; RPL6; RPS23; ROBO1; RPL12; RPS21; PSMB5; RPL9; RPS3; RPS6; RPS12; RPS15; RPS28; RPLP2; RPL14; RPL27; RPL15; RPL24; RPS27A; RPL21; RPL18A; RPL29; RPL23; RPL35A; RPL10; RPSA; RPS5; RPL7A; RPLP0; RPL18; RPL13; RPL8; RPL13A; RPL32; RPL31; RPL22; RPS4X; RPS16; RPL19; RPL38; RPS9; PFN2; RPL7; RPL34; PRKCA; RPL10A; RPL5; RPS13; RPL4; RPL3; RPS14 |
| R-HSA-9735869 | SARS-CoV-1 modulates host translation machinery | 36 | -8.57E-01 | -2.71 | 1.00E-10 | 4.42E-09 | RPS7; RPS24; RPS11; RPS29; RPS17; HNRNPA1; RPS2; FAU; RPS19; RPS27; RPS3A; RPS8; RPS25; RPS18; RPS20; RPS15A; RPS23; RPS21; RPS3; RPS6; RPS12; RPS15; RPS28; RPS27A; RPSA; RPS5; RPS4X; RPS16; RPS9; EEF1A1; RPS13; RPS14 |
| R-HSA-72766 | Translation | 287 | -6.27E-01 | -2.73 | 1.00E-10 | 4.42E-09 | RPL37A; RPL41; SRP19; SRP68; SRPRB; MRPS28; MRPS27; PTCD3; NARS2; EIF4E; EIF2S3; SSR4; MRPL49; EIF1AX; RPS7; RPS24; RPL36; EIF3K; AARS2; MRPL11; MRPL41; EIF4H; MRPL55; RPS11; MRPS9; MRPL42; EIF5; TSFM; RPL35; MRPS7; RPS29; RPS17; UBA52; RPL22L1; RPLP1; RPL37; MRPL9; MRPL32; RPS2; SRP72; EIF3I; FAU; EIF3D; RPS19; RPS27; RPL11; EIF3M; RPS3A; RPS8; RPS25; RPS18; EIF2B3; RPL39; RPS20; MRPL48; RPL27A; RPS15A; RPL6; FARSB; RPS23; RPL12; RPS21; MRPS25; RPL9; RPS3; RPS6; RPS12; MRPL38; RPS15; RPS28; RPLP2; RPL14; EEF1D; EEF2; RPL27; RPL15; RPL24; RPS27A; RPL21; EIF4B; RPL18A; EEF1B2; RPL29; RPL23; RPL35A; RPL10; RPSA; RPS5; RPL7A; RPLP0; DAP3; RPL18; RPL13; RPL8; RPL13A; RPL32; RPL31; RPL22; RPS4X; EIF3E; PPA1; RPS16; MRPS33; RPL19; RPL38; RPS9; EEF1A1; RPL7; RPL34; EIF3F; AIMP1; EIF3H; RPL10A; RPL5; EIF3L; RPS13; RPL4; RPL3; KARS1; RPS14 |
| R-HSA-72649 | Translation initiation complex formation | 58 | -7.99E-01 | -2.79 | 1.00E-10 | 4.42E-09 | EIF4E; EIF2S3; EIF1AX; RPS7; RPS24; EIF3K; EIF4H; RPS11; RPS29; RPS17; RPS2; EIF3I; FAU; EIF3D; RPS19; RPS27; EIF3M; RPS3A; RPS8; RPS25; RPS18; RPS20; RPS15A; RPS23; RPS21; RPS3; RPS6; RPS12; RPS15; RPS28; RPS27A; EIF4B; RPSA; RPS5; RPS4X; EIF3E; RPS16; RPS9; EIF3F; EIF3H; EIF3L; RPS13; RPS14 |
| R-HSA-72702 | Ribosomal scanning and start codon recognition | 58 | -8.02E-01 | -2.80 | 1.00E-10 | 4.42E-09 | EIF4E; EIF2S3; EIF1AX; RPS7; RPS24; EIF3K; EIF4H; RPS11; EIF5; RPS29; RPS17; RPS2; EIF3I; FAU; EIF3D; RPS19; RPS27; EIF3M; RPS3A; RPS8; RPS25; RPS18; RPS20; RPS15A; RPS23; RPS21; RPS3; RPS6; RPS12; RPS15; RPS28; RPS27A; EIF4B; RPSA; RPS5; RPS4X; EIF3E; RPS16; RPS9; EIF3F; EIF3H; EIF3L; RPS13; RPS14 |
| R-HSA-72662 | Activation of the mRNA upon binding of the cap-binding complex and eIFs, and subsequent binding to 43S | 59 | -7.99E-01 | -2.80 | 1.00E-10 | 4.42E-09 | EIF4E; EIF2S3; EIF1AX; RPS7; RPS24; EIF3K; EIF4H; RPS11; RPS29; RPS17; RPS2; EIF3I; FAU; EIF3D; RPS19; RPS27; EIF3M; RPS3A; RPS8; RPS25; RPS18; RPS20; RPS15A; RPS23; RPS21; RPS3; RPS6; RPS12; RPS15; RPS28; RPS27A; EIF4B; RPSA; RPS5; RPS4X; EIF3E; RPS16; RPS9; EIF3F; EIF3H; EIF3L; RPS13; RPS14 |
| R-HSA-72695 | Formation of the ternary complex, and subsequently, the 43S complex | 51 | -8.21E-01 | -2.80 | 1.00E-10 | 4.42E-09 | EIF2S3; EIF1AX; RPS7; RPS24; EIF3K; RPS11; RPS29; RPS17; RPS2; EIF3I; FAU; EIF3D; RPS19; RPS27; EIF3M; RPS3A; RPS8; RPS25; RPS18; RPS20; RPS15A; RPS23; RPS21; RPS3; RPS6; RPS12; RPS15; RPS28; RPS27A; RPSA; RPS5; RPS4X; EIF3E; RPS16; RPS9; EIF3F; EIF3H; EIF3L; RPS13; RPS14 |
| R-HSA-9010553 | Regulation of expression of SLITs and ROBOs | 159 | -7.21E-01 | -2.91 | 1.00E-10 | 4.42E-09 | PSMD6; PSMA1; PSMA3; ROBO3; RPS11; PSMD7; RPL35; RPS29; RPS17; UBA52; RPL22L1; PSMB4; RPLP1; RPL37; RPS2; FAU; RPS19; RPS27; RPL11; RPS3A; RPS8; RPS25; RPS18; RPL39; RPS20; RPL27A; RPS15A; RPL6; RPS23; ROBO1; RPL12; RPS21; PSMB5; RPL9; RPS3; RPS6; RPS12; RPS15; RPS28; RPLP2; RPL14; RPL27; RPL15; RPL24; RPS27A; RPL21; RPL18A; RPL29; RPL23; RPL35A; RPL10; RPSA; RPS5; RPL7A; RPLP0; RPL18; RPL13; RPL8; RPL13A; RPL32; RPL31; RPL22; RPS4X; RPS16; RPL19; RPL38; RPS9; RPL7; RPL34; RPL10A; RPL5; RPS13; RPL4; RPL3; RPS14 |
| R-HSA-72312 | rRNA processing | 200 | -7.01E-01 | -2.93 | 1.00E-10 | 4.42E-09 | LTV1; EXOSC9; RPS7; RPS24; RPL36; UTP14A; WDR36; UTP15; NOB1; NOL9; KRR1; NOP58; NAT10; RPS11; XRN2; NCL; NOP56; DDX21; WDR12; RPL35; RPS29; RPS17; UBA52; RPL22L1; RPLP1; SNU13; RPL37; RPS2; FAU; RPS19; RPS27; RPL11; UTP20; RPS3A; RPS8; NOL11; RPS25; RPS18; RPL39; RPS20; DKC1; RCL1; RPL27A; RPS15A; RPL6; RPS23; EXOSC7; RPL12; RPS21; UTP4; RPL9; CSNK1E; RPS3; TSR1; RPS6; RPS12; TRMT10C; RPS15; RPS28; RPLP2; RPL14; RPL27; RPL15; RPL24; RPS27A; RPL21; RPL18A; WDR43; RPL29; RPL23; RPL35A; RPL10; RPSA; RPS5; RPL7A; RPLP0; RPL18; RPL13; RPL8; RPL13A; RPL32; RPL31; RPL22; LAS1L; RPS4X; RPS16; FBL; RPL19; RPL38; RPS9; RPL7; RPL34; RPL10A; UTP18; RPL5; RPS13; RPL4; RPL3; RPS14 |
| R-HSA-8868773 | rRNA processing in the nucleus and cytosol | 190 | -7.11E-01 | -2.94 | 1.00E-10 | 4.42E-09 | LTV1; EXOSC9; RPS7; RPS24; RPL36; UTP14A; WDR36; UTP15; NOB1; NOL9; KRR1; NOP58; NAT10; RPS11; XRN2; NCL; NOP56; DDX21; WDR12; RPL35; RPS29; RPS17; UBA52; RPL22L1; RPLP1; SNU13; RPL37; RPS2; FAU; RPS19; RPS27; RPL11; UTP20; RPS3A; RPS8; NOL11; RPS25; RPS18; RPL39; RPS20; DKC1; RCL1; RPL27A; RPS15A; RPL6; RPS23; EXOSC7; RPL12; RPS21; UTP4; RPL9; CSNK1E; RPS3; TSR1; RPS6; RPS12; RPS15; RPS28; RPLP2; RPL14; RPL27; RPL15; RPL24; RPS27A; RPL21; RPL18A; WDR43; RPL29; RPL23; RPL35A; RPL10; RPSA; RPS5; RPL7A; RPLP0; RPL18; RPL13; RPL8; RPL13A; RPL32; RPL31; RPL22; LAS1L; RPS4X; RPS16; FBL; RPL19; RPL38; RPS9; RPL7; RPL34; RPL10A; UTP18; RPL5; RPS13; RPL4; RPL3; RPS14 |
| R-HSA-168255 | Influenza Infection | 152 | -7.30E-01 | -2.95 | 1.00E-10 | 4.42E-09 | RPL35; RPS29; RPS17; UBA52; RPL22L1; RPLP1; RPL37; RPS2; FAU; POLR2C; RPS19; RPS27; RPL11; RPS3A; RPS8; RPS25; RPS18; RPL39; RPS20; RPL27A; RPS15A; RPL6; RPS23; RPL12; RPS21; RPL9; RPS3; RPS6; RPS12; RAN; RPS15; RPS28; RPLP2; RPL14; HSP90AA1; RPL27; RPL15; RPL24; RPS27A; RPL21; RPL18A; RPL29; RPL23; RPL35A; RPL10; RPSA; RPS5; RPL7A; RPLP0; RPL18; RPL13; RPL8; RPL13A; IPO5; RPL32; RPL31; RPL22; RPS4X; RPS16; RPL19; RPL38; RPS9; RPL7; RPL34; RPL10A; RPL5; RPS13; RPL4; RPL3; RPS14; HSPA1B |
| R-HSA-6791226 | Major pathway of rRNA processing in the nucleolus and cytosol | 180 | -7.19E-01 | -2.96 | 1.00E-10 | 4.42E-09 | LTV1; EXOSC9; RPS7; RPS24; RPL36; UTP14A; WDR36; UTP15; NOB1; NOL9; KRR1; NOP58; RPS11; XRN2; NCL; NOP56; DDX21; WDR12; RPL35; RPS29; RPS17; UBA52; RPL22L1; RPLP1; SNU13; RPL37; RPS2; FAU; RPS19; RPS27; RPL11; UTP20; RPS3A; RPS8; NOL11; RPS25; RPS18; RPL39; RPS20; RCL1; RPL27A; RPS15A; RPL6; RPS23; EXOSC7; RPL12; RPS21; UTP4; RPL9; CSNK1E; RPS3; TSR1; RPS6; RPS12; RPS15; RPS28; RPLP2; RPL14; RPL27; RPL15; RPL24; RPS27A; RPL21; RPL18A; WDR43; RPL29; RPL23; RPL35A; RPL10; RPSA; RPS5; RPL7A; RPLP0; RPL18; RPL13; RPL8; RPL13A; RPL32; RPL31; RPL22; LAS1L; RPS4X; RPS16; FBL; RPL19; RPL38; RPS9; RPL7; RPL34; RPL10A; UTP18; RPL5; RPS13; RPL4; RPL3; RPS14 |
| R-HSA-9711097 | Cellular response to starvation | 151 | -7.41E-01 | -2.99 | 1.00E-10 | 4.42E-09 | RPL35; RPS29; RPS17; UBA52; RPL22L1; RPLP1; RPL37; RPS2; FAU; RPS19; RPS27; RPL11; RPS3A; RPS8; RPS25; RPS18; RPL39; RPS20; RPL27A; RPS15A; RPL6; RPS23; RPL12; ATP6V1G1; RPS21; RPL9; RPS3; RPS6; RPS12; RPS15; DDIT3; RPS28; RPLP2; RPL14; ATF3; RPL27; RPL15; RPL24; RPS27A; RPL21; RPL18A; RPL29; RPL23; RPL35A; RPL10; RPSA; RPS5; RPL7A; RPLP0; RPL18; RPL13; RPL8; RPL13A; RPL32; RPL31; RPL22; RPS4X; RPS16; RPL19; RPL38; RPS9; RPL7; RPL34; RPL10A; RPL5; RPS13; RPL4; RPL3; RPS14; SESN1 |
| R-HSA-168273 | Influenza Viral RNA Transcription and Replication | 133 | -7.63E-01 | -3.06 | 1.00E-10 | 4.42E-09 | RPL35; RPS29; RPS17; UBA52; RPL22L1; RPLP1; RPL37; RPS2; FAU; POLR2C; RPS19; RPS27; RPL11; RPS3A; RPS8; RPS25; RPS18; RPL39; RPS20; RPL27A; RPS15A; RPL6; RPS23; RPL12; RPS21; RPL9; RPS3; RPS6; RPS12; RPS15; RPS28; RPLP2; RPL14; HSP90AA1; RPL27; RPL15; RPL24; RPS27A; RPL21; RPL18A; RPL29; RPL23; RPL35A; RPL10; RPSA; RPS5; RPL7A; RPLP0; RPL18; RPL13; RPL8; RPL13A; IPO5; RPL32; RPL31; RPL22; RPS4X; RPS16; RPL19; RPL38; RPS9; RPL7; RPL34; RPL10A; RPL5; RPS13; RPL4; RPL3; RPS14 |
| R-HSA-1799339 | SRP-dependent cotranslational protein targeting to membrane | 110 | -7.98E-01 | -3.12 | 1.00E-10 | 4.42E-09 | RPL35; RPS29; RPS17; UBA52; RPL22L1; RPLP1; RPL37; RPS2; SRP72; FAU; RPS19; RPS27; RPL11; RPS3A; RPS8; RPS25; RPS18; RPL39; RPS20; RPL27A; RPS15A; RPL6; RPS23; RPL12; RPS21; RPL9; RPS3; RPS6; RPS12; RPS15; RPS28; RPLP2; RPL14; RPL27; RPL15; RPL24; RPS27A; RPL21; RPL18A; RPL29; RPL23; RPL35A; RPL10; RPSA; RPS5; RPL7A; RPLP0; RPL18; RPL13; RPL8; RPL13A; RPL32; RPL31; RPL22; RPS4X; RPS16; RPL19; RPL38; RPS9; RPL7; RPL34; RPL10A; RPL5; RPS13; RPL4; RPL3; RPS14 |
| R-HSA-927802 | Nonsense-Mediated Decay (NMD) | 113 | -7.97E-01 | -3.13 | 1.00E-10 | 4.42E-09 | PPP2CA; RPL35; RPS29; RPS17; UBA52; RPL22L1; RPLP1; RPL37; RPS2; FAU; RPS19; RPS27; RPL11; RPS3A; RPS8; RPS25; RPS18; RPL39; RPS20; RPL27A; RPS15A; RPL6; RPS23; RPL12; RPS21; RPL9; SMG6; RPS3; RPS6; RPS12; RPS15; RPS28; RPLP2; RPL14; RPL27; RPL15; RPL24; RPS27A; RPL21; RPL18A; RPL29; RPL23; RPL35A; RPL10; RPSA; RPS5; RPL7A; RPLP0; RPL18; RPL13; RPL8; RPL13A; RPL32; RPL31; RPL22; RPS4X; RPS16; RPL19; RPL38; RPS9; RPL7; RPL34; RPL10A; RPL5; RPS13; RPL4; RPL3; RPS14 |
| R-HSA-975957 | Nonsense Mediated Decay (NMD) enhanced by the Exon Junction Complex (EJC) | 113 | -7.97E-01 | -3.13 | 1.00E-10 | 4.42E-09 | PPP2CA; RPL35; RPS29; RPS17; UBA52; RPL22L1; RPLP1; RPL37; RPS2; FAU; RPS19; RPS27; RPL11; RPS3A; RPS8; RPS25; RPS18; RPL39; RPS20; RPL27A; RPS15A; RPL6; RPS23; RPL12; RPS21; RPL9; SMG6; RPS3; RPS6; RPS12; RPS15; RPS28; RPLP2; RPL14; RPL27; RPL15; RPL24; RPS27A; RPL21; RPL18A; RPL29; RPL23; RPL35A; RPL10; RPSA; RPS5; RPL7A; RPLP0; RPL18; RPL13; RPL8; RPL13A; RPL32; RPL31; RPL22; RPS4X; RPS16; RPL19; RPL38; RPS9; RPL7; RPL34; RPL10A; RPL5; RPS13; RPL4; RPL3; RPS14 |
| R-HSA-72613 | Eukaryotic Translation Initiation | 117 | -8.08E-01 | -3.20 | 1.00E-10 | 4.42E-09 | EIF4H; RPS11; EIF5; RPL35; RPS29; RPS17; UBA52; RPL22L1; RPLP1; RPL37; RPS2; EIF3I; FAU; EIF3D; RPS19; RPS27; RPL11; EIF3M; RPS3A; RPS8; RPS25; RPS18; EIF2B3; RPL39; RPS20; RPL27A; RPS15A; RPL6; RPS23; RPL12; RPS21; RPL9; RPS3; RPS6; RPS12; RPS15; RPS28; RPLP2; RPL14; RPL27; RPL15; RPL24; RPS27A; RPL21; EIF4B; RPL18A; RPL29; RPL23; RPL35A; RPL10; RPSA; RPS5; RPL7A; RPLP0; RPL18; RPL13; RPL8; RPL13A; RPL32; RPL31; RPL22; RPS4X; EIF3E; RPS16; RPL19; RPL38; RPS9; RPL7; RPL34; EIF3F; EIF3H; RPL10A; RPL5; EIF3L; RPS13; RPL4; RPL3; RPS14 |
| R-HSA-72737 | Cap-dependent Translation Initiation | 117 | -8.08E-01 | -3.20 | 1.00E-10 | 4.42E-09 | EIF4H; RPS11; EIF5; RPL35; RPS29; RPS17; UBA52; RPL22L1; RPLP1; RPL37; RPS2; EIF3I; FAU; EIF3D; RPS19; RPS27; RPL11; EIF3M; RPS3A; RPS8; RPS25; RPS18; EIF2B3; RPL39; RPS20; RPL27A; RPS15A; RPL6; RPS23; RPL12; RPS21; RPL9; RPS3; RPS6; RPS12; RPS15; RPS28; RPLP2; RPL14; RPL27; RPL15; RPL24; RPS27A; RPL21; EIF4B; RPL18A; RPL29; RPL23; RPL35A; RPL10; RPSA; RPS5; RPL7A; RPLP0; RPL18; RPL13; RPL8; RPL13A; RPL32; RPL31; RPL22; RPS4X; EIF3E; RPS16; RPL19; RPL38; RPS9; RPL7; RPL34; EIF3F; EIF3H; RPL10A; RPL5; EIF3L; RPS13; RPL4; RPL3; RPS14 |
| R-HSA-72706 | GTP hydrolysis and joining of the 60S ribosomal subunit | 110 | -8.18E-01 | -3.20 | 1.00E-10 | 4.42E-09 | EIF4H; RPS11; EIF5; RPL35; RPS29; RPS17; UBA52; RPL22L1; RPLP1; RPL37; RPS2; EIF3I; FAU; EIF3D; RPS19; RPS27; RPL11; EIF3M; RPS3A; RPS8; RPS25; RPS18; RPL39; RPS20; RPL27A; RPS15A; RPL6; RPS23; RPL12; RPS21; RPL9; RPS3; RPS6; RPS12; RPS15; RPS28; RPLP2; RPL14; RPL27; RPL15; RPL24; RPS27A; RPL21; EIF4B; RPL18A; RPL29; RPL23; RPL35A; RPL10; RPSA; RPS5; RPL7A; RPLP0; RPL18; RPL13; RPL8; RPL13A; RPL32; RPL31; RPL22; RPS4X; EIF3E; RPS16; RPL19; RPL38; RPS9; RPL7; RPL34; EIF3F; EIF3H; RPL10A; RPL5; EIF3L; RPS13; RPL4; RPL3; RPS14 |
| R-HSA-2408522 | Selenoamino acid metabolism | 111 | -8.16E-01 | -3.20 | 1.00E-10 | 4.42E-09 | RPL35; SECISBP2; RPS29; RPS17; UBA52; AHCY; RPL22L1; RPLP1; RPL37; RPS2; FAU; RPS19; RPS27; RPL11; RPS3A; RPS8; RPS25; RPS18; RPL39; RPS20; RPL27A; RPS15A; RPL6; RPS23; RPL12; RPS21; RPL9; RPS3; RPS6; RPS12; RPS15; RPS28; RPLP2; RPL14; RPL27; RPL15; RPL24; RPS27A; RPL21; RPL18A; RPL29; RPL23; RPL35A; RPL10; RPSA; RPS5; RPL7A; RPLP0; RPL18; RPL13; RPL8; RPL13A; RPL32; RPL31; RPL22; RPS4X; RPS16; RPL19; RPL38; RPS9; RPL7; RPL34; AIMP1; RPL10A; RPL5; RPS13; RPL4; RPL3; KARS1; RPS14 |
| R-HSA-156827 | L13a-mediated translational silencing of Ceruloplasmin expression | 109 | -8.22E-01 | -3.20 | 1.00E-10 | 4.42E-09 | RPL35; RPS29; RPS17; UBA52; RPL22L1; RPLP1; RPL37; RPS2; EIF3I; FAU; EIF3D; RPS19; RPS27; RPL11; EIF3M; RPS3A; RPS8; RPS25; RPS18; RPL39; RPS20; RPL27A; RPS15A; RPL6; RPS23; RPL12; RPS21; RPL9; RPS3; RPS6; RPS12; RPS15; RPS28; RPLP2; RPL14; RPL27; RPL15; RPL24; RPS27A; RPL21; EIF4B; RPL18A; RPL29; RPL23; RPL35A; RPL10; RPSA; RPS5; RPL7A; RPLP0; RPL18; RPL13; RPL8; RPL13A; RPL32; RPL31; RPL22; RPS4X; EIF3E; RPS16; RPL19; RPL38; RPS9; RPL7; RPL34; EIF3F; EIF3H; RPL10A; RPL5; EIF3L; RPS13; RPL4; RPL3; RPS14 |
| R-HSA-192823 | Viral mRNA Translation | 87 | -8.51E-01 | -3.23 | 1.00E-10 | 4.42E-09 | RPL35; RPS29; RPS17; UBA52; RPL22L1; RPLP1; RPL37; RPS2; FAU; RPS19; RPS27; RPL11; RPS3A; RPS8; RPS25; RPS18; RPL39; RPS20; RPL27A; RPS15A; RPL6; RPS23; RPL12; RPS21; RPL9; RPS3; RPS6; RPS12; RPS15; RPS28; RPLP2; RPL14; RPL27; RPL15; RPL24; RPS27A; RPL21; RPL18A; RPL29; RPL23; RPL35A; RPL10; RPSA; RPS5; RPL7A; RPLP0; RPL18; RPL13; RPL8; RPL13A; RPL32; RPL31; RPL22; RPS4X; RPS16; RPL19; RPL38; RPS9; RPL7; RPL34; RPL10A; RPL5; RPS13; RPL4; RPL3; RPS14 |
| R-HSA-2408557 | Selenocysteine synthesis | 91 | -8.45E-01 | -3.23 | 1.00E-10 | 4.42E-09 | RPL35; SECISBP2; RPS29; RPS17; UBA52; RPL22L1; RPLP1; RPL37; RPS2; FAU; RPS19; RPS27; RPL11; RPS3A; RPS8; RPS25; RPS18; RPL39; RPS20; RPL27A; RPS15A; RPL6; RPS23; RPL12; RPS21; RPL9; RPS3; RPS6; RPS12; RPS15; RPS28; RPLP2; RPL14; RPL27; RPL15; RPL24; RPS27A; RPL21; RPL18A; RPL29; RPL23; RPL35A; RPL10; RPSA; RPS5; RPL7A; RPLP0; RPL18; RPL13; RPL8; RPL13A; RPL32; RPL31; RPL22; RPS4X; RPS16; RPL19; RPL38; RPS9; RPL7; RPL34; RPL10A; RPL5; RPS13; RPL4; RPL3; RPS14 |
| R-HSA-975956 | Nonsense Mediated Decay (NMD) independent of the Exon Junction Complex (EJC) | 93 | -8.46E-01 | -3.24 | 1.00E-10 | 4.42E-09 | RPL35; RPS29; RPS17; UBA52; RPL22L1; RPLP1; RPL37; RPS2; FAU; RPS19; RPS27; RPL11; RPS3A; RPS8; RPS25; RPS18; RPL39; RPS20; RPL27A; RPS15A; RPL6; RPS23; RPL12; RPS21; RPL9; RPS3; RPS6; RPS12; RPS15; RPS28; RPLP2; RPL14; RPL27; RPL15; RPL24; RPS27A; RPL21; RPL18A; RPL29; RPL23; RPL35A; RPL10; RPSA; RPS5; RPL7A; RPLP0; RPL18; RPL13; RPL8; RPL13A; RPL32; RPL31; RPL22; RPS4X; RPS16; RPL19; RPL38; RPS9; RPL7; RPL34; RPL10A; RPL5; RPS13; RPL4; RPL3; RPS14 |
| R-HSA-72764 | Eukaryotic Translation Termination | 91 | -8.49E-01 | -3.25 | 1.00E-10 | 4.42E-09 | RPL35; RPS29; RPS17; UBA52; RPL22L1; RPLP1; RPL37; RPS2; FAU; RPS19; RPS27; RPL11; RPS3A; RPS8; RPS25; RPS18; RPL39; RPS20; RPL27A; RPS15A; RPL6; RPS23; RPL12; RPS21; RPL9; RPS3; RPS6; RPS12; RPS15; RPS28; RPLP2; RPL14; RPL27; RPL15; RPL24; RPS27A; RPL21; RPL18A; RPL29; RPL23; RPL35A; RPL10; RPSA; RPS5; RPL7A; RPLP0; RPL18; RPL13; RPL8; RPL13A; RPL32; RPL31; RPL22; RPS4X; RPS16; RPL19; RPL38; RPS9; RPL7; RPL34; RPL10A; RPL5; RPS13; RPL4; RPL3; RPS14 |
| R-HSA-9633012 | Response of EIF2AK4 (GCN2) to amino acid deficiency | 99 | -8.43E-01 | -3.26 | 1.00E-10 | 4.42E-09 | RPL35; RPS29; RPS17; UBA52; RPL22L1; RPLP1; RPL37; RPS2; FAU; RPS19; RPS27; RPL11; RPS3A; RPS8; RPS25; RPS18; RPL39; RPS20; RPL27A; RPS15A; RPL6; RPS23; RPL12; RPS21; RPL9; RPS3; RPS6; RPS12; RPS15; DDIT3; RPS28; RPLP2; RPL14; ATF3; RPL27; RPL15; RPL24; RPS27A; RPL21; RPL18A; RPL29; RPL23; RPL35A; RPL10; RPSA; RPS5; RPL7A; RPLP0; RPL18; RPL13; RPL8; RPL13A; RPL32; RPL31; RPL22; RPS4X; RPS16; RPL19; RPL38; RPS9; RPL7; RPL34; RPL10A; RPL5; RPS13; RPL4; RPL3; RPS14 |
| R-HSA-72689 | Formation of a pool of free 40S subunits | 99 | -8.44E-01 | -3.26 | 1.00E-10 | 4.42E-09 | RPL35; RPS29; RPS17; UBA52; RPL22L1; RPLP1; RPL37; RPS2; EIF3I; FAU; EIF3D; RPS19; RPS27; RPL11; EIF3M; RPS3A; RPS8; RPS25; RPS18; RPL39; RPS20; RPL27A; RPS15A; RPL6; RPS23; RPL12; RPS21; RPL9; RPS3; RPS6; RPS12; RPS15; RPS28; RPLP2; RPL14; RPL27; RPL15; RPL24; RPS27A; RPL21; RPL18A; RPL29; RPL23; RPL35A; RPL10; RPSA; RPS5; RPL7A; RPLP0; RPL18; RPL13; RPL8; RPL13A; RPL32; RPL31; RPL22; RPS4X; EIF3E; RPS16; RPL19; RPL38; RPS9; RPL7; RPL34; EIF3F; EIF3H; RPL10A; RPL5; EIF3L; RPS13; RPL4; RPL3; RPS14 |
| R-HSA-156902 | Peptide chain elongation | 87 | -8.61E-01 | -3.27 | 1.00E-10 | 4.42E-09 | RPL35; RPS29; RPS17; UBA52; RPL22L1; RPLP1; RPL37; RPS2; FAU; RPS19; RPS27; RPL11; RPS3A; RPS8; RPS25; RPS18; RPL39; RPS20; RPL27A; RPS15A; RPL6; RPS23; RPL12; RPS21; RPL9; RPS3; RPS6; RPS12; RPS15; RPS28; RPLP2; RPL14; EEF2; RPL27; RPL15; RPL24; RPS27A; RPL21; RPL18A; RPL29; RPL23; RPL35A; RPL10; RPSA; RPS5; RPL7A; RPLP0; RPL18; RPL13; RPL8; RPL13A; RPL32; RPL31; RPL22; RPS4X; RPS16; RPL19; RPL38; RPS9; EEF1A1; RPL7; RPL34; RPL10A; RPL5; RPS13; RPL4; RPL3; RPS14 |
| R-HSA-156842 | Eukaryotic Translation Elongation | 90 | -8.59E-01 | -3.27 | 1.00E-10 | 4.42E-09 | RPL35; RPS29; RPS17; UBA52; RPL22L1; RPLP1; RPL37; RPS2; FAU; RPS19; RPS27; RPL11; RPS3A; RPS8; RPS25; RPS18; RPL39; RPS20; RPL27A; RPS15A; RPL6; RPS23; RPL12; RPS21; RPL9; RPS3; RPS6; RPS12; RPS15; RPS28; RPLP2; RPL14; EEF1D; EEF2; RPL27; RPL15; RPL24; RPS27A; RPL21; RPL18A; EEF1B2; RPL29; RPL23; RPL35A; RPL10; RPSA; RPS5; RPL7A; RPLP0; RPL18; RPL13; RPL8; RPL13A; RPL32; RPL31; RPL22; RPS4X; RPS16; RPL19; RPL38; RPS9; EEF1A1; RPL7; RPL34; RPL10A; RPL5; RPS13; RPL4; RPL3; RPS14 |

**Supplementary Table 5B:** *ALL LOW vs MIXED (Main difference: Reservoir Total)*

| **ID** | **Description** | **Set Size** | **Enrichment Score (ES)** | **Normalized Enrichment Score (NES)** | **P-Value** | **Adjusted P-Value** | **Genes** |
| --- | --- | --- | --- | --- | --- | --- | --- |
| R-HSA-983189 | Kinesins | 46 | 6.25E-01 | 2.03E+00 | 4.14E-05 | 1.78E-03 | KIF3B; TUBB4A; KIFC1; KIF18B; KIF22; KIF21B; KIF21A; CENPE; RACGAP1; KIFC2; KIF15; KIF23; KIF20A; KLC2; KIF5A |
| R-HSA-187687 | Signalling to ERKs | 31 | 6.29E-01 | 1.89E+00 | 7.53E-04 | 2.61E-02 | RAPGEF1; NTRK1; SHC2; MAPK1; HRAS; SOS1; MAPK3; RALA; YWHAB; RALGDS; CRKL; KRAS; MAPK11; CRK; KIDINS220; GRB2; MAP2K2; MAPK14 |
| R-HSA-8856688 | Golgi-to-ER retrograde transport | 117 | 4.75E-01 | 1.81E+00 | 5.17E-05 | 2.15E-03 | KIF3B; TUBB4A; NAPB; KIFC1; BICD2; KIF18B; DCTN2; KIF22; KIF21B; DYNC1H1; KIF21A; GBF1; SURF4; CENPE; RACGAP1; KIFC2; ACTR10; KIF15; KIF23; COPB2; KIF20A; DYNC1LI1; KLC2; KIF5A; PLA2G6; COPG1; DCTN1; ARFGAP1; PAFAH1B1; COPA; TUBA4A; ZW10; ACTR1A; RAB3GAP2; KLC4; ARFGAP3; ARF3; KIF4A; KDELR1; DYNC1LI2; KIF3A; KIF11; COPZ1; RAB1B; KLC1; DCTN3; TMED2; RAB6A; RINT1; CAPZA1; KIF2A; TMED9 |
| R-HSA-6811434 | COPI-dependent Golgi-to-ER retrograde traffic | 85 | 4.83E-01 | 1.75E+00 | 3.82E-04 | 1.50E-02 | KIF3B; TUBB4A; NAPB; KIFC1; KIF18B; KIF22; KIF21B; KIF21A; GBF1; SURF4; CENPE; RACGAP1; KIFC2; KIF15; KIF23; COPB2; KIF20A; KLC2; KIF5A; COPG1; ARFGAP1; COPA; TUBA4A; ZW10; KLC4; ARFGAP3; ARF3; KIF4A; KDELR1; KIF3A; KIF11; COPZ1; RAB1B; KLC1; TMED2; RINT1; KIF2A; TMED9 |
| R-HSA-199977 | ER to Golgi Anterograde Transport | 139 | 4.26E-01 | 1.67E+00 | 4.62E-04 | 1.73E-02 | TUBB4A; NAPB; COG2; DCTN2; LMAN2L; SEC23IP; TRAPPC2B; MIA2; TBC1D20; DYNC1H1; GOLGA2; SEC16A; GBF1; PPP6R1; ACTR10; TMEM115; PREB; COPB2; SEC24D; DYNC1LI1; SPTAN1; COPG1; LMAN1; LMAN2; ANKRD28; DCTN1; ARFGAP1; GOLGB1; TRAPPC6B; F8; TRAPPC5; SEC23A; TRAPPC3; TRAPPC1; SEC24C; CNIH3; YKT6; COPA; TUBA4A; ACTR1A; SEC24A; CTSC; STX5; ARFGAP3; SEC22C; SPTB; ARF3; COG3; KDELR1; DYNC1LI2; COPZ1; RAB1B; DCTN3; TGFA; TMED2 |
| R-HSA-198933 | Immunoregulatory interactions between a Lymphoid and a non-Lymphoid cell | 114 | 4.31E-01 | 1.64E+00 | 9.64E-04 | 3.05E-02 | CD8A; VCAM1; CD8B; NPDC1; ITGA4; CD81; CRTAM; HLA-C; SLAMF6; ITGB7; CD3G; SIGLEC12; ITGAL; KLRD1; KLRK1; KLRK1; SFTPD; SH2D1A; LILRB5; CD200R1 |
| R-HSA-948021 | Transport to the Golgi and subsequent modification | 162 | 3.93E-01 | 1.58E+00 | 1.17E-03 | 3.49E-02 | TUBB4A; NAPB; B4GALT2; MGAT3; COG2; DCTN2; MAN1A1; LMAN2L; SEC23IP; ST8SIA6; TRAPPC2B; MIA2; TBC1D20; DYNC1H1; GOLGA2; SEC16A; GBF1; PPP6R1; ACTR10; TMEM115; PREB; COPB2; SEC24D; B4GALT5; DYNC1LI1; SPTAN1; COPG1; LMAN1; CHST10; LMAN2; ANKRD28; DCTN1; ST6GAL1; ARFGAP1; GOLGB1; TRAPPC6B; F8; TRAPPC5; SEC23A; TRAPPC3; TRAPPC1; SEC24C; CNIH3; B4GALT3; YKT6; COPA; TUBA4A; ACTR1A; SEC24A; CTSC; STX5; ARFGAP3; SEC22C; SPTB; ARF3; COG3; KDELR1; DYNC1LI2; COPZ1; RAB1B; DCTN3; TGFA; TMED2; ST3GAL4 |
| R-HSA-6811442 | Intra-Golgi and retrograde Golgi-to-ER traffic | 184 | 3.81E-01 | 1.55E+00 | 8.21E-04 | 2.78E-02 | KIF3B; TUBB4A; NAPB; KIFC1; COG2; BICD2; KIF18B; DCTN2; MAN1A1; KIF22; KIF21B; GOLGA5; DYNC1H1; KIF21A; GBF1; SURF4; CENPE; RACGAP1; KIFC2; ACTR10; KIF15; RAB41; KIF23; COPB2; STX6; KIF20A; M6PR; DYNC1LI1; KLC2; KIF5A; PLA2G6; COPG1; DCTN1; ARFGAP1; PAFAH1B1; VPS52; YKT6; RAB36; TRIP11; COPA; SCOC; TUBA4A; ZW10; ACTR1A; RAB3GAP2; IGF2R; STX5; KLC4; ARFGAP3; CYTH3; ARF3; COG3; KIF4A; KDELR1; DYNC1LI2; KIF3A; KIF11; COPZ1; ARFIP2; RAB1B; KLC1; ARFRP1; DCTN3; TMED2; GOLGA1; RAB6A; VAMP4; RINT1; CAPZA1; GOSR1; KIF2A |
| R-HSA-446203 | Asparagine N-linked glycosylation | 274 | 3.62E-01 | 1.53E+00 | 1.82E-04 | 7.39E-03 | TUBB4A; NAPB; B4GALT2; ST8SIA1; MGAT3; COG2; DCTN2; ALG1; SLC35C1; MAN1A1; SLC35A1; LMAN2L; STT3A; SEC23IP; ST8SIA6; TRAPPC2B; MIA2; TBC1D20; MOGS; GMPPB; UGGT1; DYNC1H1; GOLGA2; SEC16A; GBF1; PPP6R1; SYVN1; ACTR10; TMEM115; PREB; COPB2; SEC24D; B4GALT5; DYNC1LI1; SPTAN1; ALG2; SEL1L; GFPT2; MLEC; ALG5; COPG1; ENGASE; LMAN1; CHST10; RNF5; ALG14; DPAGT1; LMAN2; ANKRD28; DCTN1; ST6GAL1; ARFGAP1; RNF185; GOLGB1; TRAPPC6B; F8; TRAPPC5; SEC23A; TRAPPC3; ST8SIA4; TRAPPC1; SEC24C; CNIH3; ST6GALNAC2; B4GALT3; YKT6; PMM1; COPA; MAGT1; TUBA4A; ACTR1A; DERL2; FUOM; SEC24A; CTSC; STX5; GLB1; ARFGAP3; SEC22C; SPTB; ARF3; EDEM2; COG3; ST3GAL2; KDELR1; DYNC1LI2; ST6GALNAC1; MVD; COPZ1; RAB1B; DCTN3; TGFA; TMED2; ST3GAL4 |
| R-HSA-9675108 | Nervous system development | 470 | -3.08E-01 | -1.36E+00 | 9.57E-04 | 3.05E-02 | NCAM1; DAB1; ARHGEF11; RPS11; RPS28; SLIT1; RPS8; TUBA1A; RPS27; CDC42; RPS21; PLCG1; SEMA6A; TUBB1; ANK1; RPLP0; RPS12; RPL26; MYH10; EPHB4; PSMD7; PRNP; RPS6KA2; ABLIM1; RPS15; LAMB1; RPLP2; RPL35; RPS6KA5; RPS24; RPL36AL; RPL13A; RPS16; MYL9; CDK5R1; RPL18; ABLIM2; ANK3; PSMB1; RPL3; RPS25; ABL2; IRS2; ADGRV1; RPS17; CACNA1C; EFNA1; DLG1; PSMA6; RPL41; RGMB; PSMB3; RPS4X; RPL4; ADAM10; RPL27; YES1; RPS27A; RPL15; RPL37; RPL39L; EPHB3; FAU; RND1; RPL10; ITGA2; NEO1; RPL5; RPS3A; RPL38; MYL6; SEMA4D; PSMB7; SRGAP3; RPL21; LHX4; PFN2; HSP90AA1; RPS5; ITSN1; NRCAM; RPL11; RPL19; PSMA3; RPS23; RPL24; RPL29; RPL35A; RPS13; RPL23; SIAH1; RPSA; SEMA5A; SCN4B; RPL31; NAB2; RPL32; RPS9; PRKCA; RPS15A; RPL12; RPL10A; PSME2; RPS14; RPL14; TRPC1; SRGAP1; RPL22; HSP90AB1; PTK2; EGR2; CACNA1I; RPL9; RPL7; CACNA1H; RPL34; ROBO1; TIAM1; HSPA8; EPHB6; EPHA1 |
| R-HSA-422475 | Axon guidance | 452 | -3.14E-01 | -1.38E+00 | 1.43E-03 | 4.09E-02 | NCAM1; DAB1; ARHGEF11; RPS11; RPS28; SLIT1; RPS8; TUBA1A; RPS27; CDC42; RPS21; PLCG1; SEMA6A; TUBB1; ANK1; RPLP0; RPS12; RPL26; MYH10; EPHB4; PSMD7; PRNP; RPS6KA2; ABLIM1; RPS15; LAMB1; RPLP2; RPL35; RPS6KA5; RPS24; RPL36AL; RPL13A; RPS16; MYL9; CDK5R1; RPL18; ABLIM2; ANK3; PSMB1; RPL3; RPS25; ABL2; IRS2; RPS17; CACNA1C; EFNA1; DLG1; PSMA6; RPL41; RGMB; PSMB3; RPS4X; RPL4; ADAM10; RPL27; YES1; RPS27A; RPL15; RPL37; RPL39L; EPHB3; FAU; RND1; RPL10; ITGA2; NEO1; RPL5; RPS3A; RPL38; MYL6; SEMA4D; PSMB7; SRGAP3; RPL21; LHX4; PFN2; HSP90AA1; RPS5; ITSN1; NRCAM; RPL11; RPL19; PSMA3; RPS23; RPL24; RPL29; RPL35A; RPS13; RPL23; SIAH1; RPSA; SEMA5A; SCN4B; RPL31; RPL32; RPS9; PRKCA; RPS15A; RPL12; RPL10A; PSME2; RPS14; RPL14; TRPC1; SRGAP1; RPL22; HSP90AB1; PTK2; CACNA1I; RPL9; RPL7; CACNA1H; RPL34; ROBO1; TIAM1; HSPA8; EPHB6; EPHA1 |
| R-HSA-913531 | Interferon Signaling | 233 | -3.68E-01 | -1.52E+00 | 7.53E-04 | 2.61E-02 | IRF8; PTPN2; IFNGR2; RPS27A; BST2; HLA-A; OASL; SEH1L; ABCE1; NUP54; HLA-DPA1; IFIT5; FANCC; UBE2L6; IRF1; STAT1; IFIT1; OAS3; IFI35; HLA-DRB1; ICAM1; IFITM3; HLA-DRA; GBP1; IFI6; HERC5; CAMK2D; IFIT2; MX1; TRIM22; FCGR1A; NPM1; IFI30; HSPA8; ISG15; IFIT3; RSAD2; TRIM2; EGR1; HSPA1B |
| R-HSA-9692914 | SARS-CoV-1-host interactions | 93 | -4.36E-01 | -1.60E+00 | 1.53E-03 | 4.30E-02 | IFIH1; RPS18; RPS3; UBA52; RPS6; NFKBIA; YWHAE; RPS20; HNRNPA1; TOMM70; RPS11; RPS28; RPS8; RPS27; RPS21; RPS12; RPS15; RPS24; RPS16; RPS25; STING1; RPS17; EEF1A1; RPS4X; RPS27A; BST2; FAU; RPS3A; RPS5; RPS23; RPS13; RPSA; RPS9; RPS15A; RPS14; NLRP3; NPM1; RCAN3; PALS1 |
| R-HSA-72766 | Translation | 287 | -3.90E-01 | -1.66E+00 | 1.68E-05 | 7.93E-04 | SRPRB; AARS2; MRPL1; RPS11; RPS28; MRPS7; MRPL38; RPS8; RPS27; EIF1AX; RPS21; FARSB; MRPL48; EIF2S3; MRPL42; RPLP0; RPS12; RPL26; DAP3; RPS15; MRPS23; MRPS24; RPLP2; RPL35; RPS24; RPL36AL; EIF3I; RPL13A; RPS16; RPL18; RPL3; MRPS25; RPS25; EIF3L; EEF1D; RPS17; EIF5; EIF3F; RPL41; EIF3M; EEF1A1; RPS4X; RPL4; RPL27; EIF3D; RPS27A; RPL15; RPL37; RPL39L; FAU; RPL10; RPL5; MRPL11; RPS3A; RPL38; RPL21; RPS5; RPL11; RPL19; RPS23; MRPS33; EEF1B2; RPL24; RPL29; RPL35A; RPS13; WARS1; RPL23; RPSA; RPL31; RPL32; RPS9; RPS15A; RPL12; EIF3H; RPL10A; RPS14; RPL14; RPL22; AIMP1; EIF3E; RPL9; RPL7; RPL34; KARS1; PPA1 |
| R-HSA-2122947 | NOTCH1 Intracellular Domain Regulates Transcription | 44 | -5.44E-01 | -1.75E+00 | 1.26E-03 | 3.66E-02 | UBA52; CCNC; MAML3; HDAC2; KAT2A; HDAC10; HDAC4; TLE3; HIF1A; RPS27A; TLE2; MYC; HES1; CUL1; SKP1; MAML2; NBEA |
| R-HSA-71291 | Metabolism of amino acids and derivatives | 301 | -4.23E-01 | -1.81E+00 | 1.76E-07 | 8.85E-06 | RPL13; HAAO; RPS20; PSME1; HIBCH; PSMB4; DCT; RPL39; SERINC1; ACAT1; ALDH6A1; SLC3A2; PPM1K; RPS11; RPS28; RPS8; RPS27; GNMT; RPS21; AZIN1; RPLP0; RPS12; RPL26; PSMD7; HYKK; RPS15; RPLP2; RPL35; RPS24; RPL36AL; RPL13A; RPS16; RPL18; PSMB1; ASL; RPL3; RPS25; GAMT; FAH; RPS17; PSMA6; SLC5A5; RPL41; SECISBP2; PSMB3; RPS4X; RPL4; RPL27; GPT; AMD1; RPS27A; RPL15; RPL37; RPL39L; FAU; RPL10; RPL5; RPS3A; RPL38; ALDH7A1; PSMB7; BCKDHB; MTAP; SAT1; RPL21; SCLY; HGD; GCAT; RPS5; ADI1; RPL11; RPL19; PSMA3; RPS23; RPL24; RPL29; RPL35A; NAALAD2; RPS13; RPL23; RPSA; RPL31; RPL32; RPS9; DBH; RPS15A; RPL12; MCCC1; RPL10A; PSME2; RPS14; RPL14; ACMSD; RPL22; AIMP1; SERINC5; RPL9; RPL7; RPL34; KARS1; TPH1; SARDH; AASS |
| R-HSA-9660826 | Purinergic signaling in leishmaniasis infection | 25 | -6.55E-01 | -1.86E+00 | 9.81E-04 | 3.05E-02 | HMOX1; P2RX4; HSP90AB1; NLRP3; APP; IL1B |
| R-HSA-9664424 | Cell recruitment (pro-inflammatory response) | 25 | -6.55E-01 | -1.86E+00 | 9.81E-04 | 3.05E-02 | HMOX1; P2RX4; HSP90AB1; NLRP3; APP; IL1B |
| R-HSA-3371571 | HSF1-dependent transactivation | 19 | -6.96E-01 | -1.86E+00 | 1.06E-03 | 3.24E-02 | HSP90AA1; CAMK2D; HSP90AB1; HSPA8; DNAJB1; HSPA1B |
| R-HSA-9031628 | NGF-stimulated transcription | 31 | -6.39E-01 | -1.90E+00 | 5.16E-04 | 1.88E-02 | CDK5R1; ID2; TRIB1; ID1; FOSB; NAB2; EGR3; SGK1; EGR2; RRAD; TPH1; ID3; EGR1 |
| R-HSA-3371568 | Attenuation phase | 13 | -8.04E-01 | -1.96E+00 | 3.92E-04 | 1.50E-02 | HSP90AA1; HSP90AB1; HSPA8; DNAJB1; HSPA1B |
| R-HSA-909733 | Interferon alpha/beta signaling | 63 | -5.75E-01 | -1.98E+00 | 2.86E-05 | 1.30E-03 | STAT2; IFITM1; OAS2; XAF1; HLA-F; EIF2AK2; IFNAR2; IRF7; OAS1; IRF9; MX2; GBP2; IRF8; RPS27A; BST2; HLA-A; OASL; ABCE1; IFIT5; IRF1; STAT1; IFIT1; OAS3; IFI35; IFITM3; IFI6; IFIT2; MX1; ISG15; IFIT3; RSAD2; EGR1 |
| R-HSA-9754678 | SARS-CoV-2 modulates host translation machinery | 50 | -6.24E-01 | -2.05E+00 | 3.23E-05 | 1.43E-03 | RPS18; SNRPB; RPS3; RPS6; RPS20; SNRPD2; RPS11; RPS28; RPS8; RPS27; RPS21; SNRPG; RPS12; RPS15; RPS24; RPS16; RPS25; RPS17; RPS4X; RPS27A; FAU; RPS3A; RPS5; SNRPE; RPS23; RPS13; GEMIN4; RPSA; RPS9; RPS15A; RPS14 |
| R-HSA-376176 | Signaling by ROBO receptors | 199 | -5.02E-01 | -2.06E+00 | 4.62E-09 | 2.81E-07 | GSPT2; NRP1; RPS18; RPL28; PSMB5; PSMB9; RPS3; UBA52; SRC; RPL37A; RPL7A; RPS6; ELOB; RPL13; RPS20; PSME1; PSMB4; RPL39; RPS11; RPS28; SLIT1; RPS8; RPS27; CDC42; RPS21; RPLP0; RPS12; RPL26; PSMD7; RPS15; RPLP2; RPL35; RPS24; RPL36AL; RPL13A; RPS16; RPL18; PSMB1; RPL3; RPS25; ABL2; RPS17; PSMA6; RPL41; PSMB3; RPS4X; RPL4; RPL27; RPS27A; RPL15; RPL37; RPL39L; FAU; RPL10; RPL5; RPS3A; RPL38; PSMB7; SRGAP3; RPL21; LHX4; PFN2; RPS5; RPL11; RPL19; PSMA3; RPS23; RPL24; RPL29; RPL35A; RPS13; RPL23; RPSA; RPL31; RPL32; RPS9; PRKCA; RPS15A; RPL12; RPL10A; PSME2; RPS14; RPL14; SRGAP1; RPL22; RPL9; RPL7; RPL34; ROBO1 |
| R-HSA-8868773 | rRNA processing in the nucleus and cytosol | 190 | -5.32E-01 | -2.17E+00 | 1.00E-10 | 6.63E-09 | RPS18; NOP2; RPL28; RPS3; UBA52; RPL37A; DDX49; MPHOSPH10; RPL7A; BUD23; UTP20; EXOSC9; PNO1; RPS6; RPL13; RPS20; KRR1; RPL39; NOP58; EXOSC5; IMP3; RIOK3; RPS11; RPS28; RPS8; RPS27; RPS21; LAS1L; RPLP0; RPS12; RPL26; LTV1; RPS15; UTP4; RPLP2; RPL35; RPS24; RPL36AL; RPL13A; RPS16; UTP14A; RPL18; RPL3; RPS25; NCL; EXOSC7; EXOSC8; RPS17; RPL41; DDX21; NOC4L; RPS4X; FBL; RPL4; TEX10; RPL27; RPS27A; RPL15; RPL37; RPL39L; FAU; RPL10; RPL5; RPS3A; RPL38; NOB1; NOL11; RPP14; RPL21; RPS5; RPL11; RPL19; RPS23; RPL24; RPL29; RPL35A; RPS13; RPL23; RPSA; UTP18; RPL31; RPL32; RPS9; ERI1; PDCD11; RPS15A; RPL12; RPL10A; RPS14; RPL14; RCL1; RPL22; RPL9; RPL7; RPL34; WDR43 |
| R-HSA-72312 | rRNA processing | 200 | -5.30E-01 | -2.17E+00 | 1.00E-10 | 6.63E-09 | RPS18; NOP2; RPL28; RPS3; UBA52; RPL37A; DDX49; MPHOSPH10; RPL7A; BUD23; UTP20; EXOSC9; PNO1; RPS6; RPL13; RPS20; KRR1; RPL39; NOP58; EXOSC5; IMP3; RIOK3; RPS11; RPS28; RPS8; RPS27; RPS21; LAS1L; RPLP0; RPS12; RPL26; LTV1; RPS15; UTP4; RPLP2; RPL35; RPS24; RPL36AL; RPL13A; RPS16; UTP14A; RPL18; RPL3; RPS25; NCL; EXOSC7; EXOSC8; RPS17; RPL41; DDX21; NOC4L; RPS4X; FBL; RPL4; TEX10; RPL27; RPS27A; RPL15; RPL37; RPL39L; FAU; RPL10; RPL5; RPS3A; RPL38; NOB1; NOL11; RPP14; RPL21; RPS5; RPL11; RPL19; RPS23; RPL24; RPL29; RPL35A; RPS13; RPL23; PRORP; RPSA; UTP18; RPL31; RPL32; RPS9; ERI1; PDCD11; RPS15A; RPL12; RPL10A; RPS14; RPL14; TRMT10C; RCL1; RPL22; RPL9; RPL7; RPL34; WDR43 |
| R-HSA-9735869 | SARS-CoV-1 modulates host translation machinery | 36 | -7.16E-01 | -2.21E+00 | 1.27E-06 | 6.16E-05 | RPS18; RPS3; RPS6; RPS20; HNRNPA1; RPS11; RPS28; RPS8; RPS27; RPS21; RPS12; RPS15; RPS24; RPS16; RPS25; RPS17; EEF1A1; RPS4X; RPS27A; FAU; RPS3A; RPS5; RPS23; RPS13; RPSA; RPS9; RPS15A; RPS14 |
| R-HSA-6791226 | Major pathway of rRNA processing in the nucleolus and cytosol | 180 | -5.50E-01 | -2.21E+00 | 1.00E-10 | 6.63E-09 | RPS3; UBA52; RPL37A; DDX49; MPHOSPH10; RPL7A; BUD23; UTP20; EXOSC9; PNO1; RPS6; RPL13; RPS20; KRR1; RPL39; NOP58; EXOSC5; IMP3; RIOK3; RPS11; RPS28; RPS8; RPS27; RPS21; LAS1L; RPLP0; RPS12; RPL26; LTV1; RPS15; UTP4; RPLP2; RPL35; RPS24; RPL36AL; RPL13A; RPS16; UTP14A; RPL18; RPL3; RPS25; NCL; EXOSC7; EXOSC8; RPS17; RPL41; DDX21; NOC4L; RPS4X; FBL; RPL4; TEX10; RPL27; RPS27A; RPL15; RPL37; RPL39L; FAU; RPL10; RPL5; RPS3A; RPL38; NOB1; NOL11; RPP14; RPL21; RPS5; RPL11; RPL19; RPS23; RPL24; RPL29; RPL35A; RPS13; RPL23; RPSA; UTP18; RPL31; RPL32; RPS9; ERI1; PDCD11; RPS15A; RPL12; RPL10A; RPS14; RPL14; RCL1; RPL22; RPL9; RPL7; RPL34; WDR43 |
| R-HSA-9010553 | Regulation of expression of SLITs and ROBOs | 159 | -5.63E-01 | -2.21E+00 | 1.00E-10 | 6.63E-09 | RPS18; RPL28; PSMB5; PSMB9; RPS3; UBA52; RPL37A; RPL7A; RPS6; ELOB; RPL13; RPS20; PSME1; PSMB4; RPL39; RPS11; RPS28; SLIT1; RPS8; RPS27; RPS21; RPLP0; RPS12; RPL26; PSMD7; RPS15; RPLP2; RPL35; RPS24; RPL36AL; RPL13A; RPS16; RPL18; PSMB1; RPL3; RPS25; RPS17; PSMA6; RPL41; PSMB3; RPS4X; RPL4; RPL27; RPS27A; RPL15; RPL37; RPL39L; FAU; RPL10; RPL5; RPS3A; RPL38; PSMB7; RPL21; LHX4; RPS5; RPL11; RPL19; PSMA3; RPS23; RPL24; RPL29; RPL35A; RPS13; RPL23; RPSA; RPL31; RPL32; RPS9; RPS15A; RPL12; RPL10A; PSME2; RPS14; RPL14; RPL22; RPL9; RPL7; RPL34; ROBO1 |
| R-HSA-1799339 | SRP-dependent cotranslational protein targeting to membrane | 110 | -5.99E-01 | -2.25E+00 | 5.02E-10 | 3.19E-08 | RPS18; RPL28; RPS3; UBA52; RPL37A; SRP72; RPL7A; RPS6; RPL13; RPS20; RPL39; SRPRB; RPS11; RPS28; RPS8; RPS27; RPS21; RPLP0; RPS12; RPL26; RPS15; RPLP2; RPL35; RPS24; RPL36AL; RPL13A; RPS16; RPL18; RPL3; RPS25; RPS17; RPL41; RPS4X; RPL4; RPL27; RPS27A; RPL15; RPL37; RPL39L; FAU; RPL10; RPL5; RPS3A; RPL38; RPL21; RPS5; RPL11; RPL19; RPS23; RPL24; RPL29; RPL35A; RPS13; RPL23; RPSA; RPL31; RPL32; RPS9; RPS15A; RPL12; RPL10A; RPS14; RPL14; RPL22; RPL9; RPL7; RPL34 |
| R-HSA-72662 | Activation of the mRNA upon binding of the cap-binding complex and eIFs, and subsequent binding to 43S | 59 | -6.64E-01 | -2.26E+00 | 4.46E-08 | 2.33E-06 | RPS11; RPS28; RPS8; RPS27; EIF1AX; RPS21; EIF2S3; RPS12; RPS15; RPS24; EIF3I; RPS16; RPS25; EIF3L; RPS17; EIF3F; EIF3M; RPS4X; EIF3D; RPS27A; FAU; RPS3A; RPS5; RPS23; RPS13; RPSA; RPS9; RPS15A; EIF3H; RPS14; EIF3E |
| R-HSA-72649 | Translation initiation complex formation | 58 | -6.68E-01 | -2.27E+00 | 3.16E-08 | 1.71E-06 | RPS18; EIF4E; RPS3; RPS6; RPS20; RPS11; RPS28; RPS8; RPS27; EIF1AX; RPS21; EIF2S3; RPS12; RPS15; RPS24; EIF3I; RPS16; RPS25; EIF3L; RPS17; EIF3F; EIF3M; RPS4X; EIF3D; RPS27A; FAU; RPS3A; RPS5; RPS23; RPS13; RPSA; RPS9; RPS15A; EIF3H; RPS14; EIF3E |
| R-HSA-168255 | Influenza Infection | 152 | -5.78E-01 | -2.27E+00 | 1.00E-10 | 6.63E-09 | NUP88; RPL13; RPS20; RPL39; RAN; EIF2AK2; RPS11; RPS28; RPS8; RPS27; RPS21; IPO5; RPLP0; RPS12; RPL26; POLR2C; RPS15; RPLP2; RPL35; RPS24; RPL36AL; RPL13A; RPS16; RPL18; RPL3; RPS25; RPS17; RPL41; RPS4X; RPL4; POLR2G; RPL27; RPS27A; RPL15; RPL37; RPL39L; FAU; SEH1L; RPL10; RPL5; RPS3A; RPL38; NUP54; RPL21; HSP90AA1; RPS5; RPL11; RPL19; RPS23; RPL24; RPL29; RPL35A; RPS13; RPL23; RPSA; RPL31; RPL32; RPS9; RPS15A; RPL12; RPL10A; RPS14; RPL14; RPL22; RPL9; RPL7; RPL34; ISG15; HSPA1B |
| R-HSA-168273 | Influenza Viral RNA Transcription and Replication | 133 | -5.92E-01 | -2.27E+00 | 1.00E-10 | 6.63E-09 | RPS11; RPS28; RPS8; RPS27; RPS21; IPO5; RPLP0; RPS12; RPL26; POLR2C; RPS15; RPLP2; RPL35; RPS24; RPL36AL; RPL13A; RPS16; RPL18; RPL3; RPS25; RPS17; RPL41; RPS4X; RPL4; POLR2G; RPL27; RPS27A; RPL15; RPL37; RPL39L; FAU; SEH1L; RPL10; RPL5; RPS3A; RPL38; NUP54; RPL21; HSP90AA1; RPS5; RPL11; RPL19; RPS23; RPL24; RPL29; RPL35A; RPS13; RPL23; RPSA; RPL31; RPL32; RPS9; RPS15A; RPL12; RPL10A; RPS14; RPL14; RPL22; RPL9; RPL7; RPL34 |
| R-HSA-72702 | Ribosomal scanning and start codon recognition | 58 | -6.73E-01 | -2.28E+00 | 2.18E-08 | 1.27E-06 | RPS11; RPS28; RPS8; RPS27; EIF1AX; RPS21; EIF2S3; RPS12; RPS15; RPS24; EIF3I; RPS16; RPS25; EIF3L; RPS17; EIF5; EIF3F; EIF3M; RPS4X; EIF3D; RPS27A; FAU; RPS3A; RPS5; RPS23; RPS13; RPSA; RPS9; RPS15A; EIF3H; RPS14; EIF3E |
| R-HSA-927802 | Nonsense-Mediated Decay (NMD) | 113 | -6.14E-01 | -2.31E+00 | 1.00E-10 | 6.63E-09 | GSPT2; RPS18; RPL28; RPS3; UBA52; RPL37A; RPL7A; RPS6; RPL13; RPS20; PPP2CA; SMG6; PPP2R2A; RPL39; RPS11; RPS28; RPS8; RPS27; RPS21; RPLP0; RPS12; RPL26; RPS15; RPLP2; RPL35; RPS24; RPL36AL; RPL13A; RPS16; RPL18; RPL3; RPS25; RPS17; RPL41; RPS4X; RPL4; RPL27; RPS27A; RPL15; RPL37; RPL39L; FAU; RPL10; RPL5; RPS3A; RPL38; RPL21; RPS5; RPL11; RPL19; RPS23; RPL24; RPL29; RPL35A; RPS13; RPL23; RPSA; RPL31; RPL32; RPS9; RPS15A; RPL12; RPL10A; RPS14; RPL14; RPL22; RPL9; RPL7; RPL34 |
| R-HSA-975957 | Nonsense Mediated Decay (NMD) enhanced by the Exon Junction Complex (EJC) | 113 | -6.14E-01 | -2.31E+00 | 1.00E-10 | 6.63E-09 | GSPT2; RPS18; RPL28; RPS3; UBA52; RPL37A; RPL7A; RPS6; RPL13; RPS20; PPP2CA; SMG6; PPP2R2A; RPL39; RPS11; RPS28; RPS8; RPS27; RPS21; RPLP0; RPS12; RPL26; RPS15; RPLP2; RPL35; RPS24; RPL36AL; RPL13A; RPS16; RPL18; RPL3; RPS25; RPS17; RPL41; RPS4X; RPL4; RPL27; RPS27A; RPL15; RPL37; RPL39L; FAU; RPL10; RPL5; RPS3A; RPL38; RPL21; RPS5; RPL11; RPL19; RPS23; RPL24; RPL29; RPL35A; RPS13; RPL23; RPSA; RPL31; RPL32; RPS9; RPS15A; RPL12; RPL10A; RPS14; RPL14; RPL22; RPL9; RPL7; RPL34 |
| R-HSA-72695 | Formation of the ternary complex, and subsequently, the 43S complex | 51 | -6.97E-01 | -2.32E+00 | 2.89E-08 | 1.62E-06 | RPS11; RPS28; RPS8; RPS27; EIF1AX; RPS21; EIF2S3; RPS12; RPS15; RPS24; EIF3I; RPS16; RPS25; EIF3L; RPS17; EIF3F; EIF3M; RPS4X; EIF3D; RPS27A; FAU; RPS3A; RPS5; RPS23; RPS13; RPSA; RPS9; RPS15A; EIF3H; RPS14; EIF3E |
| R-HSA-9711097 | Cellular response to starvation | 151 | -5.92E-01 | -2.33E+00 | 1.00E-10 | 6.63E-09 | RPS18; RPL28; ATP6V1D; RPS3; UBA52; RPL37A; RPL7A; LAMTOR4; RPS6; RPL13; FLCN; RPS20; RPL39; ATF4; RPS11; RPS28; RPS8; RPS27; RPS21; EIF2S3; RPLP0; RPS12; RPL26; ATP6V0E2; RPS15; RPLP2; CEBPB; RPL35; RPS24; RPL36AL; RPL13A; RPS16; RPL18; RPL3; RPS25; ATP6V1F; RPS17; RPL41; RPS4X; RPL4; RPL27; RPS27A; RPL15; RPL37; RPL39L; FAU; SEH1L; RPL10; RPL5; RPS3A; RPL38; RPL21; RPS5; ATP6V1G1; RPL11; RPL19; RPS23; RPL24; RPL29; RPL35A; ITFG2; RPS13; RPL23; RPSA; RPL31; ATF3; RPL32; RPS9; RPS15A; RPL12; RPL10A; RPS14; RPL14; FNIP2; RPL22; RPL9; RPL7; DDIT3; RPL34; SESN1 |
| R-HSA-2408522 | Selenoamino acid metabolism | 111 | -6.21E-01 | -2.33E+00 | 1.00E-10 | 6.63E-09 | RPS11; RPS28; RPS8; RPS27; GNMT; RPS21; RPLP0; RPS12; RPL26; RPS15; RPLP2; RPL35; RPS24; RPL36AL; RPL13A; RPS16; RPL18; RPL3; RPS25; RPS17; RPL41; SECISBP2; RPS4X; RPL4; RPL27; RPS27A; RPL15; RPL37; RPL39L; FAU; RPL10; RPL5; RPS3A; RPL38; RPL21; SCLY; RPS5; RPL11; RPL19; RPS23; RPL24; RPL29; RPL35A; RPS13; RPL23; RPSA; RPL31; RPL32; RPS9; RPS15A; RPL12; RPL10A; RPS14; RPL14; RPL22; AIMP1; RPL9; RPL7; RPL34; KARS1 |
| R-HSA-2408557 | Selenocysteine synthesis | 91 | -6.81E-01 | -2.50E+00 | 1.00E-10 | 6.63E-09 | RPS18; RPL28; RPS3; UBA52; RPL37A; RPL7A; RPS6; RPL13; RPS20; RPL39; RPS11; RPS28; RPS8; RPS27; RPS21; RPLP0; RPS12; RPL26; RPS15; RPLP2; RPL35; RPS24; RPL36AL; RPL13A; RPS16; RPL18; RPL3; RPS25; RPS17; RPL41; SECISBP2; RPS4X; RPL4; RPL27; RPS27A; RPL15; RPL37; RPL39L; FAU; RPL10; RPL5; RPS3A; RPL38; RPL21; RPS5; RPL11; RPL19; RPS23; RPL24; RPL29; RPL35A; RPS13; RPL23; RPSA; RPL31; RPL32; RPS9; RPS15A; RPL12; RPL10A; RPS14; RPL14; RPL22; RPL9; RPL7; RPL34 |
| R-HSA-975956 | Nonsense Mediated Decay (NMD) independent of the Exon Junction Complex (EJC) | 93 | -6.88E-01 | -2.52E+00 | 1.00E-10 | 6.63E-09 | GSPT2; RPS18; RPL28; RPS3; UBA52; RPL37A; RPL7A; RPS6; RPL13; RPS20; RPL39; RPS11; RPS28; RPS8; RPS27; RPS21; RPLP0; RPS12; RPL26; RPS15; RPLP2; RPL35; RPS24; RPL36AL; RPL13A; RPS16; RPL18; RPL3; RPS25; RPS17; RPL41; RPS4X; RPL4; RPL27; RPS27A; RPL15; RPL37; RPL39L; FAU; RPL10; RPL5; RPS3A; RPL38; RPL21; RPS5; RPL11; RPL19; RPS23; RPL24; RPL29; RPL35A; RPS13; RPL23; RPSA; RPL31; RPL32; RPS9; RPS15A; RPL12; RPL10A; RPS14; RPL14; RPL22; RPL9; RPL7; RPL34 |
| R-HSA-9633012 | Response of EIF2AK4 (GCN2) to amino acid deficiency | 99 | -6.89E-01 | -2.54E+00 | 1.00E-10 | 6.63E-09 | RPS18; RPL28; RPS3; UBA52; RPL37A; RPL7A; RPS6; RPL13; RPS20; RPL39; ATF4; RPS11; RPS28; RPS8; RPS27; RPS21; EIF2S3; RPLP0; RPS12; RPL26; RPS15; RPLP2; CEBPB; RPL35; RPS24; RPL36AL; RPL13A; RPS16; RPL18; RPL3; RPS25; RPS17; RPL41; RPS4X; RPL4; RPL27; RPS27A; RPL15; RPL37; RPL39L; FAU; RPL10; RPL5; RPS3A; RPL38; RPL21; RPS5; RPL11; RPL19; RPS23; RPL24; RPL29; RPL35A; RPS13; RPL23; RPSA; RPL31; ATF3; RPL32; RPS9; RPS15A; RPL12; RPL10A; RPS14; RPL14; RPL22; RPL9; RPL7; DDIT3; RPL34 |
| R-HSA-72764 | Eukaryotic Translation Termination | 91 | -6.93E-01 | -2.54E+00 | 1.00E-10 | 6.63E-09 | GSPT2; RPS18; RPL28; RPS3; UBA52; RPL37A; RPL7A; RPS6; RPL13; RPS20; RPL39; RPS11; RPS28; RPS8; RPS27; RPS21; RPLP0; RPS12; RPL26; RPS15; RPLP2; RPL35; RPS24; RPL36AL; RPL13A; RPS16; RPL18; RPL3; RPS25; RPS17; RPL41; RPS4X; RPL4; RPL27; RPS27A; RPL15; RPL37; RPL39L; FAU; RPL10; RPL5; RPS3A; RPL38; RPL21; RPS5; RPL11; RPL19; RPS23; RPL24; RPL29; RPL35A; RPS13; RPL23; RPSA; RPL31; RPL32; RPS9; RPS15A; RPL12; RPL10A; RPS14; RPL14; RPL22; RPL9; RPL7; RPL34 |
| R-HSA-72613 | Eukaryotic Translation Initiation | 117 | -6.74E-01 | -2.55E+00 | 1.00E-10 | 6.63E-09 | RPS11; RPS28; RPS8; RPS27; EIF1AX; RPS21; EIF2S3; RPLP0; RPS12; RPL26; RPS15; RPLP2; RPL35; RPS24; RPL36AL; EIF3I; RPL13A; RPS16; RPL18; RPL3; RPS25; EIF3L; RPS17; EIF5; EIF3F; RPL41; EIF3M; RPS4X; RPL4; RPL27; EIF3D; RPS27A; RPL15; RPL37; RPL39L; FAU; RPL10; RPL5; RPS3A; RPL38; RPL21; RPS5; RPL11; RPL19; RPS23; RPL24; RPL29; RPL35A; RPS13; RPL23; RPSA; RPL31; RPL32; RPS9; RPS15A; RPL12; EIF3H; RPL10A; RPS14; RPL14; RPL22; EIF3E; RPL9; RPL7; RPL34 |
| R-HSA-72737 | Cap-dependent Translation Initiation | 117 | -6.74E-01 | -2.55E+00 | 1.00E-10 | 6.63E-09 | RPS11; RPS28; RPS8; RPS27; EIF1AX; RPS21; EIF2S3; RPLP0; RPS12; RPL26; RPS15; RPLP2; RPL35; RPS24; RPL36AL; EIF3I; RPL13A; RPS16; RPL18; RPL3; RPS25; EIF3L; RPS17; EIF5; EIF3F; RPL41; EIF3M; RPS4X; RPL4; RPL27; EIF3D; RPS27A; RPL15; RPL37; RPL39L; FAU; RPL10; RPL5; RPS3A; RPL38; RPL21; RPS5; RPL11; RPL19; RPS23; RPL24; RPL29; RPL35A; RPS13; RPL23; RPSA; RPL31; RPL32; RPS9; RPS15A; RPL12; EIF3H; RPL10A; RPS14; RPL14; RPL22; EIF3E; RPL9; RPL7; RPL34 |
| R-HSA-192823 | Viral mRNA Translation | 87 | -7.03E-01 | -2.56E+00 | 1.00E-10 | 6.63E-09 | RPS18; RPL28; RPS3; UBA52; RPL37A; RPL7A; RPS6; RPL13; RPS20; RPL39; RPS11; RPS28; RPS8; RPS27; RPS21; RPLP0; RPS12; RPL26; RPS15; RPLP2; RPL35; RPS24; RPL36AL; RPL13A; RPS16; RPL18; RPL3; RPS25; RPS17; RPL41; RPS4X; RPL4; RPL27; RPS27A; RPL15; RPL37; RPL39L; FAU; RPL10; RPL5; RPS3A; RPL38; RPL21; RPS5; RPL11; RPL19; RPS23; RPL24; RPL29; RPL35A; RPS13; RPL23; RPSA; RPL31; RPL32; RPS9; RPS15A; RPL12; RPL10A; RPS14; RPL14; RPL22; RPL9; RPL7; RPL34 |
| R-HSA-156842 | Eukaryotic Translation Elongation | 90 | -7.04E-01 | -2.58E+00 | 1.00E-10 | 6.63E-09 | RPS18; RPL28; RPS3; UBA52; RPL37A; RPL7A; RPS6; RPL13; RPS20; RPL39; RPS11; RPS28; RPS8; RPS27; RPS21; RPLP0; RPS12; RPL26; RPS15; RPLP2; RPL35; RPS24; RPL36AL; RPL13A; RPS16; RPL18; RPL3; RPS25; EEF1D; RPS17; RPL41; EEF1A1; RPS4X; RPL4; RPL27; RPS27A; RPL15; RPL37; RPL39L; FAU; RPL10; RPL5; RPS3A; RPL38; RPL21; RPS5; RPL11; RPL19; RPS23; EEF1B2; RPL24; RPL29; RPL35A; RPS13; RPL23; RPSA; RPL31; RPL32; RPS9; RPS15A; RPL12; RPL10A; RPS14; RPL14; RPL22; RPL9; RPL7; RPL34 |
| R-HSA-156902 | Peptide chain elongation | 87 | -7.09E-01 | -2.59E+00 | 1.00E-10 | 6.63E-09 | RPS18; RPL28; RPS3; UBA52; RPL37A; RPL7A; RPS6; RPL13; RPS20; RPL39; RPS11; RPS28; RPS8; RPS27; RPS21; RPLP0; RPS12; RPL26; RPS15; RPLP2; RPL35; RPS24; RPL36AL; RPL13A; RPS16; RPL18; RPL3; RPS25; RPS17; RPL41; EEF1A1; RPS4X; RPL4; RPL27; RPS27A; RPL15; RPL37; RPL39L; FAU; RPL10; RPL5; RPS3A; RPL38; RPL21; RPS5; RPL11; RPL19; RPS23; RPL24; RPL29; RPL35A; RPS13; RPL23; RPSA; RPL31; RPL32; RPS9; RPS15A; RPL12; RPL10A; RPS14; RPL14; RPL22; RPL9; RPL7; RPL34 |
| R-HSA-156827 | L13a-mediated translational silencing of Ceruloplasmin expression | 109 | -6.92E-01 | -2.59E+00 | 1.00E-10 | 6.63E-09 | RPS11; RPS28; RPS8; RPS27; EIF1AX; RPS21; EIF2S3; RPLP0; RPS12; RPL26; RPS15; RPLP2; RPL35; RPS24; RPL36AL; EIF3I; RPL13A; RPS16; RPL18; RPL3; RPS25; EIF3L; RPS17; EIF3F; RPL41; EIF3M; RPS4X; RPL4; RPL27; EIF3D; RPS27A; RPL15; RPL37; RPL39L; FAU; RPL10; RPL5; RPS3A; RPL38; RPL21; RPS5; RPL11; RPL19; RPS23; RPL24; RPL29; RPL35A; RPS13; RPL23; RPSA; RPL31; RPL32; RPS9; RPS15A; RPL12; EIF3H; RPL10A; RPS14; RPL14; RPL22; EIF3E; RPL9; RPL7; RPL34 |
| R-HSA-72706 | GTP hydrolysis and joining of the 60S ribosomal subunit | 110 | -6.93E-01 | -2.60E+00 | 1.00E-10 | 6.63E-09 | RPS11; RPS28; RPS8; RPS27; EIF1AX; RPS21; EIF2S3; RPLP0; RPS12; RPL26; RPS15; RPLP2; RPL35; RPS24; RPL36AL; EIF3I; RPL13A; RPS16; RPL18; RPL3; RPS25; EIF3L; RPS17; EIF5; EIF3F; RPL41; EIF3M; RPS4X; RPL4; RPL27; EIF3D; RPS27A; RPL15; RPL37; RPL39L; FAU; RPL10; RPL5; RPS3A; RPL38; RPL21; RPS5; RPL11; RPL19; RPS23; RPL24; RPL29; RPL35A; RPS13; RPL23; RPSA; RPL31; RPL32; RPS9; RPS15A; RPL12; EIF3H; RPL10A; RPS14; RPL14; RPL22; EIF3E; RPL9; RPL7; RPL34 |
| R-HSA-72689 | Formation of a pool of free 40S subunits | 99 | -7.09E-01 | -2.61E+00 | 1.00E-10 | 6.63E-09 | RPS11; RPS28; RPS8; RPS27; EIF1AX; RPS21; RPLP0; RPS12; RPL26; RPS15; RPLP2; RPL35; RPS24; RPL36AL; EIF3I; RPL13A; RPS16; RPL18; RPL3; RPS25; EIF3L; RPS17; EIF3F; RPL41; EIF3M; RPS4X; RPL4; RPL27; EIF3D; RPS27A; RPL15; RPL37; RPL39L; FAU; RPL10; RPL5; RPS3A; RPL38; RPL21; RPS5; RPL11; RPL19; RPS23; RPL24; RPL29; RPL35A; RPS13; RPL23; RPSA; RPL31; RPL32; RPS9; RPS15A; RPL12; EIF3H; RPL10A; RPS14; RPL14; RPL22; EIF3E; RPL9; RPL7; RPL34 |

**Supplementary Table 5C:** *MIXED vs ALL HIGH (Main difference: Reservoir Intactness)*

| **ID** | **Description** | **Set Size** | **Enrichment Score (ES)** | **Normalized Enrichment Score (NES)** | **P-Value** | **Adjusted P-Value** | **Genes** |
| --- | --- | --- | --- | --- | --- | --- | --- |
| R-HSA-909733 | Interferon alpha/beta signaling | 63 | 8.26E-01 | 2.55 | 1.00E-10 | 4.55E-09 | IFI27; RSAD2; IFIT1; IFIT3; ISG15; IFI6; MX2; OAS3; OAS1; XAF1; USP18; IFI35; IFITM3; OASL; ISG20; OAS2; MX1; STAT2; IFITM1; EIF2AK2; IFITM2; IFIT2; IRF7; IRF9; TYK2; IRF2; IFNAR1; STAT1; GBP2; HLA-E; BST2 |
| R-HSA-913531 | Interferon Signaling | 233 | 6.05E-01 | 2.10 | 1.00E-10 | 4.55E-09 | IFI27; RSAD2; IFIT1; IFIT3; ISG15; IFI6; MX2; OAS3; OAS1; XAF1; HERC5; USP18; IFI35; IFITM3; OASL; ISG20; OAS2; TRIM38; MX1; STAT2; IFITM1; EIF2AK2; TRIM22; IFITM2; IFIT2; IRF7; EIF4G3; TRIM25; UBE2L6; GBP4; FANCL; SP100; TRIM5; IRF9; TYK2; GBP5; RIGI; HLA-DRB1; PML; IRF2; IFNAR1; IFNGR2; STAT1; MAPK1; GBP2; HLA-E; BST2; TRIM21 |
| R-HSA-6783783 | Interleukin-10 signaling | 37 | 7.33E-01 | 2.07 | 9.50E-06 | 3.37E-04 | TIMP1; CXCL2; FPR1; CD86; IL10RB; TYK2; CCR2; IL1R1; PTGS2; FCER2; IL1RN; LIF; TNFRSF1B; CCR1; IL10RA; TNFRSF1A; CXCL1; CCR5; IL1R2 |
| R-HSA-9833110 | RSV-host interactions | 79 | 6.42E-01 | 2.04 | 1.71E-06 | 6.38E-05 | ISG15; SDC2; HERC5; OAS2; STAT2; AGRN; EIF2AK2; TRIM25; UBE2L6; IFIH1; TLR4; TLR6; TYK2; RIGI; CX3CR1; MED12; MED11; IFNAR1; TLR2; MED22; GPC4; TLR7 |
| R-HSA-5357769 | Caspase activation via extrinsic apoptotic signalling pathway | 24 | 7.42E-01 | 1.92 | 2.96E-04 | 7.06E-03 | UNC5B; TLR4; DAPK3; DAPK2; DAPK1; CFLAR; APPL1; FASLG; TNFSF10; LY96; UNC5A; CD14 |
| R-HSA-9664407 | Parasite infection | 57 | 6.31E-01 | 1.91 | 9.11E-05 | 2.41E-03 | VAV2; MYO1C; NCKAP1L; VAV3; SRC; LYN; HCK; ARPC2; MYO9B; ARPC1B; BAIAP2; PTK2; ABL1; MAPK1; CRK; SYK; FCGR3A; WAS; FGR; VAV1; ARPC4; ARPC5 |
| R-HSA-9664417 | Leishmania phagocytosis | 57 | 6.31E-01 | 1.91 | 9.11E-05 | 2.41E-03 | VAV2; MYO1C; NCKAP1L; VAV3; SRC; LYN; HCK; ARPC2; MYO9B; ARPC1B; BAIAP2; PTK2; ABL1; MAPK1; CRK; SYK; FCGR3A; WAS; FGR; VAV1; ARPC4; ARPC5 |
| R-HSA-9664422 | FCGR3A-mediated phagocytosis | 57 | 6.31E-01 | 1.91 | 9.11E-05 | 2.41E-03 | VAV2; MYO1C; NCKAP1L; VAV3; SRC; LYN; HCK; ARPC2; MYO9B; ARPC1B; BAIAP2; PTK2; ABL1; MAPK1; CRK; SYK; FCGR3A; WAS; FGR; VAV1; ARPC4; ARPC5 |
| R-HSA-877300 | Interferon gamma signaling | 84 | 5.94E-01 | 1.91 | 6.27E-05 | 1.83E-03 | OAS3; OAS1; OASL; OAS2; TRIM38; TRIM22; IRF7; TRIM25; GBP4; SP100; TRIM5; IRF9; GBP5; HLA-DRB1; PML; IRF2; IFNGR2; STAT1; MAPK1; GBP2; HLA-E; TRIM21; CIITA; GBP1; TRIM14; FCGR1A; IRF6; HLA-A; RAF1; HLA-DQA2; HLA-DPB1; ICAM1; NCAM1; HLA-DRA |
| R-HSA-1169408 | ISG15 antiviral mechanism | 72 | 6.06E-01 | 1.90 | 4.67E-05 | 1.42E-03 | IFIT1; ISG15; MX2; HERC5; USP18; MX1; EIF2AK2; EIF4G3; TRIM25; UBE2L6; RIGI |
| R-HSA-9833109 | Evasion by RSV of host interferon responses | 21 | 7.47E-01 | 1.89 | 9.17E-04 | 1.76E-02 | STAT2; EIF2AK2; TRIM25; IFIH1; TYK2; RIGI; IFNAR1 |
| R-HSA-198933 | Immunoregulatory interactions between a Lymphoid and a non-Lymphoid cell | 114 | 5.73E-01 | 1.89 | 8.10E-06 | 2.95E-04 | SIGLEC1; IFITM1; SH2D1B; SELL; OSCAR; TREML2; CD19; LILRB3; LILRB3; KLRD1; CD300A; PILRA; SIGLEC12; CD200; LILRB2; MICA; PILRB; CD22; HLA-E; FCGR3A; CD1D; FCGR2B; KIR2DL4; COL1A1; KLRC1; LAIR1; ITGB2; LILRA5; MADCAM1; LILRB1; SLAMF6; COLEC12; CD300E; FCGR1A; CD40; NCR1; HLA-A; KLRF1; TREM1; SIGLEC6; SIGLEC10; KLRK1; KLRK1; CD247; SIGLEC7; KIR2DL3; ICAM1 |
| R-HSA-9820952 | Respiratory Syncytial Virus Infection Pathway | 100 | 5.72E-01 | 1.89 | 1.67E-05 | 5.41E-04 | ISG15; SDC2; HERC5; OAS2; STAT2; AGRN; EIF2AK2; TRIM25; UBE2L6; IFIH1; TLR4; TLR6; TYK2; RIGI; CX3CR1; MED12; MED11; IFNAR1; TLR2; MED22; RAB5B; GPC4; TLR7; FURIN |
| R-HSA-1222556 | ROS and RNS production in phagocytes | 29 | 6.93E-01 | 1.87 | 5.47E-04 | 1.21E-02 | SLC11A1; TCIRG1; CYBA; RAC2; ATP6V0D1; NCF2; HVCN1; NCF4; NCF1; ATP6V0E1; ATP6V1F; ATP6V1C1; ATP6V1A |
| R-HSA-2029480 | Fcgamma receptor (FCGR) dependent phagocytosis | 82 | 5.83E-01 | 1.87 | 6.76E-05 | 1.93E-03 | VAV2; LIMK1; FCGR2A; PLCG2; MYO1C; NCKAP1L; VAV3; SRC; LYN; HCK; ARPC2; MYO9B; ARPC1B; BAIAP2; PTK2; ABL1; MAPK1; CRK; PRKCE; SYK; FCGR3A; ITPR1; AHCYL1; WAS; FGR; VAV1; ARPC4; ARPC5 |
| R-HSA-2029481 | FCGR activation | 12 | 8.33E-01 | 1.85 | 3.75E-04 | 8.67E-03 | FCGR2A; SRC; LYN; HCK; SYK; FCGR3A; FGR; FCGR1A; CD247 |
| R-HSA-9658195 | Leishmania infection | 142 | 5.38E-01 | 1.80 | 1.58E-05 | 5.22E-04 | DVL3; VAV2; RHBDF2; FCGR2A; PLCG2; ADCY4; MYO1C; NCKAP1L; VAV3; SRC; MEFV; LYN; GNG2; HCK; ARPC2; MYO9B; ARPC1B; BAIAP2; NFKB2; DPEP3; CYBA; PTK2; ABL1; MAPK1; CRK; MAPK14; SYK; C3AR1; FCGR3A; CALM3; ITPR1; FURIN; AHCYL1; GNGT2; GNB1; WAS; FGR; GNG5; PRKAR1A; VAV1; ARPC4; ARPC5 |
| R-HSA-9824443 | Parasitic Infection Pathways | 142 | 5.38E-01 | 1.80 | 1.58E-05 | 5.22E-04 | DVL3; VAV2; RHBDF2; FCGR2A; PLCG2; ADCY4; MYO1C; NCKAP1L; VAV3; SRC; MEFV; LYN; GNG2; HCK; ARPC2; MYO9B; ARPC1B; BAIAP2; NFKB2; DPEP3; CYBA; PTK2; ABL1; MAPK1; CRK; MAPK14; SYK; C3AR1; FCGR3A; CALM3; ITPR1; FURIN; AHCYL1; GNGT2; GNB1; WAS; FGR; GNG5; PRKAR1A; VAV1; ARPC4; ARPC5 |
| R-HSA-1912420 | Pre-NOTCH Processing in Golgi | 18 | 7.34E-01 | 1.78 | 2.29E-03 | 4.03E-02 | LFNG; NOTCH1; ST3GAL6; ATP2A1; NOTCH3; ST3GAL4; NOTCH2; ATP2A3; FURIN; SEL1L |
| R-HSA-139853 | Elevation of cytosolic Ca2+ levels | 11 | 8.19E-01 | 1.78 | 1.22E-03 | 2.28E-02 | ORAI2; P2RX5; P2RX1; TRPC3; ITPR1 |
| R-HSA-202733 | Cell surface interactions at the vascular wall | 102 | 5.36E-01 | 1.76 | 1.14E-04 | 2.96E-03 | OLR1; SDC2; SLC16A3; FN1; ITGAX; ITGAV; SELL; SRC; CD58; LYN; SIRPA; PECAM1; CD74; CEACAM3; INPP5D; VPREB3; SELPLG; CD99L2; F11R; COL1A1; ITGB2; PPIL2; SPN; FCER1G; TREM1; ATP1B3; CEACAM1; BSG; ITGAM; PTPN6; SOS1; SHC1; PF4V1; PROCR; CD84; MERTK; TGFB1; SDC3 |
| R-HSA-2029482 | Regulation of actin dynamics for phagocytic cup formation | 59 | 5.69E-01 | 1.73 | 1.48E-03 | 2.70E-02 | VAV2; LIMK1; FCGR2A; MYO1C; NCKAP1L; VAV3; ARPC2; MYO9B; ARPC1B; BAIAP2; PTK2; ABL1; MAPK1; CRK; SYK; FCGR3A; WAS; VAV1; ARPC4; ARPC5 |
| R-HSA-1169410 | Antiviral mechanism by IFN-stimulated genes | 139 | 5.18E-01 | 1.73 | 8.80E-05 | 2.41E-03 | IFIT1; ISG15; MX2; OAS3; OAS1; HERC5; USP18; OASL; OAS2; MX1; EIF2AK2; EIF4G3; TRIM25; UBE2L6; FANCL; RIGI |
| R-HSA-1630316 | Glycosaminoglycan metabolism | 90 | 5.30E-01 | 1.72 | 9.16E-04 | 1.76E-02 | SDC2; CHST15; AGRN; B3GNT7; SLC35B3; ST3GAL6; CHST2; HPSE; DSE; NDST1; ST3GAL4; CHST11; B3GALT6; CHSY1; SLC9A1; B4GALT6; GPC4; HS3ST3B1; HYAL1; HGSNAT; GNS; B4GALT4; B4GALT5; CHST12; ST3GAL2; GLB1; B3GAT1; NAGLU; HS3ST1; HAS1; GUSB; HEXB; CHPF2; SLC26A2 |
| R-HSA-9662851 | Anti-inflammatory response favouring Leishmania parasite infection | 64 | 5.49E-01 | 1.70 | 2.46E-03 | 4.12E-02 | RHBDF2; FCGR2A; PLCG2; ADCY4; SRC; LYN; GNG2; HCK; DPEP3; MAPK14; SYK; FCGR3A; CALM3; ITPR1; FURIN; AHCYL1; GNGT2; GNB1; FGR; GNG5; PRKAR1A; GNG7; FCGR1A; CD247 |
| R-HSA-9664433 | Leishmania parasite growth and survival | 64 | 5.49E-01 | 1.70 | 2.46E-03 | 4.12E-02 | RHBDF2; FCGR2A; PLCG2; ADCY4; SRC; LYN; GNG2; HCK; DPEP3; MAPK14; SYK; FCGR3A; CALM3; ITPR1; FURIN; AHCYL1; GNGT2; GNB1; FGR; GNG5; PRKAR1A; GNG7; FCGR1A; CD247 |
| R-HSA-194138 | Signaling by VEGF | 100 | 5.09E-01 | 1.68 | 1.50E-03 | 2.70E-02 | PTK2B; ITGAV; VAV2; FLT1; NCKAP1L; VAV3; SRC; AXL; AKT1; BAIAP2; CYBA; CTNND1; PTK2; CRK; NRP2; SH2D2A; MAPK14; CALM3; ITPR1; AHCYL1; VEGFB; PGF; NCF2; NCF4; VAV1; NCF1; CTNNA1 |
| R-HSA-168898 | Toll-like Receptor Cascades | 156 | 4.95E-01 | 1.67 | 1.93E-04 | 4.68E-03 | LGMN; USP18; RIPK3; TLR1; MEF2A; PLCG2; IRAK2; IRF7; MEF2C; TLR4; MAP2K4; TLR6; S100A1; BIRC2; MAPK9; ALPK1; NFKB2; AGER; IRAK3; MAPK1; TLR2; TLR10; MAPK14; RPS6KA1; TLR8; FOS; DNM2; TLR7; TANK; MYD88; IRAK1; ITGB2; IKBIP; IRAK4; CTSB; LY96; UNC93B1; IKBKG; NKIRAS2; CD14; TBK1; TP53; NOD2; N4BP1; CTSL |
| R-HSA-6798695 | Neutrophil degranulation | 448 | 4.68E-01 | 1.67 | 2.03E-08 | 7.78E-07 | OLR1; ALOX5; ITGAX; CD53; ITGAV; FCGR3B; NBEAL2; DBNL; FCGR2A; LPCAT1; TNFAIP6; ADAM8; FPR2; SELL; SLC11A1; CXCR2; P2RX1; OSCAR; NCKAP1L; ADGRG3; GPR84; FPR1; HSPA6; CD58; ARHGAP9; DOK3; HPSE; CRISPLD2; IRAG2; MMP25; CR1; AMPD3; SIRPA; IGF2R; LILRB3; LILRB3; NHLRC3; TCIRG1; NFAM1; CTSD; SLCO4C1; PFKL; CD300A; NCSTN; CYBA; PRG2; MME; QSOX1; PSEN1; MGAM; PECAM1; FABP5; OSTF1; MAPK1; TLR2; LRG1; MCEMP1; LILRB2; APAF1; MOSPD2; FUCA1; FTH1; CEACAM3; TMEM179B; MANBA; SLC2A5; RAB5B; SNAP23; TNFRSF1B; CPNE1; MAPK14; PTPRN2; ADGRE3; PTPRJ; C3AR1; CD93; BST2; SLC15A4; SIRPB1; TIMP2; CXCR1; MNDA; CD55; LAIR1; FGR; HGSNAT; SYNGR1; GNS; CXCL1; DNAJC5; ERP44; HVCN1; ITGB2; S100A11; SLC44A2; GRN; CTSB; ORM2; ARPC5; VAT1; PKM; PLAC8; CDA; IDH1; UNC13D; SERPINB1; MVP; CD14; RAB24; GCA; SNAP29; RAB31; QPCT; CTSA; TMEM30A; LAMP1; VNN1; CNN2; FCER1G; ATP11A; LAMP2; RAB27A; FGL2; PADI2; CLEC4D; RAB10; TRAPPC1; GLB1; AGPAT2; CEACAM1; IQGAP1; ADA2; CYSTM1 |
| R-HSA-166016 | Toll Like Receptor 4 (TLR4) Cascade | 137 | 4.84E-01 | 1.62 | 8.59E-04 | 1.69E-02 | USP18; RIPK3; TLR1; MEF2A; PLCG2; IRAK2; IRF7; MEF2C; TLR4; MAP2K4; TLR6; S100A1; BIRC2; MAPK9; ALPK1; NFKB2; AGER; IRAK3; MAPK1; TLR2; MAPK14; RPS6KA1; FOS; DNM2; TANK; MYD88; IRAK1; ITGB2; IKBIP; IRAK4; LY96; IKBKG; NKIRAS2; CD14; TBK1; TP53; NOD2; N4BP1 |
| R-HSA-1989781 | PPARA activates gene expression | 103 | 4.87E-01 | 1.61 | 2.92E-03 | 4.78E-02 | HELZ2; PPARA; FADS1; ABCB4; APOA2; ESRRA; AHRR; RXRA; ACSL1; MED12; MED11; RGL1; PEX11A; RXRB; NCOA3; MED22; PPARG; NCOR2; SP1; ACADM; CPT2; TBL1XR1; MED14; TBL1X; ABCA1; G0S2; MTF1; CDK19; NFYC; CHD9; HDAC3; SLC27A1; MED13 |
| R-HSA-400206 | Regulation of lipid metabolism by PPARalpha | 105 | 4.84E-01 | 1.60 | 2.41E-03 | 4.12E-02 | HELZ2; PPARA; FADS1; ABCB4; APOA2; ESRRA; AHRR; RXRA; ACSL1; MED12; MED11; RGL1; PEX11A; RXRB; NCOA3; MED22; PPARG; NCOR2; SP1; ACADM; CPT2; TBL1XR1; MED14; TBL1X; ABCA1; G0S2; MTF1; CDK19; NFYC; CHD9; HDAC3; SLC27A1; MED13 |
| R-HSA-9013149 | RAC1 GTPase cycle | 172 | 4.43E-01 | 1.51 | 2.92E-03 | 4.78E-02 | DOCK4; MCAM; ARHGAP31; VAV2; ARAP1; ABR; NCKAP1L; DOCK5; VAV3; ARHGAP9; ARAP3; MYO9B; GMIP; BAIAP2L1; PLEKHG2; BAIAP2; PLEKHG1; CYBA; ARHGAP30; ARHGAP27; SNAP23; FMNL1; PREX1; TRIO; FARP2; SRGAP2; SRGAP1; WAS; NCF2; ARHGAP42; NCF4; VAV1; DOCK8; NCF1; SOS2; DEPDC1B; PKN2; SWAP70; ARHGEF11; LBR; ARHGEF7; DEF6; PKN1; IQGAP1; GIT2; PIK3R3; RALBP1; MCF2; FAM13B; RAB7A; ARHGAP20; PARD6A; ARHGAP25; BRK1; SOS1; WIPF1; ; TAOK3; VRK2; ARHGAP23; PAK6; PAK6; NHS; PLEKHG3; FAM13A; VANGL1; PLEKHG6; IQGAP3; EMD; ARHGAP39 |
| R-HSA-76002 | Platelet activation, signaling and aggregation | 213 | 4.23E-01 | 1.47 | 2.42E-03 | 4.12E-02 | SERPING1; FN1; TIMP1; CD109; VAV2; PLCG2; VAV3; THBS1; SRC; AKT1; LYN; GNG2; CYRIB; LGALS3BP; SERPINE1; APLP2; PTK2; QSOX1; GTPBP2; PECAM1; MAPK1; CRK; VTI1B; PRKCE; DGKK; TOR4A; TRPC3; STX4; MAPK14; SYK; NHLRC2; CALM3; FAM3C; STXBP3; ITPR1; VEGFB; RAC2; GNGT2; COL1A1; SRGN; GNB1; GNG5; VAV1; DGKD; MANF; ORM2; PFN1; MAGED2; ARRB1; CD9; GNG7; ARRB2; FCER1G; SCCPDH; GNA12; LAMP2; STXBP2; RAF1; PIK3R3; ACTN4; CD63; PRKCB; HGF; RAPGEF3; PTPN6; RAP1A; APBB1IP; TBXA2R; LCP2; SOS1; PDPK1; SHC1; ITPR2; TLN1; MAPK3; F2R; SERPINA1; CAP1; GNB2; TGFB1; F5; DAGLA |
| R-HSA-109582 | Hemostasis | 472 | 4.10E-01 | 1.46 | 2.96E-05 | 9.25E-04 | OLR1; DOCK4; SERPING1; SDC2; SLC16A3; ORAI2; FN1; TIMP1; ITGAX; ITGAV; CD109; KIF2C; VAV2; P2RX5; PLCG2; SELL; P2RX1; CABLES2; DOCK5; VAV3; THBS1; MFN1; SRC; CD58; CDK2; AKT1; LYN; GNG2; CYRIB; ATP2A1; LGALS3BP; PDE2A; SERPINE1; SIRPA; IRF2; APLP2; PTK2; QSOX1; IRAG1; ABL1; GTPBP2; PECAM1; MAPK1; CRK; PHF21A; VTI1B; PRKCE; CD74; DGKK; CEACAM3; INPP5D; TOR4A; TRPC3; SERPINE2; STX4; VPREB3; MAPK14; SYK; KCNMA1; NHLRC2; SELPLG; SLC8A1; ATP2A3; CD99L2; CALM3; KIF11; FAM3C; H3-3A; F11R; STXBP3; ITPR1; VEGFB; RAC2; GNGT2; COL1A1; SRGN; GNB1; FGR; GNG5; PRKAR1A; MICAL1; ITGB2; VAV1; DGKD; PPIL2; MANF; DOCK8; ORM2; ITGA1; PFN1; MAGED2; ARRB1; CD9; GNG7; KIF1B; SPN; TP53; PAFAH2; CABLES1; ARRB2; FCER1G; SCCPDH; TREM1; GNA12; LAMP2; STXBP2 |
| R-HSA-449147 | Signaling by Interleukins | 382 | 4.08E-01 | 1.45 | 4.30E-04 | 9.80E-03 | HAVCR2; USP18; ALOX5; FN1; TIMP1; ITGAX; STAT2; PTK2B; CXCL2; TXLNA; MEF2A; IRAK2; PSMB9; FPR1; MEF2C; LMNB1; LGALS9; CD86; CSF1R; AKT1; LYN; MAP2K4; IRS1; HCK; IL10RB; TYK2; CCR2; MAPK9; ALPK1; STAT6; NFKB2; IL1R1; PTGS2; AGER; FCER2; CSF3R; SMAD3; IL1RN; IRAK3; PSMB10; LIF; STAT1; MAPK1; CRK; IL2RB; PSMC4; INPP5D; INPPL1; TNFRSF1B; STX4; MAPK14; RPS6KA1; CBL; SYK; IL4R; CCR1; FASLG; PITPNA; RAPGEF1; PSMB3; FOS; IL31RA; MYD88; IRAK1; IL10RA; TNFRSF1A; IL13RA1; GAB2; POMC; CXCL1; ITGB2; IKBIP; VAV1; IRAK4; IL15; CCR5; IL1R2; STX3; PTPN12; SOS2; IKBKG; NKIRAS2; TBK1; TP53; CNN2; NOD2; PSME2; MSN; HIF1A; HNRNPA2B1; STXBP2; N4BP1; BLNK; BCL6; LCP1; PSMB8; STAT5B; ICAM1; PIK3R3; CHUK; BATF; PTPN9; TAB3; MAP3K7; CD80; GSDMD; HGF; ITGAM; PTPN6; DUSP6; IL1RAP; DUSP3; IL17RA; CXCL8; P4HB; IL17C; SOS1; SHC1; S100A12; PSME1; CASP1; POU2F1; MAP2K3; IKBKB; MAPK3; TGFB1; IL27; GSTO1; CEBPD |
| R-HSA-72306 | tRNA processing | 106 | -3.91E-01 | -1.62 | 5.84E-04 | 1.25E-02 | MTO1; CLP1; RTRAF; TRMT10C; EPRS1; NDC1; URM1; TRMT11; RAE1; NUP160; NUP188; TRMT44; GTPBP3; NUP85; TYW1; NSUN6; QTRT1; TRNT1; NUP50; TYW3; THADA; TRMT61A; CDKAL1; TRMT61B; NUP93; TRMT1; ADAT1; ELAC2; METTL1; RAN; TSEN2; DDX1; PUS7 |
| R-HSA-5696399 | Global Genome Nucleotide Excision Repair (GG-NER) | 83 | -4.06E-01 | -1.64 | 1.24E-03 | 2.29E-02 | CHD1L; RFC3; COPS3; POLD2; ACTL6A; DDB1; COPS6; RFC4; RFC1; COPS8; UBE2I; UBA52; NFRKB; COPS4; ERCC4; POLE; RPS27A; COPS2; UBE2N; INO80C; ACTR5; LIG1 |
| R-HSA-5368286 | Mitochondrial translation initiation | 87 | -4.14E-01 | -1.67 | 6.30E-04 | 1.31E-02 | MRPL42; MRPS28; MRPS35; MRPS7; MRPL54; GADD45GIP1; MRPS30; MRPL3; MRPL32; MRPL41; MRPL48; MRPS25; MRPS27; MRPL34; PTCD3; MRPS2; MRPL39; MRPL9; MRPL4; MRPL55; MRPL38; MRPS33; MRPS9; DAP3; MRPL49 |
| R-HSA-5419276 | Mitochondrial translation termination | 87 | -4.19E-01 | -1.69 | 4.83E-04 | 1.08E-02 | MRPL42; MRPS28; MRPS35; MRPS7; MRPL54; GADD45GIP1; MRPS30; MRPL3; MRPL32; MRPL41; MRPL48; MRPS25; MRPS27; MRPL34; PTCD3; MRPS2; MRPL39; MRPL9; MRPL4; MRPL55; MRPL38; MRPS33; MRPS9; DAP3; MRPL49 |
| R-HSA-5368287 | Mitochondrial translation | 93 | -4.15E-01 | -1.72 | 5.94E-04 | 1.25E-02 | MRPS35; GFM1; MRPS7; MRPL54; GADD45GIP1; MRPS30; MRPL3; MRPL32; MRPL41; MRPL48; MRPS25; MRPS27; MRPL34; PTCD3; TSFM; MRPS2; MRPL39; MRPL9; MRPL4; MRPL55; MRPL38; MRPS33; MRPS9; DAP3; MRPL49 |
| R-HSA-6781827 | Transcription-Coupled Nucleotide Excision Repair (TC-NER) | 75 | -4.35E-01 | -1.74 | 6.80E-04 | 1.39E-02 | UVSSA; RFC3; COPS3; AQR; POLD2; TCEA1; DDB1; ERCC6; COPS6; RFC4; RFC1; COPS8; POLR2C; ZNF830; UBA52; COPS4; ERCC4; POLE; RPS27A; COPS2; POLR2D; PRPF19; PPIE; LIG1 |
| R-HSA-9837999 | Mitochondrial protein degradation | 95 | -4.23E-01 | -1.75 | 1.56E-04 | 3.86E-03 | NDUFA2; OMA1; NADK2; HSPD1; NDUFB6; PDK1; ALDH18A1; ACAD8; SMDT1; ACO2; ACAT1; TRIAP1; NDUFS3; LDHD; TIMM10; APP; ALDH2; COX4I1; CLPX; SLC25A5; NDUFA13; UQCRC2; YME1L1; TFAM; IARS2; ATP5MG; ATP5F1B; LONP1; CHCHD2; NDUFV1; ATP5F1C; HSPA9; PMPCA; MRPL32; OXCT1; SUCLG2; MRPS2; GLUD1; IDH3A; ATP5F1A; HADH; ACOT2; PCCB |
| R-HSA-5696398 | Nucleotide Excision Repair | 107 | -4.28E-01 | -1.78 | 2.98E-05 | 9.25E-04 | UVSSA; CHD1L; RFC3; COPS3; AQR; POLD2; ACTL6A; TCEA1; DDB1; ERCC6; COPS6; RFC4; RFC1; COPS8; POLR2C; UBE2I; ZNF830; UBA52; NFRKB; COPS4; ERCC4; POLE; RPS27A; COPS2; UBE2N; INO80C; POLR2D; ACTR5; PRPF19; PPIE; LIG1 |
| R-HSA-5389840 | Mitochondrial translation elongation | 87 | -4.41E-01 | -1.78 | 1.50E-04 | 3.76E-03 | MRPL42; MRPS28; MRPS35; GFM1; MRPS7; MRPL54; GADD45GIP1; MRPS30; MRPL3; MRPL32; MRPL41; MRPL48; MRPS25; MRPS27; MRPL34; PTCD3; TSFM; MRPS2; MRPL39; MRPL9; MRPL4; MRPL55; MRPL38; MRPS33; MRPS9; DAP3; MRPL49 |
| R-HSA-163841 | Gamma carboxylation, hypusinylation, hydroxylation, and arylsulfatase activation | 51 | -4.81E-01 | -1.80 | 7.73E-04 | 1.54E-02 | GAS6; ZC3H15; RPS23; RWDD1; KDM8; PROC; EIF5A2; SUMF2; OGFOD1; SUMF1; RCCD1; DRG1; DPH5; RPL27A; RIOX1; RPS6; RIOX2; DPH1; DPH7; RPL8; EEF2 |
| R-HSA-72203 | Processing of Capped Intron-Containing Pre-mRNA | 280 | -3.82E-01 | -1.81 | 1.18E-08 | 4.63E-07 | AAAS; CWC25; YJU2; RANBP2; SRSF10; POLDIP3; SYF2; CWC27; RNPC3; PQBP1; WBP4; CPSF1; PCBP1; SNRPG; DNAJC8; U2AF2; SNRNP27; NXF1; MTREX; WBP11; SF3B5; SRRT; SRSF5; RBM39; ACIN1; NUDT21; SNRPD2; SYMPK; XAB2; U2SURP; DHX8; CPSF6; SNRNP25; CLP1; DHX15; EIF4E; STEEP1; SF3B2; PRPF18; SRSF11; LSM4; AQR; SNRPC; HNRNPK; NDC1; PRPF4; NXT1; SRSF7; SRSF3; SNRPF; PPIG; WDR33; RAE1; BCAS2; HNRNPD; NUP160; PRKRIP1; SNIP1; HNRNPU; NUP188; DHX35; DDX39B; SNW1; HTATSF1; CWC15; NUP85; SRSF6; PRPF8; LSM5; CRNKL1; SDE2; RNPS1; POLR2C; SRRM1; C9orf78; SRSF8; PCBP2; HNRNPA1; HNRNPH1; NUP50; SNRPD1; ZNF830; SNRPB; SF3A1; PPP1R8; DDX5; THOC3; SNRNP70; SAP18; SNRPA1; SF3A2; SRRM2; RBMX; SNRPE; NUP93; DHX9; DDX42; HSPA8; HNRNPM; WDR70; SNU13; SF1; SF3B1; POLR2D; PRPF19; PPIE |
| R-HSA-9692914 | SARS-CoV-1-host interactions | 93 | -4.41E-01 | -1.83 | 1.16E-04 | 2.97E-03 | RPS17; FAU; TOMM70; RPS11; RPS3A; PPIG; RPS23; YWHAZ; NPM1; RELA; RUNX1; KPNA2; RPS7; RCAN3; RPS25; PPIA; PCBP2; HNRNPA1; RPS27; UBE2I; UBA52; RPS8; PALS1; RPS27L; RPSA; RPS29; RPS20; RPS21; RPS18; YWHAQ; RPS12; RPS5; RPS27A; RPS15; RPS9; RPS6; RPS28; RPS3; RPS4X; RPS19; RPS2; RPS16; EEF1A1; RPS13; RPS14; YWHAG |
| R-HSA-8949613 | Cristae formation | 31 | -5.79E-01 | -1.88 | 9.64E-04 | 1.82E-02 | ATP5F1D; ATP5ME; ATP5PB; ATP5MG; ATP5F1B; ATP5MC2; ATP5F1C; TMEM11; HSPA9; APOO; ATP5MC3; CHCHD6; MICOS10; ATP5F1A; SAMM50 |
| R-HSA-72172 | mRNA Splicing | 213 | -4.17E-01 | -1.88 | 1.00E-08 | 4.18E-07 | SYF2; CWC27; RNPC3; PQBP1; WBP4; PCBP1; SNRPG; DNAJC8; U2AF2; SNRNP27; MTREX; WBP11; SF3B5; SRRT; SRSF5; RBM39; ACIN1; SNRPD2; XAB2; U2SURP; DHX8; SNRNP25; DHX15; STEEP1; SF3B2; PRPF18; SRSF11; LSM4; AQR; SNRPC; HNRNPK; PRPF4; SRSF7; SRSF3; SNRPF; PPIG; BCAS2; HNRNPD; PRKRIP1; SNIP1; HNRNPU; DHX35; DDX39B; SNW1; HTATSF1; CWC15; SRSF6; PRPF8; LSM5; CRNKL1; SDE2; RNPS1; POLR2C; SRRM1; C9orf78; SRSF8; PCBP2; HNRNPA1; HNRNPH1; SNRPD1; ZNF830; SNRPB; SF3A1; PPP1R8; DDX5; SNRNP70; SAP18; SNRPA1; SF3A2; SRRM2; RBMX; SNRPE; DHX9; DDX42; HSPA8; HNRNPM; WDR70; SNU13; SF1; SF3B1; POLR2D; PRPF19; PPIE |
| R-HSA-379726 | Mitochondrial tRNA aminoacylation | 20 | -6.47E-01 | -1.89 | 1.65E-03 | 2.93E-02 | IARS2; AARS2; LARS2; EARS2; QARS1; MARS2; RARS2; VARS2; PPA2; NARS2; KARS1 |
| R-HSA-72163 | mRNA Splicing - Major Pathway | 205 | -4.21E-01 | -1.91 | 6.11E-09 | 2.62E-07 | SYF2; CWC27; PQBP1; WBP4; PCBP1; SNRPG; DNAJC8; U2AF2; SNRNP27; MTREX; WBP11; SF3B5; SRRT; SRSF5; RBM39; ACIN1; SNRPD2; XAB2; U2SURP; DHX8; DHX15; STEEP1; SF3B2; PRPF18; SRSF11; LSM4; AQR; SNRPC; HNRNPK; PRPF4; SRSF7; SRSF3; SNRPF; PPIG; BCAS2; HNRNPD; PRKRIP1; SNIP1; HNRNPU; DHX35; DDX39B; SNW1; HTATSF1; CWC15; SRSF6; PRPF8; LSM5; CRNKL1; SDE2; RNPS1; POLR2C; SRRM1; C9orf78; SRSF8; PCBP2; HNRNPA1; HNRNPH1; SNRPD1; ZNF830; SNRPB; SF3A1; PPP1R8; DDX5; SNRNP70; SAP18; SNRPA1; SF3A2; SRRM2; RBMX; SNRPE; DHX9; DDX42; HSPA8; HNRNPM; WDR70; SNU13; SF1; SF3B1; POLR2D; PRPF19; PPIE |
| R-HSA-3371571 | HSF1-dependent transactivation | 19 | -6.63E-01 | -1.93 | 7.10E-04 | 1.44E-02 | CAMK2G; HSPA1A; HSP90AA1; HSP90AB1; HSPA8; DNAJB1; PTGES3 |
| R-HSA-71291 | Metabolism of amino acids and derivatives | 301 | -4.16E-01 | -1.98 | 1.00E-10 | 4.55E-09 | RPS17; PCBD1; MTAP; FAU; EPRS1; PSMD7; IVD; RPS11; RPS3A; QARS1; RPL35; MRI1; ENOPH1; RPS23; MTR; GRHPR; AIMP2; AFMID; GCDH; RPL14; GLS2; RPS7; PSMD13; PSMC3; RPL36; SERINC1; DBH; PSMC5; AZIN2; ALDH9A1; SLC25A12; PSMB2; AMD1; RPS25; ETHE1; PSMB4; MTRR; RPS27; CKMT2; UBA52; RPS8; RPS27L; RPL22; RPL24; PSMD6; AHCY; PSMD1; AMDHD1; PSMA1; RPL34; RPL39; RPL23; RPSA; GLUD2; RPS29; RPL32; RPL7; RPS20; RPL29; RPL31; RPL35A; RPS21; RPL21; RPS18; RPL22L1; AASS; RPL27; RPL15; RPS12; GLUD1; RPS5; RPLP1; RPS27A; PSMD3; RPS15; RPS9; SERINC5; RPL27A; AIMP1; ALDH7A1; RPL19; RPLP2; RPL10; PSMB5; DLST; RPS6; RPS28; RPS3; RPL10A; RPL6; PSMA7; RPL38; RPS4X; RPS19; RPL18; RPS2; RPL13A; KARS1; RPLP0; RPS16; GAMT; RPL13; RPS13; RPS14; RPL7A; RPL5; RPL8; RPL18A; RPL4; RPL3 |
| R-HSA-3371568 | Attenuation phase | 13 | -7.56E-01 | -1.99 | 5.84E-04 | 1.25E-02 | HSPA1A; HSP90AA1; HSP90AB1; HSPA8; DNAJB1; PTGES3 |
| R-HSA-9629569 | Protein hydroxylation | 19 | -6.86E-01 | -2.00 | 3.38E-04 | 7.93E-03 | ZC3H15; RPS23; RWDD1; KDM8; OGFOD1; RCCD1; DRG1; RPL27A; RIOX1; RPS6; RIOX2; RPL8 |
| R-HSA-1268020 | Mitochondrial protein import | 61 | -5.18E-01 | -2.02 | 5.05E-05 | 1.50E-03 | NDUFB8; ACO2; SLC25A13; HSCB; LDHD; TIMM10; TIMM23; CYC1; GRPEL1; TOMM7; TOMM70; ATP5F1B; BCS1L; CHCHD2; TIMM50; TOMM22; SLC25A12; PMPCB; HSPA9; PMPCA; GRPEL2; TIMM8A; TOMM20; ATP5F1A; SAMM50; FXN; TOMM40 |
| R-HSA-6790901 | rRNA modification in the nucleus and cytosol | 60 | -5.46E-01 | -2.13 | 1.20E-05 | 4.16E-04 | RRP36; WDR43; NOL11; KRR1; NOP58; RPS7; DDX47; IMP4; WDR36; DHX37; HEATR1; UTP15; NOP56; NAT10; BMS1; EMG1; UTP20; UTP4; RPS9; DKC1; RPS6; SNU13; RRP7A; FBL; RPS2; UTP18; RPS14 |
| R-HSA-376176 | Signaling by ROBO receptors | 199 | -5.16E-01 | -2.32 | 1.00E-10 | 4.55E-09 | RPS7; PSMD13; PSMC3; RPL36; PSMC5; PSMB2; RPS25; RNPS1; CDC42; PSMB4; RPS27; UBA52; RPS8; RPS27L; RPL22; RPL24; PSMD6; PSMD1; PSMA1; RPL34; RPL39; RPL23; RPSA; RPS29; RPL32; RPL7; RPS20; RPL29; RPL31; NCK1; RPL35A; RPS21; RPL21; RPS18; RPL22L1; RPL27; RPL15; RPS12; ROBO3; RPS5; RPLP1; RPS27A; PSMD3; RPS15; RPS9; RPL27A; RPL19; RPLP2; RPL10; PSMB5; PRKCA; RPS6; RPS28; RPS3; RPL10A; RPL6; PSMA7; PRKAR2A; RPL38; PFN2; RPS4X; RPS19; RPL18; RPS2; RPL13A; RPLP0; RPS16; RPL13; RPS13; RPS14; RPL7A; RPL5; RPL8; RPL18A; RPL4; RPL3 |
| R-HSA-9735869 | SARS-CoV-1 modulates host translation machinery | 36 | -7.48E-01 | -2.49 | 2.86E-09 | 1.26E-07 | RPS17; FAU; RPS11; RPS3A; RPS23; RPS7; RPS25; HNRNPA1; RPS27; RPS8; RPS27L; RPSA; RPS29; RPS20; RPS21; RPS18; RPS12; RPS5; RPS27A; RPS15; RPS9; RPS6; RPS28; RPS3; RPS4X; RPS19; RPS2; RPS16; EEF1A1; RPS13; RPS14 |
| R-HSA-9754678 | SARS-CoV-2 modulates host translation machinery | 50 | -6.69E-01 | -2.50 | 1.04E-08 | 4.23E-07 | RPS17; FAU; RPS11; RPS3A; SNRPF; GEMIN4; RPS23; RPS7; RPS25; RPS27; SNRPD1; SNRPB; RPS8; RPS27L; DDX20; RPSA; RPS29; RPS20; SNRPE; RPS21; RPS18; RPS12; RPS5; RPS27A; RPS15; RPS9; RPS6; RPS28; RPS3; RPS4X; RPS19; RPS2; RPS16; RPS13; RPS14 |
| R-HSA-9010553 | Regulation of expression of SLITs and ROBOs | 159 | -6.13E-01 | -2.69 | 1.00E-10 | 4.55E-09 | RPS7; PSMD13; PSMC3; RPL36; PSMC5; PSMB2; RPS25; RNPS1; PSMB4; RPS27; UBA52; RPS8; RPS27L; RPL22; RPL24; PSMD6; PSMD1; PSMA1; RPL34; RPL39; RPL23; RPSA; RPS29; RPL32; RPL7; RPS20; RPL29; RPL31; RPL35A; RPS21; RPL21; RPS18; RPL22L1; RPL27; RPL15; RPS12; ROBO3; RPS5; RPLP1; RPS27A; PSMD3; RPS15; RPS9; RPL27A; RPL19; RPLP2; RPL10; PSMB5; RPS6; RPS28; RPS3; RPL10A; RPL6; PSMA7; RPL38; RPS4X; RPS19; RPL18; RPS2; RPL13A; RPLP0; RPS16; RPL13; RPS13; RPS14; RPL7A; RPL5; RPL8; RPL18A; RPL4; RPL3 |
| R-HSA-9711097 | Cellular response to starvation | 151 | -6.04E-01 | -2.69 | 1.00E-10 | 4.55E-09 | RPS27; KPTN; UBA52; RPS8; RPS27L; RPL22; RPL24; SESN1; RPL34; RPL39; RPL23; RPSA; RPS29; RPL32; RPL7; RPS20; RPL29; RPL31; RPL35A; RPS21; RPL21; RPS18; RPL22L1; RPL27; RPL15; RPS12; RPS5; RPLP1; RPS27A; RPS15; RPS9; RPL27A; RPL19; RPLP2; RPL10; RPS6; RPS28; RPS3; RPL10A; RPL6; RPL38; RPS4X; RPS19; RPL18; RPS2; RPL13A; RPLP0; RPS16; RPL13; RPS13; RPS14; RPL7A; RPL5; RPL8; RPL18A; RPL4; RPL3 |
| R-HSA-168255 | Influenza Infection | 152 | -6.06E-01 | -2.70 | 1.00E-10 | 4.55E-09 | RPS17; KPNA3; FAU; NDC1; RPS11; RPS3A; RPL35; RPS23; HSPA1A; RAE1; NUP160; NUP188; RPL14; KPNA2; NUP85; RPS7; RPL36; KPNA5; RPS25; POLR2C; RPS27; NUP50; UBA52; RPS8; RPS27L; RPL22; RPL24; HSP90AA1; RPL34; RPL39; RPL23; RPSA; RPS29; RPL32; RPL7; RPS20; RPL29; RPL31; NUP93; RPL35A; RPS21; RPL21; RPS18; RPL22L1; RPL27; RPL15; RPS12; RPS5; RPLP1; RPS27A; RPS15; RPS9; RPL27A; RAN; RPL19; RPLP2; RPL10; RPS6; RPS28; RPS3; RPL10A; RPL6; GRSF1; RPL38; RPS4X; RPS19; RPL18; RPS2; RPL13A; RPLP0; POLR2D; RPS16; IPO5; RPL13; RPS13; RPS14; RPL7A; RPL5; RPL8; RPL18A; RPL4; RPL3 |
| R-HSA-8868773 | rRNA processing in the nucleus and cytosol | 190 | -6.08E-01 | -2.71 | 1.00E-10 | 4.55E-09 | EXOSC9; RPS17; WDR43; FAU; NOL11; RPS11; RPS3A; RPL35; RIOK1; RPS23; BOP1; KRR1; NOP58; C1D; RIOK2; RPL14; RPS7; RPL36; DDX47; IMP4; WDR36; DHX37; HEATR1; RPS25; UTP15; NOP56; XRN2; RPS27; EXOSC7; UBA52; RPS8; NAT10; BMS1; EXOSC10; RPS27L; RPL22; WDR12; RPL24; EMG1; UTP20; RPL34; RPL39; RPL23; RPSA; RPS29; NOL9; RPL32; RPL7; RPS20; RPL29; UTP4; RPL31; RPL35A; RPS21; RPL21; RPS18; RPL22L1; RPL27; RPL15; RPS12; RPS5; EXOSC6; RPLP1; RPS27A; RPS15; RPS9; RPL27A; DKC1; RPL19; RPLP2; RPL10; RPS6; RPS28; RPS3; RPL10A; RPL6; RPL38; SNU13; RPS4X; RRP7A; FBL; RPS19; RPL18; RPS2; UTP18; RPL13A; RPLP0; RPS16; LAS1L; RPL13; RPS13; RPS14; RPL7A; CSNK1E; TSR1; RPL5; RPL8; RPL18A; RPL4; RPL3 |
| R-HSA-6791226 | Major pathway of rRNA processing in the nucleolus and cytosol | 180 | -6.14E-01 | -2.71 | 1.00E-10 | 4.55E-09 | EXOSC9; RPS17; WDR43; FAU; NOL11; RPS11; RPS3A; RPL35; RIOK1; RPS23; BOP1; KRR1; NOP58; C1D; RIOK2; RPL14; RPS7; RPL36; DDX47; IMP4; WDR36; DHX37; HEATR1; RPS25; UTP15; NOP56; XRN2; RPS27; EXOSC7; UBA52; RPS8; BMS1; EXOSC10; RPS27L; RPL22; WDR12; RPL24; EMG1; UTP20; RPL34; RPL39; RPL23; RPSA; RPS29; NOL9; RPL32; RPL7; RPS20; RPL29; UTP4; RPL31; RPL35A; RPS21; RPL21; RPS18; RPL22L1; RPL27; RPL15; RPS12; RPS5; EXOSC6; RPLP1; RPS27A; RPS15; RPS9; RPL27A; RPL19; RPLP2; RPL10; RPS6; RPS28; RPS3; RPL10A; RPL6; RPL38; SNU13; RPS4X; RRP7A; FBL; RPS19; RPL18; RPS2; UTP18; RPL13A; RPLP0; RPS16; LAS1L; RPL13; RPS13; RPS14; RPL7A; CSNK1E; TSR1; RPL5; RPL8; RPL18A; RPL4; RPL3 |
| R-HSA-72312 | rRNA processing | 200 | -6.02E-01 | -2.72 | 1.00E-10 | 4.55E-09 | TRMT10C; EXOSC9; RPS17; WDR43; FAU; NOL11; RPS11; RPS3A; RPL35; RIOK1; RPS23; BOP1; KRR1; NOP58; C1D; RIOK2; RPL14; MRM3; RPS7; RPL36; DDX47; IMP4; WDR36; DHX37; HEATR1; RPS25; UTP15; NOP56; XRN2; RPS27; EXOSC7; UBA52; RPS8; NAT10; BMS1; EXOSC10; RPS27L; RPL22; WDR12; RPL24; EMG1; UTP20; RPL34; RPL39; RPL23; RPSA; RPS29; NOL9; RPL32; RPL7; RPS20; RPL29; UTP4; RPL31; RPL35A; RPS21; RPL21; RPS18; RPL22L1; RPL27; RPL15; RPS12; ELAC2; RPS5; EXOSC6; RPLP1; RPS27A; RPS15; RPS9; RPL27A; DKC1; RPL19; RPLP2; RPL10; RPS6; RPS28; RPS3; RPL10A; RPL6; RPL38; SNU13; RPS4X; RRP7A; FBL; RPS19; RPL18; RPS2; UTP18; RPL13A; RPLP0; RPS16; LAS1L; RPL13; RPS13; RPS14; RPL7A; CSNK1E; TSR1; RPL5; RPL8; RPL18A; RPL4; RPL3 |
| R-HSA-72766 | Translation | 287 | -5.91E-01 | -2.77 | 1.00E-10 | 4.55E-09 | RPS17; LARS2; FAU; EPRS1; SRP68; EIF4G1; EARS2; EIF3D; MRPL42; EIF3A; EIF3C; RPS11; RPS3A; GSPT1; QARS1; RPL35; EIF4A2; MRPS28; MARS2; RPS23; SSR2; EIF3M; MRPS35; AIMP2; RPL14; GFM1; SRP19; RARS2; MRPS7; MRPL54; RPS7; RPL36; EIF3I; GADD45GIP1; RPS25; PPA1; EIF3K; EIF3B; SSR4; MRPS30; MRPL3; RPS27; EIF3E; UBA52; MRPL32; RPS8; RPS27L; RPL22; MRPL41; SRP72; MRPL48; RPL24; MRPS25; MRPS27; MRPL34; PTCD3; FARSB; RPL34; TSFM; VARS2; RPL39; MRPS2; RPL23; RPSA; RPS29; RPL32; RPL7; EEF1B2; MRPL39; RPS20; RPL29; RPL31; RPL35A; EIF4H; RPS21; RPL21; RPS18; MRPL9; PPA2; RPL22L1; RPL27; RPL15; RPS12; RPS5; MRPL4; RPLP1; RPS27A; RPS15; RPS9; RPL27A; EEF1D; MRPL55; MRPL38; MRPS33; AIMP1; EIF2B3; RPL19; RPLP2; RPL10; NARS2; RPS6; MRPS9; RPS28; RPS3; RPL10A; EIF3H; RPL6; SSR3; RPL38; RPS4X; RPS19; RPL18; RPS2; RPL13A; KARS1; DAP3; RPLP0; MRPL49; RPS16; EEF1A1; EIF3F; RPL13; RPS13; RPS14; RPL7A; RPL5; RPL8; RPL18A; EEF2; RPL4; EIF3L; EIF4B; RPL3 |
| R-HSA-72695 | Formation of the ternary complex, and subsequently, the 43S complex | 51 | -7.41E-01 | -2.78 | 1.00E-10 | 4.55E-09 | RPS17; FAU; EIF3D; EIF3A; EIF3C; RPS11; RPS3A; RPS23; EIF3M; RPS7; EIF3I; RPS25; EIF3K; EIF3B; RPS27; EIF3E; RPS8; RPS27L; RPSA; RPS29; RPS20; RPS21; RPS18; RPS12; RPS5; RPS27A; RPS15; RPS9; RPS6; RPS28; RPS3; EIF3H; RPS4X; RPS19; RPS2; RPS16; EIF3F; RPS13; RPS14; EIF3L |
| R-HSA-72702 | Ribosomal scanning and start codon recognition | 58 | -7.30E-01 | -2.81 | 1.00E-10 | 4.55E-09 | RPS17; FAU; EIF4G1; EIF3D; EIF3A; EIF3C; RPS11; RPS3A; EIF4A2; RPS23; EIF3M; RPS7; EIF3I; RPS25; EIF3K; EIF3B; RPS27; EIF3E; RPS8; RPS27L; RPSA; RPS29; RPS20; EIF4H; RPS21; RPS18; RPS12; RPS5; RPS27A; RPS15; RPS9; RPS6; RPS28; RPS3; EIF3H; RPS4X; RPS19; RPS2; RPS16; EIF3F; RPS13; RPS14; EIF3L; EIF4B |
| R-HSA-72649 | Translation initiation complex formation | 58 | -7.32E-01 | -2.82 | 1.00E-10 | 4.55E-09 | RPS17; FAU; EIF4G1; EIF3D; EIF3A; EIF3C; RPS11; RPS3A; EIF4A2; RPS23; EIF3M; RPS7; EIF3I; RPS25; EIF3K; EIF3B; RPS27; EIF3E; RPS8; RPS27L; RPSA; RPS29; RPS20; EIF4H; RPS21; RPS18; RPS12; RPS5; RPS27A; RPS15; RPS9; RPS6; RPS28; RPS3; EIF3H; RPS4X; RPS19; RPS2; RPS16; EIF3F; RPS13; RPS14; EIF3L; EIF4B |
| R-HSA-72662 | Activation of the mRNA upon binding of the cap-binding complex and eIFs, and subsequent binding to 43S | 59 | -7.29E-01 | -2.84 | 1.00E-10 | 4.55E-09 | RPS17; FAU; EIF4G1; EIF3D; EIF3A; EIF3C; RPS11; RPS3A; EIF4A2; RPS23; EIF3M; RPS7; EIF3I; RPS25; EIF3K; EIF3B; RPS27; EIF3E; RPS8; RPS27L; RPSA; RPS29; RPS20; EIF4H; RPS21; RPS18; RPS12; RPS5; RPS27A; RPS15; RPS9; RPS6; RPS28; RPS3; EIF3H; RPS4X; RPS19; RPS2; RPS16; EIF3F; RPS13; RPS14; EIF3L; EIF4B |
| R-HSA-2408522 | Selenoamino acid metabolism | 111 | -7.09E-01 | -3.02 | 1.00E-10 | 4.55E-09 | RPS27; UBA52; RPS8; RPS27L; RPL22; RPL24; AHCY; RPL34; RPL39; RPL23; RPSA; RPS29; RPL32; RPL7; RPS20; RPL29; RPL31; RPL35A; RPS21; RPL21; RPS18; RPL22L1; RPL27; RPL15; RPS12; RPS5; RPLP1; RPS27A; RPS15; RPS9; RPL27A; AIMP1; RPL19; RPLP2; RPL10; RPS6; RPS28; RPS3; RPL10A; RPL6; RPL38; RPS4X; RPS19; RPL18; RPS2; RPL13A; KARS1; RPLP0; RPS16; RPL13; RPS13; RPS14; RPL7A; RPL5; RPL8; RPL18A; RPL4; RPL3 |
| R-HSA-927802 | Nonsense-Mediated Decay (NMD) | 113 | -7.16E-01 | -3.06 | 1.00E-10 | 4.55E-09 | RPS7; RPL36; PPP2CA; PNRC2; SMG8; RPS25; RNPS1; DCP1A; RPS27; UBA52; RPS8; RPS27L; RPL22; RPL24; RPL34; RPL39; RPL23; RPSA; RPS29; RPL32; RPL7; RPS20; RPL29; RPL31; RPL35A; RPS21; RPL21; RPS18; RPL22L1; RPL27; RPL15; RPS12; RPS5; RPLP1; RPS27A; RPS15; RPS9; RPL27A; SMG6; RPL19; RPLP2; RPL10; RPS6; RPS28; RPS3; RPL10A; RPL6; RPL38; RPS4X; RPS19; RPL18; RPS2; RPL13A; RPLP0; RPS16; RPL13; RPS13; RPS14; RPL7A; RPL5; RPL8; RPL18A; RPL4; RPL3 |
| R-HSA-975957 | Nonsense Mediated Decay (NMD) enhanced by the Exon Junction Complex (EJC) | 113 | -7.16E-01 | -3.06 | 1.00E-10 | 4.55E-09 | RPS7; RPL36; PPP2CA; PNRC2; SMG8; RPS25; RNPS1; DCP1A; RPS27; UBA52; RPS8; RPS27L; RPL22; RPL24; RPL34; RPL39; RPL23; RPSA; RPS29; RPL32; RPL7; RPS20; RPL29; RPL31; RPL35A; RPS21; RPL21; RPS18; RPL22L1; RPL27; RPL15; RPS12; RPS5; RPLP1; RPS27A; RPS15; RPS9; RPL27A; SMG6; RPL19; RPLP2; RPL10; RPS6; RPS28; RPS3; RPL10A; RPL6; RPL38; RPS4X; RPS19; RPL18; RPS2; RPL13A; RPLP0; RPS16; RPL13; RPS13; RPS14; RPL7A; RPL5; RPL8; RPL18A; RPL4; RPL3 |
| R-HSA-168273 | Influenza Viral RNA Transcription and Replication | 133 | -6.94E-01 | -3.07 | 1.00E-10 | 4.55E-09 | RPS27; NUP50; UBA52; RPS8; RPS27L; RPL22; RPL24; HSP90AA1; RPL34; RPL39; RPL23; RPSA; RPS29; RPL32; RPL7; RPS20; RPL29; RPL31; NUP93; RPL35A; RPS21; RPL21; RPS18; RPL22L1; RPL27; RPL15; RPS12; RPS5; RPLP1; RPS27A; RPS15; RPS9; RPL27A; RPL19; RPLP2; RPL10; RPS6; RPS28; RPS3; RPL10A; RPL6; GRSF1; RPL38; RPS4X; RPS19; RPL18; RPS2; RPL13A; RPLP0; POLR2D; RPS16; IPO5; RPL13; RPS13; RPS14; RPL7A; RPL5; RPL8; RPL18A; RPL4; RPL3 |
| R-HSA-1799339 | SRP-dependent cotranslational protein targeting to membrane | 110 | -7.31E-01 | -3.09 | 1.00E-10 | 4.55E-09 | SSR4; RPS27; UBA52; RPS8; RPS27L; RPL22; SRP72; RPL24; RPL34; RPL39; RPL23; RPSA; RPS29; RPL32; RPL7; RPS20; RPL29; RPL31; RPL35A; RPS21; RPL21; RPS18; RPL22L1; RPL27; RPL15; RPS12; RPS5; RPLP1; RPS27A; RPS15; RPS9; RPL27A; RPL19; RPLP2; RPL10; RPS6; RPS28; RPS3; RPL10A; RPL6; SSR3; RPL38; RPS4X; RPS19; RPL18; RPS2; RPL13A; RPLP0; RPS16; RPL13; RPS13; RPS14; RPL7A; RPL5; RPL8; RPL18A; RPL4; RPL3 |
| R-HSA-9633012 | Response of EIF2AK4 (GCN2) to amino acid deficiency | 99 | -7.44E-01 | -3.11 | 1.00E-10 | 4.55E-09 | RPS27; UBA52; RPS8; RPS27L; RPL22; RPL24; RPL34; RPL39; RPL23; RPSA; RPS29; RPL32; RPL7; RPS20; RPL29; RPL31; RPL35A; RPS21; RPL21; RPS18; RPL22L1; RPL27; RPL15; RPS12; RPS5; RPLP1; RPS27A; RPS15; RPS9; RPL27A; RPL19; RPLP2; RPL10; RPS6; RPS28; RPS3; RPL10A; RPL6; RPL38; RPS4X; RPS19; RPL18; RPS2; RPL13A; RPLP0; RPS16; RPL13; RPS13; RPS14; RPL7A; RPL5; RPL8; RPL18A; RPL4; RPL3 |
| R-HSA-72613 | Eukaryotic Translation Initiation | 117 | -7.31E-01 | -3.14 | 1.00E-10 | 4.55E-09 | RPS17; FAU; EIF4G1; EIF3D; EIF3A; EIF3C; RPS11; RPS3A; RPL35; EIF4A2; RPS23; EIF3M; RPL14; RPS7; RPL36; EIF3I; RPS25; EIF3K; EIF3B; RPS27; EIF3E; UBA52; RPS8; RPS27L; RPL22; RPL24; RPL34; RPL39; RPL23; RPSA; RPS29; RPL32; RPL7; RPS20; RPL29; RPL31; RPL35A; EIF4H; RPS21; RPL21; RPS18; RPL22L1; RPL27; RPL15; RPS12; RPS5; RPLP1; RPS27A; RPS15; RPS9; RPL27A; EIF2B3; RPL19; RPLP2; RPL10; RPS6; RPS28; RPS3; RPL10A; EIF3H; RPL6; RPL38; RPS4X; RPS19; RPL18; RPS2; RPL13A; RPLP0; RPS16; EIF3F; RPL13; RPS13; RPS14; RPL7A; RPL5; RPL8; RPL18A; RPL4; EIF3L; EIF4B; RPL3 |
| R-HSA-72737 | Cap-dependent Translation Initiation | 117 | -7.31E-01 | -3.14 | 1.00E-10 | 4.55E-09 | RPS17; FAU; EIF4G1; EIF3D; EIF3A; EIF3C; RPS11; RPS3A; RPL35; EIF4A2; RPS23; EIF3M; RPL14; RPS7; RPL36; EIF3I; RPS25; EIF3K; EIF3B; RPS27; EIF3E; UBA52; RPS8; RPS27L; RPL22; RPL24; RPL34; RPL39; RPL23; RPSA; RPS29; RPL32; RPL7; RPS20; RPL29; RPL31; RPL35A; EIF4H; RPS21; RPL21; RPS18; RPL22L1; RPL27; RPL15; RPS12; RPS5; RPLP1; RPS27A; RPS15; RPS9; RPL27A; EIF2B3; RPL19; RPLP2; RPL10; RPS6; RPS28; RPS3; RPL10A; EIF3H; RPL6; RPL38; RPS4X; RPS19; RPL18; RPS2; RPL13A; RPLP0; RPS16; EIF3F; RPL13; RPS13; RPS14; RPL7A; RPL5; RPL8; RPL18A; RPL4; EIF3L; EIF4B; RPL3 |
| R-HSA-72706 | GTP hydrolysis and joining of the 60S ribosomal subunit | 110 | -7.48E-01 | -3.16 | 1.00E-10 | 4.55E-09 | RPS17; FAU; EIF4G1; EIF3D; EIF3A; EIF3C; RPS11; RPS3A; RPL35; EIF4A2; RPS23; EIF3M; RPL14; RPS7; RPL36; EIF3I; RPS25; EIF3K; EIF3B; RPS27; EIF3E; UBA52; RPS8; RPS27L; RPL22; RPL24; RPL34; RPL39; RPL23; RPSA; RPS29; RPL32; RPL7; RPS20; RPL29; RPL31; RPL35A; EIF4H; RPS21; RPL21; RPS18; RPL22L1; RPL27; RPL15; RPS12; RPS5; RPLP1; RPS27A; RPS15; RPS9; RPL27A; RPL19; RPLP2; RPL10; RPS6; RPS28; RPS3; RPL10A; EIF3H; RPL6; RPL38; RPS4X; RPS19; RPL18; RPS2; RPL13A; RPLP0; RPS16; EIF3F; RPL13; RPS13; RPS14; RPL7A; RPL5; RPL8; RPL18A; RPL4; EIF3L; EIF4B; RPL3 |
| R-HSA-2408557 | Selenocysteine synthesis | 91 | -7.66E-01 | -3.16 | 1.00E-10 | 4.55E-09 | RPS27; UBA52; RPS8; RPS27L; RPL22; RPL24; RPL34; RPL39; RPL23; RPSA; RPS29; RPL32; RPL7; RPS20; RPL29; RPL31; RPL35A; RPS21; RPL21; RPS18; RPL22L1; RPL27; RPL15; RPS12; RPS5; RPLP1; RPS27A; RPS15; RPS9; RPL27A; RPL19; RPLP2; RPL10; RPS6; RPS28; RPS3; RPL10A; RPL6; RPL38; RPS4X; RPS19; RPL18; RPS2; RPL13A; RPLP0; RPS16; RPL13; RPS13; RPS14; RPL7A; RPL5; RPL8; RPL18A; RPL4; RPL3 |
| R-HSA-192823 | Viral mRNA Translation | 87 | -7.83E-01 | -3.16 | 1.00E-10 | 4.55E-09 | RPS27; UBA52; RPS8; RPS27L; RPL22; RPL24; RPL34; RPL39; RPL23; RPSA; RPS29; RPL32; RPL7; RPS20; RPL29; RPL31; RPL35A; RPS21; RPL21; RPS18; RPL22L1; RPL27; RPL15; RPS12; RPS5; RPLP1; RPS27A; RPS15; RPS9; RPL27A; RPL19; RPLP2; RPL10; RPS6; RPS28; RPS3; RPL10A; RPL6; GRSF1; RPL38; RPS4X; RPS19; RPL18; RPS2; RPL13A; RPLP0; RPS16; RPL13; RPS13; RPS14; RPL7A; RPL5; RPL8; RPL18A; RPL4; RPL3 |
| R-HSA-975956 | Nonsense Mediated Decay (NMD) independent of the Exon Junction Complex (EJC) | 93 | -7.62E-01 | -3.16 | 1.00E-10 | 4.55E-09 | RPS27; UBA52; RPS8; RPS27L; RPL22; RPL24; RPL34; RPL39; RPL23; RPSA; RPS29; RPL32; RPL7; RPS20; RPL29; RPL31; RPL35A; RPS21; RPL21; RPS18; RPL22L1; RPL27; RPL15; RPS12; RPS5; RPLP1; RPS27A; RPS15; RPS9; RPL27A; RPL19; RPLP2; RPL10; RPS6; RPS28; RPS3; RPL10A; RPL6; RPL38; RPS4X; RPS19; RPL18; RPS2; RPL13A; RPLP0; RPS16; RPL13; RPS13; RPS14; RPL7A; RPL5; RPL8; RPL18A; RPL4; RPL3 |
| R-HSA-72764 | Eukaryotic Translation Termination | 91 | -7.67E-01 | -3.17 | 1.00E-10 | 4.55E-09 | RPS27; UBA52; RPS8; RPS27L; RPL22; RPL24; RPL34; RPL39; RPL23; RPSA; RPS29; RPL32; RPL7; RPS20; RPL29; RPL31; RPL35A; RPS21; RPL21; RPS18; RPL22L1; RPL27; RPL15; RPS12; RPS5; RPLP1; RPS27A; RPS15; RPS9; RPL27A; RPL19; RPLP2; RPL10; RPS6; RPS28; RPS3; RPL10A; RPL6; RPL38; RPS4X; RPS19; RPL18; RPS2; RPL13A; RPLP0; RPS16; RPL13; RPS13; RPS14; RPL7A; RPL5; RPL8; RPL18A; RPL4; RPL3 |
| R-HSA-156827 | L13a-mediated translational silencing of Ceruloplasmin expression | 109 | -7.51E-01 | -3.17 | 1.00E-10 | 4.55E-09 | RPS17; FAU; EIF4G1; EIF3D; EIF3A; EIF3C; RPS11; RPS3A; RPL35; EIF4A2; RPS23; EIF3M; RPL14; RPS7; RPL36; EIF3I; RPS25; EIF3K; EIF3B; RPS27; EIF3E; UBA52; RPS8; RPS27L; RPL22; RPL24; RPL34; RPL39; RPL23; RPSA; RPS29; RPL32; RPL7; RPS20; RPL29; RPL31; RPL35A; EIF4H; RPS21; RPL21; RPS18; RPL22L1; RPL27; RPL15; RPS12; RPS5; RPLP1; RPS27A; RPS15; RPS9; RPL27A; RPL19; RPLP2; RPL10; RPS6; RPS28; RPS3; RPL10A; EIF3H; RPL6; RPL38; RPS4X; RPS19; RPL18; RPS2; RPL13A; RPLP0; RPS16; EIF3F; RPL13; RPS13; RPS14; RPL7A; RPL5; RPL8; RPL18A; RPL4; EIF3L; EIF4B; RPL3 |
| R-HSA-72689 | Formation of a pool of free 40S subunits | 99 | -7.62E-01 | -3.18 | 1.00E-10 | 4.55E-09 | RPS17; FAU; EIF3D; EIF3A; EIF3C; RPS11; RPS3A; RPL35; RPS23; EIF3M; RPL14; RPS7; RPL36; EIF3I; RPS25; EIF3K; EIF3B; RPS27; EIF3E; UBA52; RPS8; RPS27L; RPL22; RPL24; RPL34; RPL39; RPL23; RPSA; RPS29; RPL32; RPL7; RPS20; RPL29; RPL31; RPL35A; RPS21; RPL21; RPS18; RPL22L1; RPL27; RPL15; RPS12; RPS5; RPLP1; RPS27A; RPS15; RPS9; RPL27A; RPL19; RPLP2; RPL10; RPS6; RPS28; RPS3; RPL10A; EIF3H; RPL6; RPL38; RPS4X; RPS19; RPL18; RPS2; RPL13A; RPLP0; RPS16; EIF3F; RPL13; RPS13; RPS14; RPL7A; RPL5; RPL8; RPL18A; RPL4; EIF3L; RPL3 |
| R-HSA-156902 | Peptide chain elongation | 87 | -7.89E-01 | -3.19 | 1.00E-10 | 4.55E-09 | RPS27; UBA52; RPS8; RPS27L; RPL22; RPL24; RPL34; RPL39; RPL23; RPSA; RPS29; RPL32; RPL7; RPS20; RPL29; RPL31; RPL35A; RPS21; RPL21; RPS18; RPL22L1; RPL27; RPL15; RPS12; RPS5; RPLP1; RPS27A; RPS15; RPS9; RPL27A; RPL19; RPLP2; RPL10; RPS6; RPS28; RPS3; RPL10A; RPL6; RPL38; RPS4X; RPS19; RPL18; RPS2; RPL13A; RPLP0; RPS16; EEF1A1; RPL13; RPS13; RPS14; RPL7A; RPL5; RPL8; RPL18A; EEF2; RPL4; RPL3 |
| R-HSA-156842 | Eukaryotic Translation Elongation | 90 | -7.91E-01 | -3.26 | 1.00E-10 | 4.55E-09 | RPS27; UBA52; RPS8; RPS27L; RPL22; RPL24; RPL34; RPL39; RPL23; RPSA; RPS29; RPL32; RPL7; EEF1B2; RPS20; RPL29; RPL31; RPL35A; RPS21; RPL21; RPS18; RPL22L1; RPL27; RPL15; RPS12; RPS5; RPLP1; RPS27A; RPS15; RPS9; RPL27A; EEF1D; RPL19; RPLP2; RPL10; RPS6; RPS28; RPS3; RPL10A; RPL6; RPL38; RPS4X; RPS19; RPL18; RPS2; RPL13A; RPLP0; RPS16; EEF1A1; RPL13; RPS13; RPS14; RPL7A; RPL5; RPL8; RPL18A; EEF2; RPL4; RPL3 |

**Supplementary Table 6 | Ex vivo cytokine production results in the 2000HIV study.**

The table reports cytokine levels from stimulated peripheral blood in the discovery (n=1,002) and validation (n=189) cohorts, comparing multi-omics endotypes (MIXED vs. ALL LOW, MIXED vs. ALL HIGH, ALL HIGH vs. ALL LOW). Data include cytokines measured after stimulation with various agents (Poly I:C, LPS, Imiquimod, IL-1α, HIV-ENV, CMV, S. pneumoniae, E. coli, S. aureus, M. tuberculosis, C. albicans conidia/hyphae, PHA) for 24 hours or 7 days. Columns include stimulus, cytokine, P-values for discovery and validation cohorts, estimates (standardized beta coefficients) for discovery and validation cohorts, endotype comparison, and stimulation time. Cytokine levels were analyzed using rank-based regression (Rfit package v0.24.6, Bent1 transformation for skewness) in R (v4.3.0), adjusting for confounders selected through a two-step process: (1) principal component analysis (PCA) on raw omics data using FactoMineR (v2.4), with the first 10 principal components regressed against potential confounders (age, sex, ethnicity [first genetic principal component], collection center, seasonality [sine/cosine functions], time to laboratory preprocessing, COVID-19 vaccination status, HIV-related variables [e.g., cART duration, CD4 nadir]); (2) confounders causing >10% change in beta coefficients for omics-clinical associations were selected (age, sex, seasonality, collection center, time to laboratory preprocessing, first genetic principal component). Significant associations required FDR-adjusted P<0.05 in the discovery cohort, nominal P<0.05 in the validation cohort, and consistent effect direction. Missing cytokine-stimulus pairs (e.g., IL-10/Poly I:C, S. aureus/IL-5) were excluded. Only validated results with consistent effect direction and significance are reported. cART, combination antiretroviral therapy; FDR, false discovery rate. See **Methods** for details.

| Stimulus | Cytokine | P-Value Discovery | P-Value Validation | Discovery Estimate | Validation Estimate | Comparison | Stimulation Time |
| --- | --- | --- | --- | --- | --- | --- | --- |
| PHA | IL5 | 0.000642496 | 0.525504573 | 0.081901865 | 0.045258497 | MIXED_vs_ALL LOW | 7d |
| C.alb.con | IL5 | 0.002130996 | 0.687253105 | 0.102162637 | 0.028166388 | MIXED_vs_ALL LOW | 7d |
| PHA | IL17 | 0.00145937 | 0.932200211 | -0.097410177 | -0.007754255 | MIXED_vs_ALL HIGH | 7d |
| S.pneu | IFNy | 0.022030367 | 0.695800954 | -0.061175737 | -0.02142275 | MIXED_vs_ALL HIGH | 7d |
| E.coli | IFNy | 0.04483173 | 0.868268105 | -0.055855 | -0.009336816 | MIXED_vs_ALL HIGH | 7d |
| PHA | IL10 | 3.50262E-05 | 0.90074307 | -0.127373861 | -0.010739419 | ALL HIGH_vs_ALL LOW | 7d |
| S.pneu | IL17 | 0.00058044 | 0.926961049 | -0.113889641 | -0.00783772 | ALL HIGH_vs_ALL LOW | 7d |
| PHA | IL5 | 0.001031579 | 0.425230875 | 0.097169927 | 0.064173736 | ALL HIGH_vs_ALL LOW | 7d |
| Spneu | MIP1a | 0.014147243 | 0.112140707 | -0.110174607 | -0.177163412 | ALL HIGH_vs_ALL LOW | 24h |
| MTB | IFNy | 0.015585815 | 0.802088815 | -0.040561734 | -0.006451547 | ALL HIGH_vs_ALL LOW | 7d |
| S.pneu | IFNy | 0.023172054 | 0.069498922 | -0.067576144 | -0.121318722 | ALL HIGH_vs_ALL LOW | 7d |
| E.coli | IFNy | 0.024835 | 0.691395286 | -0.067040305 | -0.027531166 | ALL HIGH_vs_ALL LOW | 7d |
| E.coli | IL22 | 0.025010397 | 0.789954887 | -0.057428431 | -0.013470059 | ALL HIGH_vs_ALL LOW | 7d |

**Supplementary Table 7 | Baseline demographic characteristics of the discovery and validation cohorts in the 2000HIV study.** Baseline demographic characteristics for the 2000HIV cohort (e.g. Age, Sex, Ethnicity). Data are stratified by study groups: MIXED (discovery=416, validation=67), ALL LOW (discovery=345, validation=83), and ALL HIGH (discovery=241, validation=39). Continuous variables are presented as mean (standard deviation) for normally distributed data (e.g., age) or median [interquartile range] for non-normally distributed data (e.g., CMV IgG levels). Categorical variables are reported as number (percentage). Statistical comparisons were performed using the Kruskal–Wallis test for non-normally distributed continuous variables and the chi-squared test (or Fisher’s exact test where appropriate) for categorical variables, with Bonferroni correction for multiple testing (P<0.05) using the compareGroups package in R (v4.3.0). P values are provided for overall group comparisons and pairwise comparisons (MIXED vs. ALL LOW, MIXED vs. ALL HIGH, ALL LOW vs. ALL HIGH). Hepatitis B status was defined as follows: previous infection (positive anti-HBc antibodies), vaccinated (positive anti-HBs antibodies per medical history), chronic treated (PCR-negative HBV-DNA, per inclusion criteria), or antibodies-negative (negative anti-HBV antibodies). Hepatitis C status was defined by a positive HCV test in medical history, regardless of treatment or spontaneous clearance. Ethnicity was self-reported per NIH guidelines (White, Black, Asian, Hispanic, Native American, or Mixed, with Mixed defined as two grandparents of at least two different ethnic ancestries). Sex refers to biological sex at birth. Collection centres are abbreviated as EMC (Erasmus MC, Rotterdam), OLV (OLVG, Amsterdam), RUMC (Radboud UMC, Nijmegen), and ETZ (Elizabeth-Tweesteden, Tilburg). Collection timing relative to the COVID-19 pandemic in the Netherlands was categorized as before (enrolled prior to 12 March 2020) or after (enrolled after 7 June 2020), with inclusions paused between these dates. COVID-19 history and vaccination status (one or more doses before baseline) were patient-reported. HIV controller status was categorized as follows: HIC Persistent (elite controllers without loss of control), including viremic (HIV-positive, ≥5 years without cART, HIV-RNA always <10,000 copies/ml, and stable CD4 ≥500 cells/µL for >75% of measurements) and non-viremic (HIV-positive, ≥1 year without cART, ≥3 consecutive HIV-RNA <75 copies/ml spanning ≥12 months, and stable CD4 ≥500 cells/µL for >75% of measurements) subgroups; HIC Transient (elite controllers who lost control, defined as CD4 <500 cells/mm³ or HIV-RNA >10,000 copies/ml); and Non-HIV Controller (non-controlling PLHIV on cART, reference group). Some individuals were marked NA for controller status if they had low viral load and high CD4 before cART but initiated cART before meeting HIC Persistent criteria. Immunological response was defined as responder or non-responder based on study criteria. CMV, cytomegalovirus; IgG, immunoglobulin G; IU, international units.

*Discovery Cohort*

|  | **MIXED (N=416)** | **ALL LOW (N=345)** | **ALL HIGH (N=241)** | **p.overall** | **p.MIXED vs ALL LOW** | **p.MIXED vs ALL HIGH** | **p.ALL LOW vs ALL HIGH** |
| --- | --- | --- | --- | --- | --- | --- | --- |
| Age | 52.4 (10.5) | 48.2 (13.0) | 51.8 (12.1) | <0.001 | <0.001 | 0.830 | 0.001 |
| Sex: |  |  |  | 0.616 | 0.715 | 0.715 | 0.715 |
| Male | 348 (83.7%) | 283 (82.0%) | 205 (85.1%) |  |  |  |  |
| Female | 68 (16.3%) | 62 (18.0%) | 36 (14.9%) |  |  |  |  |
| Ethnicity: |  |  |  | 0.756 | 0.839 | 0.839 | 0.839 |
| Asian | 26 (6.25%) | 15 (4.35%) | 13 (5.39%) |  |  |  |  |
| Black | 50 (12.0%) | 44 (12.8%) | 25 (10.4%) |  |  |  |  |
| Hispanic | 15 (3.61%) | 16 (4.64%) | 6 (2.49%) |  |  |  |  |
| Mixed | 33 (7.93%) | 33 (9.57%) | 19 (7.88%) |  |  |  |  |
| Native American | 0 (0.00%) | 0 (0.00%) | 0 (0.00%) |  |  |  |  |
| White | 292 (70.2%) | 237 (68.7%) | 178 (73.9%) |  |  |  |  |
| Currently Smoking: |  |  |  | 0.501 | 0.633 | 0.795 | 0.633 |
| Non-Smoking | 254 (66.5%) | 200 (63.3%) | 148 (67.9%) |  |  |  |  |
| Smoking | 128 (33.5%) | 116 (36.7%) | 70 (32.1%) |  |  |  |  |
| Collection Center: |  |  |  | 0.014 | 0.907 | 0.010 | 0.010 |
| EMC | 147 (35.3%) | 127 (36.8%) | 88 (36.5%) |  |  |  |  |
| OLV | 196 (47.1%) | 160 (46.4%) | 133 (55.2%) |  |  |  |  |
| ETZ | 0 (0.00%) | 0 (0.00%) | 0 (0.00%) |  |  |  |  |
| RUMC | 73 (17.5%) | 58 (16.8%) | 20 (8.30%) |  |  |  |  |
| Season of Baseline Visit: |  |  |  | 0.119 | 0.312 | 0.196 | 0.262 |
| Winter | 64 (15.4%) | 45 (13.0%) | 20 (8.30%) |  |  |  |  |
| Spring | 68 (16.3%) | 54 (15.7%) | 46 (19.1%) |  |  |  |  |
| Summer | 162 (38.9%) | 157 (45.5%) | 103 (42.7%) |  |  |  |  |
| Autumn | 122 (29.3%) | 89 (25.8%) | 72 (29.9%) |  |  |  |  |
| CMV IgG (IupermL) | 684 [400;  900] | 548 [323;  809] | 693 [413;  913] | 0.001 | 0.002 | 0.610 | 0.002 |
| CMV IgG Serology: |  |  |  | 0.147 | 0.262 | 1.000 | 0.262 |
| Negative | 23 (5.54%) | 30 (8.72%) | 13 (5.39%) |  |  |  |  |
| Positive | 392 (94.5%) | 314 (91.3%) | 228 (94.6%) |  |  |  |  |
| Hepatitis B Status: |  |  |  | 0.143 | 0.194 | 0.519 | 0.194 |
| Antibodies Negative | 59 (14.6%) | 58 (17.3%) | 40 (17.3%) |  |  |  |  |
| Chronic Treated | 11 (2.72%) | 7 (2.08%) | 10 (4.33%) |  |  |  |  |
| Previous Infection | 152 (37.5%) | 100 (29.8%) | 84 (36.4%) |  |  |  |  |
| Vaccinated | 183 (45.2%) | 171 (50.9%) | 97 (42.0%) |  |  |  |  |
| Hepatitis C | 44 (10.6%) | 35 (10.1%) | 19 (7.88%) | 0.513 | 0.940 | 0.648 | 0.648 |
| Treated for Hepatitis C | 37 (86.0%) | 34 (97.1%) | 15 (78.9%) | 0.055 | 0.183 | 0.479 | 0.140 |
| Collection During Pandemic |  |  |  | 1.000 | 1.000 | 1.000 | 1.000 |
| Before | 4 (0.96%) | 4 (1.16%) | 2 (0.83%) |  |  |  |  |
| After | 412 (99.0%) | 341 (98.8%) | 239 (99.2%) |  |  |  |  |
| COVID19 |  |  |  | 0.525 | 0.620 | 1.000 | 0.620 |
| No | 358 (86.1%) | 288 (83.5%) | 208 (86.3%) |  |  |  |  |
| Yes | 58 (13.9%) | 57 (16.5%) | 33 (13.7%) |  |  |  |  |
| Received COVID19 Vaccination |  |  |  | 0.204 | 0.228 | 0.879 | 0.228 |
| No | 288 (69.2%) | 221 (64.1%) | 169 (70.1%) |  |  |  |  |
| Yes | 128 (30.8%) | 124 (35.9%) | 72 (29.9%) |  |  |  |  |
| HIV Controller Status: |  |  |  | <0.001 | <0.001 | 0.707 | <0.001 |
| HIC Persistent | 3 (0.72%) | 36 (10.9%) | 2 (0.83%) |  |  |  |  |
| HIC Transiend | 13 (3.14%) | 18 (5.47%) | 5 (2.07%) |  |  |  |  |
| Non-HIV Controller | 398 (96.1%) | 275 (83.6%) | 234 (97.1%) |  |  |  |  |
| Immunological Response: |  |  |  | 0.067 | 0.370 | 0.353 | 0.107 |
| Responder | 330 (95.4%) | 295 (97.0%) | 172 (92.5%) |  |  |  |  |
| Non-Responder | 16 (4.62%) | 9 (2.96%) | 14 (7.53%) |  |  |  |  |

*Validation Cohort*

|  | **MIXED (N=67)** | **ALL LOW (N=83)** | **ALL HIGH (N=39)** | **p.overall** | **p.MIXED vs ALL LOW** | **p.MIXED vs ALL HIGH** | **p.ALL LOW vs ALL HIGH** |
| --- | --- | --- | --- | --- | --- | --- | --- |
| Age | 55.0 (9.89) | 52.0 (12.2) | 55.8 (9.66) | 0.112 | 0.208 | 0.938 | 0.175 |
| Sex: |  |  |  | 0.206 | 0.850 | 0.365 | 0.365 |
| Male | 55 (82.1%) | 66 (79.5%) | 36 (92.3%) |  |  |  |  |
| Female | 12 (17.9%) | 17 (20.5%) | 3 (7.69%) |  |  |  |  |
| Ethnicity: |  |  |  | 0.328 | 0.441 | 0.573 | 0.441 |
| Asian | 0 (0.00%) | 3 (3.61%) | 1 (2.56%) |  |  |  |  |
| Black | 6 (8.96%) | 6 (7.23%) | 3 (7.69%) |  |  |  |  |
| Hispanic | 0 (0.00%) | 0 (0.00%) | 0 (0.00%) |  |  |  |  |
| Mixed | 1 (1.49%) | 0 (0.00%) | 2 (5.13%) |  |  |  |  |
| Native American | 1 (1.49%) | 0 (0.00%) | 0 (0.00%) |  |  |  |  |
| White | 59 (88.1%) | 74 (89.2%) | 33 (84.6%) |  |  |  |  |
| Currently Smoking: |  |  |  | 0.405 | 0.562 | 0.971 | 0.562 |
| Non-Smoking | 38 (63.3%) | 54 (72.0%) | 20 (60.6%) |  |  |  |  |
| Smoking | 22 (36.7%) | 21 (28.0%) | 13 (39.4%) |  |  |  |  |
| Collection Center: |  |  |  | . | . | . | . |
| EMC | 0 (0.00%) | 0 (0.00%) | 0 (0.00%) |  |  |  |  |
| OLV | 0 (0.00%) | 0 (0.00%) | 0 (0.00%) |  |  |  |  |
| ETZ | 67 (100%) | 83 (100%) | 39 (100%) |  |  |  |  |
| RUMC | 0 (0.00%) | 0 (0.00%) | 0 (0.00%) |  |  |  |  |
| Season of Baseline Visit: |  |  |  | 0.611 | 0.640 | 0.640 | 0.800 |
| Winter | 4 (5.97%) | 10 (12.0%) | 3 (7.69%) |  |  |  |  |
| Spring | 21 (31.3%) | 27 (32.5%) | 15 (38.5%) |  |  |  |  |
| Summer | 29 (43.3%) | 36 (43.4%) | 18 (46.2%) |  |  |  |  |
| Autumn | 13 (19.4%) | 10 (12.0%) | 3 (7.69%) |  |  |  |  |
| CMV IgG (IupermL) | 700 [471;  910] | 719 [411;  946] | 734 [438;  891] | 0.858 | 0.972 | 0.972 | 0.972 |
| CMV IgG Serology: |  |  |  | 0.383 | 0.500 | 1.000 | 0.500 |
| Negative | 4 (5.97%) | 10 (12.2%) | 2 (5.13%) |  |  |  |  |
| Positive | 63 (94.0%) | 72 (87.8%) | 37 (94.9%) |  |  |  |  |
| Hepatitis B Status: |  |  |  | 0.016 | 0.007 | 0.158 | 0.718 |
| Antibodies Negative | 2 (4.35%) | 13 (19.4%) | 5 (18.5%) |  |  |  |  |
| Chronic Treated | 1 (2.17%) | 2 (2.99%) | 1 (3.70%) |  |  |  |  |
| Previous Infection | 27 (58.7%) | 18 (26.9%) | 10 (37.0%) |  |  |  |  |
| Vaccinated | 16 (34.8%) | 34 (50.7%) | 11 (40.7%) |  |  |  |  |
| Hepatitis C | 8 (11.9%) | 4 (4.82%) | 3 (7.69%) | 0.263 | 0.585 | 0.743 | 0.743 |
| Treated for Hepatitis C | 6 (85.7%) | 4 (100%) | 3 (100%) | 1.000 | 1.000 | 1.000 | . |
| Collection During Pandemic |  |  |  | . | . | . | . |
| Before | 0 (0.00%) | 0 (0.00%) | 0 (0.00%) |  |  |  |  |
| After | 67 (100%) | 83 (100%) | 39 (100%) |  |  |  |  |
| COVID19 |  |  |  | 0.235 | 0.612 | 0.612 | 0.455 |
| No | 54 (80.6%) | 72 (86.7%) | 29 (74.4%) |  |  |  |  |
| Yes | 13 (19.4%) | 11 (13.3%) | 10 (25.6%) |  |  |  |  |
| Received COVID19 Vaccination |  |  |  | 0.209 | 0.861 | 0.315 | 0.315 |
| No | 47 (70.1%) | 56 (67.5%) | 21 (53.8%) |  |  |  |  |
| Yes | 20 (29.9%) | 27 (32.5%) | 18 (46.2%) |  |  |  |  |
| HIV Controller Status: |  |  |  | 0.057 | 0.195 | 1.000 | 0.195 |
| HIC Persistent | 0 (0.00%) | 1 (1.25%) | 0 (0.00%) |  |  |  |  |
| HIC Transient | 1 (1.49%) | 7 (8.75%) | 0 (0.00%) |  |  |  |  |
| Non-HIV Controller | 66 (98.5%) | 72 (90.0%) | 38 (100%) |  |  |  |  |
| Immunological Response: |  |  |  | 0.003 | 0.007 | 1.000 | 0.019 |
| Responder | 44 (83.0%) | 69 (98.6%) | 28 (84.8%) |  |  |  |  |
| Non-Responder | 9 (17.0%) | 1 (1.43%) | 5 (15.2%) |  |  |  |  |

**Supplementary Table 8 | Baseline clinical characteristics of the discovery and validation cohorts in the 2000HIV study.**

Baseline demographic characteristics for the 2000HIV cohort (e.g. Age, Sex, Ethnicity). Data are stratified by study groups: MIXED (discovery=416, validation=67), ALL LOW (discovery=345, validation=83), and ALL HIGH (discovery=241, validation=39). Continuous variables are presented as mean (standard deviation) for normally distributed data (e.g., age) or median [interquartile range] for non-normally distributed data (e.g., CMV IgG levels). Categorical variables are reported as number (percentage). Statistical comparisons were performed using the Kruskal–Wallis test for non-normally distributed continuous variables and the chi-squared test (or Fisher’s exact test where appropriate) for categorical variables, with Bonferroni correction for multiple testing (P<0.05) using the compareGroups package in R (v4.3.0). P values are provided for overall group comparisons and pairwise comparisons (MIXED vs. ALL LOW, MIXED vs. ALL HIGH, ALL LOW vs. ALL HIGH). HIV duration and cART duration reflect years since HIV diagnosis and initiation of combination antiretroviral therapy (cART), respectively. CD4 nadir represents the lowest CD4 T cell count ever recorded (10⁹ cells/L), excluding isolated low measurements due to sample issues or concurrent infections. Viral load zenith indicates the highest recorded HIV-RNA measurement (copies/ml), including values from acute/recent HIV infection, with detection limits recorded if applicable. Latest CD4 and CD8 counts (10⁹ cells/L) and CD4/CD8 ratio were measured at baseline. Latest viral load was quantifiable (>40 copies/ml) or undetectable (unmeasurable or <40 copies/ml, considered equivalent in clinical settings). Intact and total HIV DNA copies per million CD4 T cells were measured, with logarithmic transformations applied for analysis. All measurements were extracted from baseline clinical records (NCT03994835). cART, combination antiretroviral therapy.

*Discovery Cohort*

|  | **Mixed (N=416)** | **All Low (N=345)** | **All High (N=241)** | **p.overall** | **p.MIXED vs ALL LOW** | **p.MIXED vs ALL HIGH** | **p.ALL LOW vs ALL HIGH** |
| --- | --- | --- | --- | --- | --- | --- | --- |
| BMI | 25.5 (4.34) | 25.5 (4.43) | 25.2 (4.11) | 0.513 | 0.987 | 0.509 | 0.618 |
| HIV Duration (Years) | 15.8 (8.15) | 12.8 (7.90) | 14.0 (8.33) | <0.001 | <0.001 | 0.019 | 0.185 |
| cART Duration (Years) | 12.9 (6.53) | 10.4 (6.70) | 11.6 (6.73) | <0.001 | <0.001 | 0.040 | 0.101 |
| CD4 Nadir | 0.22 [0.12;0.31] | 0.37 [0.25;0.55] | 0.19 [0.08;0.30] | <0.001 | <0.001 | 0.028 | <0.001 |
| Viral Load Zenith | 112601 [57500;  315900] | 45900 [11500;  114000] | 153500 [71860;  374449] | <0.001 | <0.001 | 0.030 | <0.001 |
| Viral Load Zenith (log) | 17.0 (2.18) | 15.1 (3.40) | 17.4 (2.33) | <0.001 | 0.000 | 0.267 | 0.000 |
| Latest CD4/CD8 | 0.92 (0.49) | 1.08 (0.51) | 0.83 (0.41) | <0.001 | <0.001 | 0.065 | <0.001 |
| Latest CD4 | 0.74 (0.28) | 0.84 (0.34) | 0.69 (0.28) | <0.001 | <0.001 | 0.171 | <0.001 |
| Latest CD8 | 0.95 (0.45) | 0.90 (0.45) | 0.97 (0.49) | 0.142 | 0.335 | 0.769 | 0.142 |
| Latest Viral Load (log) | 0.03 (0.17) | 0.02 (0.15) | 0.05 (0.22) | 0.201 | 0.809 | 0.405 | 0.178 |
| Intact HIV DNA Copies/Million CD4 T-cells | 20.0 [6.00;37.0] | 4.00 [3.00;9.00] | 144 [91.5;244] | <0.001 | <0.001 | <0.001 | <0.001 |
| Total HIV DNA Copies/Million CD4 T-cells | 840 [587;1264] | 165 [79.0;264] | 1492 [890;2251] | <0.001 | <0.001 | <0.001 | <0.001 |
| Intact HIV DNA Copies/Million CD4 T-cells (log) | 4.20 (1.28) | 2.94 (0.89) | 7.34 (1.07) | <0.001 | 0.000 | 0.000 | 0.000 |
| Total HIV DNA Copies/Million CD4 T-cells (log) | 9.76 (0.88) | 7.09 (1.26) | 10.5 (1.19) | <0.001 | 0.000 | 0.000 | 0.000 |

*Validation Cohort*

|  | **Mixed (N=67)** | **All Low (N=83)** | **All High (N=39)** | **p.overall** | **p.MIXED vs ALL LOW** | **p.MIXED vs ALL HIGH** | **p.ALL LOW vs ALL HIGH** |
| --- | --- | --- | --- | --- | --- | --- | --- |
| BMI | 26.1 (4.28) | 26.0 (4.87) | 26.3 (3.94) | 0.934 | 0.978 | 0.981 | 0.930 |
| HIV Duration (Years) | 14.6 (7.59) | 13.3 (7.88) | 11.6 (8.70) | 0.176 | 0.581 | 0.152 | 0.519 |
| cART Duration (Years) | 12.7 (6.69) | 11.1 (7.20) | 9.53 (6.53) | 0.075 | 0.346 | 0.065 | 0.478 |
| CD4 Nadir | 0.22 [0.12;0.30] | 0.30 [0.22;0.43] | 0.17 [0.08;0.35] | 0.001 | 0.002 | 0.729 | 0.018 |
| Viral Load Zenith | 200000 [94531;  369000] | 102000 [28536;  266582] | 200000 [120000;  585322] | 0.003 | 0.015 | 0.339 | 0.009 |
| Viral Load Zenith (log) | 17.5 (1.71) | 16.6 (2.63) | 17.9 (1.68) | 0.010 | 0.067 | 0.718 | 0.017 |
| Latest CD4/CD8 | . | . | . | . | . | . | . |
| Latest CD4 | 0.67 (0.28) | 0.74 (0.28) | 0.67 (0.28) | 0.183 | 0.227 | 1.000 | 0.343 |
| Latest CD8 | . | . | . | . | . | . | . |
| Latest Viral Load (log) | 0.06 (0.24) | 0.02 (0.15) | 0.03 (0.16) | 0.477 | 0.489 | 0.646 | 0.999 |
| Intact HIV DNA Copies/Million CD4 T-cells | 26.0 [7.00;49.0] | 7.00 [4.00;14.8] | 168 [122;245] | <0.001 | <0.001 | <0.001 | <0.001 |
| Total HIV DNA Copies/Million CD4 T-cells | 1039 [762;1444] | 283 [173;410] | 1677 [1018;2267] | <0.001 | <0.001 | 0.004 | <0.001 |
| Intact HIV DNA Copies/Million CD4 T-cells (log) | 4.53 (1.36) | 3.35 (1.08) | 7.52 (0.91) | <0.001 | <0.001 | <0.001 | <0.001 |
| Total HIV DNA Copies/Million CD4 T-cells (log) | 10.1 (0.77) | 7.93 (1.05) | 10.6 (0.85) | <0.001 | <0.001 | 0.023 | <0.001 |

**Supplementary Table 9 | Antiretroviral therapy (cART) characteristics of the discovery cohort in the 2000HIV study.** Baseline demographic characteristics for the 2000HIV cohort (e.g. Age, Sex, Ethnicity). Data are stratified by study groups: MIXED (discovery=416, validation=67), ALL LOW (discovery=345, validation=83), and ALL HIGH (discovery=241, validation=39). Continuous variables are presented as mean (standard deviation) for normally distributed data (e.g., age) or median [interquartile range] for non-normally distributed data (e.g., CMV IgG levels). Categorical variables are reported as number (percentage). Statistical comparisons were performed using the Kruskal–Wallis test for non-normally distributed continuous variables and the chi-squared test (or Fisher’s exact test where appropriate) for categorical variables, with Bonferroni correction for multiple testing (P<0.05) using the compareGroups package in R (v4.3.0). P values are provided for overall group comparisons and pairwise comparisons (MIXED vs. ALL LOW, MIXED vs. ALL HIGH, ALL LOW vs. ALL HIGH). Categorical variables indicate current use (yes/no) of cART drug classes at baseline: nucleoside reverse transcriptase inhibitors (NRTIs; e.g., abacavir, lamivudine, emtricitabine, excluding tenofovir), nucleotide reverse transcriptase inhibitors (NtRTIs; e.g., tenofovir alafenamide, tenofovir disoproxil), non-nucleoside reverse transcriptase inhibitors (NNRTIs; e.g., efavirenz), protease inhibitors (e.g., darunavir), CCR5 inhibitors (e.g., maraviroc), and integrase inhibitors (e.g., dolutegravir). cART interruption was defined as a treatment break of >3 consecutive months after initiation. Early cART was defined as initiation within 1 month of confirmed acute HIV infection (confirmed by a negative HIV test <6 months prior or incomplete immunoblot at diagnosis). Cumulative exposure to specific drugs (3TC, lamivudine; ABC, abacavir; DTG, dolutegravir; EFV, efavirenz; FTC, emtricitabine; RTV, ritonavir; TAF, tenofovir alafenamide; TDF, tenofovir disoproxil) and overall ART exposure were calculated as total duration in years prior to study inclusion, summed across regimens per drug, with the inclusion date as the end date for ongoing regimens. Only drugs with at least 1 month of cumulative exposure were included. Data were extracted from baseline clinical records (NCT03994835). cART, combination antiretroviral therapy.

*Discovery Cohort*

|  | **MIXED (N=416)** | **ALL LOW (N=345)** | **ALL HIGH (N=241)** | **P overall** | **p.MIXED vs ALL LOW** | **p.MIXED vs ALL HIGH** | **p.ALL LOW vs ALL HIGH** |
| --- | --- | --- | --- | --- | --- | --- | --- |
| CART NRTIs: |  |  |  | 0.141 | 0.305 | 0.641 | 0.305 |
| No | 22 (5.29%) | 27 (7.83%) | 10 (4.15%) |  |  |  |  |
| Yes | 394 (94.7%) | 318 (92.2%) | 231 (95.9%) |  |  |  |  |
| CART NtRTIs: |  |  |  | 0.621 | 0.791 | 0.816 | 0.791 |
| No | 131 (31.5%) | 117 (33.9%) | 73 (30.3%) |  |  |  |  |
| Yes | 285 (68.5%) | 228 (66.1%) | 168 (69.7%) |  |  |  |  |
| CART NNRTIs: |  |  |  | 0.160 | 0.221 | 0.793 | 0.344 |
| No | 229 (55.0%) | 213 (61.7%) | 136 (56.4%) |  |  |  |  |
| Yes | 187 (45.0%) | 132 (38.3%) | 105 (43.6%) |  |  |  |  |
| CART Protease inhibitors: |  |  |  | 0.038 | 0.047 | 0.327 | 0.327 |
| No | 364 (87.5%) | 321 (93.0%) | 218 (90.5%) |  |  |  |  |
| Yes | 52 (12.5%) | 24 (6.96%) | 23 (9.54%) |  |  |  |  |
| CART CCR5 inhibitor: |  |  |  | . | . | . | . |
| No | 416 (100%) | 345 (100%) | 241 (100%) |  |  |  |  |
| Yes | 0 (0.00%) | 0 (0.00%) | 0 (0.00%) |  |  |  |  |
| CART Integrase inhibitors: |  |  |  | 0.626 | 0.821 | 0.821 | 0.821 |
| No | 211 (50.7%) | 163 (47.2%) | 117 (48.5%) |  |  |  |  |
| Yes | 205 (49.3%) | 182 (52.8%) | 124 (51.5%) |  |  |  |  |
| CART Interrupted: |  |  |  | 0.268 | 0.316 | 0.911 | 0.316 |
| No | 380 (91.3%) | 315 (94.0%) | 216 (90.8%) |  |  |  |  |
| Yes | 36 (8.65%) | 20 (5.97%) | 22 (9.24%) |  |  |  |  |
| Early cART: |  |  |  | <0.001 | 0.002 | 0.913 | 0.028 |
| No | 302 (97.1%) | 250 (89.9%) | 164 (96.5%) |  |  |  |  |
| Yes | 9 (2.89%) | 28 (10.1%) | 6 (3.53%) |  |  |  |  |
| No ART exposure | 3.28 (4.61) | 3.65 (4.95) | 3.22 (4.65) | 0.611 | 0.668 | 0.992 | 0.684 |
| 3TC exposure | 7.20 (6.22) | 6.13 (5.69) | 5.96 (5.39) | 0.044 | 0.108 | 0.079 | 0.953 |
| ABC exposure | 6.05 (4.73) | 5.30 (4.68) | 4.98 (4.49) | 0.169 | 0.373 | 0.180 | 0.864 |
| DTG exposure | 3.29 (2.06) | 3.08 (1.91) | 2.76 (2.11) | 0.096 | 0.609 | 0.077 | 0.386 |
| EFV exposure | 5.52 (4.67) | 5.85 (4.64) | 6.11 (5.17) | 0.633 | 0.853 | 0.621 | 0.931 |
| FTC exposure | 8.07 (4.14) | 6.77 (4.00) | 7.60 (4.23) | 0.001 | <0.001 | 0.402 | 0.075 |
| RTV exposure | 6.63 (5.70) | 4.73 (4.31) | 5.74 (5.39) | 0.016 | 0.012 | 0.423 | 0.397 |
| TAF exposure | 2.94 (1.33) | 2.90 (1.25) | 2.86 (1.28) | 0.866 | 0.955 | 0.857 | 0.967 |
| TDF exposure | 7.78 (4.38) | 6.43 (4.60) | 7.49 (4.51) | 0.001 | 0.001 | 0.749 | 0.033 |

*Validation Cohort*

|  | **MIXED (N=67)** | **ALL LOW (N=83)** | **ALL HIGH (N=39)** | **P overall** | **p.MIXED vs ALL LOW** | **p.MIXED vs ALL HIGH** | **p.ALL LOW vs ALL HIGH** |
| --- | --- | --- | --- | --- | --- | --- | --- |
| CART NRTIs: |  |  |  | 1.000 | 1.000 | 1.000 | 1.000 |
| No | 3 (4.48%) | 4 (4.82%) | 1 (2.56%) |  |  |  |  |
| Yes | 64 (95.5%) | 79 (95.2%) | 38 (97.4%) |  |  |  |  |
| CART NtRTIs: |  |  |  | 0.187 | 1.000 | 0.227 | 0.227 |
| No | 20 (29.9%) | 25 (30.1%) | 6 (15.4%) |  |  |  |  |
| Yes | 47 (70.1%) | 58 (69.9%) | 33 (84.6%) |  |  |  |  |
| CART NNRTIs: |  |  |  | 0.239 | 0.477 | 1.000 | 0.482 |
| No | 42 (62.7%) | 62 (74.7%) | 25 (64.1%) |  |  |  |  |
| Yes | 25 (37.3%) | 21 (25.3%) | 14 (35.9%) |  |  |  |  |
| CART Protease inhibitors: |  |  |  | 0.219 | 0.316 | 0.316 | 1.000 |
| No | 57 (85.1%) | 77 (92.8%) | 37 (94.9%) |  |  |  |  |
| Yes | 10 (14.9%) | 6 (7.23%) | 2 (5.13%) |  |  |  |  |
| CART CCR5 inhibitor: |  |  |  | 1.000 | 1.000 | . | 1.000 |
| No | 67 (100%) | 82 (98.8%) | 39 (100%) |  |  |  |  |
| Yes | 0 (0.00%) | 1 (1.20%) | 0 (0.00%) |  |  |  |  |
| CART Integrase inhibitors: |  |  |  | 0.066 | 0.095 | 0.588 | 0.482 |
| No | 29 (43.3%) | 21 (25.3%) | 14 (35.9%) |  |  |  |  |
| Yes | 38 (56.7%) | 62 (74.7%) | 25 (64.1%) |  |  |  |  |
| CART Interrupted: |  |  |  | 0.728 | 1.000 | 0.792 | 0.792 |
| No | 61 (91.0%) | 76 (91.6%) | 34 (87.2%) |  |  |  |  |
| Yes | 6 (8.96%) | 7 (8.43%) | 5 (12.8%) |  |  |  |  |
| Early cART: |  |  |  | 1.000 | 1.000 | 1.000 | 1.000 |
| No | 35 (89.7%) | 43 (91.5%) | 17 (89.5%) |  |  |  |  |
| Yes | 4 (10.3%) | 4 (8.51%) | 2 (10.5%) |  |  |  |  |
| No ART exposure | 1.15 (1.27) | 3.44 (4.03) | 3.21 (4.31) | 0.197 | 0.186 | 0.382 | 0.985 |
| 3TC exposure | 8.75 (6.70) | 8.54 (6.78) | 4.55 (4.19) | 0.039 | 0.988 | 0.052 | 0.051 |
| ABC exposure | 6.58 (5.61) | 6.62 (5.33) | 3.86 (3.11) | 0.280 | 1.000 | 0.325 | 0.284 |
| DTG exposure | 3.85 (2.06) | 3.59 (1.95) | 3.35 (1.80) | 0.758 | 0.887 | 0.751 | 0.928 |
| EFV exposure | 6.85 (4.93) | 6.72 (5.34) | 6.78 (6.13) | 0.996 | 0.996 | 0.999 | 1.000 |
| FTC exposure | 8.10 (3.81) | 6.79 (4.02) | 7.09 (4.37) | 0.185 | 0.173 | 0.464 | 0.935 |
| RTV exposure | 6.46 (5.88) | 5.01 (5.60) | 4.59 (4.63) | 0.473 | 0.601 | 0.535 | 0.972 |
| TAF exposure | 2.51 (1.02) | 2.62 (1.09) | 2.50 (1.12) | 0.869 | 0.893 | 1.000 | 0.911 |
| TDF exposure | 7.98 (3.95) | 6.72 (4.22) | 6.96 (4.16) | 0.245 | 0.239 | 0.502 | 0.961 |

**Supplementary Table 10 | Association of immune–reservoir endotypes with clinical comorbidities (odds ratios).**

Unadjusted and adjusted odds ratios (ORs) with 95% confidence intervals are shown for logistic regression models assessing associations between endotype membership and clinical comorbidities across all three pairwise comparisons (All Low vs. Mixed, Mixed vs. All High, All Low vs. All High). Analyses were performed in the pooled cohort (n = 1,191), with binary outcomes encoded as “Yes/No.” Only complete cases were included for each model. Covariates were selected per outcome using a data-driven change-in-estimate strategy, retaining variables that altered the OR by ≥10% in any exposure group; the covariates included in each final adjusted model are listed in the rightmost column. Right-skewed continuous variables were log-transformed, multicollinearity was addressed by removing highly correlated variables (Pearson r>0.5), and final models demonstrated acceptable collinearity (VIF<5). Rare outcomes (e.g., hyperthyroidism) were excluded due to model instability. Comorbidities were assessed from patient medical records (NCT03994835) and include cardiovascular disorders (e.g., hypertension, myocardial infarction, stroke, angina pectoris), endocrine-metabolic disorders (e.g., hypercholesterolemia, hypertriglyceridemia, hyperthyroidism), gastroenterological diseases (e.g., liver steatosis, graded S0 [no steatosis] to S1 or higher [CAP >=263 dB/m] via fibroscan), pulmonary diseases (e.g., COPD, asthma), central nervous system disorders (e.g., epilepsy, multiple sclerosis), psychiatric disorders (e.g., depression, anxiety), musculoskeletal disorders (e.g., osteoporosis, osteoarthritis), and non-AIDS-defining cancers (e.g., prostate, breast, melanoma). Carotid plaque was defined as focal intima-media thickness (IMT) >1.5 mm or 1.5× mean IMT, with bilateral plaque indicating presence in both carotid arteries. Mean IMT was calculated as the average of three measurements per carotid artery (mm). Liver fibrosis was graded as F0–F1 (LSM <7 kPa) or F2 or higher (LSM >= 7 kPa) via fibroscan. LDL and HDL cholesterol levels were measured in mmol/L. Residual viremia (RV) consisted of viral load <40 copies/mL but detectable. Data were complemented by diagnostic subcategories (e.g., specific cancer types, causes of liver cirrhosis) when available from medical records. AIDS-defining malignancies and opportunistic infections Includes the occurrence of both before and after ART. AIDS-defining ‘other events” include infections such as HIV encephalopathy, PML, and wasting syndrome. CAP, controlled attenuation parameter; IMT, intima-media thickness; LSM, liver stiffness measurement.

|  |  |  |  |  |  | Unadjusted*^1^* | | | Adjusted | | | |
| --- | --- | --- | --- | --- | --- | --- | --- | --- | --- | --- | --- | --- |
|  | Comparison | N (Ref) | Cases (Ref) | N (Exp) | Cases (Exp) | OR (95% CI) unadj. | P unadj | P* unadj | OR (95% CI) adj. | P adj | P* adj | Confounders Adjusted For |
| Cardiovascular Diseases | | | | | | | | | | | | |
| Cardiovascular Disorder (All) | All Low vs Mixed | 428 | 119 | 483 | 161 | 1.30 (0.98–1.73) | 7.13e-02 | **†** | 0.87 (0.62–1.23) | 4.31e-01 |  | AGE + HIV.DURATION + CMV.IgG.IUpermL + SMOKING.CURRENT |
| Cardiovascular Disorder (All) | Mixed vs All High | 483 | 161 | 280 | 101 | 1.13 (0.83–1.54) | 4.43e-01 |  | 0.98 (0.68–1.42) | 9.20e-01 |  | AGE + HIV.DURATION + CMV.IgG.IUpermL + SMOKING.CURRENT |
| Cardiovascular Disorder (All) | All Low vs All High | 428 | 119 | 280 | 101 | 1.47 (1.06–2.02) | 2.04e-02 | ***** | 0.86 (0.58–1.28) | 4.56e-01 |  | AGE + HIV.DURATION + CMV.IgG.IUpermL + SMOKING.CURRENT |
| Stroke | All Low vs Mixed | 428 | 11 | 483 | 13 | 1.05 (0.46–2.41) | 9.09e-01 |  | 0.94 (0.40–2.26) | 8.84e-01 |  | AGE + HIV.DURATION + SMOKING.CURRENT |
| Stroke | Mixed vs All High | 483 | 13 | 280 | 16 | 2.19 (1.04–4.70) | 3.96e-02 | ***** | 1.90 (0.85–4.24) | 1.16e-01 |  | AGE + HIV.DURATION + SMOKING.CURRENT |
| Stroke | All Low vs All High | 428 | 11 | 280 | 16 | 2.30 (1.06–5.16) | 3.73e-02 | ***** | 1.74 (0.74–4.21) | 2.07e-01 |  | AGE + HIV.DURATION + SMOKING.CURRENT |
| Angina Pectoris | All Low vs Mixed | 428 | 7 | 483 | 14 | 1.80 (0.74–4.78) | 2.11e-01 |  | 2.04 (0.75–6.48) | 1.85e-01 |  | AGE + HIV.DURATION + CENTER + SMOKING.CURRENT |
| Angina Pectoris | Mixed vs All High | 483 | 14 | 280 | 13 | 1.63 (0.75–3.54) | 2.13e-01 |  | 1.12 (0.46–2.63) | 7.98e-01 |  | AGE + HIV.DURATION + CENTER + SMOKING.CURRENT |
| Angina Pectoris | All Low vs All High | 428 | 7 | 280 | 13 | 2.93 (1.18–7.88) | 2.38e-02 | ***** | 2.44 (0.83–8.14) | 1.18e-01 |  | AGE + HIV.DURATION + CENTER + SMOKING.CURRENT |
| Plaque | All Low vs Mixed | 428 | 187 | 483 | 274 | 1.69 (1.30–2.20) | 8.97e-05 | ******* | 1.41 (1.05–1.91) | 2.35e-02 | ***** | AGE + SMOKING.CURRENT |
| Plaque | Mixed vs All High | 483 | 274 | 280 | 159 | 1.00 (0.74–1.35) | 9.88e-01 |  | 1.08 (0.77–1.51) | 6.77e-01 |  | AGE + SMOKING.CURRENT |
| Plaque | All Low vs All High | 428 | 187 | 280 | 159 | 1.69 (1.25–2.30) | 6.81e-04 | ******* | 1.51 (1.06–2.16) | 2.21e-02 | ***** | AGE + SMOKING.CURRENT |
| Malignancies & AIDS Events | | | | | | | | | | | | |
| AIDS-Defining Malignancies (All) | All Low vs Mixed | 428 | 18 | 483 | 41 | 2.11 (1.21–3.82) | 1.01e-02 | ***** | 2.03 (1.16–3.69) | 1.64e-02 | ***** | HIV.DURATION |
| AIDS-Defining Malignancies (All) | Mixed vs All High | 483 | 41 | 280 | 23 | 0.96 (0.56–1.63) | 8.95e-01 |  | 0.95 (0.55–1.62) | 8.59e-01 |  | HIV.DURATION |
| AIDS-Defining Malignancies (All) | All Low vs All High | 428 | 18 | 280 | 23 | 2.04 (1.08–3.90) | 2.82e-02 | ***** | 1.99 (1.06–3.82) | 3.39e-02 | ***** | HIV.DURATION |
| Post-cART Malignancies | All Low vs Mixed | 428 | 21 | 483 | 37 | 1.61 (0.93–2.84) | 9.18e-02 | **†** | 1.25 (0.71–2.26) | 4.46e-01 |  | AGE + HIV.DURATION + CMV.IgG.IUpermL |
| Post-cART Malignancies | Mixed vs All High | 483 | 37 | 280 | 19 | 0.88 (0.49–1.54) | 6.55e-01 |  | 0.89 (0.48–1.59) | 6.94e-01 |  | AGE + HIV.DURATION + CMV.IgG.IUpermL |
| Post-cART Malignancies | All Low vs All High | 428 | 21 | 280 | 19 | 1.41 (0.74–2.68) | 2.92e-01 |  | 1.14 (0.59–2.19) | 6.99e-01 |  | AGE + HIV.DURATION + CMV.IgG.IUpermL |
| AIDS-Defining Opportunistic Infections (All) | All Low vs Mixed | 428 | 39 | 483 | 61 | 1.44 (0.95–2.22) | 9.14e-02 | **†** | 1.20 (0.78–1.87) | 4.08e-01 |  | HIV.DURATION |
| AIDS-Defining Opportunistic Infections (All) | Mixed vs All High | 483 | 61 | 280 | 59 | 1.85 (1.25–2.74) | 2.22e-03 | ****** | 1.97 (1.32–2.93) | 8.71e-04 | ******* | HIV.DURATION |
| AIDS-Defining Opportunistic Infections (All) | All Low vs All High | 428 | 39 | 280 | 59 | 2.66 (1.73–4.15) | 1.11e-05 | ******* | 2.61 (1.68–4.09) | 2.32e-05 | ******* | HIV.DURATION |
| Post-cART Opportunistic Infections | All Low vs Mixed | 428 | 34 | 483 | 71 | 2.00 (1.31–3.11) | 1.68e-03 | ****** | 1.61 (1.04–2.55) | 3.74e-02 | ***** | HIV.DURATION + CMV.IgG.IUpermL |
| Post-cART Opportunistic Infections | Mixed vs All High | 483 | 71 | 280 | 38 | 0.91 (0.59–1.39) | 6.68e-01 |  | 0.95 (0.60–1.47) | 8.14e-01 |  | HIV.DURATION + CMV.IgG.IUpermL |
| Post-cART Opportunistic Infections | All Low vs All High | 428 | 34 | 280 | 38 | 1.82 (1.12–2.98) | 1.66e-02 | ***** | 1.53 (0.91–2.57) | 1.06e-01 |  | HIV.DURATION + CMV.IgG.IUpermL |
| Other AIDS-Defining Events (All) | All Low vs Mixed | 428 | 24 | 483 | 41 | 1.56 (0.93–2.66) | 9.40e-02 | **†** | 1.42 (0.83–2.46) | 2.03e-01 |  | AGE + HIV.DURATION + CENTER |
| Other AIDS-Defining Events (All) | Mixed vs All High | 483 | 41 | 280 | 39 | 1.74 (1.09–2.78) | 1.92e-02 | ***** | 1.76 (1.08–2.86) | 2.25e-02 | ***** | AGE + HIV.DURATION + CENTER |
| Other AIDS-Defining Events (All) | All Low vs All High | 428 | 24 | 280 | 39 | 2.72 (1.61–4.70) | 2.28e-04 | ******* | 2.68 (1.55–4.70) | 4.69e-04 | ******* | AGE + HIV.DURATION + CENTER |
| Endocrine & Metabolic | | | | | | | | | | | | |
| Endocrine/Metabolic Disorder (All) | All Low vs Mixed | 428 | 101 | 483 | 152 | 1.49 (1.11–2.00) | 8.28e-03 | ****** | 1.11 (0.80–1.56) | 5.27e-01 |  | AGE + HIV.DURATION + CMV.IgG.IUpermL |
| Endocrine/Metabolic Disorder (All) | Mixed vs All High | 483 | 152 | 280 | 88 | 1.00 (0.73–1.37) | 9.91e-01 |  | 0.96 (0.66–1.37) | 8.08e-01 |  | AGE + HIV.DURATION + CMV.IgG.IUpermL |
| Endocrine/Metabolic Disorder (All) | All Low vs All High | 428 | 101 | 280 | 88 | 1.48 (1.06–2.08) | 2.16e-02 | ***** | 1.09 (0.74–1.60) | 6.75e-01 |  | AGE + HIV.DURATION + CMV.IgG.IUpermL |
| Hypercholesterolemia | All Low vs Mixed | 428 | 93 | 483 | 133 | 1.37 (1.01–1.86) | 4.32e-02 | ***** | 1.07 (0.76–1.49) | 7.13e-01 |  | AGE + HIV.DURATION |
| Hypercholesterolemia | Mixed vs All High | 483 | 133 | 280 | 76 | 0.98 (0.70–1.36) | 9.07e-01 |  | 1.01 (0.70–1.45) | 9.73e-01 |  | AGE + HIV.DURATION |
| Hypercholesterolemia | All Low vs All High | 428 | 93 | 280 | 76 | 1.34 (0.94–1.90) | 9.90e-02 | **†** | 1.08 (0.73–1.59) | 6.86e-01 |  | AGE + HIV.DURATION |
| Gastroenterology | | | | | | | | | | | | |
| Gastroenterology Disease (All) | All Low vs Mixed | 428 | 80 | 483 | 127 | 1.55 (1.13–2.14) | 6.48e-03 | ****** | 1.40 (1.01–1.94) | 4.15e-02 | ***** | HIV.DURATION |
| Gastroenterology Disease (All) | Mixed vs All High | 483 | 127 | 280 | 67 | 0.88 (0.62–1.24) | 4.70e-01 |  | 0.94 (0.66–1.33) | 7.30e-01 |  | HIV.DURATION |
| Gastroenterology Disease (All) | All Low vs All High | 428 | 80 | 280 | 67 | 1.37 (0.95–1.97) | 9.37e-02 | **†** | 1.32 (0.91–1.92) | 1.41e-01 |  | HIV.DURATION |
| Musculoskeletal | | | | | | | | | | | | |
| Musculoskeletal Disorder (All) | All Low vs Mixed | 428 | 37 | 483 | 64 | 1.61 (1.06–2.49) | 2.82e-02 | ***** | 1.35 (0.87–2.12) | 1.80e-01 |  | AGE + HIV.DURATION |
| Musculoskeletal Disorder (All) | Mixed vs All High | 483 | 64 | 280 | 43 | 1.19 (0.78–1.80) | 4.20e-01 |  | 1.25 (0.81–1.92) | 3.11e-01 |  | AGE + HIV.DURATION |
| Musculoskeletal Disorder (All) | All Low vs All High | 428 | 37 | 280 | 43 | 1.92 (1.20–3.07) | 6.43e-03 | ****** | 1.69 (1.04–2.78) | 3.52e-02 | ***** | AGE + HIV.DURATION |
| Osteoporosis | All Low vs Mixed | 428 | 10 | 483 | 18 | 1.62 (0.75–3.68) | 2.29e-01 |  | 1.25 (0.56–2.91) | 5.92e-01 |  | AGE + HIV.DURATION + CMV.IgG.IUpermL |
| Osteoporosis | Mixed vs All High | 483 | 18 | 280 | 16 | 1.57 (0.78–3.13) | 2.03e-01 |  | 1.54 (0.73–3.22) | 2.52e-01 |  | AGE + HIV.DURATION + CMV.IgG.IUpermL |
| Osteoporosis | All Low vs All High | 428 | 10 | 280 | 16 | 2.53 (1.15–5.86) | 2.36e-02 | ***** | 1.87 (0.80–4.50) | 1.51e-01 |  | AGE + HIV.DURATION + CMV.IgG.IUpermL |
| Viremia | | | | | | | | | | | | |
| Residual Viremia (Inclusion) | All Low vs Mixed | 428 | 86 | 483 | 182 | 2.40 (1.79–3.26) | 9.51e-09 | ******* | 2.65 (1.95–3.61) | 5.70e-10 | ******* | HIV.DURATION |
| Residual Viremia (Inclusion) | Mixed vs All High | 483 | 182 | 280 | 123 | 1.30 (0.96–1.75) | 8.98e-02 | **†** | 1.23 (0.90–1.66) | 1.88e-01 |  | HIV.DURATION |
| Residual Viremia (Inclusion) | All Low vs All High | 428 | 86 | 280 | 123 | 3.12 (2.24–4.36) | 2.61e-11 | ******* | 3.27 (2.33–4.60) | 7.74e-12 | ******* | HIV.DURATION |
| Residual Viremia (2-Year Follow-Up) | All Low vs Mixed | 428 | 65 | 483 | 138 | 2.23 (1.61–3.12) | 1.76e-06 | ******* | 2.66 (1.87–3.83) | 8.04e-08 | ******* | CMV.IgG.IUpermL + HIV.DURATION |
| Residual Viremia (2-Year Follow-Up) | Mixed vs All High | 483 | 138 | 280 | 100 | 1.39 (1.01–1.90) | 4.04e-02 | ***** | 1.34 (0.97–1.85) | 7.61e-02 | **†** | CMV.IgG.IUpermL + HIV.DURATION |
| Residual Viremia (2-Year Follow-Up) | All Low vs All High | 428 | 65 | 280 | 100 | 3.10 (2.17–4.46) | 6.91e-10 | ******* | 3.48 (2.38–5.14) | 1.77e-10 | ******* | CMV.IgG.IUpermL + HIV.DURATION |
| *^1^*† p<0.1, * p<0.05, ** p<0.01, *** p<0.001 (Wald test) | | | | | | | | | | | | |

**Supplementary Table 11 | Confounder-checking sensitivity analysis for endotype–comorbidity associations (odds ratios).** Each row shows the odds ratio (OR) and 95% confidence interval (CI) for the endotype effect after individually adding one potential confounder to the base model (Outcome ∼ Endotype). Three pairwise comparisons are shown: Mixed vs. All Low, All High vs. All Low, and All High vs. Mixed. Analyses were performed in the pooled cohort (n = 1,191). Only complete cases were included for each model. Firth’s penalised logistic regression was used automatically for outcomes with fewer than 20 events. Outcomes were restricted to those reaching nominal significance (P < 0.05) in the main analysis (Supplementary Table 10). The same nine outcomes are shown for all comparisons. Covariates: Age, years at time of study visit. Sex at Birth, biological sex (male/female). BMI, body mass index (kg/m²) at baseline. CD4 Nadir, lowest recorded CD4⁺ T-cell count (×10⁹ cells/L) prior to ART initiation; used as a proxy for pre-ART immunological disease severity. Latest CD4 Count, most recent CD4⁺ T-cell count (×10⁹ cells/L) at the study visit. Hepatitis C, history of hepatitis C virus co-infection (yes/no). Study Center, participating clinical center (Radboudumc, EMC, OLVG, ETZ). COVID-19 Vaccination, COVID-19 vaccination status at the time of the study visit. TB History (pre- or post-cART), active tuberculosis diagnosis recorded before or after cART initiation (34 pre-cART events, 11 post-cART events in the full cohort). BCG Vaccination, self-reported history of BCG vaccination from questionnaire data (n = 265 vaccinated out of 1,658 with questionnaire data). *Pneumococcal Vaccination, vaccination against Streptococcus pneumoniae recorded during the 2-year follow-up window; this variable reflects vaccination status during follow-up only and does not capture baseline vaccination history. ART, antiretroviral therapy; CAP, controlled attenuation parameter; CI, confidence interval; IMT, intima-media thickness; LSM, liver stiffness measurement; OR, odds ratio.

### **A) Outcome: Residual Viremia (At Inclusion)**

| **Model** | **N** | **OR Mixed vs All Low** | **95% CI** | **N** | **OR All High vs All Low** | **95% CI** | **N** | **OR All High vs Mixed** | **95% CI** |
| --- | --- | --- | --- | --- | --- | --- | --- | --- | --- |
| **(base model)** | **911** | **2.4** | **1.79-3.26** | **708** | **3.12** | **2.24-4.36** | **763** | **1.3** | **0.96-1.75** |
| + Cohort (Discovery/Validation) | 911 | 2.55 | 1.89-3.48 | 708 | 3.31 | 2.36-4.66 | 763 | 1.3 | 0.96-1.76 |
| + Age | 911 | 2.39 | 1.77-3.25 | 708 | 3.12 | 2.23-4.39 | 763 | 1.29 | 0.96-1.75 |
| + Sex at Birth | 911 | 2.4 | 1.78-3.25 | 708 | 3.07 | 2.2-4.3 | 763 | 1.3 | 0.96-1.75 |
| + BMI | 911 | 2.4 | 1.79-3.26 | 708 | 3.12 | 2.24-4.37 | 763 | 1.3 | 0.97-1.76 |
| + Currently Smoking | 833 | 2.38 | 1.74-3.27 | 642 | 3.12 | 2.2-4.45 | 693 | 1.32 | 0.96-1.81 |
| + Ethnicity | 911 | 2.43 | 1.8-3.29 | 708 | 3.12 | 2.23-4.37 | 763 | 1.3 | 0.97-1.76 |
| + HIV Duration | 911 | 2.65 | 1.95-3.61 | 708 | 3.27 | 2.33-4.6 | 763 | 1.23 | 0.9-1.66 |
| + Time on ART | 893 | 2.56 | 1.88-3.5 | 688 | 3.12 | 2.22-4.4 | 761 | 1.21 | 0.89-1.65 |
| + CD4 Nadir | 890 | 2.69 | 1.94-3.76 | 695 | 3.32 | 2.3-4.83 | 749 | 1.32 | 0.98-1.79 |
| + Latest CD4 Count | 905 | 2.49 | 1.84-3.4 | 703 | 3.27 | 2.32-4.64 | 756 | 1.29 | 0.95-1.74 |
| + Hepatitis C | 911 | 2.42 | 1.8-3.27 | 708 | 3.13 | 2.25-4.39 | 763 | 1.29 | 0.96-1.74 |
| + Study Center | 911 | 2.56 | 1.89-3.49 | 708 | 3.31 | 2.35-4.68 | 763 | 1.29 | 0.95-1.75 |
| + COVID-19 Vaccination | 911 | 2.41 | 1.79-3.26 | 708 | 3.11 | 2.23-4.36 | 763 | 1.3 | 0.96-1.75 |
| + TB History (pre- or post-cART) | 911 | 2.4 | 1.79-3.26 | 708 | 3.14 | 2.25-4.41 | 763 | 1.3 | 0.96-1.75 |
| + BCG Vaccination | 780 | 2.2 | 1.6-3.06 | 592 | 2.83 | 1.97-4.09 | 642 | 1.29 | 0.93-1.8 |
| + Pneumococcal Vaccination (2Y follow-up)* | 692 | 2.46 | 1.75-3.48 | 527 | 3.12 | 2.13-4.61 | 573 | 1.27 | 0.9-1.8 |

### **B) Outcome: Residual Viremia (2-Years Follow-up)**

| **Model** | **N** | **OR Mixed vs All Low** | **95% CI** | **N** | **OR All High vs All Low** | **95% CI** | **N** | **OR All High vs Mixed** | **95% CI** |
| --- | --- | --- | --- | --- | --- | --- | --- | --- | --- |
| **(base model)** | **911** | **2.23** | **1.61-3.12** | **708** | **3.1** | **2.17-4.46** | **763** | **1.39** | **1.01-1.9** |
| + Cohort (Discovery/Validation) | 911 | 2.47 | 1.77-3.49 | 708 | 3.49 | 2.41-5.1 | 763 | 1.4 | 1.02-1.92 |
| + Age | 911 | 2.22 | 1.6-3.11 | 708 | 3.08 | 2.15-4.45 | 763 | 1.39 | 1.01-1.9 |
| + Sex at Birth | 911 | 2.24 | 1.62-3.13 | 708 | 3.07 | 2.15-4.42 | 763 | 1.39 | 1.01-1.9 |
| + BMI | 911 | 2.24 | 1.61-3.13 | 708 | 3.1 | 2.17-4.45 | 763 | 1.41 | 1.02-1.92 |
| + Currently Smoking | 833 | 2.15 | 1.54-3.03 | 642 | 2.88 | 1.98-4.2 | 693 | 1.34 | 0.96-1.87 |
| + Ethnicity | 911 | 2.26 | 1.63-3.17 | 708 | 3.14 | 2.19-4.53 | 763 | 1.4 | 1.02-1.92 |
| + HIV Duration | 911 | 2.38 | 1.7-3.34 | 708 | 3.14 | 2.19-4.52 | 763 | 1.37 | 1-1.88 |
| + Time on ART | 893 | 2.3 | 1.65-3.24 | 688 | 2.99 | 2.09-4.32 | 761 | 1.35 | 0.98-1.85 |
| + CD4 Nadir | 890 | 2.32 | 1.63-3.33 | 695 | 3.12 | 2.1-4.66 | 749 | 1.39 | 1.01-1.91 |
| + Latest CD4 Count | 905 | 2.34 | 1.68-3.3 | 703 | 3.32 | 2.29-4.84 | 756 | 1.42 | 1.03-1.95 |
| + Hepatitis C | 911 | 2.24 | 1.62-3.13 | 708 | 3.09 | 2.16-4.45 | 763 | 1.38 | 1.01-1.89 |
| + Study Center | 911 | 2.49 | 1.78-3.52 | 708 | 3.36 | 2.31-4.92 | 763 | 1.36 | 0.98-1.87 |
| + COVID-19 Vaccination | 911 | 2.2 | 1.59-3.08 | 708 | 3.09 | 2.16-4.45 | 763 | 1.4 | 1.02-1.92 |
| + TB History (pre- or post-cART) | 911 | 2.23 | 1.61-3.12 | 708 | 3.1 | 2.17-4.46 | 763 | 1.39 | 1.01-1.91 |
| + BCG Vaccination | 780 | 1.99 | 1.41-2.84 | 592 | 2.71 | 1.84-4.01 | 642 | 1.34 | 0.95-1.9 |
| + Pneumococcal Vaccination (2Y follow-up)* | 692 | 2.21 | 1.53-3.22 | 527 | 2.72 | 1.8-4.14 | 573 | 1.23 | 0.85-1.77 |

### **C) Outcome: Carotid Plaque**

| **Model** | **N** | **OR Mixed vs All Low** | **95% CI** | **N** | **OR All High vs All Low** | **95% CI** | **N** | **OR All High vs Mixed** | **95% CI** |
| --- | --- | --- | --- | --- | --- | --- | --- | --- | --- |
| **(base model)** | **911** | **1.69** | **1.3-2.2** | **708** | **1.69** | **1.25-2.3** | **763** | **1** | **0.74-1.35** |
| + Cohort (Discovery/Validation) | 911 | 1.68 | 1.29-2.18 | 708 | 1.68 | 1.24-2.28 | 763 | 1 | 0.74-1.35 |
| + Age | 911 | 1.37 | 1.03-1.83 | 708 | 1.43 | 1.02-2 | 763 | 1.05 | 0.76-1.45 |
| + Sex at Birth | 911 | 1.69 | 1.3-2.19 | 708 | 1.67 | 1.23-2.26 | 763 | 1 | 0.74-1.35 |
| + BMI | 911 | 1.69 | 1.3-2.2 | 708 | 1.7 | 1.26-2.31 | 763 | 1 | 0.74-1.34 |
| + Currently Smoking | 833 | 1.75 | 1.33-2.31 | 642 | 1.73 | 1.25-2.38 | 693 | 0.99 | 0.72-1.35 |
| + Ethnicity | 911 | 1.69 | 1.3-2.21 | 708 | 1.67 | 1.23-2.27 | 763 | 0.99 | 0.74-1.34 |
| + HIV Duration | 911 | 1.55 | 1.19-2.03 | 708 | 1.67 | 1.23-2.27 | 763 | 1.09 | 0.81-1.48 |
| + Time on ART | 893 | 1.52 | 1.16-2 | 688 | 1.64 | 1.2-2.25 | 761 | 1.12 | 0.82-1.52 |
| + CD4 Nadir | 890 | 1.47 | 1.1-1.95 | 695 | 1.43 | 1.03-1.99 | 749 | 0.99 | 0.73-1.34 |
| + Latest CD4 Count | 905 | 1.76 | 1.35-2.3 | 703 | 1.73 | 1.26-2.36 | 756 | 1 | 0.74-1.36 |
| + Hepatitis C | 911 | 1.69 | 1.3-2.2 | 708 | 1.71 | 1.26-2.32 | 763 | 1 | 0.74-1.34 |
| + Study Center | 911 | 1.77 | 1.34-2.34 | 708 | 1.6 | 1.16-2.2 | 763 | 0.92 | 0.67-1.26 |
| + COVID-19 Vaccination | 911 | 1.72 | 1.32-2.25 | 708 | 1.73 | 1.28-2.36 | 763 | 1 | 0.74-1.35 |
| + TB History (pre- or post-cART) | 911 | 1.69 | 1.3-2.2 | 708 | 1.69 | 1.25-2.3 | 763 | 0.99 | 0.74-1.34 |
| + BCG Vaccination | 780 | 1.84 | 1.39-2.45 | 592 | 1.78 | 1.28-2.5 | 642 | 0.98 | 0.7-1.36 |
| + Pneumococcal Vaccination (2Y follow-up)* | 692 | 1.69 | 1.24-2.31 | 527 | 1.64 | 1.14-2.37 | 573 | 0.98 | 0.69-1.4 |

### **D) Outcome: Musculoskeletal Disorders**

| **Model** | **N** | **OR Mixed vs All Low** | **95% CI** | **N** | **OR All High vs All Low** | **95% CI** | **N** | **OR All High vs Mixed** | **95% CI** |
| --- | --- | --- | --- | --- | --- | --- | --- | --- | --- |
| **(base model)** | **911** | **1.61** | **1.06-2.49** | **708** | **1.92** | **1.2-3.07** | **763** | **1.19** | **0.78-1.8** |
| + Cohort (Discovery/Validation) | 911 | 1.69 | 1.1-2.62 | 708 | 1.93 | 1.21-3.09 | 763 | 1.19 | 0.78-1.8 |
| + Age | 911 | 1.41 | 0.92-2.21 | 708 | 1.66 | 1.03-2.7 | 763 | 1.18 | 0.77-1.81 |
| + Sex at Birth | 911 | 1.66 | 1.08-2.58 | 708 | 2.01 | 1.25-3.24 | 763 | 1.21 | 0.79-1.84 |
| + BMI | 911 | 1.61 | 1.06-2.49 | 708 | 1.9 | 1.19-3.05 | 763 | 1.18 | 0.77-1.79 |
| + Currently Smoking | 833 | 1.56 | 1.01-2.45 | 642 | 1.86 | 1.15-3.05 | 693 | 1.19 | 0.77-1.84 |
| + Ethnicity | 911 | 1.63 | 1.07-2.52 | 708 | 1.89 | 1.18-3.05 | 763 | 1.18 | 0.77-1.79 |
| + HIV Duration | 911 | 1.41 | 0.91-2.19 | 708 | 1.84 | 1.13-2.99 | 763 | 1.31 | 0.85-2 |
| + Time on ART | 893 | 1.34 | 0.87-2.09 | 688 | 1.8 | 1.11-2.94 | 761 | 1.32 | 0.86-2.02 |
| + CD4 Nadir | 890 | 1.32 | 0.84-2.1 | 695 | 1.25 | 0.76-2.08 | 749 | 1.16 | 0.76-1.77 |
| + Latest CD4 Count | 905 | 1.62 | 1.05-2.52 | 703 | 2.01 | 1.24-3.28 | 756 | 1.24 | 0.8-1.89 |
| + Hepatitis C | 911 | 1.61 | 1.06-2.49 | 708 | 1.92 | 1.2-3.08 | 763 | 1.19 | 0.78-1.8 |
| + Study Center | 911 | 1.69 | 1.1-2.62 | 708 | 1.93 | 1.2-3.13 | 763 | 1.16 | 0.75-1.76 |
| + COVID-19 Vaccination | 911 | 1.59 | 1.04-2.46 | 708 | 1.91 | 1.19-3.06 | 763 | 1.19 | 0.78-1.8 |
| + TB History (pre- or post-cART) | 911 | 1.62 | 1.06-2.5 | 708 | 1.85 | 1.15-2.97 | 763 | 1.18 | 0.77-1.79 |
| + BCG Vaccination | 780 | 1.49 | 0.95-2.35 | 592 | 1.89 | 1.15-3.12 | 642 | 1.26 | 0.8-1.97 |
| + Pneumococcal Vaccination (2Y follow-up)* | 692 | 1.67 | 1.02-2.77 | 527 | 2.18 | 1.27-3.76 | 573 | 1.31 | 0.81-2.1 |

### **E) Outcome: AIDS-defining Opportunistic Infections**

| **Model** | **N** | **OR Mixed vs All Low** | **95% CI** | **N** | **OR All High vs All Low** | **95% CI** | **N** | **OR All High vs Mixed** | **95% CI** |
| --- | --- | --- | --- | --- | --- | --- | --- | --- | --- |
| **(base model)** | **911** | **1.44** | **0.95-2.22** | **708** | **2.66** | **1.73-4.15** | **763** | **1.85** | **1.25-2.74** |
| + Cohort (Discovery/Validation) | 911 | 1.46 | 0.95-2.24 | 708 | 2.71 | 1.75-4.23 | 763 | 1.85 | 1.25-2.74 |
| + Age | 911 | 1.31 | 0.86-2.03 | 708 | 2.46 | 1.59-3.85 | 763 | 1.86 | 1.25-2.77 |
| + Sex at Birth | 911 | 1.46 | 0.96-2.25 | 708 | 2.69 | 1.74-4.2 | 763 | 1.88 | 1.26-2.79 |
| + BMI | 911 | 1.44 | 0.95-2.22 | 708 | 2.65 | 1.72-4.13 | 763 | 1.84 | 1.24-2.73 |
| + Currently Smoking | 833 | 1.67 | 1.07-2.64 | 642 | 2.91 | 1.83-4.68 | 693 | 1.75 | 1.16-2.63 |
| + Ethnicity | 911 | 1.42 | 0.93-2.2 | 708 | 2.68 | 1.73-4.18 | 763 | 1.87 | 1.26-2.79 |
| + HIV Duration | 911 | 1.2 | 0.78-1.87 | 708 | 2.61 | 1.68-4.09 | 763 | 1.97 | 1.32-2.93 |
| + Time on ART | 893 | 1.13 | 0.73-1.76 | 688 | 2.53 | 1.62-4 | 761 | 2.07 | 1.38-3.1 |
| + CD4 Nadir | 890 | 0.64 | 0.39-1.05 | 695 | 1.03 | 0.62-1.72 | 749 | 1.51 | 0.94-2.4 |
| + Latest CD4 Count | 905 | 1.22 | 0.79-1.9 | 703 | 2.42 | 1.55-3.81 | 756 | 1.82 | 1.22-2.71 |
| + Hepatitis C | 911 | 1.45 | 0.95-2.24 | 708 | 2.66 | 1.73-4.15 | 763 | 1.86 | 1.25-2.76 |
| + Study Center | 911 | 1.45 | 0.95-2.24 | 708 | 2.77 | 1.79-4.35 | 763 | 1.82 | 1.22-2.7 |
| + COVID-19 Vaccination | 911 | 1.45 | 0.95-2.23 | 708 | 2.69 | 1.74-4.19 | 763 | 1.85 | 1.25-2.75 |
| + TB History (pre- or post-cART) | 911 | 1.47 | 0.95-2.28 | 708 | 2.52 | 1.61-3.96 | 763 | 1.73 | 1.16-2.59 |
| + BCG Vaccination | 780 | 1.53 | 0.97-2.44 | 592 | 2.49 | 1.54-4.07 | 642 | 1.64 | 1.06-2.53 |
| + Pneumococcal Vaccination (2Y follow-up)* | 692 | 1.47 | 0.91-2.42 | 527 | 2.61 | 1.57-4.39 | 573 | 1.78 | 1.12-2.81 |

### **F) Outcome: AIDS-defining Malignancies**

| **Model** | **N** | **OR Mixed vs All Low** | **95% CI** | **N** | **OR All High vs All Low** | **95% CI** | **N** | **OR All High vs Mixed** | **95% CI** |
| --- | --- | --- | --- | --- | --- | --- | --- | --- | --- |
| **(base model)** | **911** | **2.11** | **1.21-3.82** | **708** | **2.04** | **1.08-3.9** | **763** | **0.96** | **0.56-1.63** |
| + Cohort (Discovery/Validation) | 911 | 2.13 | 1.22-3.85 | 708 | 2.02 | 1.07-3.88 | 763 | 0.96 | 0.56-1.63 |
| + Age | 911 | 1.9 | 1.09-3.45 | 708 | 1.93 | 1.02-3.71 | 763 | 0.97 | 0.56-1.64 |
| + Sex at Birth | 911 | 2.09 | 1.2-3.78 | 708 | 1.97 | 1.04-3.77 | 763 | 0.94 | 0.54-1.59 |
| + BMI | 911 | 2.11 | 1.21-3.83 | 708 | 2.05 | 1.09-3.93 | 763 | 0.98 | 0.57-1.66 |
| + Currently Smoking | 833 | 2.12 | 1.2-3.91 | 642 | 2 | 1.03-3.91 | 693 | 0.94 | 0.53-1.62 |
| + Ethnicity | 911 | 2.1 | 1.2-3.82 | 708 | 2.05 | 1.09-3.94 | 763 | 0.96 | 0.56-1.64 |
| + HIV Duration | 911 | 2.03 | 1.16-3.69 | 708 | 1.99 | 1.06-3.82 | 763 | 0.95 | 0.55-1.62 |
| + Time on ART | 893 | 1.82 | 1.04-3.31 | 688 | 1.9 | 1-3.64 | 761 | 1.01 | 0.58-1.71 |
| + CD4 Nadir | 890 | 1.34 | 0.75-2.47 | 695 | 1.18 | 0.6-2.35 | 749 | 0.88 | 0.5-1.49 |
| + Latest CD4 Count | 905 | 1.85 | 1.05-3.37 | 703 | 2 | 1.05-3.88 | 756 | 0.93 | 0.54-1.58 |
| + Hepatitis C | 911 | 2.1 | 1.21-3.8 | 708 | 2.05 | 1.09-3.93 | 763 | 0.97 | 0.56-1.64 |
| + Study Center | 911 | 2.13 | 1.21-3.86 | 708 | 1.95 | 1.03-3.76 | 763 | 0.94 | 0.54-1.61 |
| + COVID-19 Vaccination | 911 | 2.11 | 1.21-3.81 | 708 | 2.02 | 1.07-3.86 | 763 | 0.97 | 0.56-1.64 |
| + TB History (pre- or post-cART) | 911 | 2.11 | 1.21-3.83 | 708 | 2.01 | 1.07-3.86 | 763 | 0.93 | 0.53-1.58 |
| + BCG Vaccination | 780 | 2.3 | 1.26-4.39 | 592 | 1.91 | 0.93-3.96 | 642 | 0.83 | 0.45-1.49 |
| + Pneumococcal Vaccination (2Y follow-up)* | 692 | 2.67 | 1.4-5.46 | 527 | 2.09 | 0.96-4.66 | 573 | 0.76 | 0.39-1.4 |

### **G) Outcome: Other AIDS-defining Comorbidities**

| **Model** | **N** | **OR Mixed vs All Low** | **95% CI** | **N** | **OR All High vs All Low** | **95% CI** | **N** | **OR All High vs Mixed** | **95% CI** |
| --- | --- | --- | --- | --- | --- | --- | --- | --- | --- |
| **(base model)** | **911** | **1.56** | **0.93-2.66** | **708** | **2.72** | **1.61-4.7** | **763** | **1.74** | **1.09-2.78** |
| + Cohort (Discovery/Validation) | 911 | 1.61 | 0.96-2.76 | 708 | 2.86 | 1.68-4.97 | 763 | 1.75 | 1.09-2.79 |
| + Age | 911 | 1.38 | 0.82-2.37 | 708 | 2.47 | 1.45-4.27 | 763 | 1.75 | 1.1-2.8 |
| + Sex at Birth | 911 | 1.56 | 0.94-2.67 | 708 | 2.71 | 1.6-4.68 | 763 | 1.75 | 1.09-2.79 |
| + BMI | 911 | 1.56 | 0.93-2.67 | 708 | 2.72 | 1.61-4.69 | 763 | 1.76 | 1.1-2.81 |
| + Currently Smoking | 833 | 1.7 | 0.99-3 | 642 | 2.77 | 1.58-4.97 | 693 | 1.63 | 0.99-2.66 |
| + Ethnicity | 911 | 1.54 | 0.92-2.64 | 708 | 2.79 | 1.64-4.83 | 763 | 1.76 | 1.1-2.81 |
| + HIV Duration | 911 | 1.51 | 0.9-2.59 | 708 | 2.72 | 1.61-4.69 | 763 | 1.64 | 1.02-2.63 |
| + Time on ART | 893 | 1.35 | 0.8-2.32 | 688 | 2.58 | 1.52-4.46 | 761 | 1.74 | 1.08-2.78 |
| + CD4 Nadir | 890 | 0.71 | 0.4-1.29 | 695 | 1.01 | 0.55-1.86 | 749 | 1.4 | 0.84-2.34 |
| + Latest CD4 Count | 905 | 1.26 | 0.74-2.17 | 703 | 2.2 | 1.28-3.85 | 756 | 1.64 | 1.02-2.64 |
| + Hepatitis C | 911 | 1.57 | 0.94-2.68 | 708 | 2.72 | 1.61-4.69 | 763 | 1.72 | 1.08-2.74 |
| + Study Center | 911 | 1.62 | 0.96-2.78 | 708 | 3.03 | 1.77-5.3 | 763 | 1.92 | 1.19-3.09 |
| + COVID-19 Vaccination | 911 | 1.56 | 0.93-2.66 | 708 | 2.7 | 1.6-4.66 | 763 | 1.75 | 1.1-2.8 |
| + TB History (pre- or post-cART) | 911 | 1.56 | 0.93-2.66 | 708 | 2.63 | 1.55-4.54 | 763 | 1.71 | 1.07-2.74 |
| + BCG Vaccination | 780 | 1.49 | 0.86-2.67 | 592 | 2.33 | 1.29-4.26 | 642 | 1.55 | 0.91-2.63 |
| + Pneumococcal Vaccination (2Y follow-up)* | 692 | 1.25 | 0.69-2.29 | 527 | 2.33 | 1.27-4.33 | 573 | 1.87 | 1.06-3.28 |

### **H) Outcome: Gastroenterology Disease**

| **Model** | **N** | **OR Mixed vs All Low** | **95% CI** | **N** | **OR All High vs All Low** | **95% CI** | **N** | **OR All High vs Mixed** | **95% CI** |
| --- | --- | --- | --- | --- | --- | --- | --- | --- | --- |
| **(base model)** | **911** | **1.55** | **1.13-2.14** | **708** | **1.37** | **0.95-1.97** | **763** | **0.88** | **0.62-1.24** |
| + Cohort (Discovery/Validation) | 911 | 1.49 | 1.08-2.05 | 708 | 1.31 | 0.9-1.89 | 763 | 0.88 | 0.62-1.24 |
| + Age | 911 | 1.41 | 1.02-1.95 | 708 | 1.23 | 0.85-1.79 | 763 | 0.88 | 0.62-1.24 |
| + Sex at Birth | 911 | 1.54 | 1.13-2.13 | 708 | 1.34 | 0.93-1.93 | 763 | 0.88 | 0.62-1.23 |
| + BMI | 911 | 1.55 | 1.13-2.14 | 708 | 1.39 | 0.96-2 | 763 | 0.9 | 0.63-1.26 |
| + Currently Smoking | 833 | 1.45 | 1.05-2.03 | 642 | 1.27 | 0.86-1.86 | 693 | 0.87 | 0.6-1.25 |
| + Ethnicity | 911 | 1.58 | 1.15-2.18 | 708 | 1.39 | 0.96-2.01 | 763 | 0.87 | 0.61-1.22 |
| + HIV Duration | 911 | 1.4 | 1.01-1.94 | 708 | 1.32 | 0.91-1.92 | 763 | 0.94 | 0.66-1.33 |
| + Time on ART | 893 | 1.41 | 1.02-1.96 | 688 | 1.33 | 0.91-1.93 | 761 | 0.96 | 0.68-1.35 |
| + CD4 Nadir | 890 | 1.38 | 0.98-1.95 | 695 | 1.21 | 0.81-1.8 | 749 | 0.92 | 0.65-1.29 |
| + Latest CD4 Count | 905 | 1.55 | 1.13-2.15 | 703 | 1.32 | 0.9-1.93 | 756 | 0.87 | 0.62-1.23 |
| + Hepatitis C | 911 | 1.54 | 1.12-2.13 | 708 | 1.44 | 0.98-2.1 | 763 | 0.92 | 0.65-1.29 |
| + Study Center | 911 | 1.48 | 1.08-2.05 | 708 | 1.26 | 0.86-1.83 | 763 | 0.92 | 0.64-1.29 |
| + COVID-19 Vaccination | 911 | 1.53 | 1.11-2.1 | 708 | 1.36 | 0.94-1.96 | 763 | 0.88 | 0.63-1.24 |
| + TB History (pre- or post-cART) | 911 | 1.55 | 1.13-2.14 | 708 | 1.38 | 0.95-1.99 | 763 | 0.88 | 0.62-1.23 |
| + BCG Vaccination | 780 | 1.52 | 1.08-2.13 | 592 | 1.38 | 0.93-2.05 | 642 | 0.91 | 0.62-1.31 |
| + Pneumococcal Vaccination (2Y follow-up)* | 692 | 1.47 | 1.02-2.13 | 527 | 1.09 | 0.7-1.69 | 573 | 0.75 | 0.49-1.12 |

**Supplementary Table 12 | Post-2015 Diagnosis Sensitivity Analysis.** Restricted to participants diagnosed with HIV after 2015, who started ART immediately (immediate ART guidelines), minimizing pre-ART disease accumulation. Unadjusted and adjusted (Age, Sex, BMI, Smoking, Ethnicity, HIV Duration, Time on ART, Center, Cohort) logistic regression models. Firth's penalized logistic regression used for outcomes with <20 events.

| **Outcome** | **Model** | **N** | **OR Mixed vs All Low** | **95% CI** | **p** | **N** | **OR All High vs All Low** | **95% CI** | **p** | **N** | **OR All High vs Mixed** | **95% CI** | **p** | **Covariates used** |
| --- | --- | --- | --- | --- | --- | --- | --- | --- | --- | --- | --- | --- | --- | --- |
| **Residual viremia (baseline)** | Unadjusted | 154 | 2 | 1-4.03 | **0.049** | 157 | 5.03 | 2.55-10.24 | <0.001 | 119 | 2.51 | 1.21-5.32 | **0.015** | — |
|  | Adjusted (main model covariates) | 154 | 2 | 1-4.03 | **0.049** | 157 | 5.03 | 2.55-10.24 | <0.001 | 119 | 2.51 | 1.21-5.32 | **0.015** | — |
|  | Sensitivity: + Age only | 154 | 1.74 | 0.85-3.57 | 0.130 | 157 | 4.68 | 2.35-9.59 | <0.001 | 119 | 2.51 | 1.21-5.32 | **0.015** | AGE |
|  | Sensitivity: + CD4 Nadir only | 152 | 1.69 | 0.8-3.57 | 0.168 | 155 | 3.62 | 1.66-8.07 | **0.001** | 119 | 2.52 | 1.2-5.43 | **0.016** | CD4_NADIR |
| **Residual viremia (2Y follow-up)** | Unadjusted | 154 | 2.85 | 1.37-6 | **0.005** | 157 | 3.01 | 1.47-6.28 | **0.003** | 119 | 1.06 | 0.51-2.21 | 0.883 | — |
|  | Adjusted (main model covariates) | 154 | 2.85 | 1.37-6 | **0.005** | 157 | 3.01 | 1.47-6.28 | **0.003** | 119 | 1.06 | 0.51-2.21 | 0.883 | — |
|  | Sensitivity: + Age only | 154 | 2.5 | 1.18-5.36 | **0.017** | 157 | 2.79 | 1.35-5.89 | **0.006** | 119 | 1.07 | 0.51-2.25 | 0.849 | AGE |
|  | Sensitivity: + CD4 Nadir only | 152 | 2.58 | 1.18-5.74 | **0.018** | 155 | 2.86 | 1.24-6.75 | **0.015** | 119 | 0.97 | 0.46-2.06 | 0.942 | CD4_NADIR |
| **Carotid plaque** | Unadjusted | 154 | 1.21 | 0.62-2.34 | 0.576 | 157 | 1.27 | 0.66-2.43 | 0.477 | 119 | 1.05 | 0.51-2.17 | 0.899 | — |
|  | Adjusted (main model covariates) | 137 | 0.46 | 0.17-1.19 | 0.119 | 139 | 0.58 | 0.22-1.46 | 0.257 | 108 | 1.48 | 0.58-3.92 | 0.414 | AGE, HIV_DURATION, CENTER, CMV_IgG_IUpermL |
|  | Sensitivity: + Age only | 154 | 0.69 | 0.31-1.49 | 0.357 | 157 | 0.76 | 0.34-1.66 | 0.502 | 119 | 1.15 | 0.53-2.51 | 0.729 | AGE |
|  | Sensitivity: + CD4 Nadir only | 152 | 1.21 | 0.59-2.47 | 0.601 | 155 | 1.22 | 0.57-2.6 | 0.603 | 119 | 0.95 | 0.45-2 | 0.900 | CD4_NADIR |
| **Musculoskeletal disorders** | Unadjusted | 154 | 1.68 | 0.35-8.2 | 0.503 | 157 | 1.6 | 0.33-7.77 | 0.546 | 119 | 0.95 | 0.19-4.65 | 0.946 | — |
|  | Adjusted (main model covariates) | 154 | 1.68 | 0.35-8.2 | 0.503 | 157 | 1.6 | 0.33-7.77 | 0.546 | 119 | 0.95 | 0.19-4.65 | 0.946 | — |
|  | Sensitivity: + Age only | 154 | 1.22 | 0.24-6.06 | 0.802 | 157 | 1.13 | 0.22-5.65 | 0.881 | 119 | 0.99 | 0.2-4.89 | 0.994 | AGE |
|  | Sensitivity: + CD4 Nadir only | 152 | 1.43 | 0.25-7.85 | 0.675 | 155 | 0.57 | 0.08-3.83 | 0.557 | 119 | 0.89 | 0.18-4.43 | 0.884 | CD4_NADIR |
| **AIDS-defining opportunistic infections** | Unadjusted | 154 | 0.99 | 0.09-7.6 | 0.989 | 157 | 9.55 | 2.72-50.3 | <0.001 | 119 | 9.68 | 2.21-91.09 | **0.001** | — |
|  | Adjusted (main model covariates) | 154 | 0.99 | 0.09-7.6 | 0.989 | 157 | 9.55 | 2.72-50.3 | <0.001 | 119 | 9.68 | 2.21-91.09 | **0.001** | — |
|  | Sensitivity: + Age only | 154 | 0.79 | 0.07-6.49 | 0.829 | 157 | 8.49 | 2.39-44.99 | **0.001** | 119 | 10.23 | 2.32-96.68 | **0.001** | AGE |
|  | Sensitivity: + CD4 Nadir only | 152 | 0.16 | 0.01-1.97 | 0.135 | 155 | 0.69 | 0.06-6.22 | 0.734 | 119 | 3.46 | 0.53-38.1 | 0.200 | CD4_NADIR |
| **AIDS-defining malignancies** | Unadjusted | 154 | 28.11 | 3.31-3674.84 | **0.001** | 157 | 15.1 | 1.57-2015.36 | **0.015** | 119 | 0.54 | 0.14-1.79 | 0.314 | — |
|  | Adjusted (main model covariates) | 154 | 28.11 | 3.31-3674.84 | **0.001** | 157 | 15.1 | 1.57-2015.36 | **0.015** | 119 | 0.54 | 0.14-1.79 | 0.314 | — |
|  | Sensitivity: + Age only | 154 | 20.31 | 2.36-2659.12 | **0.003** | 157 | 24.94 | 2.42-3394.04 | **0.004** | 119 | 0.55 | 0.15-1.81 | 0.325 | AGE |
|  | Sensitivity: + CD4 Nadir only | 152 | 9.18 | 0.98-1216.34 | 0.053 | 155 | 2.2 | 0.16-345.31 | 0.609 | 119 | 0.23 | 0.05-0.93 | **0.039** | CD4_NADIR |
| **Other AIDS-defining events** | Unadjusted | 154 | 3.12 | 0.67-18.39 | 0.147 | 157 | 8.61 | 2.42-45.62 | **0.001** | 119 | 2.76 | 0.92-9.73 | 0.072 | — |
|  | Adjusted (main model covariates) | 154 | 3.12 | 0.67-18.39 | 0.147 | 157 | 8.61 | 2.42-45.62 | **0.001** | 119 | 2.76 | 0.92-9.73 | 0.072 | — |
|  | Sensitivity: + Age only | 154 | 2.51 | 0.52-15.16 | 0.252 | 157 | 7.82 | 2.17-41.76 | **0.001** | 119 | 2.77 | 0.92-9.76 | 0.070 | AGE |
|  | Sensitivity: + CD4 Nadir only | 152 | 0.72 | 0.11-4.92 | 0.721 | 155 | 0.94 | 0.15-6.36 | 0.948 | 119 | 1.28 | 0.32-5.29 | 0.728 | CD4_NADIR |
| **Post-cART opportunistic infections** | Unadjusted | 154 | 6.55 | 1.26-65.04 | **0.024** | 157 | 2.68 | 0.35-29.75 | 0.337 | 119 | 0.41 | 0.07-1.78 | 0.237 | — |
|  | Adjusted (main model covariates) | 154 | 6.55 | 1.26-65.04 | **0.024** | 157 | 2.68 | 0.35-29.75 | 0.337 | 119 | 0.41 | 0.07-1.78 | 0.237 | — |
|  | Sensitivity: + Age only | 154 | 7.06 | 1.28-73 | **0.024** | 157 | 2.71 | 0.34-31.25 | 0.342 | 119 | 0.4 | 0.07-1.74 | 0.225 | AGE |
|  | Sensitivity: + CD4 Nadir only | 152 | 4.56 | 0.73-49.74 | 0.108 | 155 | 0.21 | 0.02-2.95 | 0.220 | 119 | 0.36 | 0.06-1.62 | 0.187 | CD4_NADIR |
| **Gastrointestinal disease** | Unadjusted | 154 | 1.75 | 0.63-4.9 | 0.279 | 157 | 1.43 | 0.49-4.08 | 0.499 | 119 | 0.82 | 0.28-2.36 | 0.708 | — |
|  | Adjusted (main model covariates) | 154 | 1.75 | 0.63-4.9 | 0.279 | 157 | 1.43 | 0.49-4.08 | 0.499 | 119 | 0.82 | 0.28-2.36 | 0.708 | — |
|  | Sensitivity: + Age only | 154 | 1.24 | 0.42-3.59 | 0.692 | 157 | 1.22 | 0.41-3.53 | 0.717 | 119 | 0.85 | 0.29-2.46 | 0.756 | AGE |
|  | Sensitivity: + CD4 Nadir only | 152 | 1.87 | 0.63-5.62 | 0.257 | 155 | 1.73 | 0.51-5.86 | 0.372 | 119 | 0.9 | 0.3-2.64 | 0.840 | CD4_NADIR |

Post-2015 subset: participants diagnosed after 2015, who started ART immediately. Three separate pairwise binary logistic regression models per comparison, matching the main analysis approach. Blue rows = adjusted (same covariates as main analysis). Orange rows = age-only sensitivity. Green rows = CD4 nadir-only sensitivity. Bold p-values = p<0.05. Firth's penalised logistic regression used for outcomes with <20 events.

**Supplementary Table 13 | Mediation Analysis for the Association Between Endotypes and Comorbidities.**

This table summarizes causal mediation analyses assessing whether total HIV DNA (log-transformed) mediates the association between pairwise endotype comparisons and clinical comorbidities. For each outcome and comparison, the table reports results from both the model-based product-of-coefficients approach (a×b) and the potential-outcomes framework implemented in the mediate() function (Imai et al., 2010). Columns include the total effect of endotype on the outcome (Y~X), the a-path (X→M) and b-path (M→Y), the direct effect (X adjusted for M), the manual indirect effect (a×b) with bootstrap confidence intervals, and the average causal mediation effect (ACME), average direct effect (ADE), total effect, and proportion mediated estimated by mediate() using 5,000 bootstrap simulations and the nonparametric potential-outcomes framework. Confounders correspond to those used in the adjusted odds-ratio models for each outcome. Significance of mediation is based on ACME (95% CI excluding 0). The final column summarizes whether mediation was complete, partial, absent, or inconsistent (suppression). Analyses were conducted using complete-case data from the pooled cohort (n=1,191). Significance of mediation was determined using ACME (p<0.05 or 95% CI excluding 0). Cases where ACME was significant but ADE remained non-zero (p<0.05) were classified as partial mediation, whereas non-significant ADE indicated complete mediation. All mediation results were compiled across adjusted and unadjusted models, with complete-case data used for each mediator–outcome pair. Analyses were conducted in R (v4.3.0).M = Mediator (total HIV DNA), X = Predictor (Endotype), Y = Outcome (Comorbidity “Yes”/”No”).

| Comparison | Run | Confounders | Y ~ X (total) | X -> M (a) | M -> Y (b) | Y ~ X + M (X direct) | Manual IE (a*b) | Manual IE boot CI | Prop mediated (manual) | ACME (Imai bootstrap) | ACME significant? | ADE (Imai bootstrap) | ADE significant? | Total (Imai bootstrap) | Mediate prop (Imai bootstrap) | Conclusion |
| --- | --- | --- | --- | --- | --- | --- | --- | --- | --- | --- | --- | --- | --- | --- | --- | --- |
| Carotid Plaque | | | | | | | | | | | | | | | | |
| All Low vs Mixed | Unadjusted | AGE, SMOKING.CURRENT | 1.690 (95% CI 1.300–2.197), p=0.000 | 2.546 (SE 0.071), p=0.000 | 1.174 (95% CI 1.036–1.330), p=0.012 | 1.128 (95% CI 0.751–1.695), p=0.561 | 0.408 (log-odds), OR=1.504 (95% CI 1.094–2.068) | Boot CI [0.091, 0.736] | 77.8% | 0.101 (95% CI 0.026–0.180), p=0.010 | TRUE | 0.030 (95% CI -0.070–0.130), p=0.581 | FALSE | 0.130 (95% CI 0.065–0.195), p=0.000 | 0.772920193 | Significant complete mediation |
| All Low vs Mixed | Adjusted | AGE, SMOKING.CURRENT | 1.414 (95% CI 1.048–1.908), p=0.024 | 2.521 (SE 0.076), p=0.000 | 1.114 (95% CI 0.967–1.284), p=0.136 | 1.079 (95% CI 0.679–1.716), p=0.748 | 0.272 (log-odds), OR=1.313 (95% CI 0.918–1.878) | Boot CI [-0.095, 0.665] | 78.6% | 0.057 (95% CI -0.019–0.130), p=0.146 | FALSE | 0.016 (95% CI -0.079–0.116), p=0.730 | FALSE | 0.073 (95% CI 0.010–0.135), p=0.022 | 0.781804276 | No significant mediation (ACME not significant) |
| Mixed vs All High | Unadjusted | AGE, SMOKING.CURRENT | 1.002 (95% CI 0.745–1.349), p=0.988 | 0.668 (SE 0.074), p=0.000 | 1.190 (95% CI 1.026–1.380), p=0.021 | 0.893 (95% CI 0.653–1.222), p=0.480 | 0.116 (log-odds), OR=1.123 (95% CI 1.014–1.244) | Boot CI [0.019, 0.223] | 5011.8% (>100%) | 0.028 (95% CI 0.005–0.054), p=0.020 | TRUE | -0.027 (95% CI -0.105–0.050), p=0.483 | FALSE | 0.001 (95% CI -0.073–0.076), p=0.992 | 35.523294895 | Significant complete mediation |
| Mixed vs All High | Adjusted | AGE, SMOKING.CURRENT | 1.075 (95% CI 0.765–1.512), p=0.677 | 0.652 (SE 0.078), p=0.000 | 1.175 (95% CI 0.994–1.389), p=0.059 | 0.972 (95% CI 0.680–1.390), p=0.877 | 0.105 (log-odds), OR=1.111 (95% CI 0.993–1.242) | Boot CI [-0.003, 0.227] | 144.8% (>100%) | 0.022 (95% CI -0.001–0.046), p=0.060 | FALSE | -0.006 (95% CI -0.078–0.069), p=0.886 | FALSE | 0.016 (95% CI -0.054–0.088), p=0.635 | 1.368224312 | No significant mediation (ACME not significant) |
| All Low vs All High | Unadjusted | AGE, SMOKING.CURRENT | 1.694 (95% CI 1.250–2.295), p=0.001 | 3.214 (SE 0.094), p=0.000 | 1.125 (95% CI 0.995–1.273), p=0.061 | 1.162 (95% CI 0.708–1.908), p=0.553 | 0.380 (log-odds), OR=1.462 (95% CI 0.982–2.176) | Boot CI [-0.015, 0.803] | 72.0% | 0.094 (95% CI -0.005–0.194), p=0.059 | FALSE | 0.037 (95% CI -0.084–0.161), p=0.564 | FALSE | 0.131 (95% CI 0.056–0.206), p=0.002 | 0.717626252 | No significant mediation (ACME not significant) |
| All Low vs All High | Adjusted | AGE, SMOKING.CURRENT | 1.513 (95% CI 1.061–2.158), p=0.022 | 3.156 (SE 0.098), p=0.000 | 0.994 (95% CI 0.860–1.150), p=0.940 | 1.540 (95% CI 0.863–2.747), p=0.144 | -0.018 (log-odds), OR=0.983 (95% CI 0.621–1.556) | Boot CI [-0.501, 0.459] | -4.2% | -0.004 (95% CI -0.097–0.092), p=0.947 | FALSE | 0.087 (95% CI -0.030–0.203), p=0.153 | FALSE | 0.083 (95% CI 0.011–0.156), p=0.024 | -0.042294796 | No significant mediation (ACME not significant) |
| Residual Viremia (At Inclusion) | | | | | | | | | | | | | | | | |
| All Low vs Mixed | Unadjusted | HIV.DURATION | 2.405 (95% CI 1.782–3.245), p=0.000 | 2.546 (SE 0.071), p=0.000 | 1.308 (95% CI 1.126–1.519), p=0.000 | 1.248 (95% CI 0.785–1.984), p=0.348 | 0.683 (log-odds), OR=1.979 (95% CI 1.349–2.903) | Boot CI [0.321, 1.083] | 77.8% | 0.135 (95% CI 0.063–0.206), p=0.000 | TRUE | 0.043 (95% CI -0.047–0.134), p=0.348 | FALSE | 0.178 (95% CI 0.119–0.236), p=0.000 | 0.756051772 | Significant complete mediation |
| All Low vs Mixed | Adjusted | HIV.DURATION | 2.647 (95% CI 1.946–3.602), p=0.000 | 2.499 (SE 0.072), p=0.000 | 1.354 (95% CI 1.162–1.576), p=0.000 | 1.270 (95% CI 0.794–2.031), p=0.318 | 0.756 (log-odds), OR=2.131 (95% CI 1.452–3.126) | Boot CI [0.403, 1.148] | 77.7% | 0.147 (95% CI 0.078–0.217), p=0.000 | TRUE | 0.046 (95% CI -0.044–0.134), p=0.321 | FALSE | 0.193 (95% CI 0.134–0.251), p=0.000 | 0.761507506 | Significant complete mediation |
| Mixed vs All High | Unadjusted | HIV.DURATION | 1.296 (95% CI 0.961–1.748), p=0.090 | 0.668 (SE 0.074), p=0.000 | 1.238 (95% CI 1.066–1.438), p=0.005 | 1.124 (95% CI 0.818–1.543), p=0.471 | 0.143 (log-odds), OR=1.153 (95% CI 1.039–1.281) | Boot CI [0.039, 0.264] | 55.1% | 0.034 (95% CI 0.009–0.061), p=0.007 | TRUE | 0.028 (95% CI -0.053–0.105), p=0.488 | FALSE | 0.062 (95% CI -0.014–0.134), p=0.107 | 0.549945501 | Significant complete mediation |
| Mixed vs All High | Adjusted | HIV.DURATION | 1.226 (95% CI 0.905–1.660), p=0.188 | 0.684 (SE 0.074), p=0.000 | 1.264 (95% CI 1.086–1.471), p=0.002 | 1.047 (95% CI 0.759–1.444), p=0.779 | 0.160 (log-odds), OR=1.174 (95% CI 1.052–1.309) | Boot CI [0.056, 0.282] | 78.5% | 0.038 (95% CI 0.013–0.065), p=0.002 | TRUE | 0.011 (95% CI -0.064–0.089), p=0.778 | FALSE | 0.048 (95% CI -0.023–0.124), p=0.188 | 0.776905298 | Significant complete mediation |
| All Low vs All High | Unadjusted | HIV.DURATION | 3.116 (95% CI 2.231–4.351), p=0.000 | 3.214 (SE 0.094), p=0.000 | 1.263 (95% CI 1.089–1.464), p=0.002 | 1.503 (95% CI 0.857–2.637), p=0.155 | 0.750 (log-odds), OR=2.116 (95% CI 1.312–3.414) | Boot CI [0.298, 1.253] | 66.0% | 0.155 (95% CI 0.058–0.253), p=0.001 | TRUE | 0.084 (95% CI -0.030–0.201), p=0.159 | FALSE | 0.239 (95% CI 0.171–0.307), p=0.000 | 0.649758549 | Significant complete mediation |
| All Low vs All High | Adjusted | HIV.DURATION | 3.268 (95% CI 2.328–4.588), p=0.000 | 3.191 (SE 0.092), p=0.000 | 1.350 (95% CI 1.156–1.577), p=0.000 | 1.292 (95% CI 0.726–2.299), p=0.384 | 0.958 (log-odds), OR=2.607 (95% CI 1.584–4.293) | Boot CI [0.503, 1.489] | 80.9% | 0.194 (95% CI 0.100–0.286), p=0.000 | TRUE | 0.051 (95% CI -0.061–0.165), p=0.404 | FALSE | 0.245 (95% CI 0.175–0.311), p=0.000 | 0.792366011 | Significant complete mediation |
| Residual Viremia (2 Years Follow-Up) | | | | | | | | | | | | | | | | |
| All Low vs Mixed | Unadjusted | CMV.IgG.IUpermL, HIV.DURATION | 2.234 (95% CI 1.607–3.106), p=0.000 | 2.546 (SE 0.071), p=0.000 | 1.421 (95% CI 1.202–1.682), p=0.000 | 0.954 (95% CI 0.573–1.590), p=0.858 | 0.895 (log-odds), OR=2.448 (95% CI 1.591–3.766) | Boot CI [0.470, 1.361] | 111.4% (>100%) | 0.144 (95% CI 0.078–0.214), p=0.000 | TRUE | -0.008 (95% CI -0.088–0.075), p=0.859 | FALSE | 0.137 (95% CI 0.083–0.190), p=0.000 | 1.054897308 | Significant complete mediation |
| All Low vs Mixed | Adjusted | CMV.IgG.IUpermL, HIV.DURATION | 2.660 (95% CI 1.861–3.802), p=0.000 | 2.489 (SE 0.073), p=0.000 | 1.409 (95% CI 1.176–1.688), p=0.000 | 1.168 (95% CI 0.676–2.019), p=0.577 | 0.853 (log-odds), OR=2.348 (95% CI 1.494–3.689) | Boot CI [0.422, 1.356] | 87.2% | 0.138 (95% CI 0.066–0.213), p=0.000 | TRUE | 0.025 (95% CI -0.066–0.110), p=0.608 | FALSE | 0.163 (95% CI 0.107–0.214), p=0.000 | 0.845970941 | Significant complete mediation |
| Mixed vs All High | Unadjusted | CMV.IgG.IUpermL, HIV.DURATION | 1.389 (95% CI 1.014–1.902), p=0.040 | 0.668 (SE 0.074), p=0.000 | 1.489 (95% CI 1.265–1.754), p=0.000 | 1.060 (95% CI 0.755–1.488), p=0.737 | 0.266 (log-odds), OR=1.305 (95% CI 1.153–1.477) | Boot CI [0.150, 0.412] | 81.0% | 0.056 (95% CI 0.032–0.083), p=0.000 | TRUE | 0.012 (95% CI -0.061–0.084), p=0.759 | FALSE | 0.068 (95% CI -0.003–0.140), p=0.058 | 0.820856005 | Significant complete mediation |
| Mixed vs All High | Adjusted | CMV.IgG.IUpermL, HIV.DURATION | 1.340 (95% CI 0.970–1.853), p=0.076 | 0.704 (SE 0.076), p=0.000 | 1.536 (95% CI 1.294–1.823), p=0.000 | 0.987 (95% CI 0.694–1.404), p=0.942 | 0.302 (log-odds), OR=1.353 (95% CI 1.180–1.550) | Boot CI [0.177, 0.453] | 103.1% (>100%) | 0.064 (95% CI 0.038–0.095), p=0.000 | TRUE | -0.003 (95% CI -0.077–0.070), p=0.914 | FALSE | 0.061 (95% CI -0.012–0.133), p=0.104 | 1.045492082 | Significant complete mediation |
| All Low vs All High | Unadjusted | CMV.IgG.IUpermL, HIV.DURATION | 3.103 (95% CI 2.165–4.446), p=0.000 | 3.214 (SE 0.094), p=0.000 | 1.457 (95% CI 1.230–1.727), p=0.000 | 0.970 (95% CI 0.520–1.809), p=0.923 | 1.211 (log-odds), OR=3.355 (95% CI 1.935–5.818) | Boot CI [0.680, 1.836] | 106.9% (>100%) | 0.212 (95% CI 0.118–0.303), p=0.000 | TRUE | -0.005 (95% CI -0.109–0.105), p=0.919 | FALSE | 0.207 (95% CI 0.141–0.271), p=0.000 | 1.025767201 | Significant complete mediation |
| All Low vs All High | Adjusted | CMV.IgG.IUpermL, HIV.DURATION | 3.484 (95% CI 2.374–5.112), p=0.000 | 3.191 (SE 0.095), p=0.000 | 1.466 (95% CI 1.222–1.760), p=0.000 | 1.073 (95% CI 0.551–2.090), p=0.837 | 1.221 (log-odds), OR=3.391 (95% CI 1.886–6.096) | Boot CI [0.670, 1.903] | 97.8% | 0.209 (95% CI 0.113–0.308), p=0.000 | TRUE | 0.012 (95% CI -0.099–0.124), p=0.842 | FALSE | 0.221 (95% CI 0.156–0.290), p=0.000 | 0.946321621 | Significant complete mediation |
| Post-cART Opportunistic Infections | | | | | | | | | | | | | | | | |
| All Low vs Mixed | Unadjusted | HIV.DURATION, CMV.IgG.IUpermL | 1.997 (95% CI 1.297–3.074), p=0.002 | 2.546 (SE 0.071), p=0.000 | 1.290 (95% CI 1.044–1.594), p=0.018 | 1.068 (95% CI 0.551–2.073), p=0.845 | 0.649 (log-odds), OR=1.913 (95% CI 1.115–3.281) | Boot CI [0.101, 1.258] | 93.8% | 0.063 (95% CI 0.007–0.123), p=0.024 | TRUE | 0.006 (95% CI -0.062–0.072), p=0.868 | FALSE | 0.069 (95% CI 0.026–0.110), p=0.001 | 0.906323119 | Significant complete mediation |
| All Low vs Mixed | Adjusted | HIV.DURATION, CMV.IgG.IUpermL | 1.612 (95% CI 1.028–2.527), p=0.037 | 2.489 (SE 0.073), p=0.000 | 1.314 (95% CI 1.046–1.651), p=0.019 | 0.851 (95% CI 0.428–1.691), p=0.645 | 0.680 (log-odds), OR=1.975 (95% CI 1.117–3.490) | Boot CI [0.099, 1.351] | 142.5% (>100%) | 0.067 (95% CI 0.009–0.134), p=0.020 | TRUE | -0.016 (95% CI -0.087–0.052), p=0.664 | FALSE | 0.051 (95% CI 0.008–0.096), p=0.023 | 1.315461533 | Significant complete mediation |
| Mixed vs All High | Unadjusted | HIV.DURATION, CMV.IgG.IUpermL | 0.911 (95% CI 0.596–1.393), p=0.668 | 0.668 (SE 0.074), p=0.000 | 1.220 (95% CI 0.991–1.502), p=0.061 | 0.791 (95% CI 0.503–1.245), p=0.312 | 0.133 (log-odds), OR=1.142 (95% CI 0.991–1.316) | Boot CI [-0.015, 0.297] | Suppression / inconsistent mediation (opposite signs) | 0.016 (95% CI -0.002–0.035), p=0.074 | FALSE | -0.028 (95% CI -0.084–0.027), p=0.300 | FALSE | -0.012 (95% CI -0.063–0.038), p=0.621 | -1.316731854 | No significant mediation (ACME not significant) |
| Mixed vs All High | Adjusted | HIV.DURATION, CMV.IgG.IUpermL | 0.948 (95% CI 0.608–1.478), p=0.814 | 0.704 (SE 0.076), p=0.000 | 1.208 (95% CI 0.968–1.508), p=0.094 | 0.816 (95% CI 0.504–1.321), p=0.409 | 0.133 (log-odds), OR=1.142 (95% CI 0.975–1.339) | Boot CI [-0.025, 0.319] | -249.1% | 0.016 (95% CI -0.004–0.037), p=0.115 | FALSE | -0.024 (95% CI -0.085–0.036), p=0.436 | FALSE | -0.008 (95% CI -0.061–0.046), p=0.755 | -1.909172887 | No significant mediation (ACME not significant) |
| All Low vs All High | Unadjusted | HIV.DURATION, CMV.IgG.IUpermL | 1.820 (95% CI 1.115–2.969), p=0.017 | 3.214 (SE 0.094), p=0.000 | 1.129 (95% CI 0.914–1.393), p=0.260 | 1.236 (95% CI 0.540–2.832), p=0.616 | 0.389 (log-odds), OR=1.476 (95% CI 0.750–2.904) | Boot CI [-0.305, 1.159] | 65.0% | 0.037 (95% CI -0.028–0.111), p=0.250 | FALSE | 0.020 (95% CI -0.069–0.101), p=0.642 | FALSE | 0.057 (95% CI 0.008–0.105), p=0.020 | 0.646249120 | No significant mediation (ACME not significant) |
| All Low vs All High | Adjusted | HIV.DURATION, CMV.IgG.IUpermL | 1.529 (95% CI 0.914–2.560), p=0.106 | 3.191 (SE 0.095), p=0.000 | 1.012 (95% CI 0.808–1.267), p=0.917 | 1.473 (95% CI 0.613–3.540), p=0.387 | 0.038 (log-odds), OR=1.039 (95% CI 0.507–2.131) | Boot CI [-0.684, 0.851] | 9.0% | 0.003 (95% CI -0.064–0.079), p=0.922 | FALSE | 0.035 (95% CI -0.057–0.121), p=0.454 | FALSE | 0.039 (95% CI -0.008–0.087), p=0.104 | 0.090181962 | No significant mediation (ACME not significant) |
| AIDS-Defining Malignancies | | | | | | | | | | | | | | | | |
| All Low vs Mixed | Unadjusted | HIV.DURATION | 2.113 (95% CI 1.195–3.737), p=0.010 | 2.546 (SE 0.071), p=0.000 | 1.722 (95% CI 1.288–2.302), p=0.000 | 0.578 (95% CI 0.240–1.388), p=0.220 | 1.384 (log-odds), OR=3.990 (95% CI 1.898–8.384) | Boot CI [0.515, 2.349] | 185.0% (>100%) | 0.084 (95% CI 0.027–0.155), p=0.002 | TRUE | -0.035 (95% CI -0.111–0.024), p=0.286 | FALSE | 0.048 (95% CI 0.017–0.079), p=0.004 | 1.736309136 | Significant complete mediation |
| All Low vs Mixed | Adjusted | HIV.DURATION | 2.026 (95% CI 1.138–3.607), p=0.016 | 2.499 (SE 0.072), p=0.000 | 1.713 (95% CI 1.280–2.293), p=0.000 | 0.574 (95% CI 0.239–1.377), p=0.213 | 1.345 (log-odds), OR=3.840 (95% CI 1.846–7.986) | Boot CI [0.454, 2.308] | 190.5% (>100%) | 0.081 (95% CI 0.022–0.149), p=0.004 | TRUE | -0.036 (95% CI -0.108–0.025), p=0.290 | FALSE | 0.046 (95% CI 0.013–0.078), p=0.005 | 1.783762490 | Significant complete mediation |
| Mixed vs All High | Unadjusted | HIV.DURATION | 0.965 (95% CI 0.566–1.644), p=0.895 | 0.668 (SE 0.074), p=0.000 | 1.462 (95% CI 1.126–1.898), p=0.004 | 0.726 (95% CI 0.407–1.294), p=0.277 | 0.254 (log-odds), OR=1.289 (95% CI 1.073–1.547) | Boot CI [0.069, 0.462] | Suppression / inconsistent mediation (opposite signs) | 0.019 (95% CI 0.005–0.034), p=0.005 | TRUE | -0.024 (95% CI -0.069–0.022), p=0.298 | FALSE | -0.005 (95% CI -0.046–0.037), p=0.796 | -3.831374733 | Significant complete mediation |
| Mixed vs All High | Adjusted | HIV.DURATION | 0.952 (95% CI 0.557–1.629), p=0.859 | 0.684 (SE 0.074), p=0.000 | 1.465 (95% CI 1.129–1.901), p=0.004 | 0.715 (95% CI 0.400–1.277), p=0.256 | 0.261 (log-odds), OR=1.298 (95% CI 1.077–1.565) | Boot CI [0.077, 0.464] | -534.6% | 0.020 (95% CI 0.005–0.035), p=0.004 | TRUE | -0.025 (95% CI -0.070–0.020), p=0.270 | FALSE | -0.006 (95% CI -0.046–0.037), p=0.782 | -3.512066861 | Significant complete mediation |
| All Low vs All High | Unadjusted | HIV.DURATION | 2.038 (95% CI 1.079–3.851), p=0.028 | 3.214 (SE 0.094), p=0.000 | 1.339 (95% CI 1.005–1.785), p=0.046 | 0.806 (95% CI 0.266–2.441), p=0.702 | 0.939 (log-odds), OR=2.558 (95% CI 1.014–6.453) | Boot CI [-0.047, 2.094] | 131.9% (>100%) | 0.054 (95% CI -0.004–0.138), p=0.067 | FALSE | -0.013 (95% CI -0.108–0.057), p=0.715 | FALSE | 0.041 (95% CI 0.005–0.078), p=0.024 | 1.315597877 | No significant mediation (ACME not significant) |
| All Low vs All High | Adjusted | HIV.DURATION | 1.995 (95% CI 1.054–3.775), p=0.034 | 3.191 (SE 0.092), p=0.000 | 1.304 (95% CI 0.973–1.747), p=0.076 | 0.868 (95% CI 0.284–2.648), p=0.803 | 0.846 (log-odds), OR=2.330 (95% CI 0.914–5.939) | Boot CI [-0.148, 1.985] | 122.5% (>100%) | 0.047 (95% CI -0.008–0.132), p=0.095 | FALSE | -0.008 (95% CI -0.102–0.060), p=0.812 | FALSE | 0.039 (95% CI 0.002–0.078), p=0.038 | 1.210501679 | No significant mediation (ACME not significant) |
| Endocrine/Metabolic Disorders | | | | | | | | | | | | | | | | |
| All Low vs Mixed | Unadjusted | AGE, HIV.DURATION, CMV.IgG.IUpermL | 1.487 (95% CI 1.108–1.996), p=0.008 | 2.546 (SE 0.071), p=0.000 | 1.104 (95% CI 0.960–1.271), p=0.165 | 1.158 (95% CI 0.733–1.829), p=0.530 | 0.253 (log-odds), OR=1.288 (95% CI 0.901–1.842) | Boot CI [-0.104, 0.599] | 63.8% | 0.050 (95% CI -0.018–0.120), p=0.148 | FALSE | 0.029 (95% CI -0.063–0.124), p=0.566 | FALSE | 0.079 (95% CI 0.021–0.138), p=0.006 | 0.633591579 | No significant mediation (ACME not significant) |
| All Low vs Mixed | Adjusted | AGE, HIV.DURATION, CMV.IgG.IUpermL | 1.115 (95% CI 0.796–1.560), p=0.527 | 2.481 (SE 0.074), p=0.000 | 0.990 (95% CI 0.843–1.164), p=0.907 | 1.141 (95% CI 0.683–1.906), p=0.615 | -0.024 (log-odds), OR=0.976 (95% CI 0.654–1.458) | Boot CI [-0.427, 0.412] | -22.0% | -0.004 (95% CI -0.071–0.065), p=0.899 | FALSE | 0.022 (95% CI -0.068–0.114), p=0.611 | FALSE | 0.018 (95% CI -0.038–0.076), p=0.520 | -0.221748775 | No significant mediation (ACME not significant) |
| Mixed vs All High | Unadjusted | AGE, HIV.DURATION, CMV.IgG.IUpermL | 0.998 (95% CI 0.727–1.370), p=0.991 | 0.668 (SE 0.074), p=0.000 | 1.001 (95% CI 0.857–1.170), p=0.986 | 0.997 (95% CI 0.714–1.392), p=0.987 | 0.001 (log-odds), OR=1.001 (95% CI 0.902–1.111) | Boot CI [-0.104, 0.101] | Suppression / inconsistent mediation (opposite signs) | 0.000 (95% CI -0.022–0.022), p=0.999 | FALSE | -0.001 (95% CI -0.071–0.068), p=0.990 | FALSE | -0.000 (95% CI -0.070–0.068), p=0.982 | -0.481710283 | No significant mediation (ACME not significant) |
| Mixed vs All High | Adjusted | AGE, HIV.DURATION, CMV.IgG.IUpermL | 0.956 (95% CI 0.666–1.372), p=0.808 | 0.704 (SE 0.076), p=0.000 | 0.925 (95% CI 0.772–1.108), p=0.396 | 1.013 (95% CI 0.689–1.488), p=0.949 | -0.055 (log-odds), OR=0.946 (95% CI 0.833–1.075) | Boot CI [-0.190, 0.075] | 122.6% (>100%) | -0.010 (95% CI -0.034–0.013), p=0.384 | FALSE | 0.002 (95% CI -0.066–0.071), p=0.973 | FALSE | -0.008 (95% CI -0.072–0.056), p=0.790 | 1.296465298 | No significant mediation (ACME not significant) |
| All Low vs All High | Unadjusted | AGE, HIV.DURATION, CMV.IgG.IUpermL | 1.484 (95% CI 1.060–2.078), p=0.022 | 3.214 (SE 0.094), p=0.000 | 1.378 (95% CI 1.182–1.605), p=0.000 | 0.540 (95% CI 0.301–0.968), p=0.038 | 1.030 (log-odds), OR=2.801 (95% CI 1.708–4.592) | Boot CI [0.559, 1.553] | 260.9% (>100%) | 0.194 (95% CI 0.111–0.277), p=0.000 | TRUE | -0.115 (95% CI -0.213–-0.011), p=0.032 | TRUE | 0.079 (95% CI 0.011–0.149), p=0.024 | 2.454422968 | Significant partial mediation |
| All Low vs All High | Adjusted | AGE, HIV.DURATION, CMV.IgG.IUpermL | 1.086 (95% CI 0.739–1.597), p=0.675 | 3.169 (SE 0.095), p=0.000 | 1.222 (95% CI 1.030–1.451), p=0.021 | 0.580 (95% CI 0.300–1.120), p=0.105 | 0.636 (log-odds), OR=1.890 (95% CI 1.097–3.254) | Boot CI [0.102, 1.211] | 771.7% (>100%) | 0.102 (95% CI 0.015–0.181), p=0.022 | TRUE | -0.087 (95% CI -0.186–0.016), p=0.105 | FALSE | 0.015 (95% CI -0.048–0.076), p=0.606 | 6.937906373 | Significant complete mediation |
| Hypercholestoremia | | | | | | | | | | | | | | | | |
| All Low vs Mixed | Unadjusted | AGE, HIV.DURATION | 1.369 (95% CI 1.010–1.856), p=0.043 | 2.546 (SE 0.071), p=0.000 | 1.052 (95% CI 0.911–1.215), p=0.489 | 1.204 (95% CI 0.750–1.931), p=0.442 | 0.129 (log-odds), OR=1.138 (95% CI 0.789–1.640) | Boot CI [-0.213, 0.480] | 41.2% | 0.024 (95% CI -0.040–0.089), p=0.467 | FALSE | 0.034 (95% CI -0.055–0.123), p=0.469 | FALSE | 0.058 (95% CI 0.004–0.113), p=0.034 | 0.410543771 | No significant mediation (ACME not significant) |
| All Low vs Mixed | Adjusted | AGE, HIV.DURATION | 1.065 (95% CI 0.761–1.491), p=0.713 | 2.482 (SE 0.072), p=0.000 | 0.946 (95% CI 0.805–1.110), p=0.495 | 1.217 (95% CI 0.731–2.025), p=0.451 | -0.139 (log-odds), OR=0.870 (95% CI 0.584–1.296) | Boot CI [-0.536, 0.269] | -219.7% | -0.021 (95% CI -0.083–0.041), p=0.494 | FALSE | 0.030 (95% CI -0.053–0.115), p=0.470 | FALSE | 0.009 (95% CI -0.044–0.063), p=0.724 | -2.424300125 | No significant mediation (ACME not significant) |
| Mixed vs All High | Unadjusted | AGE, HIV.DURATION | 0.980 (95% CI 0.704–1.364), p=0.907 | 0.668 (SE 0.074), p=0.000 | 0.995 (95% CI 0.846–1.170), p=0.954 | 0.984 (95% CI 0.695–1.393), p=0.925 | -0.003 (log-odds), OR=0.997 (95% CI 0.894–1.111) | Boot CI [-0.112, 0.097] | 16.2% | -0.001 (95% CI -0.021–0.020), p=0.975 | FALSE | -0.003 (95% CI -0.070–0.061), p=0.899 | FALSE | -0.004 (95% CI -0.070–0.059), p=0.900 | 0.162125107 | No significant mediation (ACME not significant) |
| Mixed vs All High | Adjusted | AGE, HIV.DURATION | 1.006 (95% CI 0.698–1.452), p=0.973 | 0.681 (SE 0.074), p=0.000 | 0.929 (95% CI 0.774–1.115), p=0.428 | 1.063 (95% CI 0.720–1.572), p=0.758 | -0.050 (log-odds), OR=0.951 (95% CI 0.839–1.077) | Boot CI [-0.186, 0.074] | -793.7% | -0.008 (95% CI -0.030–0.012), p=0.441 | FALSE | 0.010 (95% CI -0.053–0.074), p=0.740 | FALSE | 0.002 (95% CI -0.058–0.063), p=0.939 | -4.499442224 | No significant mediation (ACME not significant) |
| All Low vs All High | Unadjusted | AGE, HIV.DURATION | 1.342 (95% CI 0.946–1.903), p=0.099 | 3.214 (SE 0.094), p=0.000 | 1.330 (95% CI 1.138–1.555), p=0.000 | 0.543 (95% CI 0.297–0.992), p=0.047 | 0.918 (log-odds), OR=2.503 (95% CI 1.511–4.146) | Boot CI [0.450, 1.443] | 312.0% (>100%) | 0.163 (95% CI 0.080–0.245), p=0.000 | TRUE | -0.108 (95% CI -0.209–-0.005), p=0.044 | TRUE | 0.055 (95% CI -0.009–0.122), p=0.090 | 2.968413946 | Significant partial mediation |
| All Low vs All High | Adjusted | AGE, HIV.DURATION | 1.083 (95% CI 0.736–1.595), p=0.686 | 3.156 (SE 0.092), p=0.000 | 1.171 (95% CI 0.987–1.389), p=0.070 | 0.662 (95% CI 0.343–1.277), p=0.219 | 0.498 (log-odds), OR=1.646 (95% CI 0.960–2.824) | Boot CI [-0.035, 1.066] | 624.6% (>100%) | 0.073 (95% CI -0.005–0.151), p=0.066 | FALSE | -0.061 (95% CI -0.156–0.035), p=0.204 | FALSE | 0.013 (95% CI -0.044–0.070), p=0.663 | 5.769645664 | No significant mediation (ACME not significant) |
| Gastroenterology Disease | | | | | | | | | | | | | | | | |
| All Low vs Mixed | Unadjusted | HIV.DURATION | 1.552 (95% CI 1.131–2.129), p=0.006 | 2.546 (SE 0.071), p=0.000 | 0.995 (95% CI 0.858–1.153), p=0.943 | 1.573 (95% CI 0.962–2.574), p=0.071 | -0.014 (log-odds), OR=0.986 (95% CI 0.677–1.437) | Boot CI [-0.361, 0.350] | Suppression / inconsistent mediation (opposite signs) | -0.002 (95% CI -0.063–0.061), p=0.952 | FALSE | 0.078 (95% CI -0.006–0.162), p=0.068 | FALSE | 0.076 (95% CI 0.023–0.129), p=0.005 | -0.031007682 | No significant mediation (ACME not significant) |
| All Low vs Mixed | Adjusted | HIV.DURATION | 1.399 (95% CI 1.013–1.931), p=0.042 | 2.499 (SE 0.072), p=0.000 | 0.957 (95% CI 0.823–1.113), p=0.571 | 1.557 (95% CI 0.951–2.551), p=0.079 | -0.109 (log-odds), OR=0.897 (95% CI 0.615–1.307) | Boot CI [-0.458, 0.256] | -32.4% | -0.018 (95% CI -0.077–0.041), p=0.550 | FALSE | 0.075 (95% CI -0.006–0.154), p=0.077 | FALSE | 0.057 (95% CI 0.002–0.113), p=0.044 | -0.325338805 | No significant mediation (ACME not significant) |
| Mixed vs All High | Unadjusted | HIV.DURATION | 0.882 (95% CI 0.627–1.240), p=0.470 | 0.668 (SE 0.074), p=0.000 | 0.894 (95% CI 0.756–1.057), p=0.191 | 0.948 (95% CI 0.663–1.356), p=0.771 | -0.075 (log-odds), OR=0.928 (95% CI 0.829–1.039) | Boot CI [-0.188, 0.031] | 59.4% | -0.014 (95% CI -0.035–0.006), p=0.172 | FALSE | -0.010 (95% CI -0.077–0.057), p=0.745 | FALSE | -0.024 (95% CI -0.088–0.040), p=0.438 | 0.584395746 | No significant mediation (ACME not significant) |
| Mixed vs All High | Adjusted | HIV.DURATION | 0.941 (95% CI 0.666–1.330), p=0.730 | 0.684 (SE 0.074), p=0.000 | 0.871 (95% CI 0.734–1.034), p=0.115 | 1.034 (95% CI 0.718–1.490), p=0.856 | -0.094 (log-odds), OR=0.910 (95% CI 0.808–1.025) | Boot CI [-0.213, 0.020] | 154.7% (>100%) | -0.017 (95% CI -0.039–0.003), p=0.102 | FALSE | 0.006 (95% CI -0.061–0.073), p=0.854 | FALSE | -0.011 (95% CI -0.074–0.052), p=0.729 | 1.558044327 | No significant mediation (ACME not significant) |
| All Low vs All High | Unadjusted | HIV.DURATION | 1.368 (95% CI 0.948–1.974), p=0.094 | 3.214 (SE 0.094), p=0.000 | 0.983 (95% CI 0.847–1.142), p=0.824 | 1.445 (95% CI 0.789–2.645), p=0.233 | -0.054 (log-odds), OR=0.947 (95% CI 0.586–1.530) | Boot CI [-0.490, 0.413] | Suppression / inconsistent mediation (opposite signs) | -0.009 (95% CI -0.085–0.068), p=0.807 | FALSE | 0.061 (95% CI -0.038–0.161), p=0.238 | FALSE | 0.052 (95% CI -0.010–0.115), p=0.096 | -0.173233280 | No significant mediation (ACME not significant) |
| All Low vs All High | Adjusted | HIV.DURATION | 1.323 (95% CI 0.911–1.920), p=0.141 | 3.191 (SE 0.092), p=0.000 | 0.915 (95% CI 0.784–1.067), p=0.257 | 1.755 (95% CI 0.948–3.251), p=0.074 | -0.285 (log-odds), OR=0.752 (95% CI 0.460–1.231) | Boot CI [-0.740, 0.194] | -101.8% | -0.046 (95% CI -0.117–0.033), p=0.253 | FALSE | 0.091 (95% CI -0.010–0.193), p=0.083 | FALSE | 0.045 (95% CI -0.017–0.109), p=0.152 | -1.024777166 | No significant mediation (ACME not significant) |
| Musculoskeletal Disorder | | | | | | | | | | | | | | | | |
| All Low vs Mixed | Unadjusted | AGE, HIV.DURATION | 1.614 (95% CI 1.053–2.475), p=0.028 | 2.546 (SE 0.071), p=0.000 | 1.038 (95% CI 0.848–1.269), p=0.720 | 1.470 (95% CI 0.757–2.854), p=0.255 | 0.094 (log-odds), OR=1.098 (95% CI 0.658–1.834) | Boot CI [-0.359, 0.590] | 19.6% | 0.009 (95% CI -0.036–0.055), p=0.710 | FALSE | 0.037 (95% CI -0.026–0.100), p=0.237 | FALSE | 0.046 (95% CI 0.006–0.087), p=0.023 | 0.196630292 | No significant mediation (ACME not significant) |
| All Low vs Mixed | Adjusted | AGE, HIV.DURATION | 1.353 (95% CI 0.870–2.102), p=0.180 | 2.482 (SE 0.072), p=0.000 | 0.967 (95% CI 0.783–1.193), p=0.753 | 1.465 (95% CI 0.753–2.851), p=0.261 | -0.084 (log-odds), OR=0.919 (95% CI 0.545–1.551) | Boot CI [-0.579, 0.415] | -27.9% | -0.008 (95% CI -0.054–0.038), p=0.764 | FALSE | 0.035 (95% CI -0.023–0.095), p=0.242 | FALSE | 0.027 (95% CI -0.014–0.069), p=0.192 | -0.283805449 | No significant mediation (ACME not significant) |
| Mixed vs All High | Unadjusted | AGE, HIV.DURATION | 1.188 (95% CI 0.782–1.804), p=0.420 | 0.668 (SE 0.074), p=0.000 | 1.103 (95% CI 0.897–1.356), p=0.351 | 1.110 (95% CI 0.713–1.729), p=0.643 | 0.066 (log-odds), OR=1.068 (95% CI 0.930–1.227) | Boot CI [-0.069, 0.208] | 38.2% | 0.008 (95% CI -0.009–0.026), p=0.344 | FALSE | 0.013 (95% CI -0.040–0.065), p=0.643 | FALSE | 0.021 (95% CI -0.031–0.071), p=0.452 | 0.385354037 | No significant mediation (ACME not significant) |
| Mixed vs All High | Adjusted | AGE, HIV.DURATION | 1.250 (95% CI 0.812–1.924), p=0.311 | 0.681 (SE 0.074), p=0.000 | 1.061 (95% CI 0.853–1.318), p=0.596 | 1.194 (95% CI 0.751–1.899), p=0.454 | 0.040 (log-odds), OR=1.041 (95% CI 0.897–1.207) | Boot CI [-0.112, 0.192] | 18.0% | 0.005 (95% CI -0.014–0.023), p=0.599 | FALSE | 0.021 (95% CI -0.035–0.078), p=0.466 | FALSE | 0.025 (95% CI -0.026–0.080), p=0.333 | 0.184404037 | No significant mediation (ACME not significant) |
| All Low vs All High | Unadjusted | AGE, HIV.DURATION | 1.917 (95% CI 1.201–3.062), p=0.006 | 3.214 (SE 0.094), p=0.000 | 1.168 (95% CI 0.953–1.432), p=0.134 | 1.169 (95% CI 0.527–2.590), p=0.701 | 0.500 (log-odds), OR=1.648 (95% CI 0.856–3.172) | Boot CI [-0.071, 1.152] | 76.8% | 0.051 (95% CI -0.009–0.122), p=0.095 | FALSE | 0.016 (95% CI -0.063–0.091), p=0.686 | FALSE | 0.067 (95% CI 0.019–0.118), p=0.008 | 0.761117707 | No significant mediation (ACME not significant) |
| All Low vs All High | Adjusted | AGE, HIV.DURATION | 1.694 (95% CI 1.037–2.766), p=0.035 | 3.156 (SE 0.092), p=0.000 | 1.005 (95% CI 0.810–1.247), p=0.967 | 1.670 (95% CI 0.729–3.827), p=0.226 | 0.014 (log-odds), OR=1.015 (95% CI 0.514–2.004) | Boot CI [-0.666, 0.700] | 2.7% | 0.001 (95% CI -0.061–0.067), p=0.954 | FALSE | 0.048 (95% CI -0.029–0.123), p=0.236 | FALSE | 0.049 (95% CI 0.004–0.096), p=0.035 | 0.027594539 | No significant mediation (ACME not significant) |
| Stroke | | | | | | | | | | | | | | | | |
| All Low vs Mixed | Unadjusted | AGE, HIV.DURATION, SMOKING.CURRENT | 1.049 (95% CI 0.465–2.366), p=0.909 | 2.546 (SE 0.071), p=0.000 | 1.197 (95% CI 0.803–1.783), p=0.377 | 0.671 (95% CI 0.189–2.379), p=0.537 | 0.457 (log-odds), OR=1.580 (95% CI 0.572–4.363) | Boot CI [-0.510, 1.543] | 965.0% (>100%) | 0.012 (95% CI -0.013–0.052), p=0.328 | FALSE | -0.010 (95% CI -0.057–0.025), p=0.547 | FALSE | 0.001 (95% CI -0.018–0.022), p=0.844 | 7.987527257 | No significant mediation (ACME not significant) |
| All Low vs Mixed | Adjusted | AGE, HIV.DURATION, SMOKING.CURRENT | 0.938 (95% CI 0.398–2.209), p=0.884 | 2.502 (SE 0.076), p=0.000 | 1.081 (95% CI 0.710–1.647), p=0.715 | 0.779 (95% CI 0.210–2.888), p=0.708 | 0.196 (log-odds), OR=1.216 (95% CI 0.425–3.483) | Boot CI [-0.842, 1.396] | -306.3% | 0.005 (95% CI -0.022–0.040), p=0.718 | FALSE | -0.007 (95% CI -0.048–0.033), p=0.748 | FALSE | -0.001 (95% CI -0.023–0.022), p=0.976 | -3.615858596 | No significant mediation (ACME not significant) |
| Mixed vs All High | Unadjusted | AGE, HIV.DURATION, SMOKING.CURRENT | 2.191 (95% CI 1.038–4.626), p=0.040 | 0.668 (SE 0.074), p=0.000 | 1.180 (95% CI 0.822–1.694), p=0.369 | 1.950 (95% CI 0.882–4.313), p=0.099 | 0.111 (log-odds), OR=1.117 (95% CI 0.876–1.424) | Boot CI [-0.108, 0.324] | 14.1% | 0.004 (95% CI -0.004–0.013), p=0.305 | FALSE | 0.026 (95% CI -0.004–0.057), p=0.092 | FALSE | 0.030 (95% CI 0.000–0.062), p=0.049 | 0.145414323 | No significant mediation (ACME not significant) |
| Mixed vs All High | Adjusted | AGE, HIV.DURATION, SMOKING.CURRENT | 1.896 (95% CI 0.855–4.204), p=0.116 | 0.661 (SE 0.079), p=0.000 | 1.043 (95% CI 0.700–1.554), p=0.837 | 1.834 (95% CI 0.779–4.321), p=0.165 | 0.028 (log-odds), OR=1.028 (95% CI 0.790–1.338) | Boot CI [-0.242, 0.275] | 4.3% | 0.001 (95% CI -0.010–0.010), p=0.809 | FALSE | 0.023 (95% CI -0.010–0.056), p=0.171 | FALSE | 0.024 (95% CI -0.008–0.057), p=0.140 | 0.044454639 | No significant mediation (ACME not significant) |
| All Low vs All High | Unadjusted | AGE, HIV.DURATION, SMOKING.CURRENT | 2.298 (95% CI 1.050–5.027), p=0.037 | 3.214 (SE 0.094), p=0.000 | 1.330 (95% CI 0.938–1.884), p=0.109 | 0.929 (95% CI 0.240–3.595), p=0.914 | 0.915 (log-odds), OR=2.498 (95% CI 0.814–7.668) | Boot CI [0.063, 1.906] | 110.0% (>100%) | 0.035 (95% CI 0.002–0.088), p=0.036 | TRUE | -0.003 (95% CI -0.066–0.041), p=0.900 | FALSE | 0.032 (95% CI 0.001–0.064), p=0.040 | 1.093046348 | Significant complete mediation |
| All Low vs All High | Adjusted | AGE, HIV.DURATION, SMOKING.CURRENT | 1.740 (95% CI 0.736–4.114), p=0.207 | 3.158 (SE 0.098), p=0.000 | 1.076 (95% CI 0.736–1.573), p=0.706 | 1.383 (95% CI 0.318–6.017), p=0.665 | 0.231 (log-odds), OR=1.259 (95% CI 0.379–4.180) | Boot CI [-0.793, 1.386] | 41.6% | 0.008 (95% CI -0.025–0.051), p=0.628 | FALSE | 0.011 (95% CI -0.041–0.056), p=0.646 | FALSE | 0.019 (95% CI -0.011–0.051), p=0.221 | 0.416040709 | No significant mediation (ACME not significant) |
| AIDS-Defining Opportunistic Infections | | | | | | | | | | | | | | | | |
| All Low vs Mixed | Unadjusted | HIV.DURATION | 1.442 (95% CI 0.943–2.205), p=0.091 | 2.546 (SE 0.071), p=0.000 | 1.617 (95% CI 1.291–2.025), p=0.000 | 0.453 (95% CI 0.231–0.889), p=0.021 | 1.223 (log-odds), OR=3.399 (95% CI 1.910–6.048) | Boot CI [0.643, 1.864] | 334.4% (>100%) | 0.122 (95% CI 0.062–0.187), p=0.000 | TRUE | -0.080 (95% CI -0.155–-0.009), p=0.026 | TRUE | 0.041 (95% CI 0.002–0.080), p=0.040 | 2.944913668 | Significant partial mediation |
| All Low vs Mixed | Adjusted | HIV.DURATION | 1.202 (95% CI 0.777–1.861), p=0.408 | 2.499 (SE 0.072), p=0.000 | 1.564 (95% CI 1.239–1.974), p=0.000 | 0.433 (95% CI 0.220–0.852), p=0.015 | 1.117 (log-odds), OR=3.056 (95% CI 1.702–5.487) | Boot CI [0.524, 1.766] | 606.1% (>100%) | 0.104 (95% CI 0.047–0.168), p=0.000 | TRUE | -0.079 (95% CI -0.151–-0.012), p=0.021 | TRUE | 0.025 (95% CI -0.016–0.065), p=0.232 | 4.116272339 | Significant partial mediation |
| Mixed vs All High | Unadjusted | HIV.DURATION | 1.847 (95% CI 1.247–2.736), p=0.002 | 0.668 (SE 0.074), p=0.000 | 1.476 (95% CI 1.208–1.804), p=0.000 | 1.407 (95% CI 0.921–2.148), p=0.114 | 0.260 (log-odds), OR=1.297 (95% CI 1.122–1.500) | Boot CI [0.129, 0.413] | 42.4% | 0.035 (95% CI 0.017–0.056), p=0.000 | TRUE | 0.046 (95% CI -0.013–0.104), p=0.125 | FALSE | 0.082 (95% CI 0.023–0.137), p=0.005 | 0.432733558 | Significant complete mediation |
| Mixed vs All High | Adjusted | HIV.DURATION | 1.967 (95% CI 1.321–2.931), p=0.001 | 0.684 (SE 0.074), p=0.000 | 1.461 (95% CI 1.192–1.790), p=0.000 | 1.489 (95% CI 0.969–2.289), p=0.069 | 0.259 (log-odds), OR=1.296 (95% CI 1.116–1.504) | Boot CI [0.115, 0.424] | 38.3% | 0.034 (95% CI 0.015–0.057), p=0.000 | TRUE | 0.053 (95% CI -0.004–0.114), p=0.082 | FALSE | 0.087 (95% CI 0.033–0.145), p=0.001 | 0.394292574 | Significant complete mediation |
| All Low vs All High | Unadjusted | HIV.DURATION | 2.663 (95% CI 1.720–4.122), p=0.000 | 3.214 (SE 0.094), p=0.000 | 1.652 (95% CI 1.332–2.048), p=0.000 | 0.560 (95% CI 0.255–1.231), p=0.149 | 1.613 (log-odds), OR=5.016 (95% CI 2.497–10.077) | Boot CI [0.903, 2.423] | 164.6% (>100%) | 0.194 (95% CI 0.107–0.281), p=0.000 | TRUE | -0.072 (95% CI -0.178–0.030), p=0.169 | FALSE | 0.122 (95% CI 0.065–0.176), p=0.000 | 1.584279181 | Significant complete mediation |
| All Low vs All High | Adjusted | HIV.DURATION | 2.607 (95% CI 1.672–4.063), p=0.000 | 3.191 (SE 0.092), p=0.000 | 1.565 (95% CI 1.256–1.950), p=0.000 | 0.661 (95% CI 0.298–1.468), p=0.309 | 1.429 (log-odds), OR=4.175 (95% CI 2.059–8.464) | Boot CI [0.672, 2.285] | 149.2% (>100%) | 0.167 (95% CI 0.078–0.259), p=0.000 | TRUE | -0.049 (95% CI -0.152–0.045), p=0.321 | FALSE | 0.118 (95% CI 0.060–0.171), p=0.000 | 1.418944533 | Significant complete mediation |
| Other AIDS-Defining Events | | | | | | | | | | | | | | | | |
| All Low vs Mixed | Unadjusted | AGE, HIV.DURATION, CENTER | 1.561 (95% CI 0.927–2.630), p=0.094 | 2.546 (SE 0.071), p=0.000 | 1.565 (95% CI 1.195–2.051), p=0.001 | 0.530 (95% CI 0.234–1.199), p=0.127 | 1.141 (log-odds), OR=3.130 (95% CI 1.570–6.241) | Boot CI [0.519, 1.823] | 256.1% (>100%) | 0.076 (95% CI 0.031–0.138), p=0.001 | TRUE | -0.044 (95% CI -0.121–0.018), p=0.182 | FALSE | 0.032 (95% CI -0.000–0.065), p=0.050 | 2.371145752 | Significant complete mediation |
| All Low vs Mixed | Adjusted | AGE, HIV.DURATION, CENTER | 1.418 (95% CI 0.828–2.430), p=0.203 | 2.519 (SE 0.071), p=0.000 | 1.552 (95% CI 1.161–2.075), p=0.003 | 0.514 (95% CI 0.220–1.201), p=0.124 | 1.107 (log-odds), OR=3.026 (95% CI 1.452–6.306) | Boot CI [0.478, 1.905] | 316.9% (>100%) | 0.071 (95% CI 0.025–0.127), p=0.001 | TRUE | -0.044 (95% CI -0.112–0.017), p=0.180 | FALSE | 0.027 (95% CI -0.005–0.059), p=0.096 | 2.604388855 | Significant complete mediation |
| Mixed vs All High | Unadjusted | AGE, HIV.DURATION, CENTER | 1.745 (95% CI 1.095–2.779), p=0.019 | 0.668 (SE 0.074), p=0.000 | 1.405 (95% CI 1.113–1.773), p=0.004 | 1.368 (95% CI 0.829–2.257), p=0.220 | 0.227 (log-odds), OR=1.255 (95% CI 1.066–1.477) | Boot CI [0.081, 0.391] | 40.8% | 0.022 (95% CI 0.008–0.038), p=0.004 | TRUE | 0.030 (95% CI -0.017–0.076), p=0.184 | FALSE | 0.052 (95% CI 0.004–0.101), p=0.034 | 0.420512890 | Significant complete mediation |
| Mixed vs All High | Adjusted | AGE, HIV.DURATION, CENTER | 1.757 (95% CI 1.083–2.852), p=0.022 | 0.689 (SE 0.075), p=0.000 | 1.403 (95% CI 1.105–1.782), p=0.006 | 1.365 (95% CI 0.811–2.297), p=0.241 | 0.233 (log-odds), OR=1.263 (95% CI 1.063–1.500) | Boot CI [0.081, 0.422] | 41.4% | 0.022 (95% CI 0.007–0.039), p=0.002 | TRUE | 0.029 (95% CI -0.017–0.076), p=0.215 | FALSE | 0.051 (95% CI 0.005–0.099), p=0.032 | 0.428695715 | Significant complete mediation |
| All Low vs All High | Unadjusted | AGE, HIV.DURATION, CENTER | 2.724 (95% CI 1.599–4.642), p=0.000 | 3.214 (SE 0.094), p=0.000 | 2.018 (95% CI 1.530–2.660), p=0.000 | 0.306 (95% CI 0.112–0.835), p=0.021 | 2.256 (log-odds), OR=9.542 (95% CI 3.890–23.408) | Boot CI [1.507, 3.177] | 225.1% (>100%) | 0.195 (95% CI 0.114–0.285), p=0.000 | TRUE | -0.110 (95% CI -0.211–-0.014), p=0.021 | TRUE | 0.085 (95% CI 0.039–0.132), p=0.000 | 2.287179022 | Significant partial mediation |
| All Low vs All High | Adjusted | AGE, HIV.DURATION, CENTER | 2.675 (95% CI 1.541–4.644), p=0.000 | 3.198 (SE 0.092), p=0.000 | 1.962 (95% CI 1.474–2.611), p=0.000 | 0.336 (95% CI 0.119–0.946), p=0.039 | 2.155 (log-odds), OR=8.628 (95% CI 3.431–21.698) | Boot CI [1.400, 3.193] | 219.0% (>100%) | 0.177 (95% CI 0.102–0.265), p=0.000 | TRUE | -0.095 (95% CI -0.192–-0.010), p=0.026 | TRUE | 0.082 (95% CI 0.037–0.130), p=0.000 | 2.162393038 | Significant partial mediation |
| Cardiovascular Disorders | | | | | | | | | | | | | | | | |
| All Low vs Mixed | Unadjusted | AGE, HIV.DURATION, CMV.IgG.IUpermL, SMOKING.CURRENT | 1.298 (95% CI 0.978–1.724), p=0.071 | 2.546 (SE 0.071), p=0.000 | 1.107 (95% CI 0.967–1.266), p=0.141 | 1.006 (95% CI 0.647–1.563), p=0.980 | 0.258 (log-odds), OR=1.294 (95% CI 0.918–1.825) | Boot CI [-0.073, 0.618] | 98.7% | 0.054 (95% CI -0.016–0.126), p=0.137 | FALSE | 0.001 (95% CI -0.095–0.102), p=0.968 | FALSE | 0.056 (95% CI -0.004–0.118), p=0.068 | 0.978394557 | No significant mediation (ACME not significant) |
| All Low vs Mixed | Adjusted | AGE, HIV.DURATION, CMV.IgG.IUpermL, SMOKING.CURRENT | 0.872 (95% CI 0.620–1.226), p=0.431 | 2.502 (SE 0.077), p=0.000 | 1.041 (95% CI 0.884–1.225), p=0.632 | 0.791 (95% CI 0.469–1.335), p=0.380 | 0.100 (log-odds), OR=1.105 (95% CI 0.735–1.661) | Boot CI [-0.307, 0.518] | -72.6% | 0.018 (95% CI -0.056–0.090), p=0.626 | FALSE | -0.042 (95% CI -0.136–0.057), p=0.393 | FALSE | -0.024 (95% CI -0.086–0.037), p=0.449 | -0.738961429 | No significant mediation (ACME not significant) |
| Mixed vs All High | Unadjusted | AGE, HIV.DURATION, CMV.IgG.IUpermL, SMOKING.CURRENT | 1.128 (95% CI 0.829–1.537), p=0.443 | 0.668 (SE 0.074), p=0.000 | 1.025 (95% CI 0.880–1.193), p=0.753 | 1.110 (95% CI 0.802–1.537), p=0.528 | 0.016 (log-odds), OR=1.016 (95% CI 0.918–1.125) | Boot CI [-0.083, 0.113] | 13.5% | 0.004 (95% CI -0.019–0.026), p=0.752 | FALSE | 0.024 (95% CI -0.049–0.093), p=0.517 | FALSE | 0.027 (95% CI -0.044–0.094), p=0.439 | 0.134563431 | No significant mediation (ACME not significant) |
| Mixed vs All High | Adjusted | AGE, HIV.DURATION, CMV.IgG.IUpermL, SMOKING.CURRENT | 0.981 (95% CI 0.677–1.423), p=0.920 | 0.672 (SE 0.080), p=0.000 | 0.957 (95% CI 0.796–1.151), p=0.642 | 1.012 (95% CI 0.683–1.499), p=0.954 | -0.029 (log-odds), OR=0.971 (95% CI 0.858–1.099) | Boot CI [-0.161, 0.092] | 154.7% (>100%) | -0.006 (95% CI -0.029–0.017), p=0.641 | FALSE | 0.002 (95% CI -0.072–0.076), p=0.955 | FALSE | -0.003 (95% CI -0.073–0.066), p=0.934 | 1.654533809 | No significant mediation (ACME not significant) |
| All Low vs All High | Unadjusted | AGE, HIV.DURATION, CMV.IgG.IUpermL, SMOKING.CURRENT | 1.465 (95% CI 1.061–2.023), p=0.020 | 3.214 (SE 0.094), p=0.000 | 1.266 (95% CI 1.100–1.457), p=0.001 | 0.694 (95% CI 0.402–1.199), p=0.190 | 0.758 (log-odds), OR=2.133 (95% CI 1.355–3.357) | Boot CI [0.328, 1.248] | 198.3% (>100%) | 0.160 (95% CI 0.064–0.245), p=0.002 | TRUE | -0.076 (95% CI -0.178–0.038), p=0.196 | FALSE | 0.083 (95% CI 0.013–0.153), p=0.017 | 1.913691317 | Significant complete mediation |
| All Low vs All High | Adjusted | AGE, HIV.DURATION, CMV.IgG.IUpermL, SMOKING.CURRENT | 0.860 (95% CI 0.578–1.279), p=0.456 | 3.161 (SE 0.101), p=0.000 | 1.113 (95% CI 0.941–1.316), p=0.210 | 0.614 (95% CI 0.318–1.187), p=0.147 | 0.339 (log-odds), OR=1.403 (95% CI 0.826–2.383) | Boot CI [-0.194, 0.902] | -224.3% | 0.058 (95% CI -0.036–0.149), p=0.221 | FALSE | -0.083 (95% CI -0.189–0.030), p=0.155 | FALSE | -0.026 (95% CI -0.093–0.046), p=0.456 | -2.268564212 | No significant mediation (ACME not significant) |
| Angina Pectoris | | | | | | | | | | | | | | | | |
| All Low vs Mixed | Unadjusted | AGE, HIV.DURATION, CENTER, SMOKING.CURRENT | 1.795 (95% CI 0.718–4.490), p=0.211 | 2.546 (SE 0.071), p=0.000 | 1.334 (95% CI 0.852–2.089), p=0.208 | 0.886 (95% CI 0.217–3.616), p=0.866 | 0.734 (log-odds), OR=2.083 (95% CI 0.665–6.528) | Boot CI [-0.270, 1.859] | 125.4% (>100%) | 0.016 (95% CI -0.005–0.053), p=0.141 | FALSE | -0.003 (95% CI -0.048–0.030), p=0.880 | FALSE | 0.013 (95% CI -0.005–0.032), p=0.143 | 1.206290875 | No significant mediation (ACME not significant) |
| All Low vs Mixed | Adjusted | AGE, HIV.DURATION, CENTER, SMOKING.CURRENT | 2.040 (95% CI 0.711–5.848), p=0.185 | 2.541 (SE 0.075), p=0.000 | 1.206 (95% CI 0.743–1.959), p=0.448 | 1.303 (95% CI 0.277–6.129), p=0.737 | 0.477 (log-odds), OR=1.611 (95% CI 0.470–5.524) | Boot CI [-0.622, 1.645] | 66.9% | 0.009 (95% CI -0.011–0.035), p=0.372 | FALSE | 0.005 (95% CI -0.030–0.035), p=0.685 | FALSE | 0.015 (95% CI -0.004–0.034), p=0.137 | 0.640773936 | No significant mediation (ACME not significant) |
| Mixed vs All High | Unadjusted | AGE, HIV.DURATION, CENTER, SMOKING.CURRENT | 1.631 (95% CI 0.755–3.522), p=0.213 | 0.668 (SE 0.074), p=0.000 | 1.006 (95% CI 0.688–1.470), p=0.977 | 1.625 (95% CI 0.722–3.656), p=0.241 | 0.004 (log-odds), OR=1.004 (95% CI 0.779–1.294) | Boot CI [-0.210, 0.220] | 0.8% | 0.000 (95% CI -0.008–0.008), p=0.986 | FALSE | 0.017 (95% CI -0.012–0.050), p=0.245 | FALSE | 0.017 (95% CI -0.011–0.049), p=0.240 | 0.007706143 | No significant mediation (ACME not significant) |
| Mixed vs All High | Adjusted | AGE, HIV.DURATION, CENTER, SMOKING.CURRENT | 1.119 (95% CI 0.472–2.654), p=0.798 | 0.667 (SE 0.079), p=0.000 | 0.932 (95% CI 0.607–1.432), p=0.747 | 1.185 (95% CI 0.469–2.997), p=0.720 | -0.047 (log-odds), OR=0.954 (95% CI 0.716–1.271) | Boot CI [-0.348, 0.211] | -41.7% | -0.001 (95% CI -0.011–0.006), p=0.679 | FALSE | 0.005 (95% CI -0.026–0.037), p=0.787 | FALSE | 0.004 (95% CI -0.025–0.031), p=0.857 | -0.384292365 | No significant mediation (ACME not significant) |
| All Low vs All High | Unadjusted | AGE, HIV.DURATION, CENTER, SMOKING.CURRENT | 2.928 (95% CI 1.154–7.432), p=0.024 | 3.214 (SE 0.094), p=0.000 | 1.247 (95% CI 0.839–1.853), p=0.275 | 1.450 (95% CI 0.303–6.929), p=0.641 | 0.709 (log-odds), OR=2.032 (95% CI 0.568–7.267) | Boot CI [-0.268, 1.792] | 66.0% | 0.020 (95% CI -0.009–0.065), p=0.156 | FALSE | 0.011 (95% CI -0.052–0.054), p=0.657 | FALSE | 0.030 (95% CI 0.004–0.059), p=0.023 | 0.650621857 | No significant mediation (ACME not significant) |
| All Low vs All High | Adjusted | AGE, HIV.DURATION, CENTER, SMOKING.CURRENT | 2.441 (95% CI 0.798–7.471), p=0.118 | 3.207 (SE 0.097), p=0.000 | 0.979 (95% CI 0.612–1.565), p=0.928 | 2.613 (95% CI 0.409–16.698), p=0.310 | -0.069 (log-odds), OR=0.933 (95% CI 0.207–4.205) | Boot CI [-1.479, 1.306] | -7.8% | -0.002 (95% CI -0.039–0.027), p=0.894 | FALSE | 0.020 (95% CI -0.023–0.071), p=0.322 | FALSE | 0.019 (95% CI -0.005–0.043), p=0.116 | -0.081739701 | No significant mediation (ACME not significant) |
| Osteoporosis | | | | | | | | | | | | | | | | |
| All Low vs Mixed | Unadjusted | AGE, HIV.DURATION, CMV.IgG.IUpermL | 1.618 (95% CI 0.739–3.545), p=0.229 | 2.546 (SE 0.071), p=0.000 | 1.087 (95% CI 0.749–1.577), p=0.660 | 1.312 (95% CI 0.390–4.408), p=0.661 | 0.213 (log-odds), OR=1.237 (95% CI 0.480–3.191) | Boot CI [-0.686, 1.216] | 44.2% | 0.006 (95% CI -0.023–0.039), p=0.661 | FALSE | 0.008 (95% CI -0.035–0.050), p=0.684 | FALSE | 0.014 (95% CI -0.008–0.038), p=0.211 | 0.440087722 | No significant mediation (ACME not significant) |
| All Low vs Mixed | Adjusted | AGE, HIV.DURATION, CMV.IgG.IUpermL | 1.249 (95% CI 0.554–2.814), p=0.592 | 2.481 (SE 0.074), p=0.000 | 1.046 (95% CI 0.698–1.568), p=0.827 | 1.126 (95% CI 0.329–3.850), p=0.850 | 0.112 (log-odds), OR=1.118 (95% CI 0.410–3.050) | Boot CI [-0.917, 1.290] | 50.3% | 0.003 (95% CI -0.029–0.041), p=0.848 | FALSE | 0.003 (95% CI -0.040–0.048), p=0.852 | FALSE | 0.007 (95% CI -0.018–0.032), p=0.574 | 0.484684920 | No significant mediation (ACME not significant) |
| Mixed vs All High | Unadjusted | AGE, HIV.DURATION, CMV.IgG.IUpermL | 1.566 (95% CI 0.785–3.122), p=0.203 | 0.668 (SE 0.074), p=0.000 | 0.876 (95% CI 0.622–1.233), p=0.447 | 1.704 (95% CI 0.829–3.503), p=0.147 | -0.089 (log-odds), OR=0.915 (95% CI 0.728–1.151) | Boot CI [-0.308, 0.098] | Suppression / inconsistent mediation (opposite signs) | -0.004 (95% CI -0.014–0.004), p=0.376 | FALSE | 0.024 (95% CI -0.010–0.058), p=0.178 | FALSE | 0.020 (95% CI -0.013–0.052), p=0.237 | -0.203392522 | No significant mediation (ACME not significant) |
| Mixed vs All High | Adjusted | AGE, HIV.DURATION, CMV.IgG.IUpermL | 1.539 (95% CI 0.736–3.219), p=0.252 | 0.704 (SE 0.076), p=0.000 | 0.827 (95% CI 0.565–1.211), p=0.329 | 1.789 (95% CI 0.809–3.955), p=0.151 | -0.134 (log-odds), OR=0.875 (95% CI 0.668–1.146) | Boot CI [-0.400, 0.123] | -31.0% | -0.006 (95% CI -0.019–0.005), p=0.315 | FALSE | 0.025 (95% CI -0.013–0.064), p=0.215 | FALSE | 0.019 (95% CI -0.016–0.054), p=0.298 | -0.304841253 | No significant mediation (ACME not significant) |
| All Low vs All High | Unadjusted | AGE, HIV.DURATION, CMV.IgG.IUpermL | 2.533 (95% CI 1.133–5.666), p=0.024 | 3.214 (SE 0.094), p=0.000 | 1.032 (95% CI 0.740–1.438), p=0.853 | 2.291 (95% CI 0.603–8.698), p=0.223 | 0.101 (log-odds), OR=1.106 (95% CI 0.380–3.216) | Boot CI [-0.858, 1.178] | 10.8% | 0.004 (95% CI -0.042–0.045), p=0.812 | FALSE | 0.030 (95% CI -0.025–0.092), p=0.269 | FALSE | 0.034 (95% CI 0.003–0.066), p=0.031 | 0.112562613 | No significant mediation (ACME not significant) |
| All Low vs All High | Adjusted | AGE, HIV.DURATION, CMV.IgG.IUpermL | 1.867 (95% CI 0.796–4.379), p=0.151 | 3.169 (SE 0.095), p=0.000 | 0.879 (95% CI 0.612–1.262), p=0.484 | 2.825 (95% CI 0.665–11.997), p=0.159 | -0.410 (log-odds), OR=0.664 (95% CI 0.211–2.089) | Boot CI [-1.599, 0.851] | -65.6% | -0.015 (95% CI -0.071–0.028), p=0.508 | FALSE | 0.036 (95% CI -0.021–0.109), p=0.216 | FALSE | 0.021 (95% CI -0.009–0.053), p=0.163 | -0.698316455 | No significant mediation (ACME not significant) |
| Post-cART Malignancies | | | | | | | | | | | | | | | | |
| All Low vs Mixed | Unadjusted | AGE, HIV.DURATION, CMV.IgG.IUpermL | 1.608 (95% CI 0.926–2.793), p=0.092 | 2.546 (SE 0.071), p=0.000 | 1.324 (95% CI 1.007–1.743), p=0.045 | 0.807 (95% CI 0.343–1.899), p=0.624 | 0.715 (log-odds), OR=2.045 (95% CI 1.016–4.117) | Boot CI [-0.046, 1.565] | 150.6% (>100%) | 0.041 (95% CI -0.006–0.100), p=0.080 | FALSE | -0.013 (95% CI -0.078–0.041), p=0.632 | FALSE | 0.029 (95% CI -0.003–0.061), p=0.072 | 1.441924363 | No significant mediation (ACME not significant) |
| All Low vs Mixed | Adjusted | AGE, HIV.DURATION, CMV.IgG.IUpermL | 1.251 (95% CI 0.703–2.227), p=0.446 | 2.481 (SE 0.074), p=0.000 | 1.335 (95% CI 0.987–1.804), p=0.061 | 0.650 (95% CI 0.269–1.570), p=0.338 | 0.716 (log-odds), OR=2.046 (95% CI 0.968–4.327) | Boot CI [-0.182, 1.714] | 319.6% (>100%) | 0.042 (95% CI -0.008–0.105), p=0.106 | FALSE | -0.026 (95% CI -0.091–0.030), p=0.373 | FALSE | 0.016 (95% CI -0.018–0.053), p=0.362 | 2.559774739 | No significant mediation (ACME not significant) |
| Mixed vs All High | Unadjusted | AGE, HIV.DURATION, CMV.IgG.IUpermL | 0.877 (95% CI 0.494–1.558), p=0.655 | 0.668 (SE 0.074), p=0.000 | 1.174 (95% CI 0.889–1.550), p=0.259 | 0.784 (95% CI 0.426–1.443), p=0.434 | 0.107 (log-odds), OR=1.113 (95% CI 0.923–1.342) | Boot CI [-0.066, 0.297] | Suppression / inconsistent mediation (opposite signs) | 0.007 (95% CI -0.004–0.019), p=0.211 | FALSE | -0.016 (95% CI -0.060–0.024), p=0.433 | FALSE | -0.009 (95% CI -0.048–0.029), p=0.620 | -0.785446532 | No significant mediation (ACME not significant) |
| Mixed vs All High | Adjusted | AGE, HIV.DURATION, CMV.IgG.IUpermL | 0.888 (95% CI 0.492–1.603), p=0.694 | 0.704 (SE 0.076), p=0.000 | 1.173 (95% CI 0.875–1.573), p=0.285 | 0.782 (95% CI 0.413–1.482), p=0.452 | 0.112 (log-odds), OR=1.119 (95% CI 0.909–1.377) | Boot CI [-0.085, 0.323] | -94.9% | 0.008 (95% CI -0.006–0.021), p=0.246 | FALSE | -0.017 (95% CI -0.062–0.030), p=0.464 | FALSE | -0.009 (95% CI -0.049–0.033), p=0.664 | -0.848535993 | No significant mediation (ACME not significant) |
| All Low vs All High | Unadjusted | AGE, HIV.DURATION, CMV.IgG.IUpermL | 1.411 (95% CI 0.744–2.675), p=0.292 | 3.214 (SE 0.094), p=0.000 | 1.107 (95% CI 0.843–1.455), p=0.464 | 1.018 (95% CI 0.346–2.998), p=0.974 | 0.327 (log-odds), OR=1.387 (95% CI 0.577–3.335) | Boot CI [-0.626, 1.434] | 95.1% | 0.018 (95% CI -0.036–0.083), p=0.512 | FALSE | 0.001 (95% CI -0.077–0.072), p=0.980 | FALSE | 0.019 (95% CI -0.016–0.056), p=0.285 | 0.947226329 | No significant mediation (ACME not significant) |
| All Low vs All High | Adjusted | AGE, HIV.DURATION, CMV.IgG.IUpermL | 1.138 (95% CI 0.591–2.191), p=0.699 | 3.169 (SE 0.095), p=0.000 | 0.958 (95% CI 0.724–1.269), p=0.766 | 1.302 (95% CI 0.431–3.928), p=0.640 | -0.135 (log-odds), OR=0.874 (95% CI 0.359–2.126) | Boot CI [-1.110, 0.968] | -104.2% | -0.008 (95% CI -0.067–0.057), p=0.821 | FALSE | 0.015 (95% CI -0.063–0.089), p=0.704 | FALSE | 0.007 (95% CI -0.029–0.047), p=0.690 | -1.048882799 | No significant mediation (ACME not significant) |
| *^1^*Manual IE uses a*b on the log-odds scale (a from lm(M~X), b from glm(Y~X+M)). Manual % mediated = (a*b)/c on log-odds scale only when signs agree and TE != 0. mediate() ACME is on the probability scale (Imai et al). Interpret logistic-scale and probability-scale proportions with caution. | | | | | | | | | | | | | | | | |

**Supplementary Table 14 | Consensus outliers in multi-omics clustering for the 2000HIV study.**

The table reports consensus outliers identified in the discovery (n=1,230, pre-outlier removal) and validation (n=189, pre-outlier removal) cohorts during the final MoCluster run using the MOVICS pipeline. Columns include sample ID, endotype (MIXED, ALL LOW, ALL HIGH), multi-omics factor scores (PC1, PC2, PC3), and outlier status flags for three methods: univariate interquartile range (IQR, 1.5×IQR rule on z-scores of principal components), robust Mahalanobis distance (FAC MAH, Minimum Covariance Determinant, α=0.75, 99% chi-squared, df=2), and silhouette scores (Silhouette, Euclidean distance, threshold<0.1). The consensus outlier column indicates samples flagged by at least two of the three methods. Consensus outliers were identified using an iterative approach combining three methods applied to the integrated latent space (PC1, PC2, PC3): (1) Univariate IQR: Outliers detected by applying the 1.5×IQR rule to z-scores of each principal component; (2) Mahalanobis Distance: Outliers identified using Minimum Covariance Determinant (MCD)-based Mahalanobis distance on integrated factor scores; (3) Silhouette Scores: Outliers flagged based on low silhouette scores (threshold<0.1), indicating poor fit to any endotype. A sample was classified as a consensus outlier if flagged by at least two methods. This process was repeated iteratively, removing identified outliers and recalculating factor scores and outlier metrics until no further consensus outliers were detected. MoCluster hyperparameters were optimized using Silhouette, Davies-Bouldin, and Calinski-Harabasz scores. Consensus outliers (25 in discovery, 14 in validation) were removed, yielding final discovery (n=1,002) and validation (n=189) cohorts for downstream analyses. Results were visualized via violin plots, silhouette plots, 3D scatterplots, and 2D projections (Supplementary Figures 3–5). IQR, interquartile range; FAC MAH, factor Mahalanobis distance.

| ID | Endotype | PC1 | PC2 | PC3 | outlier IQR | outlier FAC MAH | Outlier Silhouette | Consensus Outlier | Cohort |
| --- | --- | --- | --- | --- | --- | --- | --- | --- | --- |
| EMC144 | ALL LOW | -0.089283094 | -0.034725949 | -0.004067685 | TRUE | TRUE | FALSE | TRUE | Discovery |
| EMC191 | MIXED | -0.01842796 | 0.027292471 | -0.01730427 | TRUE | FALSE | TRUE | TRUE | Discovery |
| EMC288 | MIXED | -0.011315058 | 0.01911296 | -0.024893915 | TRUE | TRUE | FALSE | TRUE | Discovery |
| EMC360 | ALL HIGH | -0.040444619 | 0.003274657 | -0.142475856 | TRUE | TRUE | TRUE | TRUE | Discovery |
| EMC391 | ALL HIGH | 0.040631169 | -0.003386072 | -0.142597411 | TRUE | TRUE | TRUE | TRUE | Discovery |
| EMC404 | ALL HIGH | -0.040500873 | -0.035581307 | -0.141132161 | TRUE | TRUE | TRUE | TRUE | Discovery |
| EMC408 | MIXED | 0.03534893 | -0.000709952 | -0.020247029 | TRUE | TRUE | TRUE | TRUE | Discovery |
| EMC411 | ALL HIGH | 0.029298384 | 0.002766404 | -0.142680288 | TRUE | TRUE | TRUE | TRUE | Discovery |
| EMC415 | ALL HIGH | 0.039139948 | -0.003259404 | -0.14252507 | TRUE | TRUE | TRUE | TRUE | Discovery |
| EMC438 | ALL HIGH | 0.104918781 | -0.015951473 | 0.003464899 | TRUE | TRUE | FALSE | TRUE | Discovery |
| EMC479 | ALL HIGH | 0.105038751 | -0.025337583 | -0.010016459 | TRUE | TRUE | FALSE | TRUE | Discovery |
| EMC572 | ALL LOW | -0.08502119 | -0.039294457 | 0.004378231 | TRUE | TRUE | FALSE | TRUE | Discovery |
| OLV323 | ALL HIGH | -0.052815762 | -0.020391954 | -0.141607131 | TRUE | TRUE | TRUE | TRUE | Discovery |
| OLV480 | MIXED | 0.008881904 | 0.013936189 | -0.023819538 | TRUE | TRUE | FALSE | TRUE | Discovery |
| OLV531 | ALL LOW | -0.05543357 | -0.048900302 | 0.008504549 | TRUE | TRUE | FALSE | TRUE | Discovery |
| OLV656 | ALL HIGH | 0.026636899 | -0.03590823 | -0.141378232 | TRUE | TRUE | TRUE | TRUE | Discovery |
| OLV657 | ALL HIGH | 0.009837145 | 0.001295679 | -0.14259874 | TRUE | TRUE | TRUE | TRUE | Discovery |
| OLV658 | ALL HIGH | 0.034455089 | 0.001621256 | -0.142690716 | TRUE | TRUE | TRUE | TRUE | Discovery |
| OLV659 | ALL HIGH | 0.011884745 | -0.023767546 | -0.141764601 | TRUE | TRUE | TRUE | TRUE | Discovery |
| OLV671 | ALL HIGH | -0.039741006 | -0.007160238 | -0.142142634 | TRUE | TRUE | TRUE | TRUE | Discovery |
| OLV672 | MIXED | 0.021581362 | 0.05187287 | 0.009564214 | TRUE | TRUE | FALSE | TRUE | Discovery |
| RAD146 | ALL HIGH | -0.006848229 | 0.025917416 | -0.143357634 | TRUE | TRUE | TRUE | TRUE | Discovery |
| RAD189 | ALL HIGH | -0.018187608 | 0.00953969 | -0.142784495 | TRUE | TRUE | TRUE | TRUE | Discovery |
| RAD236 | ALL LOW | -0.004183566 | 0.009292943 | 0.021346646 | TRUE | FALSE | TRUE | TRUE | Discovery |
| RAD257 | MIXED | -0.003297985 | -0.00577506 | -0.02512678 | TRUE | TRUE | FALSE | TRUE | Discovery |
| ETZ103 | ALL LOW | 0.034769247 | -0.039881985 | -0.107286973 | TRUE | TRUE | FALSE | TRUE | Validation |
| ETZ108 | ALL HIGH | -0.213816031 | 0.00656004 | -0.045597467 | TRUE | TRUE | FALSE | TRUE | Validation |
| ETZ165 | ALL LOW | 0.045235568 | 0.086648111 | -0.024037979 | TRUE | TRUE | FALSE | TRUE | Validation |
| ETZ173 | ALL HIGH | -0.032344146 | 0.041914412 | 0.076835866 | FALSE | TRUE | TRUE | TRUE | Validation |
| ETZ174 | ALL HIGH | -0.062971794 | 0.043706234 | 0.085777405 | FALSE | TRUE | TRUE | TRUE | Validation |
| ETZ195 | ALL LOW | -0.007296068 | -0.037771458 | -0.096597384 | FALSE | TRUE | TRUE | TRUE | Validation |
| ETZ206 | ALL HIGH | -0.033799862 | 0.017226517 | -0.11348422 | TRUE | FALSE | TRUE | TRUE | Validation |
| ETZ241 | ALL HIGH | -0.187047535 | 0.046697813 | -0.012425992 | TRUE | TRUE | FALSE | TRUE | Validation |
| ETZ242 | ALL HIGH | -0.070407643 | 0.020865799 | -0.129379503 | TRUE | TRUE | FALSE | TRUE | Validation |
| ETZ245 | ALL LOW | 0.18110209 | -0.022206315 | -0.086757307 | TRUE | TRUE | FALSE | TRUE | Validation |
| ETZ252 | ALL HIGH | 0.013163037 | 0.387858017 | 0.088833662 | TRUE | TRUE | TRUE | TRUE | Validation |
| ETZ265 | ALL HIGH | 0.006314095 | 0.382133563 | 0.11282501 | TRUE | TRUE | TRUE | TRUE | Validation |
| ETZ327 | ALL HIGH | -0.185184736 | -0.008995037 | -0.055635687 | TRUE | TRUE | FALSE | TRUE | Validation |
| ETZ332 | ALL HIGH | -0.218173586 | 0.004194615 | -0.024556423 | TRUE | TRUE | FALSE | TRUE | Validation |
